# Direct and indirect responses of the Arabidopsis transcriptome to an induced increase in trehalose 6-phosphate

**DOI:** 10.1101/2023.09.18.555309

**Authors:** Omri Avidan, Marina C. M. Martins, Regina Feil, Marc Lohse, Federico M. Giorgi, Armin Schlereth, John E. Lunn, Mark Stitt

**Author notes:** ^a^in Press - Consultoria e Comunicação Científica, São Paulo, Brazil (Marina C. M. Martins). ^b^targenomix GmbH, Am Mühlenberg 11, 14476 Potsdam, Germany (Marc Lohse). ^c^Department of Pharmacy and Biotechnology, <u>University of Bologna</u>, Italy (Federico M. Giorgi).

## Abstract

Trehalose 6-phosphate (Tre6P) is an essential signal metabolite that reports and regulates the level of sucrose, linking growth and development to the metabolic status. We hypothesized that Tre6P plays a role in mediating the regulation of gene expression by sucrose. To test this, we performed transcriptomic profiling on Arabidopsis plants that expressed a bacterial trehalose-6-phosphate synthase (TPS) under the control of an ethanol-inducible promoter. Induction led to a 4-fold rise in Tre6P levels, a concomitant decrease in sucrose, and significant changes of over 13,000 transcripts and two-fold or larger changes of over 5000 transcripts. Comparison with nine published responses to sugar availability allowed some of these changes to be linked to the rise in Tre6P, while others were probably due to lower sucrose or other indirect effects. Changes linked to Tre6P included repression of photosynthesis and induction of many growth-related processes including ribosome biogenesis. About 500 starvation-related genes are known to be induced by SUCROSE-NON-FERMENTING-1-RELATED KINASE 1 (SnRK1). They were largely repressed by Tre6P in a manner consistent with Tre6P acting to inhibit SnRK1. SnRK1 also represses many genes that are involved in biosynthesis and growth. These responded to Tre6P in a more complex manner, pointing to Tre6P also interacting with further C-signaling pathways. In addition, elevated Tre6P modified expression of genes encoding regulatory subunits of the SnRK1 complex and TPS class II and FLZ proteins that are thought to modulate SnRK1 function, and genes involved in the circadian clock and in TOR, light, abscisic acid and other hormone signaling.

**One sentence summary:** An induced increase in trehalose 6-phosphate levels has direct effects on gene expression via inhibition of SUCROSE-NON-FERMENTING-1-RELATED KINASE 1 and interactions with light, circadian clock and phytohormone signaling, and widespread indirect effects on gene expression from reciprocal changes in sucrose levels.

## Introduction

Trehalose 6-phosphate (Tre6P) is an essential signal metabolite in plant metabolism and development (Fichtner and Lunn, 2021). Tre6P is synthesized by TREHALOSE-6-PHOSPHATE SYNTHASE (TPS) from UDP-glucose (UDPGlc) and glucose 6-phosphate (Glc6P) and dephosphorylated by TREHALOSE-6-PHOSPHATE PHOSPHATASE (TPP) to produce trehalose (Cabib and Leloir, 1958). Whereas trehalose is a major carbon (C) storage metabolite and osmolyte in bacteria and fungi, these roles have been largely taken over by sucrose in vascular plants, allowing neo-functionalization of the trehalose biosynthesis pathway (Goddijn and Van Dun, 1999; Lunn et al., 2014; Figueroa and Lunn, 2016). In addition to canonical catalytically-active TPS proteins like AtTPS1 in Arabidopsis (*Arabidopsis thaliana*), plants possess a family of catalytically inactive TPS proteins (termed class II TPS proteins) (Leyman et al., 2001; Harthill et al., 2006; Lunn, 2007; Ramon et al., 2009; Vandesteene et al., 2010; Lunn et al., 2014; Delorge et al., 2015) and a family of TPP proteins (Leyman et al., 2001; Vandesteene et al., 2012; Kretzschmar et al., 2015). The class II TPS and TPP families have expanded independently in different plant lineages and have seven and ten members, respectively, in Arabidopsis. The embryo lethal phenotype of the Arabidopsis *tps1* mutant (Eastmond et al 2002), analysis of the phenotypes of Arabidopsis lines overexpressing bacterial TPS and bacterial TPPs (Schleupmann et al., 2003) and measurements of Tre6P levels in plants (Lunn et al., 2006) established that Tre6P acts as a signal and is essential for plant growth. Tre6P profoundly influences metabolism and growth (Lunn et al., 2014; Figueroa and Lunn, 2016; Fichtner and Lunn, 2021) and also has a major impact on development, including the floral transition (Wahl et al., 2013) inflorescence structure (Satoh-Nagasawa et al., 2006; Claeys et al., 2019, Klein et al., 2022) and lateral shoot branching (Fichtner et al., 2017, 2021). Furthermore, modification of Tre6P levels can influence crop yield (Nuccio et al., 2015; Griffiths et al., 2016; Paul et al., 2020). However, open questions remain concerning the mechanism(s) by which Tre6P acts. In the following, we investigate Tre6P signaling by analyzing the immediate response of the transcriptome to an induced increase in Tre6P.

It has been proposed that Tre6P acts as a signal for sucrose availability (Lunn et al., 2006). Tre6P levels are strongly correlated with sucrose levels in Arabidopsis, including when sucrose levels change during and after perturbing diel cycles (Lunn et al., 2006; Carillo et al., 2013; Martins et al., 2013; Sulpice et al., 2014; Wahl et al., 2013; Figueroa et al., 2016; Annunziata et al., 2017), after sugar feeding (Lunn et al., 2006; Nunes et al., 2013; Yadav et al., 2014) and in response to genetic interventions that alter sucrose levels by changing C partitioning (Lunn et al., 2006) or sucrose transport (dos Anjos et al., 2019). Correlations between sucrose and Tre6P levels have also been observed in other species (Debast et al., 2011; Martinez-Barajas et al., 2011; Henry et al., 2014; Zhang et al., 2015). The response of Tre6P to sucrose appears to be specific; responses to changes in the levels of other sugars, such as glucose and fructose, or the nitrogen (N) supply are explained by concomitant changes in sucrose levels (Yadav et al., 2014). The mechanism linking Tre6P to sucrose levels is unknown, but depends on *de novo* protein synthesis (Yadav et al., 2014) and features of the AtTPS1 protein (Fichtner et al., 2021).

Genetic interventions that alter TPS or TPP expression result in reciprocal changes of Tre6P and sucrose. This is seen both in plants that constitutively overexpress bacterial TPS or TPP (Gomez et al., 2011; Yadav et al., 2014; Nuccio et al., 2015) and in the short-term response to induced expression of bacterial TPS (Martins et al., 2013; Figueroa et al., 2016). Reciprocal changes are also seen after vascular tissue-specific overexpression of TPS (Fichtner et al., 2020; 2021). These observations imply that Tre6P inhibits sucrose production and/or stimulates sucrose consumption. Together with the positive correlation between sucrose and Tre6P levels in wild-type plants, these observations led to proposal of the sucrose:Tre6P nexus hypothesis, according to which Tre6P has a dual function as a signal of sucrose levels and a negative feedback regulator of sucrose levels (Lunn et al., 2014; Yadav et al., 2014; Figueroa and Lunn, 2016). This hypothesis has been expanded (Fichtner and Lunn, 2021) to propose that the sensitivity and response range of the relationship between sucrose and Tre6P differ between tissues and depend on developmental stage (Martinez-Barajas et al., 2011; Carillo et al., 2013; Dai et al., 2013; Wahl et al., 2013; Henry et al., 2015; Czedik-Eysenberg et al., 2016) and environmental conditions (Carillo et al., 2013).

The primary role of Tre6P may vary between source tissues that are producing and exporting sucrose, and sink tissues that are utilizing sucrose for growth or long-term storage. In source leaves in the light, induced increases in Tre6P post-translationally stimulate phospho*enol*pyruvate carboxylase (PEPC) and nitrate reductase (NR), leading to increased synthesis of organic acids and amino acids and decreased synthesis of sucrose (Figueroa et al., 2016). At night, increased Tre6P restricts starch mobilization, resulting in decreased synthesis of sucrose (Martins et al., 2013; dos Anjos et al., 2019). It was recently shown that Tre6P also restricts starch mobilization in the light (Ishihara et al., 2021). In Arabidopsis leaves, *AtTPS1* is mainly expressed in the phloem parenchyma and the companion cell-sieve element complex (Fichtner et al., 2020). Presumably, Tre6P that is formed in the phloem parenchyma moves symplastically, via plasmodesmata, into mesophyll cells to modify metabolism and adjust the supply of sucrose and amino acids for export to growing organs, whilst Tre6P that is produced in the companion cells may provide a signal linked to the movement of sucrose in the phloem. Tre6P is known to modulate long-distance signaling pathways that control developmental transitions which set up a future demand for sucrose, including the CONSTANS (CO)/FLOWERING TIME (FT) photoperiod pathway that regulates flowering (Wahl et al., 2013) and signals that regulate shoot branching (Fichtner et al., 2017; 2021).

In sink tissues, Tre6P regulates sucrose mobilization by inhibiting sucrose synthase (Fedosejevs et al., 2018) and modifying expression of sucrolytic enzymes and SWEETs (Bledsoe et al., 2017; Oszvald et al., 2018; Fichtneer et al., 2021). Tre6P stimulates central metabolism in axillary buds in a similar manner to leaves (Fichtner et al., 2017; 2021). Constitutive overexpression of a bacterial TPS in seedlings led to increased transcript abundance of genes involved in cell wall biosynthesis (Paul et al., 2010). Tre6P promotes storage product accumulation in Arabidopsis seeds by stabilizing the transcription factor WRINKLED1 (Zhai et al., 2018), and in pea seeds by inducing the auxin biosynthesis gene *TRYPTOPHAN AMINOTRANSFERASE RELATED2* (Meitzel et al., 2021). There is growing evidence that Tre6P influences further signaling pathways including light- and auxin-signaling (Paul et al., 2010), the miR156-SQUAMOSA-PROMOTER-BINDING LIKE (SPL) pathway (Wahl et al., 2013; Ponnu et al., 2020; Zhang et al., 2022) and abscisic acid (ABA) signaling (Avonce et al., 2006; Ramon et al., 2007; Gomez et al., 2010; Tian et al., 2019).

Tre6P can act by inhibiting SnRK1 (Zhang et al., 2009; Paul et al., 2020; Baena-Gonzales and Lunn, 2020). SnRK1 is the plant homolog of yeast SUCROSE-NON-FERMENTING1 (SNF1) and mammalian AMP-ACTIVATED PROTEIN KINASE (AMPK) that play a key role in low-energy signaling (Jossier et al., 2009; Hulsmans et al., 2016; Paul et al., 2020; Crepin and Rolland, 2019b; Baena-González and Lunn, 2020). The first line of evidence is that Tre6P, along with several other sugar phosphates like Glc6P, inhibits *in vitro* SnRK1 activity (Zhang et al., 2009; Debast et al., 2011; Delatte et al., 2011; Nunes et al., 2013; Coello and Martínez-Barajas, 2014; Emanuelle et al., 2015). This *in vitro* inhibition requires an unidentified protein and has only been observed in extracts from sink tissues (Zhang et al., 2009; Emmanuelle et al., 2015). Subsequent *in vitro* studies showed that Tre6P interferes with binding of SnRK1-activating kinases (SnAK1/GRIK1 and SnAK2/GRIK2) to the α subunit of SnRK1, leading to inhibition of SnRK1 activity (Glab et al., 2017; Zhai et al., 2018; Hwang et al., 2019). However, SnAK1 and 2 are expressed ubiquitously whereas *in vitro* inhibition of SnRK1 by Tre6P is only observed in extracts from young growing tissues, indicating that this may be a separate mechanism to that reported by Zhang et al. (2009). The second line of evidence is that changes of Tre6P levels *in vivo* often correlate negatively with the abundance of C-starvation induced transcripts like *DARK-INDUCED6* (*DIN6*), *DIN1*, *BRANCHED CHAIN AMINO-ACID TRANSAMINASE2* (*BCAT2*) and *EXPANSIN10* (*EXP10*) (Zhang et al., 2009; Paul et al., 2010; Henry et al., 2015; Bledsoe et al., 2017: Peixoto et al., 2021) that are induced by transient SnRK1 overexpression in mesophyll protoplasts (Baena-Gonzalez et al., 2007). Further, many of these genes are repressed in transgenic plants that constituvely overexpress a bacterial TPS (Zhang et al., 2009; Oszvald et al., 2018). Application of permeable Tre6P analogs to Arabidopsis also led to changes in transcript abundance for a subset of SnRK1 downstream target genes (Griffiths et al., 2016). Two recent findings provide further evidence for interactions between Tre6P and SnRK1. One is that TPS class II proteins physically interact with SnRK1 and can inhibit its activity (van Leene et al., 2022). The other is that whilst the strong positive correlation between Tre6P and sucrose was retained in Arabidopsis lines with increased or reduced SnRK1 expression, the response of Tre6P to sucrose was damped as SnRK1 expression was increased (Peixoto et al., 2021).

However, changes of downstream targets of SnRK1 are seen in plant material that is dominated by mature source leaves, whereas *in vitro* inhibition of SnRK1 activity by Tre6P is observed in sink tissues but not source leaves (see Baena-Gonzalez and Lunn, 2020, for discussion). Furthermore, diel changes in SnRK1 activity, based on phosphorylation of an *in-vivo* reporter protein, did not always correlate with Tre6P levels and also varied independently of the abundance of SnRK1 downstream target transcripts, indicating that the latter may not always be a faithful readout of SnRK1 activity (Avidan et al., 2023). Thus, whilst it has been established that Tre6P can act, at least in part, via inhibition of SnRK1 activity, open questions remain both concerning the molecular mechanism and when this interaction plays a major role in Tre6P and SnRK1 signaling.

Another open question concerns the relationship between Tre6P signaling and TARGET OF RAPAMYCIN (TOR), which operates in an antagonistic manner to SnRK1 to promote metabolism and growth (Lastdräger et al., 2014; Baena-González and Hanson, 2017; Wu et al., 2019, Meng et al., 2022). The TOR and SnRK1 complexes are expressed in the vasculature (Moreau et al., 2012, Williams et al., 2014), overlapping with *AtTPS1* expression (Fichtner et al., 2020), underlining the possibility of regulatory interactions between TOR, SnRK1 and Tre6P. The metabolic profiles of Arabidopsis *lst8* (*lethal with sec thirteen 8*) mutants that lack a regulatory subunit of the TOR complex exhibit similarities to those of plants with constitutively elevated Tre6P (Moreau et al., 2012). Furthermore, TOR signaling interacts closely with other signaling pathways that control metabolism and growth including brassinosteroid (BR) and ABA signaling (Zhang et al., 2016; Wang et al., 2018a; Wu et al., 2019, Meng et al., 2022) and there is long-standing evidence for an interaction between the latter and Tre6P signaling (Avonce et al., 2004; Gomez et al., 2010; Debast et al., 2011; Tian et al., 2019; Belda-Palazón et al., 2020, 2022). Further open questions include whether there are interactions between Tre6P and other sugar-signaling pathways like the transcription factor (TF) BASIC LEUCINE ZIPPER 11 (bZIP11; Hanson et al., 2008; Ma et al., 2011) that is translationally regulated by sucrose (Weise et al., 2004; Rahmani et al., 2009) and acts downstream of SnRK1 in the transcriptional orchestration of starvation responses.

Except for a recent study that investigated the impact of an induced increase of Tre6P on a small subset of SnRK1 downstream target transcripts (Peixoto et al., 2021), previous studies of the impact of Tre6P on transcript abundance have either been correlative or used transgenic lines with constitutive overexpression of TPS or TPP. As already discussed, these genetic interventions generate reciprocal changes in the levels of Tre6P and sucrose and other sugars, and result in large-scale changes in metabolism, growth and development. This makes it very difficult to distinguish between direct transcriptional responses to Tre6P, indirect responses due to a decline in the levels of sucrose and other sugars, and pleiotropic effects. In the following, we investigate the short-term response of transcript abundance to an induced increase in Tre6P. Based on the role of Tre6P as a sucrose signal, we reasoned that the direct response to Tre6P should resemble the response observed when sucrose increases in wild-type plants. By comparing the response to transiently elevated Tre6P with responses in a set of published treatments in which the levels of sucrose and other sugars were increased, we identify components of C-signaling that may be directly downstream of Tre6P, and demarcate them from indirect responses. The results revealed a complex pattern of direct and indirect responses of gene expression to elevated Tre6P, including inhibition of SnRK1 by Tre6P and links to other sugar-signaling-pathways and light, circadian clock and phytohormone signaling pathways.

## Results

### Response to an induced increase of Tre6P over an entire light or dark period

In an initial experiment, two iTPS lines (29.2, 31.3) and control alcR plants were sprayed with ethanol or water at dawn and harvested 12h later at the end of the day (ED treatment) or were sprayed at dusk and harvested 12h later at the end of the night (EN treatment) and profiled using Affymetrix ATH1 arrays (see Supplemental text for details, and Supplemental Table S1 for a list of all utilized transcriptome datasets). Differentially expressed genes (DEGs) were identified using a false discovery rate (FDR) <0.05. We termed the response to induction the ‘iTPS response’. Whilst line 29.1 showed a stronger response than line 31, their responses were strongly correlated (Supplemental Figure S1, Supplemental Table S2, R^2^ = 0.96 and 0.98 at ED and EN, respectively).

### Deconvolution of the iTPS response using the carbon response factor

Given that Tre6P is a sucrose signal, it might be expected that the iTPS response would qualitatively resemble the response to elevated sugar. However, initial inspection revealed that many genes that we know to be induced by sugars were repressed in the iTPS response and vice versa (see below for examples). A possible explanation is that an induced increase in Tre6P leads to a fall in sucrose (Martins et al., 2013; Figueroa et al., 2016) so many “Tre6P-responsive” genes might actually be responding to the fall in sucrose, rather than the rise in Tre6P per se.

To provide a global overview, we calculated a carbon response factor (CRF) for each gene, based on transcriptomics data from multiple published experiments (Supplemental Figure S2, Supplemental Dataset 2). Nine treatments were chosen that focused on short-term responses and minimized side-effects due to circadian- or light-signaling. They included addition of exogenous glucose or sucrose to starved seedlings in liquid culture under continuous low light (Osuna et al., 2007, Bläsing et al., 2005), comparison of the starchless *pgm* mutant with wild-type plants at four times in the diel cycle (Gibon et al., 2004; Bläsing et al., 2005; Usadel et al., 2008), and illumination of wild-type plants for four hours with ambient or low CO2 (Bläsing et al., 2005). We assigned transcripts to three CRF groups: group 1 (G1) contained transcripts that responded in iTPS in the same direction as to increased sugar, group 2 (G2) contained transcripts that responded in the opposite direction, and group 0 (G0) for transcripts that responded in iTPS but did not show a consistent response to sugar. A relaxed filter (CRF>log20.1) was used to maximize assignment to G1 or G2.

When the CRF values were compared with the overall iTPS response there was little similarity at ED and even less at EN (Supplemental Figure S3A-B). Transcripts assigned to G1 (48 and 24% of DEGs at ED and EN, respectively) showed a positive correlation between their CRF values and iTPS response consistent with them responding to elevated Tre6P. Transcripts assigned to G2 (23 and 57% of DEGs at ED and EN, respectively) showed a negative correlation between their CRF values and iTPS response consistent with them responding to the decrease in sugars (Supplemental Figure S3C-D, Supplemental Table S2, Supplemental Dataset S2). Some transcripts (29 and 19% of DEGs at ED and EN, respectively) were assigned to CRF group G0. This initial experiment showed that the 12-h post-induction response includes many indirect effects at ED and that these predominate at EN. We decided to focus on induction in the light and to harvest at earlier time points.

### Early response to an induced increase of Tre6P at the beginning of the day

Twenty-two-day-old iTPS29.2 and alcR plants were sprayed with 2% (v/v) ethanol or water at 0.5h after dawn and rosettes harvested 2h, 4h and 6h later. The bacterial TPS protein was detectable by immunoblotting at 2h and its abundance was higher at 4h and 6h after induction (Supplemental Figure S4A). Tre6P levels were significantly increased by 4h and continued to rise at 6h, reaching 4-fold higher levels than in controls (Figure 1A, Supplemental Dataset S3). Sucrose levels decreased significantly, falling to 70% of those in controls by 4h and remaining low at 6h (Figure 1B). Whilst Tre6P and sucrose levels were positively correlated in the three control treatments, they were negatively correlated from 2h onwards after ethanol-spraying of line 29.2 (Supplemental Figure S4B). The increase in Tre6P was accompanied by a decrease of Glc6P, Fru6P and PEP, and an increase of pyruvate, malate, fumarate, citrate, aconitate, 2-oxoglutarate and shikimate (Supplemental Figure S4C). These responses resembled those in previous studies on the iTPS lines (Martins et al., 2013; Figueroa et al., 2016; Avidan et al., 2022).

**Figure 1.**
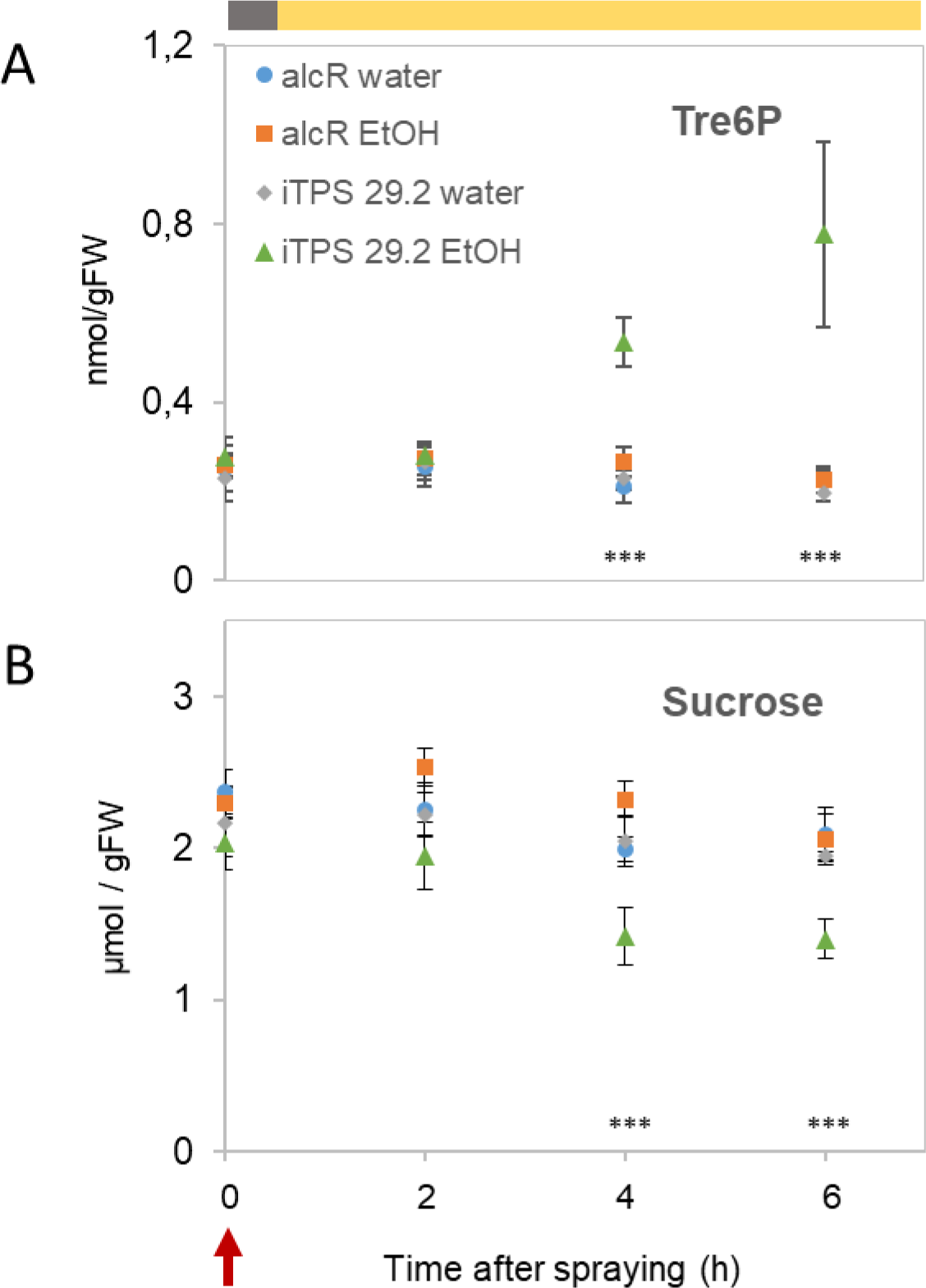
Changes of TreβPand sucrose after induction of bacterial TPS in the light. Arabidopsis İTPS29.2 and alcR plants were grown in long-day conditions (16 h light / 8 h dark, 160 µmol m^−2^ s^−1^ irradiance) for 22 days and were then sprayed at 0.5 h after dawn (Zeitgeber time 0.5, ZT0.5, red arrow) with either 2% (v/v) ethanol or water, and harvested 2, 4 and 6 h later (ZT2.5, 4.5, 6.5). The light period is indicated in the upper grey (dark) and orange (light) bar. Measurements were carried out on three controls (alcR and iTPS sprayed with water and alcR sprayed with *2%* (v/v) ethanol) and on İTPS29.2 sprayed with 2% (v/v) ethanol to induce bacterial TPS. (A) TreõP, (B) Sucrose. The results are plotted as mean ± S.D. (n = 4) (each replicate contained 4-5 whole rosettes). Significant differences (one-way ANOVA, Holm-Sidak) are denoted by asterisks (*P<0.05, **P<0.01, ***P<0.001) when the ethanol-induced iTPS samples were significantly different from all three controls in pairwise comparisons. Data for additional metabolites are shown in Supplemental Figure S4 and the original data are provided in Supplemental Dataset S3.

RNA sequencing (RNAseq) was performed on quadruplicate samples, harvested 4h and 6h after spraying. The ethanol-sprayed and water-sprayed quadruplicates were used to calculate the average log2 fold change (FC) in abundance and FDR (Benjamini and Hochberg, 1995). Elevated Tre6P led to extensive changes in transcript abundance; of 23.8K detected transcripts, over 13K passed an FDR<0.05 filter at both time points (about 55% of detected transcripts) and 5.6K and 5.4K passed a combined filter of FDR<0.05 and FC≥2 at 4h and 6h, respectively (about 23% of detected transcripts) (Figure 2A, Supplemental Dataset 4). There was high reproducibility between the 4-h and 6-h response (Figure 2A-B). The 4273 genes that passed the combined filter at both times represented 76 and 79% of the genes that passed the filter at 4h or 6h, respectively (Figure 2A, see also Supplemental Table S2). The 4-h and 6-h responses were highly correlated when the comparison was made using all detected transcripts, transcripts that passed the FDR-only filter, or transcripts that passed the combined filter (R^2^ = 0.71, 0.92 and 0.93, respectively, Figure 2B).

**Figure 2.**
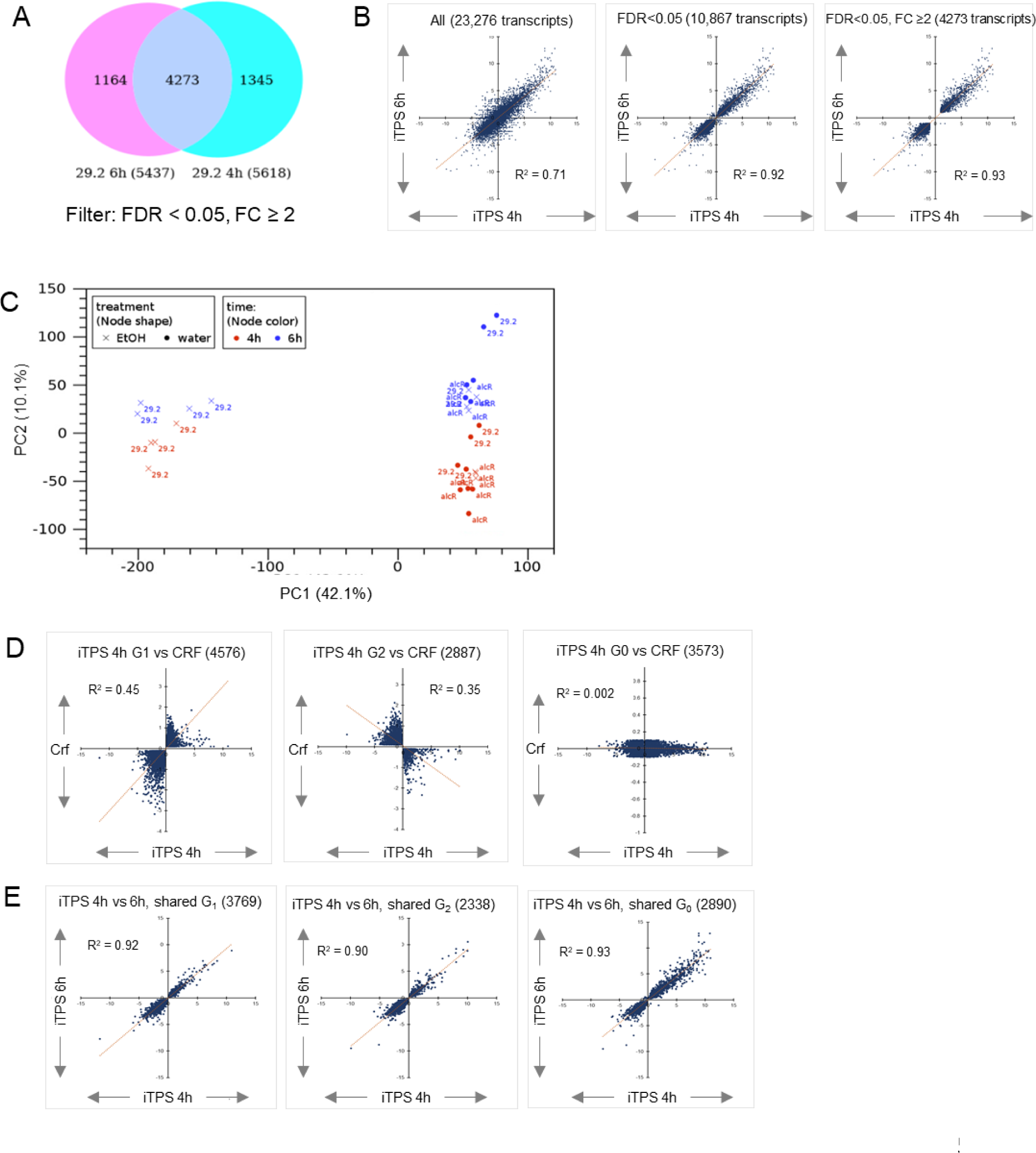
Changes of transcript abundance 4-h and 6-h after induction of bacterial TPS in the light. Arabidopsis İTPS29.2 and alcR plants we re grown and treated with ethanol or water as described in Figure 1 and RNAwas extracted for RNAseq analysis. Differentially expressed genes (DEGs) were identified by comparing the transcript abundance in the ethanol sprayed samples to their water sprayed control, using RPKM values (see Methods for full description). The VENN diagram compares DEGs that passed FDR <0.05 and fold change > 2 (FC2) filters in the 4-h and 6-h datasets; the numbers at the bottom represent total DEGs in each sample, while numbers located within circles represent shared and non­shared responses. Comparison of transcript responses at 4 h and 6 h for all 23.8K detected genes (left), the 10,867 transcripts that passed the FDR<0.05 filter (middle) and the 4273 transcripts that passed the FDR<0.05 and FC≥2 filter (right Principal component (PC) analysis performed on all detected transcripts. Genotype (29.2, alcR) is indicated in the figure, red and blue are the 4-h and 6­h treatments, circles and crosses are water- and ethanol-sprayed (see insert). Deconvoluted response to elevated Tre6P plotted against the carbon response factor (CRF) for iTPS 4-h (analogous plots for the response at 6 h are provided in Supplemental Figure S5B). The CRF summarizes the response of a given Arabidopsis gene transcript to a change in sugar levels across a set of treatments, with an increasingly positive sign indicating the average increase in abundance and an increasingly negative sign denoting the average decrease in abundance, whilst a value around zero indicatesthat transcript abundance does not respond to sugar status. Group 1 (G_a_) denotes transcripts where the iTPS response and CRF are qualitatively the same and, by inference, the iTPS response may be a direct response to elevated TreöP.Group 2 (G_2_) denotes transcripts where the iTPS response and CRF are qualitatively opposed and by inference the iTPS response is unlikely to be a direct response to elevated Tre6P. Group 0 (G_α_) denotes transcripts that respond in the iTPS response but cannot be assigned to G1 or G2 because they do not show a consistent response to changes in sugars (for details see Supplemental Figure S2 and Supplemental Dataset S2). Comparability of the response of transcript assigned to G_lř_ G_2_ and G_o_in the 4-h and 6-h data set.

To eliminate a possible effect of ethanol and off-target effects of alcR (Randall, 2021) we compared the response of TPS29.2 and alcR plants to ethanol induction. It should be noted that the alcR line contains an empty alcA promoter:OCT terminator cassette, providing a natural binding site (i.e. the alcA promoter) for the alcR protein to minimize off-target binding to endogenous genes. The effect was negligible; the number of shared DEGs between alcR and iTPS at 4h and 6h was 34 (22 in the same direction, compared to 5.6K total DEGs) and 12 (all 12 in the same direction, compared to 5.4K total DEGs), respectively (Supplemental Dataset S4). Genes with similar changes were omitted from further analyses.

Principal component (PC) analysis (Figure 2C) revealed a strong separation of ethanol-sprayed iTPS lines from the control treatments along the major PC1 axis (42% of total variance). PC2 (10% of variance) captured time-of-day-dependent separation of the 4-h and 6-h samples. Thus, an induced rise in Tre6P led to rapid and massive changes in transcript abundance.

### Dissection of iTPS 4 and 6-h response into CRF groups reveals a mix of direct and indirect responses even at early times after induction of TPS

Despite the earlier harvest times, the overall iTPS response showed no relationship to the CRF (Supplemental Figure S5A-B). To distinguish direct from indirect responses, we again assigned transcripts to CRF groups G1, G2 and G0 (see Supplemental Dataset S4 for CRF assignments). This meant that further analysis of the RNAseq dataset focused on the 22K genes present on the ATH1 array, for which multiple suitable treatments were available to allow CRF estimation. The largest subset of transcripts was assigned to G1, but substantial numbers were assigned to G2 and G0 (4576-4470, 2887-2969 and 3573-3653 at 4h and 6h, respectively; Supplemental Table S3). Positive correlations to CRF emerged for iTPS G1 and negative correlations for iTPS G2 (Figure 2D, Supplemental Figure S5B). There was strong agreement between the 4-h and 6-h response for each group (Figure 2E, R^2^ = 0.92, 0.90 and 0.93 for G1, G2 and G0, respectively). Thus, even at early time points, whilst many transcripts showed a response consistent with them responding to elevated Tre6P, the majority changed in a way indicating they were indirect responses. We compared the responses of transcripts assigned to G1, G2 and G0 in the microarray data from the ED and EN treatments with their responses in the RNAseq experiment (Supplemental Figure S5C, Supplemental Table S4). There was relatively good agreement, especially for transcripts that were repressed. Discrepancies may reflect genes that respond either transiently or slowly to an induced rise in Tre6P

Many iTPS-responsive transcripts were assigned to G0 (i.e., were apparently unresponsive to sugar availability). We investigated two technical explanations for this unexpected result. One is that the response to elevated sugar is context-dependent, i.e., transcript abundance responded in opposing ways in the nine treatments we used to calculate the CRF, so averaging leads to a very low CRF. Supplemental Figure S6A-B shows the iTPS response of the top 10 and 100 up-regulated and down-regulated transcripts individually for the nine treatments. Whilst some transcripts show marked and opposing responses, most are non-responsive across all nine treatments. A second technical explanation is that ATH1 arrays can underestimate responses for lowly-expressed genes, especially genes in large families where multiple members may cross-hybridize with the same probe set, masking any member that does respond. Inspection of the absolute abundance of transcripts assigned to G0 (Supplemental Figure S6C) revealed that whilst this may have interfered with assignment of a small proportion, the majority are expressed at levels comparable to those of transcripts assigned to G1 or G2. Biological explanations for why many iTPS-responding transcripts do not respond to changes in sugar availability will be examined in the Discussion.

### Elevated Tre6P represses genes involved in photosynthesis and gluconeogenesis, nucleotide biosynthesis and some sectors of specialized metabolism, and represses sucrose export

We next explored whether assignment of genes to G1, G2 and G0 allows areas of metabolism or cellular function to be identified that respond to elevated Tre6P, as opposed to indirect effects. We first used the PageMan tool (Usadel et al., 2006; https://mapman.gabipd.org/pageman) to visualize the response of genes assigned to different categories (BINs) in the MapMan ontology (Thimm et al., 2004; Schwache et al., 2019). All transcripts that failed to pass a filter of FDR<0.05 and log2 FC≥0.2 were set as zero before averaging the FC response in each category. We performed the analysis first at the highest level of the MapMan ontology. Different patterns emerged for G1, G2 and G0 (Figure 3). For example, in G1 many genes involved in photosynthesis and gluconeogenesis were repressed, whereas in G2 genes involved in fermentation, cell wall, lipid metabolism, N assimilation, S assimilation and specialized metabolism were repressed. This points to some sectors of metabolism being regulated by Tre6P and others by low sugars or other indirect consequences of suddenly elevating Tre6P.

**Figure 3.**
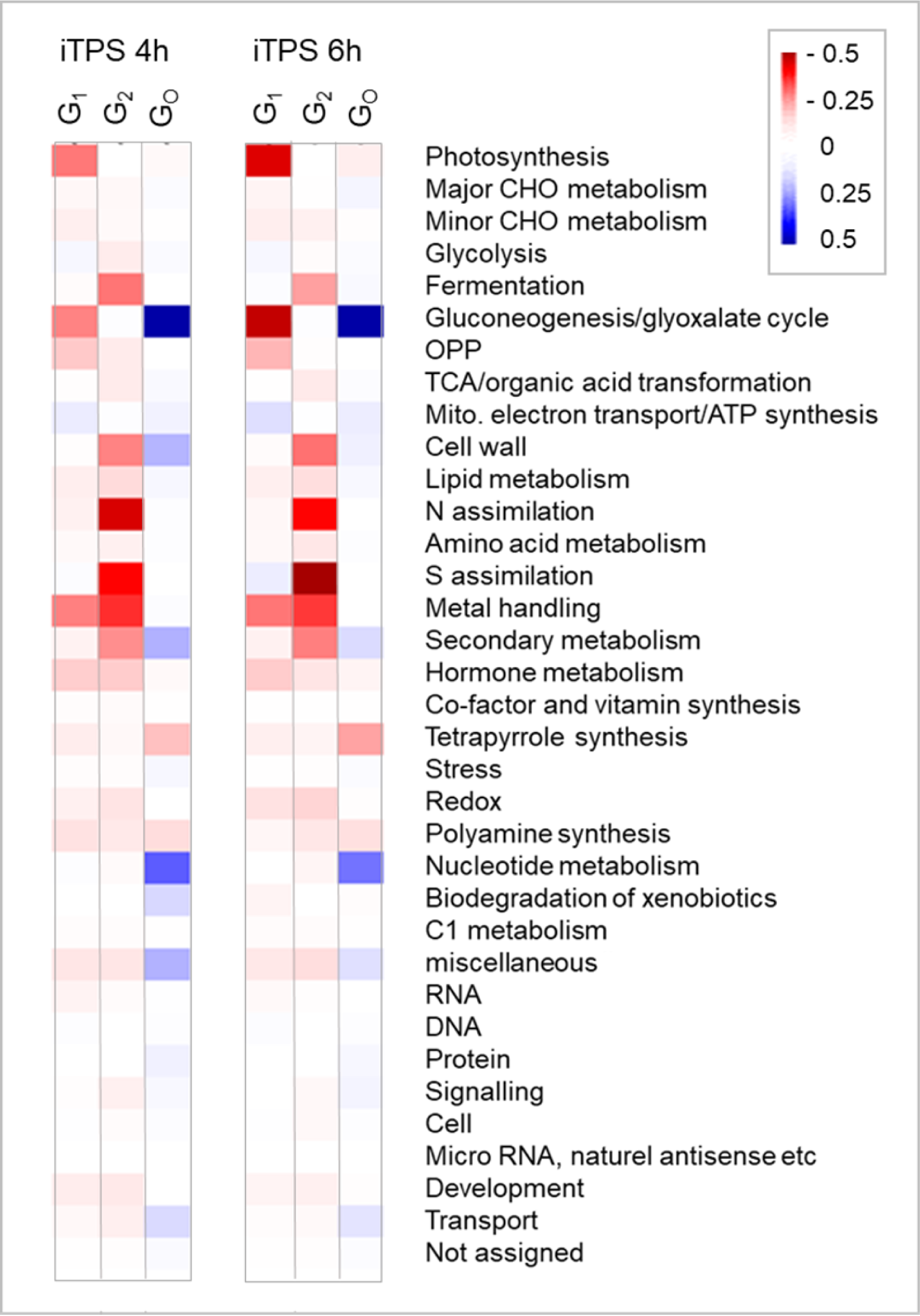
Enrichment analysis of responses at 4 h and 6 h after induction of TPS. The analysis was conducted using PageMan (Usadel et al., 2006) and MapMan software (version 3.6.0RC1; https://mapman.gabipd.org/; Ath_AGI_LOCUS_TAIR10_Aug2012). The analysis was performed separately for the sets of genes that were assigned to the CRF groups G_1_, G_2_ and G_o_ (see Supplemental Figure S2) and for the responses at 4 h and 6 h after spraying. The CFR groups are shown from left to right in the block in which the 4h and 6h response is displayed. The analyses used the log_2_FC values for all genes in a given category. These were filtered (FDR <0.05, FC ≥0.2; all values that did not pass the filter were set to zero) and all individual log_2_FC values (including the zero values) were then averaged for all genes in that category. The average log_2_FC values for each BIN (the upper category in the MapMan ontology) are displayed as a heat map (for scale see insert.) More detailed analyses in which several higher-level categories (photosynthesis, gluconeogenesis/glyoxylate, N metabolism, nucleotide metabolism, secondary metabolism, protein, cell wall) are broken down into sub-categories (subBINS) are provided in Supplemental Figure S7.

MapMan BINs group genes that participate in a given function, irrespective of whether they are involved in biosynthesis or catabolism. Some BINs also group different processes. We therefore inspected the responses in selected BINs at higher resolution (Supplemental Figure S7, see Supplemental text for details). Inspection of the response of genes assigned to CRF group G1 revealed that elevated Tre6P led to wide repression of genes for photosynthesis and related functions like tetrapyrrole, tocopherol and carotenoid biosynthesis and plastid ribosome biogenesis, and repressed gluconeogenesis and anthocyanin biosynthesis. Elevated Tre6P induced genes for nucleotide biosynthesis and (see Figure 4) cytosolic and mitochondrial ribosomal proteins and ribosome assembly factors. There were widespread changes in transcript abundance for genes involved in sucrose transport and metabolism, the control of floral induction, and the circadian clock, with many of these responses being assigned to CRF group G1 (Supplementary Figures S8-S10, see Supplemental text for details). In contrast, many genes that are involved in nitrate and ammonium assimilation, and large sectors of specialized metabolism including phenylpropanoid, flavonoid and glucosinolate biosynthesis were assigned to CRF group G2. Their responses were presumably indirect, due to the decline in sucrose or further changes in metabolism.

**Figure 4.**
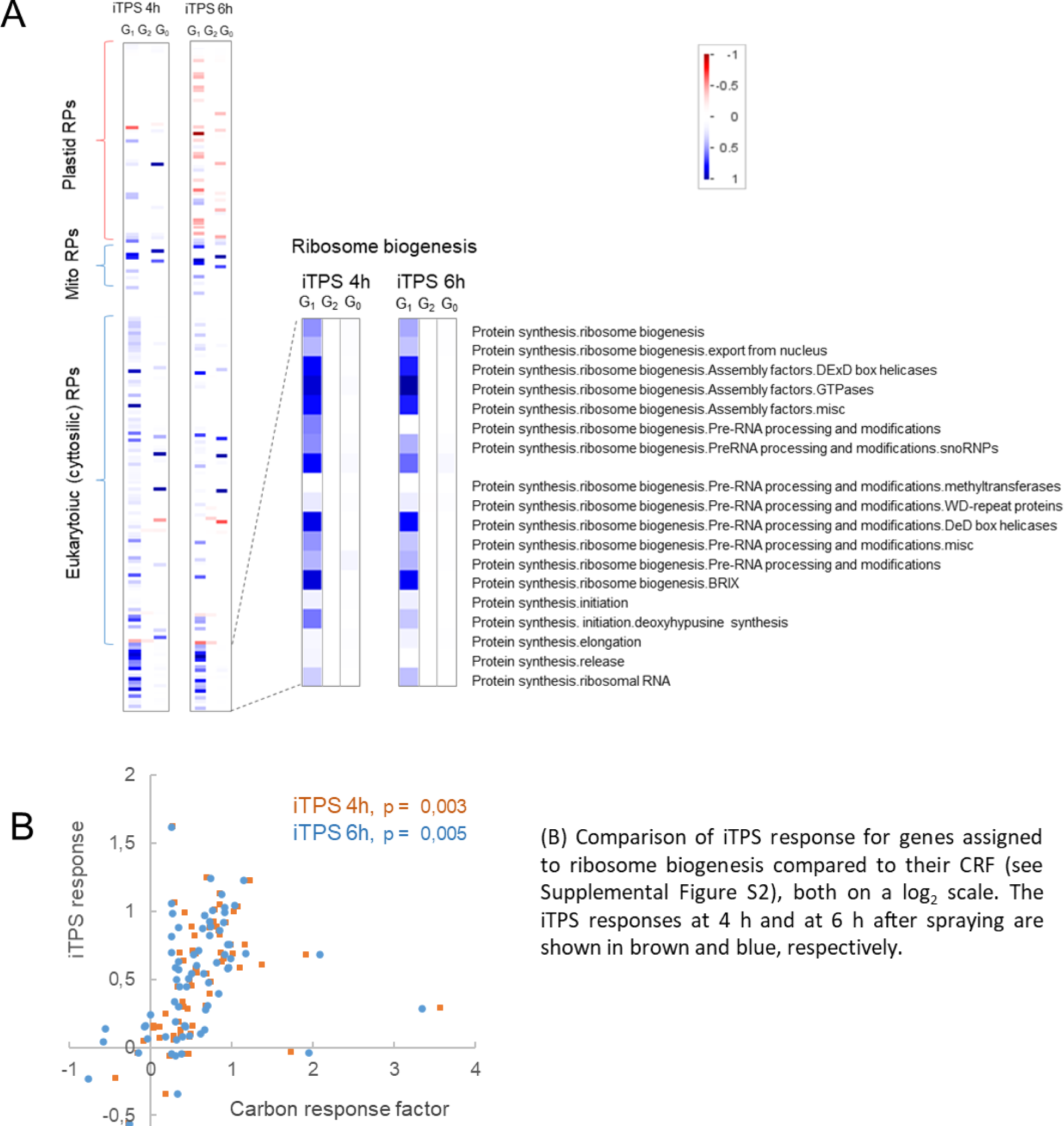
Induction of TPS leads to up-regulation of ribosome biogenesis at the transcript level. The plots show changes in transcript abundance after induction of TPS for genes assigned to ribosomal proteins, ribosome biogenesis and ribosomal RNA in the MapMan ontology. For each transcript, the response was calculated as the average change in ethanol-sprayed iTPS plants (induced) compared to water-sprayed iTPS plants (control) at 4 h or 6 h after spraying. A) Overview of genes representing ribosome proteins, ribosome biogenesis and ribosomal RNA with an expanded view of the subBINs associated with ribosome biogenesis and ribosomal RNA. The analysis was performed using PageMan (Usadel et *al.,* 2006); the shading indicates the average change in transcript abundance for genes assigned to a given subBIN. Genes that did not pass FDR<0.05 and log_2_FC >0.2 filters were excluded and their values set as zero. The display shows the average of all individual logƒC values. The heat map shows average changes calculated for each BIN or subBIN. A further, more detailed analysis in which further higher-level categories (photosynthesis, gluconeogenesis/glyoxylate, N metabolism, nucleotide metabolism, secondary metabolism, protein, cell wall) are broken down into sub-categories (subBINS) is provided in Supplemental Figure S6.

### Gene ontology analysis also highlights that Tre6P-signaling impacts on many sectors of metabolism, cellular growth and signaling pathways

As a complementary approach, we performed Gene Ontology (GO) analysis. This was done after filtering by FDR<0.05 and FC≥2 (Supplemental Figure S11) or FDR<0.05 (Supplemental Dataset S5). GO analysis confirmed many of the transcriptional responses in metabolism and cellular growth that were highlighted by PAGEMAN analysis, and highlighted further signaling responses. In the G1 gene set, the enriched upregulated categories included mitochondrial RNA modification and gene expression, as well as (in the FDR-only filtered set) protein synthesis, refolding and stability. Enriched downregulated categories included xyloglucan metabolic process, circadian clock entrainment, light responses, several hormone-related responses and (in the FDR-only filtered dataset) photosynthesis and pigments.

GO analysis confirmed that indirect effects impact other functions than those regulated by Tre6P. In the G2 gene set, the enriched upregulated GO categories were related mainly to stress, and the enriched downregulated categories included nitrate assimilation, nucleotide salvage, glucosinolate biosynthesis, flavonoid metabolism, cell wall loosening, pectin synthesis, cutin and wax biosynthesis, indole acetic acid biosynthesis, brassinosteroid biosynthesis, auxin transport, gibberellic acid and jasmonic acid signaling and (in the larger dataset) inositol biosynthesis, cell-wall biosynthesis, and phloem transport. In the CRF group G0 gene set, which respond to iTPS but not to an increase in sugar availability, enriched upregulated processes included processing and modification of RNA and DNA as well as protein post-translational modification, and downregulated processes included auxin-mediated signaling, and xylem, growth/morphogenesis, cell wall, nitrate, cytokinins, glucosinolate and defense related responses.

As summarized in Figure 5, PageMan and GO analysis of the iTPS response point to elevated Tre6P leading to repression of the photosynthetic machinery, repression of gluconeogenesis, complex changes in sucrose metabolism and transport, stimulation of nucleotide biosynthesis and stimulation of ribosome biogenesis in the cytosol and mitochondria. The increase of Tre6P is accompanied by a decrease of sucrose and changes in other central metabolites. This probably triggers many secondary changes including repression of genes involved in N and S metabolism and specialized metabolism, repression of genes involved in cell wall loosening and cell expansion, and widespread changes in signaling. It might be noted that when C availability rises in a wild-type plant, Tre6P and sugars will increase in parallel, leading to a cooperative repression of photosynthesis and induction of nutrient assimilation, and a broad induction of growth- and defense-related responses (see Discussion).

**Figure 5.**
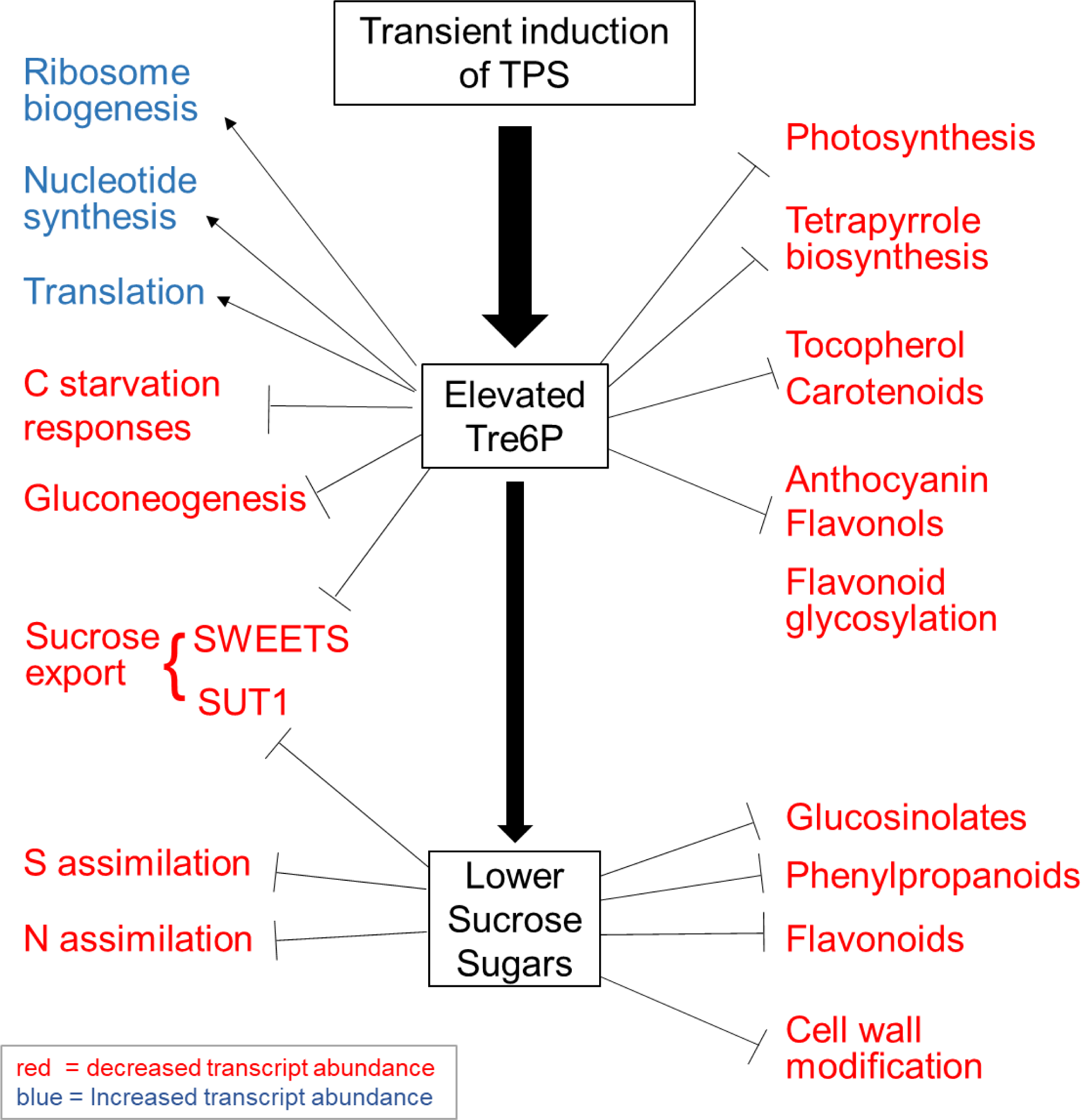
Summary of the metabolic and growth-related responses to an increase in Tre6P. Categories of genes that respond directly to elevated Tre6P were inferred from their assignment to CRF G_lř_ while categories of genes that respond to the concomitant fall in sucrose were inferred from their assignment to CRF G_2_. Up- and down-regulated gene categories are shown in blue and red, respectively, (see Supplemental Figure 52 and Supplemental Dataset 54)

### Comparison of the response to an induced increase in Tre6P with published response to constitutive overexpression of bacterial TPS

We next compared the short-term response to elevated Tre6P with a published response to constitutive overexpression of a bacterial TPS in seedlings (termed hereafter the oeTPS response) (Zhang et al., 2009; Paul et al., 2010). In this earlier study, transcript abundance was profiled using the CATMA array, which uses cDNA rather than oligonucleotide probes (Allemeersch et al., 2005). A total of 5273 genes were shortlisted as showing a response (FDR <0.05, FC≥2) and 4966 of these were found in the iTPS dataset. A scatter plot of the oeTPS and iTPS responses reveals poor agreement (Figure 6A; Supplemental Figure S12A). About half (2559) of the 4996 genes did not pass the FDR<0.05 filter in the iTPS dataset. Of the remainder, based on the shared iTPS response at 4h and 6h, 1596 were assigned to G1, 484 to G2 and 347 to G0 (Supplemental Table S5). Genes assigned to G1 showed very good agreement between the oeTPS and iTPS responses (R^2^ = 0.49, p = 7.02×10^-238^; Figure 6B, F Supplemental Figure S12B, Supplemental Table S5) with only 72 (4.5%) showing reciprocal responses. Plots of the individual G1 iTPS responses at 4h and 6h against the oeTPS response also showed good agreement (R^2^ = 0.47 and 0.51, p = 1.56×10^-255^ and 1.06×10^-297^, respectively, Supplemental Figure S12B). In contrast, there was little or no agreement between the oeTPS response and the iTPS response for genes that were assigned to G2 or GO (Figure 6C-D, Supplemental Figure S12B, Supplemental Table S5). We conclude about 30% of the responses to constitutive oeTPS were to elevated Tre6P whilst the rest were probably indirect.

**Figure 6.**
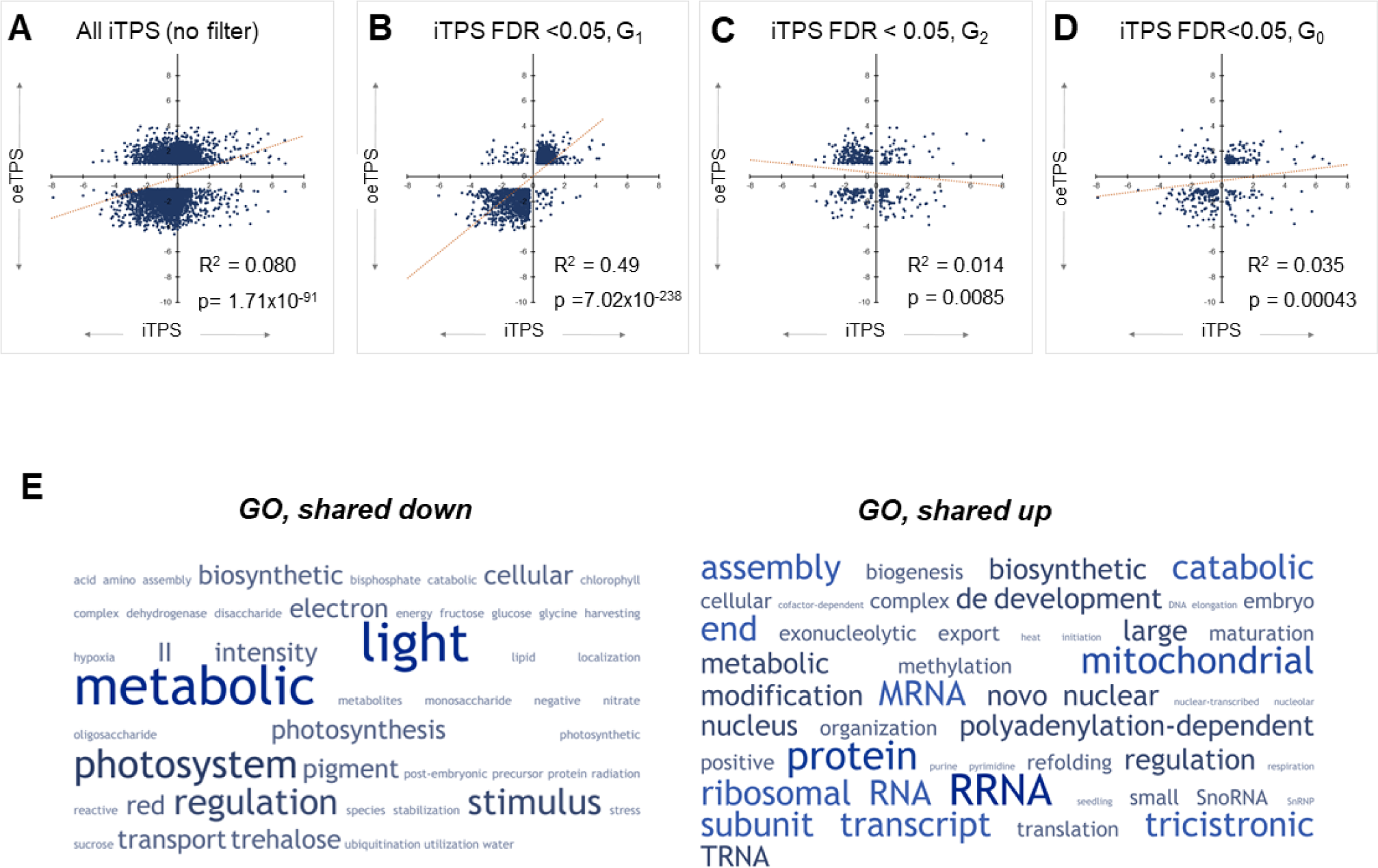
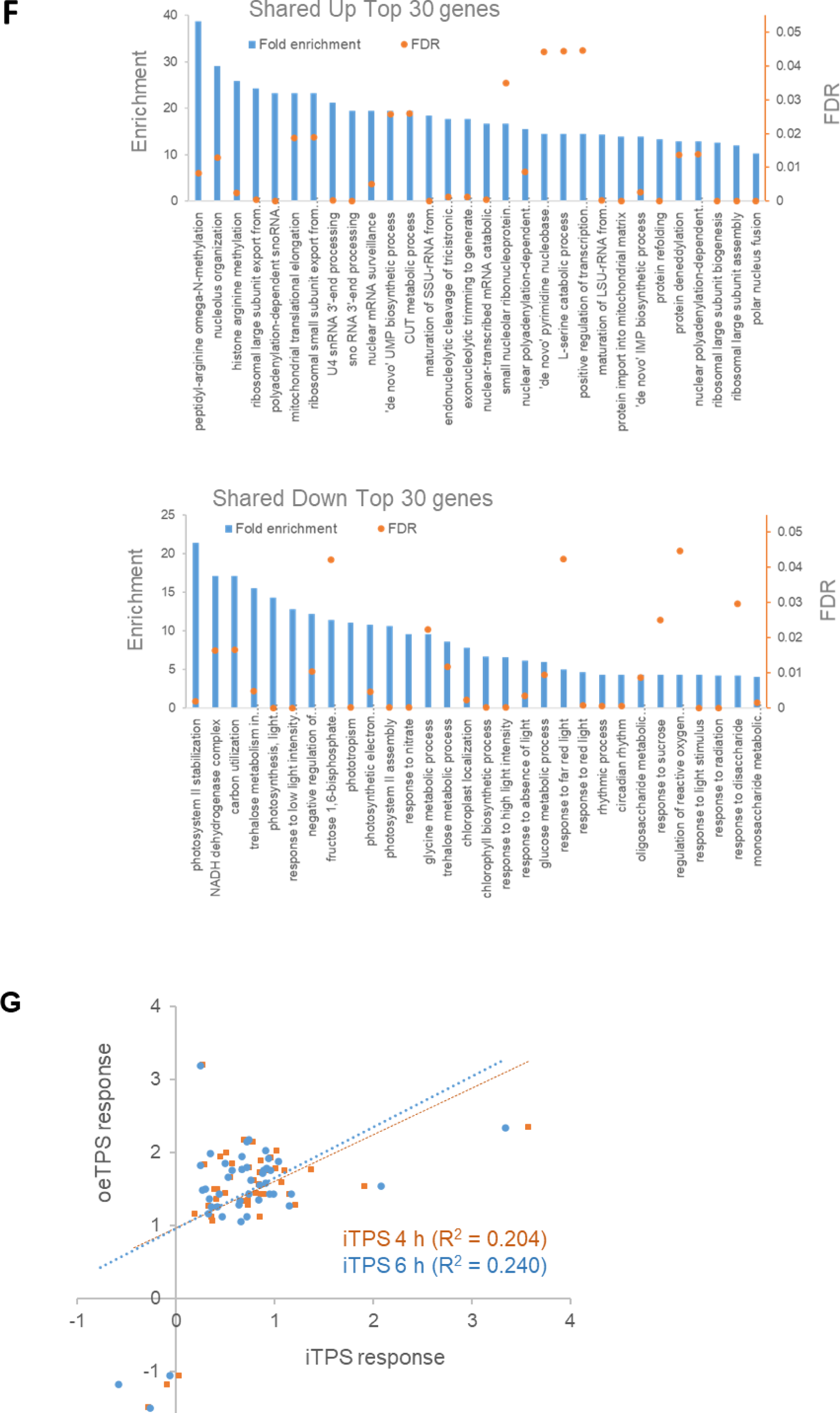
Comparison of the response to an induced increase in TreδP and constitutive overexpression of a bacterial TPS. The response of transcript abundance to constitutive overexpression of a bacterial TPS (oeTPS) is taken from Zhang et al. (2009) who harvested 7 day-old seedlings growing in liquid culture under continuous light. (A) oeTPS response plotted against the İTPS response at 4 h after induction of TPS. Of the 5.2K responsive transcripts reported by Zhang et al. (2009), 4956 were found in the FTPS response data set. No FDR filter was applied to the FTPS dataset for this plot. (B-D) oeTPS response plotted against the FTPS response for the 2437 transcripts that responded significantly (FDR<O.05) at both 4 h and 6 h after spraying (termed ‘FTPS 4-6h’ in the display). Data were plotted separately for each CRF group of genes: (B) 1596 transcripts assigned to CRF G_p_ (C) 494 transcripts that were assigned to CRF G_2_ and (D) 347 transcripts that were assigned to CRF G„. Transcripts were assigned to CRF G_1_ G_2_ and G_o_ as explained in Supplemental Figure S2. The İTPS response of transcripts in G_t_ is probably a direct response to elevated TreδP, in G_2_ to lower sugar and G_0_ to more complex interactions. Plots of oeTPSA against the individual 4-h and 6-h FTPS responses are provided in Supplemental Figure S12A-B. (E-F) Enriched pathways based on Gene Ontology. The analysis was performed for DEGs from the oeTPS data set of Zhang *et¤l.* (2009) that were assigned to G_l·_ in both FTPS datasets (4 h and 6 h). (E) This shared set of transcripts was analyzed using the TagCrowd on-line tool (https://tagcrowd.com/) to identify frequently occurring terms among the gene names and descriptions and are shown in a word map, with the font size representing the frequency. (F) Histogram depicting the fold enrichment (left y-axis) and p value (righty-axis) of the top 30 enriched processes. An analysis of all enriched processes is provided in Supplemental Figure S12D. (G) Comparison of the oeTPS (Zhang *etal.,* 2009) and FTPS responses for genes assigned to ribosome biogenesis, both plotted on a log_2_ scale. The plot shows the FTPS response at 4 h and at 6 h after spraying. The number of genes shown in this display is less that in panel B because not all of the genes in the FTPS response were present in the data set of Zhang *etal.* (2009). Although the oeTPS data of Zhang et al. (2009) showed the strong response of ribosome biogenesis, this was not explicitly noted at the time because assignment of genes to the ribosome biogenesis category was very incomplete in the ontology that they used.

Zhang et al. (2009) reported that oeTPS repressed genes involved in photosynthesis, the glyoxylate cycle and gluconeogenesis, and induced genes involved in mitochondrial electron transport, amino acid synthesis, nucleotide synthesis and protein synthesis. Our analysis of the iTPS response shows that many of these responses are due to Tre6P signaling, and reveals further Tre6P-mediated responses like the opposing response of cytosolic and mitochondrial ribosomal proteins compared to chloroplast ribosomal proteins. Paul et al. (2010) noted that seedlings with constitutive overexpression of TPS were small and stunted, leading them to investigate the response of transcripts that might contribute to this phenotype, like cell wall biosynthesis and light- and auxin signaling. These responses are partly confirmed in the iTPS response. There was strong agreement for genes involved in light-signaling; these were mainly repressed in both oeTPS and iTPS and the vast majority were assigned to CRF G1 implying repression by Tre6P (Supplemental Figure S12C, Supplemental Text). Others like cell wall modification and auxin signaling include many indirect effects (Supplemental Figure S7G, Supplemental Figure S11, see also below).

Genes that respond to a short-term elevation of Tre6P, are assigned to CRF G1, and respond in a qualitatively similar manner to constitutive oeTPS represent a very robust set of Tre6P-regulated genes. They are listed in Supplemental Dataset S6 and a GO analysis is provided in Supplemental Dataset S7 and summarized in Figure 6E-F and Supplemental Fig 12D. Downregulated processes included responses related to photosynthesis, chlorophyll synthesis and pigment metabolism, carbon utilization, monosaccharide metabolism, generation of precursor metabolites and energy, amino acid catabolism as well as various light responses and the circadian clock. The most enriched upregulated process was cellular component organization or biogenesis, which includes enrichment of categories related to ribosome biogenesis and mitochondrial biogenesis.

### Impact of elevated Tre6P on genes assigned to trehalose metabolism

We next investigated signaling pathways that might contribute to the transcriptional response to elevated Tre6P. The transient elevation of Tre6P was achieved by induced expression of a heterologous bacterial TPS. We first asked how the endogenous Tre6P pathway responds to this sudden imposed increase in Tre6P (Figure 7, Supplemental Figure S13). Tre6P is synthesized by TPS1, whilst TPS2-4 are catalytically active but only expressed at a specific stage of seed development (Delorge et al., 2014; Fichtner and Lunn, 2021). TPS1 was assigned to G2 and repressed, possibly due to the decrease in sugar. TPS5-11 are termed ‘class II’ TPSs and lack catalytic activity (Ramon et al., 2009; Lunn and Fichtner 2021). Except for *TPS7*, all class II *TPS*s were repressed and all of these except *TPS5* were assigned to G1. This observation indicates that Tre6P represses most of the class II *TPSs*. Arabidopsis possesses ten diverse TPPs (Vandesteene et al., 2012). These were assigned to G2 or G0 with two being induced and six repressed. These observations point to large-scale rewiring of Tre6P metabolism in response to an imposed increase in Tre6P (see Discussion).

**Figure 7.**
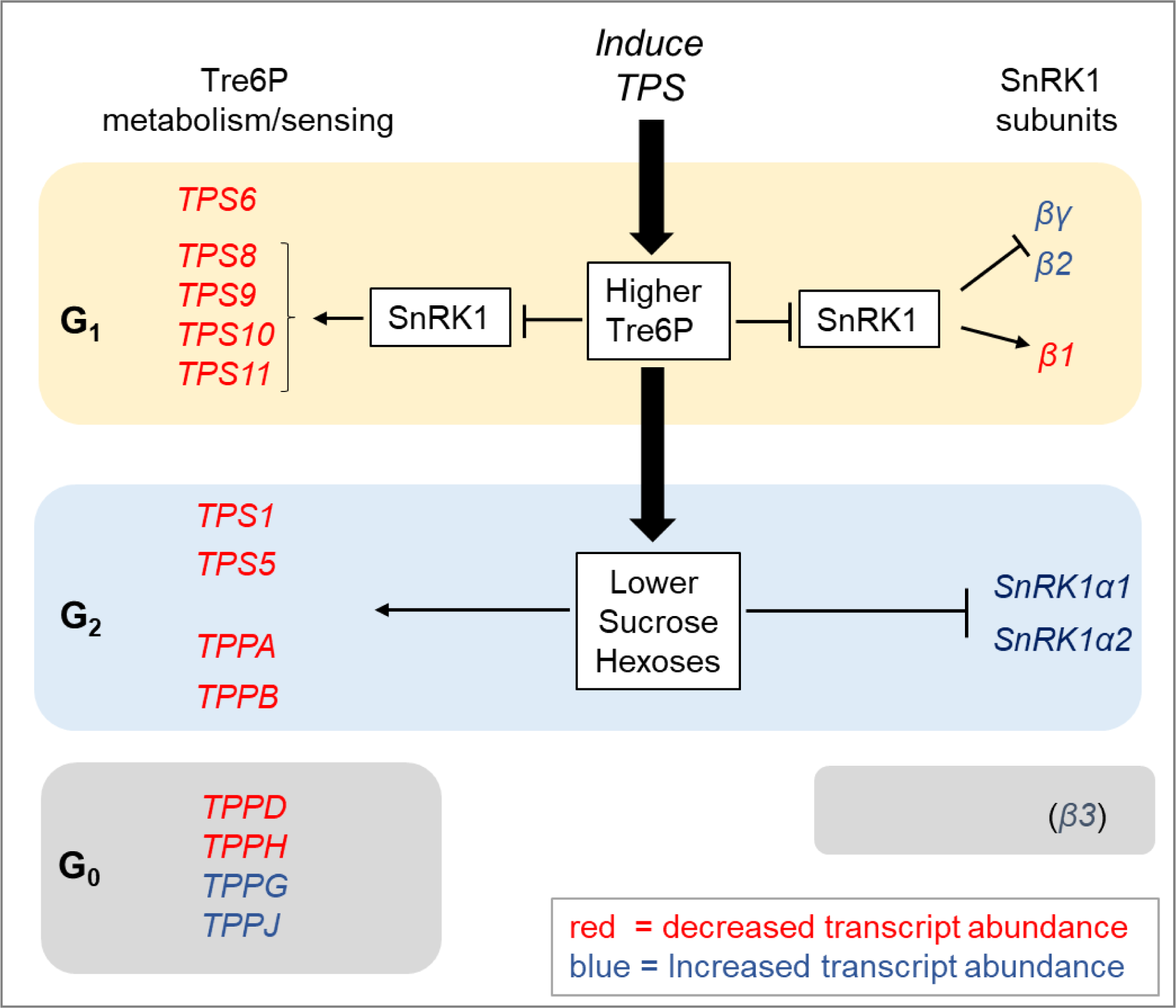
Schematic overview summarizing responses of transcript encoding proteins involved in Tre6P metabolism and subunits of the SnRKl complex. *TREHALOSE-6-PHOSPHATE SYNTHASE (TPS)* and *TREHALOSE-6-PHOSPHATE PHOSPHATASE (TPP)* genes and SnRKl subunit genes whose transcripts responded to elevated TreSP were assigned to CRF groups G_l_, G_2_ and G_o_. Based on responses to transient over-expression of SnRKlαl (Baena-González et al., 2007), it could be inferred that most of the genes in CRF group G-į were responding due to inhibition of SnRKl by TreSP. The genes in CRF group G_2_ are probably responding to the decrease in sucrose and other sugars that follows an induced rise in TreSP levels, rather than the rise in TreSP *per se.* Genes in CRF group G_o_ respond to TreSP but their expression appears not to be regulated by sugars. The response of SnRKlβ3 is shown in brackets because it inconsistent at 4 h and 6 h. Up- and down-regulated genes are shown in blue and red, respectively. The display is based on data provided in Supplemental Figures S13 and S14A.

### Impact of iTPS on SnRK1 expression

Tre6P is known to inhibit SnRK1 activity *in vitro* and it is likely that this interaction underlies at least some SnRK1 signaling functions *in vivo* (see Introduction). We therefore inspected the relationship between the iTPS response and SnRK1 signaling.

We first asked if Tre6P modifies expression of the SnRK1 subunits (Figure 7, Supplemental Figure S14A). SnRK1 is a hetero-trimeric protein complex containing one of two alternative catalytic subunits (SnRK1α1, SnRK1α2), one of three alternative regulatory β-subunits (SnRK1β1, SnRK1β2, SnRK1β3) and a regulatory SnRK1βγ subunit (Polge et al., 2008; Broeckx et al., 2016; Nietzsche et al., 2016; Wang et al., 2020). In the iTPS response, both catalytic subunits were assigned to G2 and were induced, presumably in response to falling sugar. Except for SnRK1β3, the regulatory subunits were assigned to G1 with two being induced (SnRK1β2, SnRK1βγ) and one repressed (SnRK1β1). SnRK1β3 was assigned to G0 and showed an inconsistent response, with weak induction at 4h and no significant change at 6h. These results point to low sugar inducing the catalytic subunits, whilst Tre6P modifies expression of the regulatory subunits (see Discussion).

### Comparison of the iTPS response with SnRK1 signaling

We next interrogated the iTPS response to learn if inhibition of SnRK1 plays a major role in Tre6P signaling i*n vivo*. The published response to transient *SnRK1α1* overexpression in protoplasts (Baena-Gonzalez et al., 2007; hereafter termed the tSnRK1α1 response) has been widely used to define transcriptional events downstream of SnRK1. Supplemental Figure S14B shows the iTPS response of the top 25 responders from Baena-Gonzalez et al. (2007) and four further genes that have been widely used as SnRK1 marker genes (*DIN1* and *DIN3*) or intensively studied as key players in the transcriptional regulation of C-responsive genes (*bZIP11*and *bZIP63*) (Ma et al., 2011; Mair et al., 2016). Of these 29 genes, 14 are induced by tSnRK1α1 and, of these, 12 were repressed in the iTPS response and assigned to G1, consistent with their repression being due to Tre6P inhibiting SnRK1. The exceptions were *DIN1*, whose transcript abundance did not respond significantly to iTPS, and *DIN6* that was assigned to G2 and induced. A less consistent picture emerged for the 15 genes that are repressed by tSnRK1α1. Five were assigned to G1 and induced, consistent with elevated Tre6P inhibiting SnRK1. Five were assigned to G2 and one to G0 and repressed, which is inconsistent with their response being due to inhibition of SnRK1 by Tre6P, but consistent with it being due to inhibition of SnRK1 by signals deriving from low sugars or other side-effects of the iTPS treatment. The remaining three genes did not show a significant iTPS response. Despite the more complex response of the SnRK1-repressed genes, the main conclusion from this analysis is that 17 of the 29 genes were assigned to G1 and responded in the direction that is predicted if Tre6P inhibits SnRK1 *in vivo*.

These findings encouraged us to inspect the iTPS response for all 1021 potential SnRK1 downstream target genes listed in Baena-González et al. (2007). Of these, 1004 were present in the iTPS dataset. When the complete 4-h and 6-h iTPS response was compared with the tSnRK1α1 response there was a weak negative trend (R^2^ = 0.14 and 0.21. respectively, Supplemental Table S6). Many transcripts showed a qualitatively similar rather than the expected reciprocal response (Figure 8A). When the iTPS response was deconvoluted to retain only transcripts in CRF group G1, a very strong negative correlation emerged (580 and 542 genes, R^2^ = 0.64 and 0.68, p = 7.6×10^-132^ and 7.0×10^-54^ in the 4-h and 6-h iTPS response, respectively) (Figure 8B, Supplemental Table S6). A similar picture emerged after filtering to focus on the top 100 tSnRK1α1 responders (see Supplemental Figure S14C). These genes presumably represent downstream targets where SnRK1 signaling is inhibited by elevated Tre6P. An analogous analysis with transcripts assigned to G2 yielded a relatively good positive regression (144 and 151 genes, R^2^ = 0.41 and 0.29, p = 2.9×10^-18^ and 1.44×10^-12^, respectively, Supplemental Table S6, Supplemental Figure S14D). These presumably represent genes that are downstream of SnRK1 and whose response is not counteracted by elevated Tre6P but is instead promoted by other signals, for example, low sugars. Transcripts assigned to G0 yielded a very weak relationship (22 and 22 genes, R^2^ = 0.042 and 0.074, p = 0.36 and 0.22, respectively (Supplemental Table S6, Supplemental Figure S14D,). Although these genes were not scored as sugar-responsive in our CRF analysis, many responded not only to elevated Tre6P but also to transient overexpression of SnRK1, although in a diverse manner.

**Figure 8.**
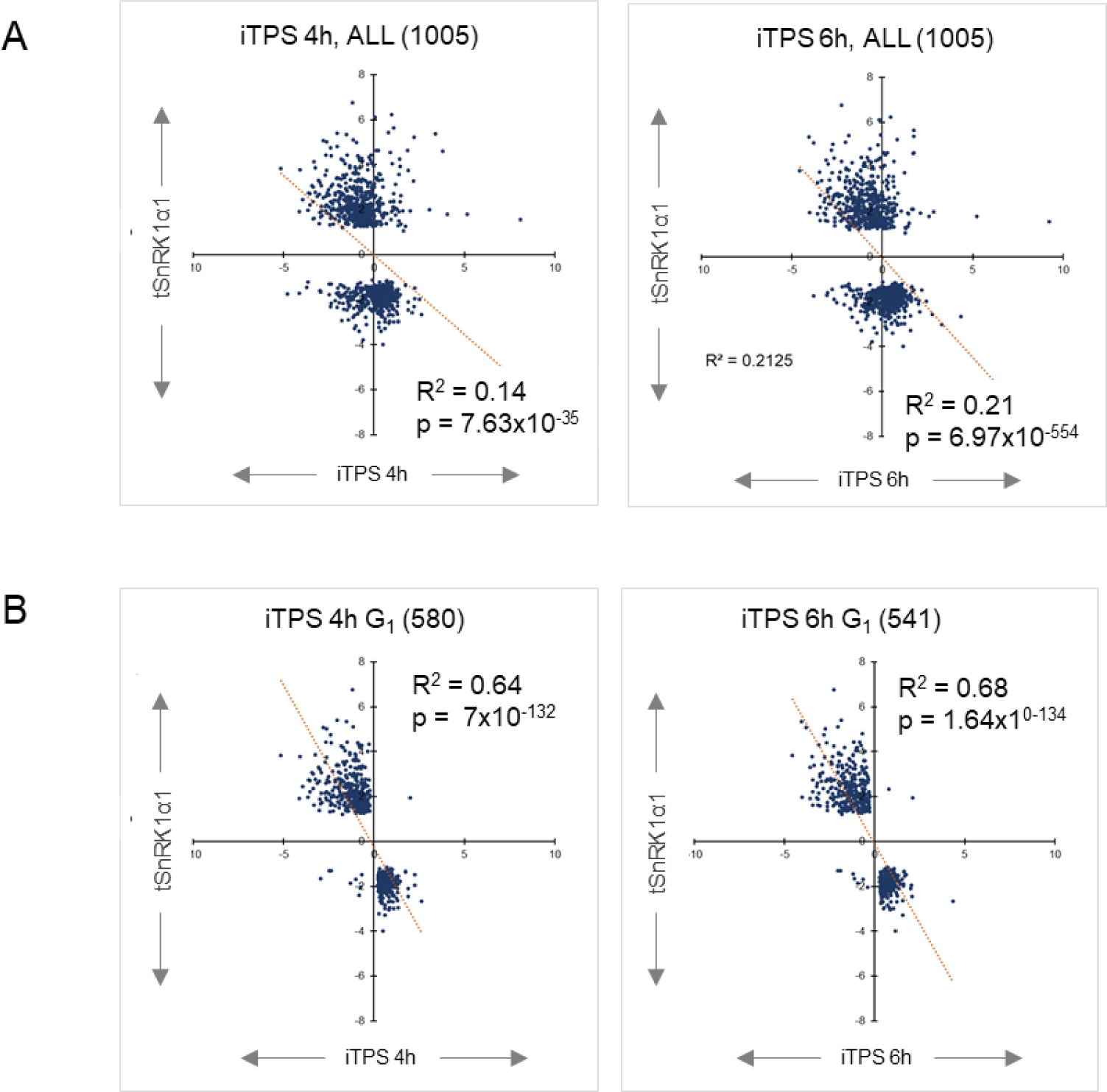
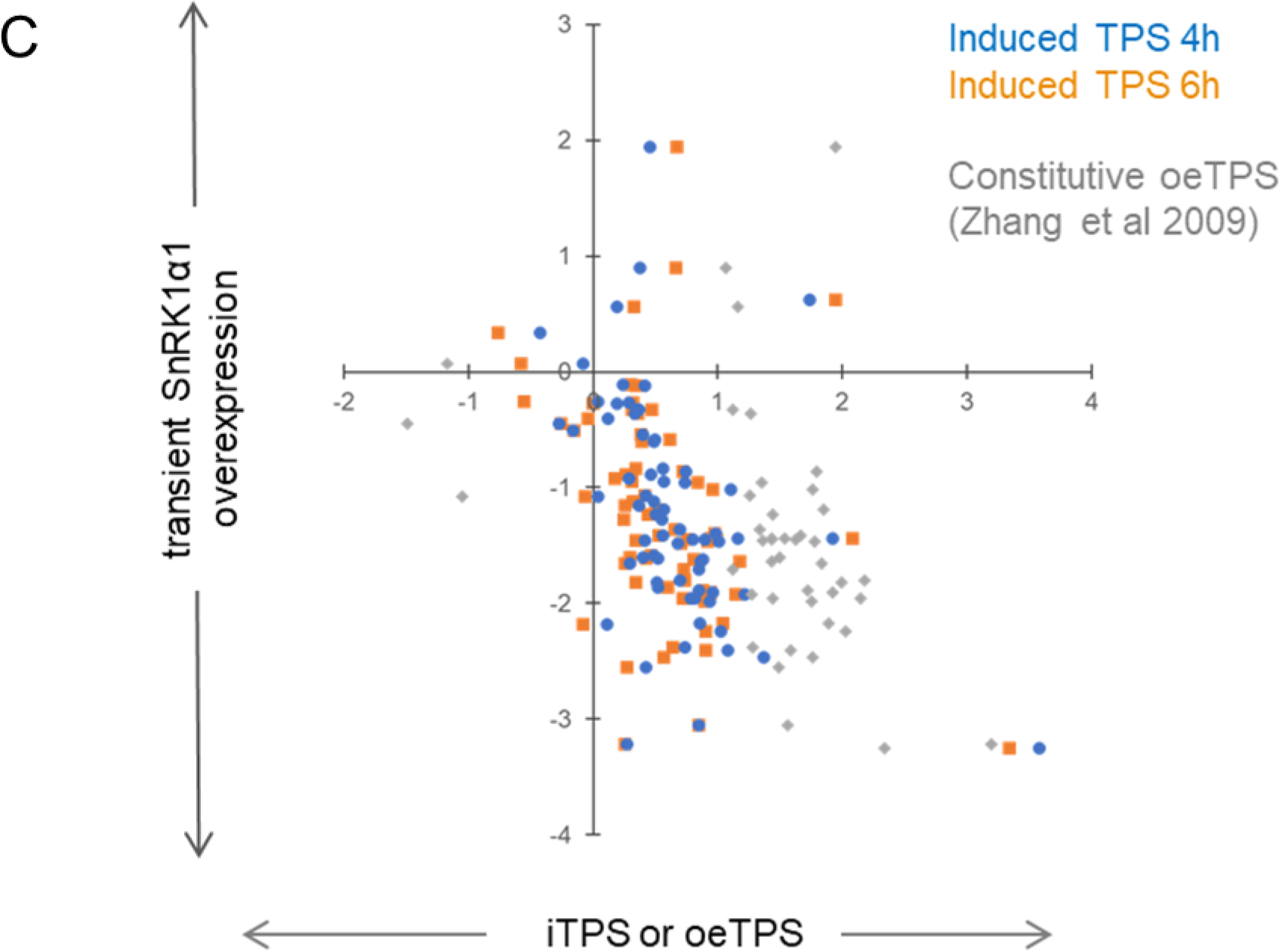
Response of SnRKl-regulated transcripts to elevation of Tre6P levels. A list of SnRKl-regulated transcripts was drawn up based on the data for the response to transient overexpression of SnRKlαl in Arabidopsis mesophyll protoplasts (Baena-Gonzalez et al., 2007, here termed the tSnRKlαl response). A total of 1001 of these transcripts was retrieved in the unfiltered iTPS data set. (A) Regression plot for all 1001 genes of the tSnRKlαl response versus the response at 4 h (left and 6 h (right) after spraying ÎTPS29.2 with ethanol. In the 4-h and 6-h ÎTPS samples, 763 and 762 transcripts, respectively, showed a qualitatively opposite response to their tSnRKαl response, whilst 242 and 243 transcripts, respectively, showed a qualitatively similar response to their tSnRKlαl response. (B) Regression plot for all 1001 DEGs of the tSnRKlαl response versus the filtered Gļ iTPS response. Transcripts were filtered (FDR > 0.05, log_2_FC ≥ 0.2) and then compared with the CRF (Supplemental Figure S2) to assign transcripts to G_l·_ (i.e., transcripts whose iTPS response is qualitatively similar to their CRF and probably a direct effect of elevated Tre6P). A total of 580 and 541 transcripts were assigned to Gj in the iTPS 4 h and iTPS 6-h data sets, respectively. Of these transcripts, at 4-h and 6-h iTPS, the vast majority (571 and 532, respectively) showed a qualitatively opposite response to their tSnRKlαl response, whilst at both times only 9 transcripts showed a qualitatively similar response to their tSnRKlαl response. Further information about these analyses and the correlations between tSnRKlαl response and transcripts assigned to iTPS CRF groups G_2_ and G_o_ is provided in Supplemental Figure S14 and Supplemental Table S6. (C) Regression plots of the tSnRKlαl response and the iTPS and oeTPSl responses of genes encoding ribosome assembly factor (next page). (C) Regression plots of the tSnRKlαl response and the iTPS and oeTPSl response of genes encoding ribosome assembly factor. The plot shows all 74 genes assigned to the subBIN ‘ribosome biogenesis’ in the MapMan TAIR10 ontology. Of these, 54 were assigned to CRF group Gļ and 10 to CRF group G_o_, respectively, in at least one of the two iTPS treatments, and only 4 were unassigned. The responses in the 4h and 6h iTPS treatments were similar and those in the oeTPS response were qualitatively similar but stronger than in the induced treatments. As reported in Baena-González et al. (2007), tSnRKlαl represses ribosome assembly genes (see also Supplemental Figure S14I). The vast majority of the changes in response to overexpression of TPS were therefore reciprocal to the response to tSnRKlαl.

Global analyses with the >500 genes assigned to CRF group G1 confirmed that there was a higher proportion of expected responses (i.e., iTPS G1 qualitatively opposite to tSnRK1α1) for genes that are induced by tSnRK1α1 than for genes that are repressed by tSnRK1α1 (Supplemental Figure S14E). This points to Tre6P playing a large role in SnRK1 signaling that represses genes, but a smaller role in SnRK1 signaling that induces genes.

We compared the iTPS and tSnRK1α1 responses in three sets of genes related to metabolism, growth and signaling; photosynthesis, light signaling and cytosolic ribosomal proteins (Supplemental Figure S14F-H). The responses tended to be reciprocal, but there were large differences in magnitude resulting in very low correlation coefficients. The strongest negative correlation (R^2^ = 0.3-0.42) was found for ribosomal proteins after filtering to focus on CRF group G1 (i.e., genes responding to elevated Tre6P). As already seen by Baena-González et al. (2007), many ribosome assembly factors are induced by sugars and repressed by tSnRK1α1 (Supplemental Figure S14I). Ribosome assembly factors were induced by an induced elevation of Tre6P and even more strongly by a constitutive increase in Tre6P, and these responses were reciprocal to the tSnRK1α1 response (Figure 8C), consistent with them resulting from Tre6P-inhibition of SnRK1.

As already mentioned, increased C availability typically leads to repression of *TPS8-TPS11* and a switch from the β1 to the β2 subunit of SnRK1. We compared the iTPS and tSnRK1α1 responses for these genes (Supplemental Figure S14J). Five TPS class II genes (*TPS6*, *TPS8, TPS9, TPS10*, *TPS11*) and the β1 and β2 subunits of SnRK1 responded in a way consistent with elevated Tre6P inhibiting SnRK1. Overall, our analyses highlight that there are close interactions at multiple levels between Tre6P- and SnRK1-signaling (see Discussion).

### Interaction with TOR signaling

In mammals, yeast and plants, TOR acts as a counterpart to AMPK/SNF1/SnRK1 to positively regulate ribosome biogenesis (Sabatini, 2017; Ryabova et al., 2018; Wu et al., 2019; Meng et al., 2022; Scarpin et al., 2022). There is emerging evidence for multi-layered interactions between TOR and SnRK1 in plants (Nukarinen et al.,2016, Wang et al., 2018; Belda-Palazon et al., 2020). We investigated whether iTPS alters the levels of transcripts that encode TOR subunits or known post-translational targets of TOR (Supplemental Figure S15, see Supplemental text for details).

There was no obvious impact of elevated Tre6P on expression of the TOR subunits (Supplemental Figure S15A), or the downstream kinases S6 KINASE (S6K) and YET ANOTHER KINASE 1 (YAK1). However, the responses of two LA-RELATED PROTEIN 1 kinase family members (LARPs) were consistent with them being induced by Tre6P (Supplemental Figure S15B). LARP proteins are involved in the TOR-LARP1-5’TOP signaling axis that regulates expression of 5’TOP mRNAs, including transcripts encoding ribosome assembly factors and ribosomal proteins (Scarpin et al., 2020; 2022). Comparison of the iTPS and tSnRK1α1 responses also indicated that Tre6P-inhibition of SnRK1 signaling leads to induction of *NUCLEOSOME ASSEMBLY PROTEIN 1* (*NAP1:1*) and *RIBOSOMAL PROTEIN S6* (*RS6)* (Supplemental Figure S15C) which jointly promote transcription of rRNA (Son et al., 2015). Overall, this may contribute to the increased expression of ribosomal proteins and assembly factors observed in the iTPS G1 response (see Figure 3, Supplemental Figure S7).

ABA is sensed via PYROBACTIN RESISTANCE/PYROBACTIN RESISTANCE-LIKE (PYR/PYL) receptors. These ABA receptor proteins are known to be phosphorylated and inactivated by TOR (Wang et al., 2018a). Transcript abundance for seven of the eight ABA receptors was decreased in the iTPS response, including four that were assigned to CRF group G1 (Supplemental Figure S15D). Some of these genes were also strongly repressed by constitutive oeTPS (Zhang et al., 2009) and most were significantly induced by tSnRK1α1 (Baena-Gonzaléz et al., 2007). These observations are consistent with Tre6P acting via inhibition of SnRK1 to repress ABA signaling. Other known downstream phosphorylation targets of TOR (Scarpin et al., 2020; Meng et al., 2022; Liao et al., 2022) were also found in the G1 iTPS response, including several initiation and elongation factors (Supplemental Figure S4E). Overall, our data point to a synergy between post-translational regulation of ribosome assembly, translation and other processes like ABA signaling by TOR, and transcriptional regulation of these processes by Tre6P-mediated signaling.

### FCS-LIKE ZINC FINGER (FLZ) family protein

FLZ family proteins (Jamsheer et al., 2015) are negative regulators of SnRK1 and are implicated in interactions between SnRK1 and TOR (Nietsch et al., 2014; 2016; Jamsheer and Lamxi, 2015; Jamsheer et al., 2015; 2018a; 2018b; 2022; Bortlik et al., 2022). Comparison of the iTPS response with the CRF and the tSnRK1α1 response added to the evidence that expression of the *FLZ* family is highly regulated by C status. It also pointed to Tre6P inhibition of SnRK1 contributing to regulation of a subset of *FLZs* that are induced in high sugar, whereas other signaling pathways are involved for *FLZs* whose expression increases in low C conditions (Supplemental Figure S16, Supplemental text).

### Interaction with brassinosteroid signaling

The emerging evidence for links between sugar- and BR-signaling (Zhang et al., 2016; 2021; Liao et al., 2022) prompted us to inspect the iTPS response of genes assigned to BR synthesis and signaling (Supplemental Figure 17 and Supplemental text). iTPS repressed genes in the biosynthesis pathway, probably via indirect effects, but elevated Tre6P had a direct impact on signaling components like *BRASSINOSTEROID INSENSITIVE 2* (*BIN2*), *BRASSINOSTEROID ENHANCED EXPRESSION1* (*BEE1*) and *BEE2*. Zhang et al. (2016) reported that eight *EXPANSIN* family members were repressed by BR-signaling. They were all repressed in the iTPS response, but probably indirectly as they were assigned to CRF group G2. This observation prompted us to examine the response of the entire *EXPANSIN* family, as well as the *XTH* family that is also involved in cell wall modification (Supplemental Figure S18). This analysis confirmed (see Supplemental Figure S7G) that the repression of genes for cell wall modification in the iTPS responses is mainly indirect, and indicated links with BR signaling.

### Comparison with bZIP11 signaling

S1 and C class bZIP proteins play an important role in low energy signaling (Dröge-Laser and Weiste, 2018). S1 bZIPs are translationally regulated by a mechanism in which sucrose acts at upstream open reading frames (uORFs) to stall ribosome progression (Wiese et al., 2004; Rahmani et al., 2009; Juntawong et al., 2014). When their translation is increased by falling sucrose, they interact with C class bZIPs to transcriptionally activate starvation responses and inhibit growth (Hanson et al., 2008; Ma et al., 2011, Dröge-Laser and Weiste, 2018). We compared the iTPS response with the published response to constitutive overexpression of bZIP11 (termed oebZIP11, data from Ma et al., 2011) (Supplemental Figure S19, see also Supplemental text). Many genes that respond to oebZIP11 also responded to iTPS, with some being assigned to CRF group G1 and others to group G2. Overall, our analyses pointed to many genes being regulated by both Tre6P/SnRK1 signaling and bZIP11 signaling, often acting in a mutually-reinforcing manner but in some cases in an antagonistic manner (see Supplemental text for details).

### Changes in expression of transcription factors

We also inspected the response of transcription factors (TFs), a subset of genes that might give independent insights into the transcriptional response to elevated Tre6P. Over 400 TFs showed a log2FC≥1, and more than 100 were assigned to CRF group G1 (see Supplemental Figure S20).

To provide an overview of the G1 response we performed both GO and STRING analyses (Supplemental Figure S21). The latter utilizes available datasets, published work, automated text mining and computational predictions from various organisms to score the likelihood of association between proteins (Szklarczyk et al., 2021). Both approaches are biased towards TFs in processes where understanding of transcriptional networks is more advanced. These analyses highlighted TFs that regulate carbohydrate and C-signaling, biosynthesis (chlorophyll, anthocyanin, glucosinolate), the circadian clock, and many signaling pathways related to light (shade avoidance, red/far red and blue light signaling), hormones (auxin, gibberellic acid, ethylene, ABA, jasmonic acid) and development (flowering, phloem or xylem development). This strengthened conclusions drawn from PageMan and GO analyses of the entire iTPS response (Figure 3, Supplemental Figures S7 and S11) including the strong interaction between Tre6P- and light-signaling. The most highly linked TF in the STRING analysis was ELONGATED HYPOCOTYL 5 (HY5), which is a master regulator of thousands of genes and coordinates light, environmental and developmental signaling (Gangappa and Botto, 2016; Dröge-Laser et al., 2018). TF analyses revealed that many C and S1 bZIP family members showed significant changes in transcript abundance in the iTPS response (Supplemental Figure S22A-B), including *bZIP1*, *bZIP2*, and *bZIP44* in the S1 subfamily and *bZIP9*, *bZIP25* and *bZIP63* in the C subfamily, with *bZIP1*, *bZIP2*, *bZIP9*, *bZIP25* and *bZIP63* being assigned to CRF group G1 indicating that they are repressed by signaling downstream of Tre6P. *bZIP1 bZIP25* and *bZIP63* showed a reciprocal response in the tSnRK1α1 dataset (Baena-González et al., 2007) consistent with SnRK1 inhibition by Tre6P. Analyses of other TF families pointed to many other cases where Tre6P may act via SnRK1 to regulate TF expression (Supplemental Figure S23).

We also performed GO and STRING analyses of the iTPS G2 response (Supplemental Figure S23, Supplemental text). They highlighted TFs assigned to water relations, N metabolism, S starvation, and secondary metabolism (glucosinolate, flavonoid, anthocyanin biosynthesis), again resembling analyses of the entire iTPS G2 response (Figure 3, Supplemental Figures S7 and S11). They also uncovered an interaction between six MYB family members that induce glucosinolate biosynthesis (Mitreiter and Gigolashvili, 2021) that were assigned to CRF group G2 and repressed, presumably as a result of the decline in sugars, and a set of MYC TFs that act in jasmonic acid signaling (Dombrecht et al., 2007; Caldo et al 2011; Schweizer et al., 2013) and were in part placed downstream of Tre6P (Supplemental Figure S25)

## Discussion

### Elevation of Tre6P levels leads to large and widespread changes in transcript abundance

Tre6P is a sucrose-signaling metabolite that plays a central role in the regulation of plant metabolism, growth and development (Fichtner and Lunn, 2021). First insights into the transcriptional response to Tre6P were provided by analyses of plants with constitutive overexpression of TPS (Zhang et al., 2009; Paul et al., 2010). However, these plants showed strong growth and developmental phenotypes (see also Schleupmann et al., 2003; Yadav et al., 2014). Furthermore, elevated Tre6P leads to major changes in metabolism including decreased levels of sugars and increased levels of organic acids and amino acids (Martins et al., 2013; Yadav et al., 2014; Figueroa et al. 2016; Ishihara et al., 2022) that are likely to trigger secondary changes in gene expression. We have used Arabidopsis plants expressing a bacterial TPS under the control of an ethanol-inducible promoter to investigate the short-term response of the transcriptome to sudden elevation of Tre6P.

The induction system was chosen to avoid artifacts due to constitutive or unphysiologically high levels of Tre6P (Martins et al., 2013; Figueroa et al. 2016). However, TPS protein is induced in a wide range of cell types. AtTPS1, the endogenous protein that synthesizes Tre6P in Arabidopsis, is mainly expressed in the companion cells and phloem parenchyma (i.e., the phloem-loading zone) and guard cells in leaves, the root vasculature, and the shoot apical meristem (Fichtner et al., 2020). Although it is likely that the Tre6P moves via plasmodesmata into neighboring cell types, there might be spatial gradients that are overridden when TPS is expressed in a less spatially-resolved manner. Despite this caveat, we argue that our inducible promoter system enables us to investigate primary responses to Tre6P more easily than in plants with constitutively elevated Tre6P.

Almost half the detected transcripts showed significant changes and >5000 transcripts showed >2-fold changes in abundance 4h after ethanol-spraying (Figure 2A), which is within 2h of the first detectable rise in Tre6P (Figure 1). This resembles the number of transcripts that show changes in abundance during diel cycles when the C supply, light and the circadian clock act in concert to regulate gene expression (Bläsing et al., 2006; Usadel et al., 2008; Flis et al., 2016). The increase in Tre6P was accompanied by a simultaneous decline in sucrose (Figure 1) and changes in the levels of other metabolites (Supplemental Figure S4) that might themselves lead to changes in gene expression. Thus, even our use of an inducible system and short sampling times did not entirely remove complications due to changes in the levels of other metabolites. This may reflect the rapid action of Tre6P on metabolism via post-translational mechanisms (Figueroa et al. 2016).

### Deconvolution of Tre6P-dependent and indirect responses

To distinguish responses to elevated Tre6P from responses to lower sucrose and other indirect effects, we compared the response to induction of TPS (iTPS) with the response to increased sugar availability. To do this we calculated an average response, which we termed a carbon response factor (CRF), across nine published treatments that increased sugar levels whilst minimizing confounding changes in light- or circadian-signaling (Supplemental Figure S2). We considered transcripts that showed a qualitatively similar response to iTPS and elevated sugar to be potential targets of Tre6P-signaling (CRF group G1), whilst transcripts that showed opposite responses were more likely to be responding to the decline in sucrose or other indirect effects (CRF group G2). Assignment of a gene to group G1 or G2 does not mean that it is regulated only by Tre6P or only by signals from sucrose or other indirect effects; it is possible that in some cases expression is regulated by both, and the observed change depends on the relative strengths of the responses to 3- to 4-fold elevated Tre6P compared to a 30% decrease in sucrose or other secondary effects. Our analysis also identified many transcripts that responded to induction of TPS but did not show an obvious response to changes in sugar levels (CRF group G0). Overall, of the responding transcripts, about 40% responded in a manner consistent with a response to elevated Tre6P and about 27% in a manner consistent with them responding to lower sucrose or other indirect effects, whilst about 33% could not be assigned to either response type. We conclude that there are massive transcriptional responses within 2h of Tre6P levels starting to rise, but also massive indirect effects.

It was unexpected that so many transcripts responded to a rise in Tre6P but did not respond to elevated sugar levels, at least, not in the treatments we used to estimate the CRF. This is partly due to two technical issues. One is that some transcripts responded to sugars in a context-dependent manner, rising in some and falling in other treatments. Another is that some responses may have been missed because the CRF was calculated using data from ATH1 arrays, which can be insensitive to changes of low-abundance transcripts. However, these technical factors explained only a small proportion of the unexpected responses (Supplemental Figure S6). It remains possible that some genes respond to sugars in other conditions. However, another explanation is that their expression is regulated in an opposing manner by Tre6P and by signals that derive from or change in concert with sucrose. In most situations in wild-type plants, such genes would not respond strongly or consistently because sucrose and Tre6P usually change in parallel (Lunn et al., 2014; Figueroa et al., 2016; see also Introduction). However, they would respond to perturbations that change the relationship between Tre6P and sucrose, or that differentially modulate Tre6P-depdendentzand -independent signaling. The link between sucrose and Tre6P can be modified by mid-term environmental changes and during developmental transitions (Carillo et al., 2013; Figueroa and Lunn, 2016; Fichtner and Lunn, 2021, see Introduction for further references). Changes in SnRK1 activity can also impact on the relationship between Tre6P and sucrose (Peixoto et al., 2021). It is worth noting that some genes in CRF group G0 were responsive to transient expression of SnRK1α1 or overexpression of bZIP11, (Supplemental Figures S14D, S19C), providing further evidence that they are regulated by sugar-signaling.

We revisited a published cDNA-based microarray study that used plants with constitutive overexpression of TPS (Zhang et al., 2009) (Figure 6, Supplemental Figure S12). In this study, about 5000 transcripts showed >2-fold changes in abundance compared to wild-type plants. In our study, about 30% of these transcripts responded to a short-term increase in Tre6P (i.e., were assigned to CRF group G1) and showed qualitatively similar changes in abundance in the induced and constitutive responses (Figure 6B). Even though the quantitative responses differed between the induced and constitutive responses, the overall agreement was highly significant (Figure 6B). This agreement is striking as different tissues (seedlings versus rosettes), growth conditions and RNA analysis techniques were employed, quite apart from it being likely that the magnitude of the change will often differ between induced and constitutive responses. There was much less overlap between the induced and the constitutive response for genes that we assigned to indirect responses; CRF groups G2 and G0 contained only 10 and 7%, respectively, of the genes listed in Zhang et al. (2009) and many of these changed in opposite directions in the induced and constitutive responses (Figure 6C-D). Over 50% of the genes that responded to constitutive overexpression of TPS did not show significant changes in the induced response and presumably reflect indirect responses to long-term elevation of Tre6P. Overall, this comparison identified about 1500 genes that respond robustly to both short- and long-term elevation of Tre6P, but also highlighted the importance of studying short-term responses and of distinguishing direct from indirect responses.

### When C availability increases, Tre6P-dependent and -independent signaling act collectively to promote biosynthesis, growth and defense

Genes whose expression was regulated by Tre6P were often from different functional categories to those whose expression was affected by the concomitant decline in sucrose and other indirect effects (summarized in Figure 5). Metabolic processes predicted to be transcriptionally regulated by Tre6P (Figure 3, Supplemental Figure S7A-E, S11) included repression of photosynthesis and associated processes like chlorophyll, tocopherol and flavanol biosynthesis, repression of gluconeogenesis, induction of nucleotide synthesis, repression of anthocyanin biosynthesis, and repression of sucrose export, especially *SWEET*s that are involved in sucrose export from source leaves. In contrast, N-assimilation, S-assimilation and large sectors of secondary metabolism, including glucosinolate, phenylpropanoid and flavonoid biosynthesis, were predicted to be mainly regulated by indirect effects like the decline in sucrose (Figure 5, Supplemental Figure S7B-F). Growth-related processes that were induced by Tre6P included mitochondrial biogenesis (Figure 6) and ribosome assembly and protein synthesis (Figures 4, 6), whereas indirect responses included repression of cell wall modification (Supplemental Figures S7G, S11, S17, S18).

In a wild-type plant, Tre6P and sucrose usually change in parallel (see Introduction). For transcripts assigned to CRF group G1, the response of wild-type plants to rising C availability is predicted to resemble their iTPS response. In contrast, for transcripts assigned to CRF group G2, the response of wild-type plants to rising C availability is predicted to be qualitatively the opposite of their iTPS response. This is illustrated in Supplemental Figure S26A, which is derived from the summary of response of different metabolic and growth processes to induced expression of TPS in Fig. 5, but with the direction of change reversed for processes assigned to CRF group G2. As C-availability rises, Tre6P-mediated and Tre6P-independent sugar signaling are predicted to act in concert to decrease investment in the photosynthetic apparatus and promote nutrient assimilation, biosynthesis, protein synthesis and cellular growth, as well as synthesis of specialized metabolites for defense purposes.

It has long been known that sugars repress genes that are involved in photosynthesis (Sheen 1990, Von Schwaeren et al., 1991, Krapp et al., 1993; Lastdrager et al., 2014). In some cases, this may be linked with changes in the C/N balance (Stitt and Krapp, 1999; Li et al., 2021; Wang et al., 2022) or pathogen signaling (Biemelt and Sonnewald, 2006; Doehlemann et al., 2008; de Haro et al 2019). Our data reveal a rapid and direct impact of sugar signaling, mediated by Tre6P.

SWEET11 and SWEET12 are involved in sucrose export from source leaves (Braun, 2022; Xue et al., 2022). It is curious that they are repressed by elevated Tre6P that, in wild-type plants, would be associated with higher sucrose in source leaves. A possible explanation is that, in wild-type plants, high sucrose in source leaves might be associated with limited utilization in sink tissues, with repression of *SWEET*s serving to increase retention of C in source leaves and promote nitrate assimilation and amino acid synthesis (Figueroa et al., 2016). Many genes involved in N assimilation and metabolism are induced by sugar (Vincentz et al., 1993; Krapp et al.,1995; Wang et al., 2000; Stitt et al., 2002; Coruzzi, 2003; Vidal et al., 2020). Our analysis of the iTPS response indicates that this is due to sugar signaling that is independent of Tre6P (Supplemental Figure S7B). As sucrose and Tre6P usually change in parallel in wild-type plants (see Introduction) it can be envisaged that Tre6P and sugar signaling act cooperatively in the transcriptional regulation of sucrose export and N assimilation. In addition, Tre6P acts post-translationally to activate NR and PEPC and stimulate amino acid synthesis (Figueroa et al., 2016) allowing rapid fine-tuning of C and N metabolism. Thus, seen in the context of the whole plant, repression of sucrose effluxers by Tre6P may be part of a cooperative network that rebalances C and N metabolism and restores growth in sink organs.

Protein synthesis is positively regulated by sugars, which promote both polysome loading and ribosome biogenesis (Juntawong and Bailey-Serres, Pal et al., 2013; Juntawong et al., 2014; Nelson and Millar, 2015; Flis et al., 2016; Ishihara et al., 2017). Translation initiation is stimulated by TOR (Lastdrager et al., 2014), and ribosome biogenesis is stimulated by TOR (Lastdrager et al., 2014; Scarpin et al., 2022) and inhibited by transient SnRK1 overexpression (Baena-González et al., 2007). Our finding that elevated Tre6P rapidly induces genes for cytosolic and mitochondrial ribosomal proteins and ribosomal assembly factors (Figure 4, Supplemental Figure S7F) reveals a link between Tre6P-signalling and ribosome biogenesis. Induction of nucleotide biosynthesis by Tre6P (see above) may also contribute to increased ribosome biogenesis, as rRNA typically represents about 80% of total RNA (Warner, 1999). Increased ribosome biogenesis will presumably promote protein synthesis and cellular growth. More studies are needed to learn if this response is due to increased ribosomal biogenesis in growing sink leaves, or if it also occurs in mature source leaves where increased protein synthesis would presumably be linked with faster protein turnover. Protein turnover represents a substantial energy cost (Ishihara et al., 2017) and it would be interesting to learn if the rate of protein turnover is regulated in response to energy supply. In contrast, genes for plastidic ribosome proteins were mainly repressed. This presumably contributes to the repression of chloroplast function and photosynthesis in sugar-replete conditions.

Induction of TPS led to broad repression of *EXPANSIN* and *XTH* family members, (Supplemental Figures S7G, S18), which will presumably decrease cell wall modification and cell expansion (Cosgrove, 2005; Kaewthai et al. 2013). However, many *EXPANSIN* and *XTH* family members were assigned to CRF group G2, indicating that their repression might be related to the decline in sucrose after induction of TPS, rather than elevated Tre6P. This provides another example where Tre6P and sugar signaling act cooperatively to regulate a higher-level process; in this case, the balance between protein synthesis and cell expansion, with Tre6P-mediated signaling promoting protein synthesis and cellular growth, whilst Tre6P-independent signaling promotes *EXPANSIN* and *XTH* expression and cell expansion. At least for *EXPANSIN*s, repression may also be related to BR signaling (Supplemental Figure S17B).

### Tre6P-dependent signaling impacts many signaling processes

Tre6P acts transcriptionally on many signaling functions including the circadian clock, light signaling, ABA signaling, auxin signaling and floral induction. These categories were highlighted both in our analyses of the total transcriptome (Figure 6E-F, Supplemental Figures S9-S12, S15) and TFs (Supplemental Figures S20-21).

Light-signaling pathways were highlighted as downstream targets of elevated Tre6P in GO analyses of the response of the total transcriptome and of TFs (Supplemental Figure S11, S12C-D, S24A). There was a remarkably conserved repression of many light-signaling components between the induced and the constitutive (Zhang et al., 2009; Paul et al., 2010) responses (Supplemental Figure S12D). STRING analysis of the responses of TFs highlighted HY5 as a major hub, and PHYTOCHROME INTERACTING FACTOR4 (PIF4) and LONG HYPOCOTYL IN FAR-RED1 (HFR1) as further hubs. HY5 is a master regulator of thousands of genes and coordinates light, environmental and developmental signaling (Gangappa and Botto, 2016; Dröge-Laser et al., 2018). Sugars are known to inhibit *PIF4* expression (Shor et al, 2017; 2018) and this may contribute to modulation of PIF4 function and growth in varying light regimes (Moraes et al., 2019). Overall, as suggested by Paul et al. (2010), Tre6P interacts with and modulates light signaling, providing a mechanism whereby C availability can modify and tune light-induced morphogenesis and growth responses. Sucrose-dependent hypocotyl elongation was prevented in *tps1* mutants and SnRK1α1 overexpressors (Simon et al., 2018). It will be interesting to learn how the interaction between Tre6P and light signaling impacts on the parallel regulation of protein synthesis by Tre6P and cell wall modification by sugars to control expansion growth and plant composition and morphology.

The circadian clock was highlighted as downstream of Tre6P signaling in GO analyses of the response of the total transcriptome response and of TFs (Figure 6F, Supplemental Figures S11, S21). There were widespread changes in the expression of core clock components, with dawn components being induced and day, dusk and evening components repressed (Supplemental Figures S10, Supplemental text). This rapid response to a 3- to 4-fold increase in Tre6P underlines the sensitivity of clock dynamics to changes in the C status and is in broad agreement with earlier studies showing that sugars regulate the expression of many clock components (Dalchau et al., 2011; Haydon et al., 2017; Shin et al., 2017; Webb et al., 2019). However, the response to elevated Tre6P differs in details from that seen in previous studies. For example, low C and Tre6P promote SnRK1-dependent action of bZIP63, leading to increased *PRR7* expression and lengthening of clock period (Haydon et al., 2013; Frank et al., 2018; Viana et al., 2021) and sudden low-light perturbations lead to lower sucrose and Tre6P, increased *PRR7* expression and a small delay in clock progression (Moraes et al., 2019). In contrast, *PRR7* did not respond in our study and the changes in other clock transcripts were consistent with a delay in clock progression by Tre6P (see Supplemental text). This may be because our experiments investigated the response to elevated Tre6P, whereas most studies of the impact of sugars on the clock have addressed the impact of C-starvation on clock dynamics. Furthermore, metabolic regulation and light signaling interact to modify clock gene expression (Shin et al., 2017; Shor et al., 2017; 2018). Modified light-signaling (see above) may contribute to the response of the circadian clock to elevated Tre6P.

There is a close interaction between sugar signaling and ABA signaling (Rolland et al., 2006; Lastdrager et al., 2014) with the arrest of root growth by high sugar levels in the medium being alleviated in mutants defective in ABA sensing or signaling. There are also interactions between Tre6P and ABA signaling (Avonce et al., 2004; Ramon et al. 2007; Debast et al., 2011; Tian et al., 2019; Belda-Palazón et al., 2020, 2022), and reduced-function mutant alleles of *tps1* are hypersensitive to ABA (Gómez et al., 2010). It is known that TOR phosphorylates and inhibits all eight members of the PYR/PYL ABA receptor family (Meng et al., 2022). Strikingly, elevated Tre6P led to decreased transcript abundance for seven of the eight family members, with four responding in a manner consistent with them being repressed by Tre6P-dependent signaling, three of the four also being repressed by constitutive overexpression of TPS (Zhang et al., 2009) and all four being induced by transient overexpression of SnRKα1 indicating that Tre6P is acting via inhibition of SnRK1(Supplemental Figure S15D). This multilevel regulation of ABA receptors by TOR, Tre6P and SnRK1 may contribute to the cross-sensitization of sugar- and ABA signaling

### Impact of elevated Tre6P on genes in the Tre6P biosynthesis and signaling pathways

Introduction of a heterologous TPS will disturb the network that regulates and balances the Tre6P level with the production and use of sucrose. *TPS1*, which encodes the main enzyme responsible for synthesizing Tre6P (see Introduction), was repressed in the iTPS response but assigned to CRF group G2, indicating an indirect response possibly due to the decline in sucrose. In wild-type plants, rising C availability will presumably induce *TPS1* expression and make a midterm contribution to the increase in Tre6P levels as sucrose rises (Figure S26B). Several class II TPSs (*TPS6, TPS8*, *TPS9*, *TPS10* and *TPS11*) were repressed by elevated Tre6P, probably acting by inhibiting SnRK1, whilst *TPS5* was repressed by indirect effects (Figure 7, Supplemental Figures S13-S14). Of the ten *TPPs* in Arabidopsis (Vandesteene et al., 2012), two were induced (*TPPG*, *TPPJ*) and six repressed. Their assignment to G2 or G0 indicates these may be indirect responses (Figure 7). Overall, these observations point to large-scale rewiring of Tre6P metabolism. In the context of the iTPS response, these transcriptional responses may partly counteract the imposed increase in Tre6P. In the context of rising C availability in wild-type plants, cooperation between Tre6P-mediated and Tre6P-independent signaling is predicted to induce *TPS1* expression and promote Tre6P synthesis, repress many *TPS* class II genes and lead to diverse responses of *TPP*s (Supplemental Figure 26B). Based on emerging evidence that the TPS class II proteins interact with and inhibit SnRK1 (van Leene et al., 2022), their repression might provide feedforward amplification within the Tre6P-SnRK1 network. However, the repression involves mainly *TPS8-11*. These are strongly induced under C-starvation and repressed as C-availability rises (Usadel et al., 2008; Cookson et al., 2016; Flis et al., 2016) and their function is poorly understood. *TPS5, TPS6* and *TPS7*, which are expressed under more benign conditions, showed smaller and more varied responses (see Supplemental Figure S8). *TPP*s also showed diverse responses, possibly reflecting their differentiated roles in different biological contexts. Overall, there is a need for a more fine-grained understanding of the function of the various TPS class II and TPP proteins.

### Interaction between Tre6P and SnRK1 signaling

Plants possess a plethora of sugar-signaling pathways including the SnRK1, TOR, HEXOKINASE1 and REGULATOR OF G-PROTEIN SIGNALING1 mediated pathways, and inhibition of the translation of S1 type bZIP TFs by sucrose binding to uORFs (Rolland et al., 2006; Fu et al., 2014; Lastdrager et al., 2014; Dröge-Laser and Weiste, 2018; Crepin and Rolland, 2019; Baena-Gonzalez and Lunn, 2020; Meng et al., 2022; Scarpin et al., 2022). We investigated how Tre6P-signaling interacts with other sugar-signaling pathways (summarized in Figures 9-10).

**Figure 9.**
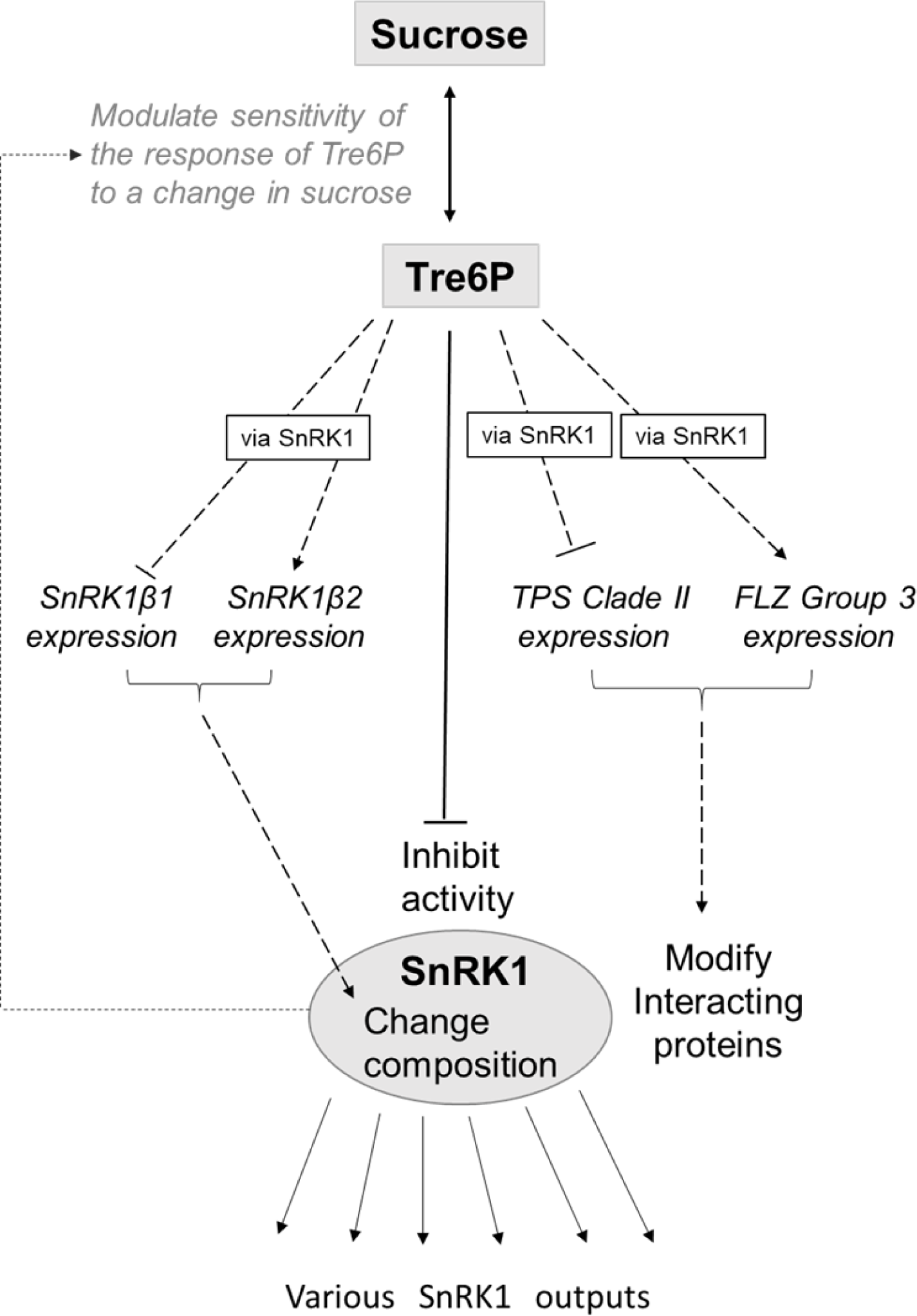
Interactions betweenTre6P and SnRKl signaling. The known inhibitory effect of TreδP in SnRKl activity is shown as a solid black line. In addition, TreöP regulates the expression of the two regulatory β-subunits of SnRKl, repressing *SnRKlβl* and inducing *SnRKlβ2,* which will probably lead o changes in SnRKl complex composition. TreδP also represses expression of several TPS Class II genes, in particular clade 2 *(TPS8, TPS9, TPS10, TPS11)* that were shown to physically interact with and at least in some cases may inhibit SnRKl activity (van Leene *et al.,* 2022). Furthermore, TreĉP induces FLZ group 3 genes, which also regulate SnRKl activity (see Nietsch et al., 2014; 2016; Jamsheer et al., 2018, 2022). These interactions are shown as dotted lines. The action of Tre6P on the SnRKl β-subunits, TPS class II and FLZ group 3 expression may be mediated via inhibition of SnRKl activity. The arrows to ‘various SnRKl outputs’ indicate that the changes in SnRKl composition and interacting proteins may modify if and how they operate. In addition, it has been shown that increased expression of tSnRKlαl dampens the response of Tre6P to sucrose (Peixoto *et* al., 2021) (grey line). TreSP-independent repression by rising sugar of the catalytic subunits *SnRKlal* and *SrıRKla2* is not shown.

**Figure 10.**
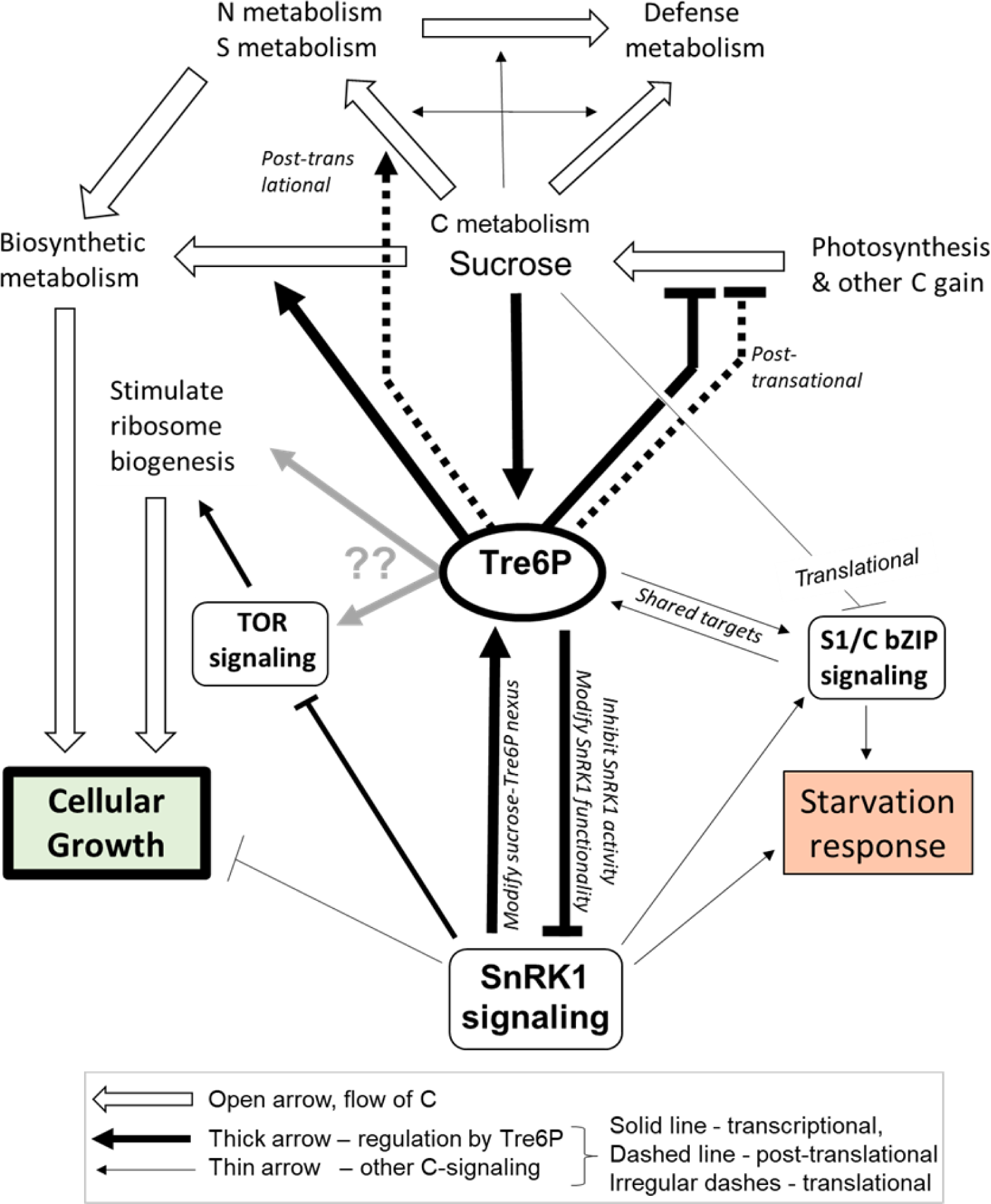
Regulation of metabolism and growth by TreβP. Flows of C, N and S are indicated by open arrows, transcriptional regulation and post-translational regulation by TreSP and thick solid arrows and thick dotted arrows, respectively. Thin solid arrows denote further C signaling. TreSP represses expression of genes encoding components of the photosynthetic machinery and post-translationally modified starch and sucrose breakdown. TreSP exerts positive transcriptional regulation on biosynthesis- and growth-related processes, in part by action via inhibition of SnRKl and via links to TOR. TreSP inhibits starvation responses via inhibition of SnRKl. The action of TreSP on SnRKl involves not only inhibition but also alteration of SnRKl composition, and modification of the expression of genes encoding TPS Class II proteins and Sļ/C FLZ proteins that interact with SnRKl and presumably modify its activity and functionality (see also Figure 9) as well as links and overlap with type Sj/C bZIP signaling. The induction of ribosome biogenesis may be at least partly explained by inhibition of SnRKl by elevated TreSP, but other mor direct links to TOR cannot be excluded (indicated by grey arrows) The action of TreSP on C metabolism is probably reinforced by other sugar-signaling pathways. TreSP is not directly involved in the transcriptional regulation of N and S assimilation, but acts at a post-translational level to promote C flux to organic acids (OA) and amino acids (AA). The synthesis of major sets of specialized metabolites like glucosinolates, phenylpropanoids and flavonoids appears to be regulated by sugar-signaling pathways other than TreSP, which may nevertheless make a small contribution (not shown). Links from sugar­signaling (largely TreSP-independent) to cell wall modification and expansion growth, and links from TreSP to light-signaling, the circadian clockand various hormone-signaling pathways are not shown.

It is known that Tre6P inhibits SnRK1 activity in extracts but the mechanism and biological function are not clear (see Introduction). We used our inducible system to re-examine the relation between Tre6P and SnRK1. Whilst there was a weak (R^2^ = 0.14-0.21) negative correlation between the response of all ∼1000 genes that responded to tSnRK1 α1 in protoplasts (Baena González et al., 2007) and their response to induced overexpression of TPS, many genes showed a similar rather than a reciprocal response (Figure 6A). The negative correlation became much stronger (R^2^ = 0.64-0.68) when the iTPS dataset was dissected to focus on the >500 transcripts that were probably responding to elevated Tre6P (Figure 6B). Good agreement was also seen for the response of a small set of strong responders in the tSnRK1α1 treatment (Supplemental Figure S14B). It is notable that the reciprocal relationship between the tSnRK1α1 response and the response to elevated Tre6P was much stronger for transcripts that were induced by tSnRK1α1 than for transcripts that were repressed by tSnRK1α1 (Supplemental Figure S14B, S14E). Many of the transcripts that are induced by tSnRK1α1 are related to the C-starvation response, whereas those that are repressed are mainly related to biosynthesis or growth. The implication is that rising Tre6P inhibits the SnRK1 starvation response in a rather consistent manner, whereas the interaction is more complex for genes that are involved in biosynthesis and growth. Analyses of four sets of genes related to metabolism, growth and signaling (photosynthesis, light signaling, cytoplasmic ribosomal proteins and ribosome assembly factors) confirmed that, although the response to elevated Tre6P and the response to overexpression of SnRK1α1 tended to be reciprocal, there were large differences in the strength of the response (Supplemental Figure S14F-H). The strongest correlation was found for ribosome proteins after filtering to focus on CRF group G1 (i.e., genes that are likely to be responding to elevated Tre6P) and for ribosome assembly factors (Figure 8C). The complex responses of many SnRK1α1-repressed genes might indicate that the Tre6P and SnRK1 responses are partly independent but could alternatively be explained by Tre6P acting to inhibit SnRK1 and further inputs acting downstream to modulate SnRK1 outputs (see also Avidan et al., 2023).

Our analyses also revealed that Tre6P exerts transcriptional control over SnRK1 expression (Supplemental Figure S14A). Whilst the response to induced overexpression of TPS was consistent with the SnRK1 catalytic subunits being induced by low sugar, expression of the regulatory subunits is regulated by Tre6P, with elevated Tre6P leading to increased abundance of *SnRK1βγ* and *SnRK1β2* transcripts and decreased abundance of the *SnRK1β1* transcript. This is reciprocal to their response to transient overexpression of SnRK1α1 (Baena-González et al., 2007, see also Supplemental Figure S14J). These observations are consistent with Tre6P acting via SnRK1 to regulate the expression of the regulatory β-subunits and, hence, SnRK1 complex composition. In wild-type plants, *SnRK1β2* typically predominates in C-replete conditions, and *SnRK1β1* is induced in low-C conditions (Usadel et al., 2008; Broeckx et al., 2016; Cookson et al., 2016; Flis et al., 2016; Peixoto and Baena-González, 2022). Our analyses indicate that this switch is due to the decline in Tre6P in low-C conditions and the resulting increase in SnRK1 activity (Supplemental Figure S26B).

TPS class II proteins were recently shown to bind SnRK1 and, in at least some cases, inhibit SnRK1 activity (van Leene et al., 2022). As discussed above, induction of *TPS8, TPS9*, *TPS10* and *TPS11* under C starvation may be due to falling Tre6P levels and might derepress SnRK1. The FLZ family proteins are also emerging as important modulators of SnRK1 signaling (Nietsch et al., 2014; 2016; Jamsheer and Lamxi, 2015; Jamsheer et al., 2015; 2018a; 2018b; 2022; Bortlik et al., 2022). Our analysis of their CRFs adds to the evidence that *FLZ* expression is strongly regulated by C status. Analysis of the response to induced overexpression of TPS pointed to Tre6P-inhibition of SnRK1 contributing to the regulation of a subset of *FLZs* that are induced in high sugar (Supplemental Figure S16, Supplemental text). It also pointed to other signaling pathways being involved in the regulation of *FLZs* whose expression increases in low-C conditions.

Overall, Tre6P signaling and SnRK1 function are closely intermeshed (Figure 9), with Tre6P acting not only to inhibit SnRK1 activity but also transcriptionally to control SnRK1 composition and the expression of two classes of proteins that interact with SnRK1. Overall, interactions with Tre6P may adjust not only SnRK1 activity *per se* but also its functionality to the prevailing conditions. In addition, increased SnRK1 expression dampens the response of Tre6P to sucrose (Peixoto et al., 2021).

### Interaction between Tre6P and other sugar-signaling pathways

We also explored interactions between Tre6P and further sugar-signaling pathways (Figure 10). A marked feature of the response to elevated Tre6P was the broad induction of ribosomal proteins and assembly factors (Figure 8C, Supplemental Figure S14H). TOR is a canonical positive regulator of ribosome biogenesis ((Sabatini, 2017; Ryabova et al., 2018; Wu et al., 2019; Meng et al., 2022; Scarpin et al., 2022). The most parsimonious explanation for the induction of ribosome biogenesis by Tre6P is that Tre6P acts via SnRK1 to regulate TOR activity. It has previously been observed that transient overexpression of SnRKα1 leads to repression of many ribosomal proteins and ribosome assembly factors (Baena-González et al. (2007); see also Figure 8C, Supplemental Figures 14H-I). Nukarinen et al. (2016) reported that SnRK1 phosphorylates the RAPTORB subunit of TOR and that loss of SnRK1α1 leads to increased phosphorylation of RPS6K, a canonical target of TOR.

To uncover other possible interactions between Tre6P and TOR, we analyzed the impact of elevated Tre6P on expression of TOR subunits and TOR phosphorylation targets (Supplemental Figure 15 and Supplemental text). Elevated Tre6P had no impact on expression of TOR subunits or the downstream kinases S6K1/2 and YAK1 (see Barrada et al., 2019; Forzani et al., 2019) but did induce two LARP kinases as well as NAP1;1 and RPS6. LARPs are involved in the TOR-LARP1-5’TOP signaling pathway that promotes expression of ribosomal proteins and assembly factors (Scarpin et al., 2020; 2022). NAP1;1 and RPS6 promote transcription of rRNA (Son et al., 2015) and their induction may contribute to the stimulation of ribosome biogenesis by elevated Tre6P. Other downstream TOR targets that were transcriptionally induced by elevated Tre6P included initiation and elongation factors (Supplemental Figure S15). As already mentioned, Tre6P repressed several members of the PYR/PYL family, which are phosphorylated and inhibited by TOR, pointing to concerted action of Tre6P and TOR signaling to inhibit ABA sensing and signaling in C-replete conditions. Reciprocal response to elevated Tre6P and transient overexpression of SnRK1a1 indicate that, in many cases, Tre6P may act via inhibition of SnRK1.

S1-type bZIP proteins like bZIP11 dimerize with S-type bZIPs to orchestrate C-starvation responses, and this action is inhibited in C-replete conditions because sucrose translationally inhibits their synthesis (Weise et al., 2004; Rahmani et al., 2009; Ma et al., 2011; Dröge-Laser and Weiste 2018, Pedrotti et al., 2018). There was considerable overlap between the iTPS response and the published response to bZIP11 overexpression (Ma et al., 2011) (Supplemental Figure 19). This was partly indirect, possibly because the decrease in sucrose allows translation of the bZIP11 protein. However, many bZIP11 targets were found in the set of transcripts assigned to CRF G1, which probably respond directly to elevated Tre6P. For most of the shared genes, the response to elevated Tre6P was opposite to the response to bZIP11 overexpression, and in many cases also opposite to the response to transient overexpression of SnRK1α1 (Supplemental Figure S19E). This observation implies that this subset of genes is regulated both by bZIP11 signaling and by Tre6P-SnRK1 signaling. It included many genes involved in starvation responses and also *TREHALASE1*, *BETA-AMYLASE9 (BAM9)*, a catalytically inactive β-amylase which together with BAM4 acts to accelerate the degradation of assimilatory starch in leaves during the night (David et al., 2021), and *NIGHT LIGHT-INDUCIBLE AND CLOCK-REGULATED2* (*LNK2*), a member of a small gene family that integrates light signaling with the circadian clock and is required for function of the dawn-phased REVEILLE4 (RVE4) and RVE8 (Xie et al., 2014). This dual layer of regulation presumably enhances responsiveness to low C. However, another set of genes showed qualitatively similar responses to elevated Tre6P and bZIP11 overexpression (Supplemental Figure S19E). This set of genes included the sucrose effluxer *SWEET12* and genes involved in gibberellic acid-, auxin- and jasmonate-signaling. In this case, dual but opposing regulation may stabilize expression or, alternatively, might allow changes in expression in conditions where SnRK1 activity and sucrose levels change independently of each other. Dual regulation of *SWEET12* expression by two sugar-signaling pathways might explain why elevated Tre6P sometime represses *SWEET12* (current study, Zhang et al., 2009) and sometimes induces *SWEET12* (Fichtner et al., 2021; Oszvald et al 2018).

### Tre6P as a component in a highly integrated signaling network that processes internal and external information

In conclusion, inducible overexpression of bacterial TPS leads to widespread changes in transcript abundance, with significant changes for almost half the genome, and >2-fold changes for about 5000 genes. Of these changes, about 40% are probably a response to elevated Tre6P but there is also a high proportion of indirect responses, probably including responses to the concomitant decline in sucrose. This complex transcriptional response mirrors the dual action of Tre6P as an upstream regulator of transcriptional responses and as a post-translational regulator of primary metabolism where it decreases sucrose synthesis and promotes sucrose utilization, with the resulting changes in metabolism leading to further layers of transcriptional responses. Many key aspects of metabolism and growth are transcriptionally regulated by signaling downstream of Tre6P, including repression of photosynthesis, changes in sucrose export, C/N interactions and enhancement of ribosome assembly and translation (Figures 9-10). There are also widespread interactions with other signaling pathways including light-signaling pathways, the circadian clock, and ABA-, auxin-, gibberellin-, brassinosteroid- and jasmonate-signaling. Mechanistically, our global analysis provides strong support for the idea that one function of Tre6P is to inhibit SnRK1 activity and starvation responses when C availability is high. Tre6P also acts via inhibition of SnRK1 to promote biosynthesis and growth in C-replete conditions, but in this case Tre6P probably also acts via additional pathways and/or the response is modulated by other factors. Our study also reveals that Tre6P exerts transcriptional control over the regulatory subunits of SnRK1 and sets of proteins that interact with SnRK1, and that Tre6P impacts on TOR signaling and S1/C bZIP signaling. It is known that over half the genes in Arabidopsis exhibit diel changes in transcript abundance, driven by changes in C availability and light- and circadian-signaling (Price et al., 2004; Bläsing et al., 2006; Usadel et al., 2007; Flis et al, 2016). Perturbation of Tre6P, which is just one component of the C-signaling network, suffices to generate equally large perturbations including crosstalk with circadian-, light- and hormone-signaling. The response to elevated Tre6P highlights, on the one hand, the sensitivity of signaling networks that plants use to integrate information about their internal metabolic status, the external conditions and the time-of-day, and, on the other hand, the robustness that is provided by multiple connections within this network. Deeper insights into this exquisite network will require integration of information about protein abundance and post-translational regulation, as well as strategies to gain the spatial resolution that is needed to understand how the network operates in different tissues and cells.

## Materials and methods

### Plant Material

Transgenic Arabidopsis (*Arabidopsis thaliana* (L.) Heyhn. accession Columbia-0) lines carrying the *p35S:AlcR*/*pAlcA:otsA* construct for ethanol-inducible of *Escherichia coli* TPS (otsA) (iTPS lines 29.2 and 31.3), and the AlcR empty vector control line were described in Martins et al. (2013) and Figueroa et al. (2016).

### Plant growth and ethanol Induction

The Arabidopsis iTPS lines were grown in two sets of experiments. For the ATH1 microarray analysis, Arabidopsis iTPS lines 29.2 and 31.3 and AlcR control lines were grown exactly as in Martins et al. (2013). Four-week-old plants were sprayed to runoff with water (mock induction control) or 2% (v/v) ethanol at dawn or dusk and harvested 12 h after spraying at the end of the day (ED) or at the end of the night (EN), respectively. Whole rosettes were excised *in situ* and immediately quenched in liquid nitrogen. Plants from one or two pots (i.e., five or 10 plants) were pooled to form one sample, ground to a fine powder at −70°C using a robotized ball mill (Stitt et al., 2007), and stored at −80°C until aliquots were extracted for transcript analysis. For the RNA-sequencing experiment, iTPS line 29.2 and AlcR seeds were sown on a 1:1 mixture of soil (Stender AG, Schermbeck, Germany; https://www.stender.de) and vermiculite in 6-cm diameter pots, covered and stratified at 4°C in the dark for 48 h before transfer to a controlled environment chamber (Percival E-36 L chamber model AR66-cL2-cLED fitted with white LED lighting, CLF Plant Climatics GmbH, Weringen, Germany; (https://www.percival-scientific.com9 with a 16-h photoperiod (160 µmol m^-2^ s^-1^ irradiance (white LEDs) and day/night temperature of 21°C/19°C. After germination, seedlings were transferred to 10-cm diameter pots (five seedlings per pot), and grown in the same conditions until 21 days after sowing (DAS). At 22 DAS, untreated control samples were harvested in the light immediately after dawn. The remaining plants were sprayed to run-off with either water (mock-induced control) or 2% (v/v) ethanol (induced) at 0.5 h after dawn and harvested 2h, 4h and 6h after spraying. Whole rosettes were harvested *in situ* and immediately quenched in liquid nitrogen. Four biological replicates were harvested per time point (each replicate contained four to five plants from one pot). Frozen plant material was ground to a fine powder using a robotized ball mill (Sulpice et al., 2014) and stored at −80°C until use.

### Extraction and measurement of metabolites

Metabolites were extracted with chloroform-methanol (Lunn et al., 2006). Tre6P, other phosphorylated intermediates and organic acids were quantified by anion-exchange high-performance liquid chromatography coupled to tandem mass spectrometry, as in Lunn et al. (2006), with modifications as in Figueroa et al. (2016). Soluble sugars were assayed enzymatically (Stitt et al., 1989). Starch was assayed enzymatically in the insoluble residue (Hendriks et al., 2003).

### RNA Extraction

Total RNA was extracted with an RNeasy Plant Mini-Kit (Qiagen, Hilden, Gemany; www.qiagen.com) according to manufacturer’s protocol. RNA concentration and purity were estimated by photometric analysis using a NanoDrop® ND-1000 UV-VIS spectrophotometer (Thermo Fisher Scientific Inc. Waltham, MA., USA; www.thermofisher.com). If necessary, RNA was diluted with water to a final concentration of 2 µg/µl. RNA integrity was confirmed by agarose (1% w/v) gel electrophoresis and by microfluidic electrophoresis using a Bioanalyzer 2100 (Agilent Technologies, Santa Clara, CA, USA:

www.agilent.com). DNA was removed by treatment with Turbo DNA-free™ DNase I enzyme (Applied Biosystems, Waltham, MA, USA; www.thermofisher.com) according to the manufacturer’s instructions. Absence of DNA contamination was confirmed by real-time PCR using intron-spanning primers for the *MADS AFFECTING FLOWERING 5* (*MAF5*; At5g65080) gene: *MAF5*-For: 5’-TTTTTTGCCCCCTTCGAATC-3’; *MAF5*-Rev: 5’-ATCTTCCGCCACCACATTGTAC-3’.

### Global gene expression profiling

For ATH1 arrays (Affymetrix ATH1 GeneChip® probe array) quality control and normalization of the microarray data were performed using Robin software (Lohse et al., 2010). To account for AlcR effects, the AlcR response was subtracted as follows. First, the difference in transcript abundance between water- and ethanol-treated plants was calculated separately for each genotype. Ethanol minus water differences were then plotted against those in the AlcR empty vector control samples. A simple linear model was generated with the ethanol minus water difference in the iTPS samples as the response variable, and the ethanol minus water difference in the AlcR empty vector samples as the predictive variable. The coefficient relating the two variables was then used to weight the data from the iTPS lines, thus normalizing them to minimize the background ethanol effect after the subtraction of the AlcR values. Differences remaining after this normalization and subtraction of the background ethanol effect could be ascribed to increased TPS expression and the resulting changes in Tre6P levels.

RNAseq analysis was performed on 2 µg of total RNA by BGI Genomics (Shenzhen, China; www.bgi.com) on quadruplicate replicates for each sample type. The service provided by BGI Genomics included library construction, sequencing, quality control and processing of the raw sequencing data (including barcode trimming and the removal of adaptor sequences, as well as of low-quality reads), generating at least 20 million paired-end reads (100 bp) for each sample. Gene mapping and statistical analysis were performed in-house using the CLC Genomics Workbench software (QIAGEN Aarhus A/S, www.qiagenbioinformatics.com) with the Araport10 and Araport11 genome release (https://www.arabidopsis.org; Cheng et al., 2016) being used for gene annotation. Expression values (RPKM; reads per kilobase of transcript per million reads mapped) were corrected for differences in library size. The CLC Genomics Workbench was used for differential gene expression analysis and FC calculations using the corrected RPKM values of ethanol *v*. water-sprayed samples.

After applying an FDR<0.05 and a FC≥2 cutoff only a small fraction of genes exhibited significant change in the same direction in the iTPS and in the alcR control (22 out of 5618 DEGs at 4h, 12 DEGs out of 5437 DEGs at 6h) after ethanol induction. These DEGs were highlighted in the full DEG list and no further normalization was carried out (see Supplemental Dataset S4). In data analyses in which a more relaxed filter (FDR<0.05, FC≥0.2) was applied to the iTPS data, more genes changed in the same direction in iTPS29.2 and alcR after ethanol induction. These genes were omitted from further analyses.

### Statistical Analysis

Technical replicates were always averaged to generate a single value for each biological replicate. The statistical analysis was restricted to biological replicates and performed using Sigma-Plot 14.5 software (Systat Software GmbH, Düsseldorf, Germany; http://www.systat.de). Significance of changes in metabolite levels was tested by one-way ANOVA using a pairwise multiple comparison procedure, with post-hoc testing by the Holm-Sidak method (p<0.05), as described in each figure legend.

### PageMan analyses

Data were analyzed with the PageMan tool (Usadel et al., 2006) using MapMan software (Thimm et al., 2004; Usadel et al., 2006; version 3.6.0RC1; https://mapman.gabipd.org/) using log2FC values and mapping assignments of Ath<u>_</u>AGI<u>_</u>LOCUS<u>_</u>TAIR10<u>_</u>Aug2012. The heat maps show the average changes in all transcripts in a given BIN or sub-BIN. Only significant changes (by CRF criteria, (FDR<0.05, FC≥0.2)) were retained and non-significant genes were assigned a zero value, before averaging across all genes in the BIN or sub-BIN.

### Gene Ontology analysis

Gene Ontology enrichment analyses were performed for significant responses only using the GO database (http://geneontology.org/), version PANTHER17.0 (release date 7^th^ February 2021).

### STRING analyses

The STRING (search tool for recurring instances of neighboring genes; Snel et al., 2000; Szklarczyk et al., 2022; http://www.bork.embl-heidelberg.de/STRING) tool was used for enrichment analysis of DEGs.

## Accession numbers

Microarray data from this article have been deposited in Gene Expression Omnibus database (http://www.ncbi.nlm.nih.gov/geo) under accession number E-MTAB-12783.

## Supporting information

Combined Supplemental Figures and Tables

## Acknowledgements

We thank Ursula Krause for skilled technical help

## Author contributions

OA, JEL and MS conceived the study. MCMM performed the ATH1 microarray experiments, ML and FMG processed the raw data and MCMM and MS analyzed the data. OA performed the RNAseq experiments and processed the raw data, AS assisted with RNA extraction, quality and integrity checks and processing, OA and RF performed the metabolite analysis. OA and MS analyzed the data and drafted the manuscript.

## Supplemental Data

**Supplemental Figure S1.** Comparison of changes in transcript abundance 12 h after induction of TPS in two independent iTPS lines, 29.2 and 31.3

**Supplemental Figure S2.** Estimation of the carbon response factor, CRF

**Supplemental Figure S3.** Comparison of the carbon response factor (CRF) and iTPS in the ED and EN treatments

**Supplemental Figure S4.** Relation between Tre6P and sucrose, and changes of further metabolites after induction in iTPS29.2 for 4 or 6-h in the light (supplemental to Fig. 1)

**Supplemental Figure S5.** Global analysis of the response of transcript abundance at 4-h and 6-h after induction in iTPS29.1 (supplemental to Figure 2)

**Supplemental Figure S6.** Changes in expression of selected G0 genes in response to various treatments by which CRF was determined, and abundance of transcripts of genes assigned to G0 Supplemental Figure S7. Visualization of selected responses in metabolism using PageMan displays (supplemental to Figure 3)

**Supplemental Figure S8.** Responses of transcripts encoding proteins involved in sucrose transport and metabolism

**Supplemental Figure S9.** Responses of selected transcripts encoding proteins involved in the regulation of floral induction

**Supplemental Figure S10.** Responses of transcripts encoding proteins involved in the circadian clock.

**Supplemental Figure S11.** Enrichment analyses using Gene Ontology.

**Supplemental Figure S12.** Comparison of the changes in transcript abundance after induced increase in Tre6P, and in Arabidopsis lines that constitutively express bacterial TPS (Supplemental to Figure 6 and Supplemental Table S5.

**Supplemental Figure S13.** Responses of transcripts encoding proteins involved in Tre6P metabolism.

**Supplemental Figure S14.** Responses of transcripts encoding SnRK1 subunits, or implicated as downstream transcriptional targets of SnRK1 signaling.

**Supplemental Figure S15.** TOR subunits, higher levels regulators of ribosome biogenesis and other TOR targets including ABA receptors.

**Supplemental Figure S16.** FCS-LIKE ZINC FINGER (FLZ) family proteins.

**Supplemental Figure S17.** Brassinosteroid signaling.

**Supplemental Figure S18.** Cell wall modification

**Supplemental Figure S19.** Comparison of iTPS with constitutive overexpression of bZIP11. Supplemental Figure S20. Transcription factors that respond to an induced increase in Tre6P levels Supplemental Figure S21. Analysis of transcription factors assigned to CRF group G1.

**Supplemental Figure S22.** C and S1 bZIP transcription factors

**Supplemental Figure S23.** Transcript for TFs that were assigned to CRF group G1 in the iTPS response: comparison of their iTPS response and in their oeAKIN10 response.

**Supplemental Figure S24.** Analysis of transcription factors assigned to CRF group G2.

**Supplemental Figure S25.** Glucosinolate metabolism and its transcriptional regulation

**Supplemental Figure S26.** Proposed interplay between Tre6P-mediated signaling and other C- signaling pathways in wild-type plants (supplemental to Figures 5 and 7)

**Supplemental Table S1.** List of transcriptome datasets used in this study

**Supplemental Table S2.** Changes in transcript abundance 12 h after induction of TPS, investigated at ED or EN (extends Supplemental Figures S1 and S3)

**Supplemental Table S3.** Summary of the number of differentially expressed genes, their assignment to CRF groups and the R2, direction and p-value of the relationship between the change in transcript abundance and the CRF value (extends Figure 2 and Supplemental Figure S5A-B)

**Supplemental Table S4.** Pairwise comparison of responses after 4 h and 6 h of induction in the light, and in the ED and EN treatments (extends Supplemental Figure S5C)

**Supplemental Table S5.** Comparison of the responses to constitutive overexpression of TPS (oeTPS) and induced overexpression of TPS (iTPS) (extends Figure 6 and Supplemental Figure S12).

**Supplemental Table S6.** Comparison of the tSnRK1α1 and iTPS responses (extends Figure 8 and Supplemental Figure S14)

**Supplemental Dataset S1.** ATH1 profiling of ED and EN treatments (underlies Supplemental Figure S1 and Supplemental Tables S2 and S4)

**Supplemental Dataset S2.** Calculation of carbon response factor (underlies Supplemental Figure S2)

**Supplemental Dataset S3.** Metabolites iTPS 4-6h (underlies Figure 1 and Supplemental Figure S3)

**Supplemental Dataset S4.** RNA seq data (underlies Figures 2-8, Supplementary Figures S5-S14, and Supplementary Tables S2-S6)

**Supplemental Dataset S5.** GO analysis of RNAseq dataset (extends Supplemental Figure S11)

**Supplemental Dataset S6.** Robust set of Tre6P-regulated genes (underlies Figure 6, Supplemental Figure S12 and Supplemental Table S5).

**Supplemental Dataset S7.** GO analysis (underlies Figure 6, Supplemental Figure S12 and Supplemental Table S6).

**Supplemental Dataset S8.** Transcription factors (underlies Supplemental Figures S20-24)

## References

Allemeersch J, Durinck S, Vanderhaeghen R, Alard P, Maes R, Seeuws K, Bogaert T, Coddens K, Deschouwer K, Van Hummelen P, et al. Benchmarking the CATMA microarray. A novel tool for Arabidopsis transcriptome analysis. Plant Physiol 2005:137:588–601

Annunziata MG, Apelt F, Carillo P, Krause U, Feil R, Mengin V, Lauxmann MA, Köhl K, Nikoloski Z, Stitt M, Lunn JE. Getting back to nature: a reality check for experiments in controlled environments. J Exp Bot 2017:68:4463–4477

Avidan O, Moraes TA, Mengin VA, Feil R, Rolland F, Stitt M, Lunn JE. *In-vivo* protein kinase activity of SnRK1 fluctuates in Arabidopsis rosettes during light-dark cycles. Plant Physiol 2023:192:387–408 10.1093/plphys/kiad066

Avonce N, Leyman B, Mascorro-Gallardo JO, Van Dijck P, Thevelein JM, Iturriaga G. The Arabidopsis trehalose-6-P synthase *AtTPS1* gene is a regulator of glucose, abscisic acid, and stress signaling. Plant Physiol 2004:136:3649–3655

Baena-González E, Rolland F, Thevelein JM, Sheen J. A central integrator of transcription networks in plant stress and energy signaling. Nature 2007:448:938–942

Baena-González E, Hanson J. Shaping plant development through the SnRK1–TOR metabolic regulators. Curr Opin Plant Biol 2017:35:152–155

Baena-González E, Lunn JE. SnRK1 and trehalose 6-phosphate – two ancient pathways converge to regulate plant metabolism and growth. Curr Opin Plant Biol 2020:55:52–59

Barrada A, Djendli M, Desnos T, Mercier R, Robaglia C, Montané MH, Menand B. A TOR–YAK1 signaling axis controls cell cycle, meristem activity and plant growth in Arabidopsis. Development 2019:146: dev171298

Belda-Palazón B, Adamo M, Valerio C, Ferreira LJ, Confraria A, Reis-Barata D, Rodrigues A, Meyer C, Rodriguez PL, Baena-González E. A dual function of SnRK2 kinases in the regulation of SnRK1 and plant growth. Nature Plants 2020:6:1345–1353

Belda-Palazón B, Costa M, Beeckman T, Rolland F, Baena-González E. ABA represses TOR and root meristem activity through nuclear exit of the SnRK1 kinase. Proc Natl Acad Sci USA 2022:19:e2204862119

Benjamini Y, Hochberg Y. Controlling the false discovery rate: a practical and powerful approach to multiple testing. J Royal Stat Soc Ser B. 1995:5:289–300

Bernard SM, Habash DZ. The importance of cytosolic glutamine synthetase in nitrogen assimilation and recycling. New Phytol 2009:182(3):608–320

Biemelt S, Sonnewald U-Plant-microbe interactions to probe regulation of plant carbon metabolism. J Plant Physiol. 2006:163(3):307–318 doi: 10.1016/j.jplph.2005.10.011.

Bläsing O, Gibon Y, Günther M, Höhne M, Osuna D, Thimm T, Scheible W-R, Morcuende R, Stitt M. Sugars and circadian regulation make major contributions to the global regulation of diurnal gene expression in Arabidopsis. Plant Cell 2005:17:3257–3281

Bledsoe SW, Henry C, Griffiths CA, Paul MJ, Feil R, Lunn JE, Stitt M, Lagrimini LM. The role of Tre6P and SnRK1 in maize early kernel development and events leading to stress-induced kernel abortion. BMC Plant Biol 2017:17:1–17

Bortlik J, Jost Lühle J, Alseekh S, Weiste C, Fernie AR, Dröge-Laser W, Börnke F. DUF581-9 (At2g44670; FLZ3) negatively regulates SnRK1 activity by interference with T-loop phosphorylation. bioRxiv 2022: 10.1101/2022.03.17.484690

Braun DM. Phloem loading and unloading of sucrose: what a long, strange trip from source to sink. Annu Rev Plant Biol 2022:73:553–58

Broeckx T, Hulsmans S, Rolland F. The plant energy sensor: evolutionary conservation and divergence of SnRK1 structure, regulation, and function. J Exp Bot 2016:67:6215–6252. doi: 10.1093/jxb/erw416.

Cabib E, Leloir LF. The biosynthesis of trehalose phosphate. J Biol Chem. 1958:231(1):259–275

Carillo P, Feil R, Gibon Y, Satoh-Nagasawa N, Jackson D, Bläsing OE, Stitt M, Lunn JE. A fluorometric assay for trehalose in the picomole range. Plant Methods 2013:9:21

Cheng C-Y, Krishnakumar V, Chan AP, Thibaud-Nissen F, Schobel S, Town CD. Araport11: a complete reannotation of the *Arabidopsis thaliana* reference genome. Plant J 2017:89:789–804 10.1111/tpj.13415

Claeys H, Vi S, Xu X, Satoh-Nagasawa N, Eveland A, Goldshmidt A, Feil R, Beggs G, Sakai H, Brennan R, Lunn JE, Jackson D. Control of meristem determinacy by trehalose-6-phosphate phosphatases is uncoupled from enzymatic activity. Nature Plants 2019:5:e352.

Coello P, Martínez-Barajas E. The activity of SnRK1 is increased in *Phaseolus vulgaris* seeds in response to a reduced nutrient supply. Front Plant Sci 2014:5:1–7

Cookson S. Yadav UP, Klie S, Morcuende RM, Usadel, B, Lunn JE, Stitt, M. Temporal kinetics of the transcriptional response to carbon depletion and sucrose readdition in Arabidopsis seedlings. Plant Cell Environ 2016:39:768–786

Coruzzi GM. Primary N-assimilation into Amino Acids in Arabidopsis. Arabidopsis Book, 2003:2: e0010

Cosgrove DJ. Growth of the plant cell wall. Nat Rev Mol Cell Biol 2005:6(11):850–861 doi: 10.1038/nrm1746

Crepin N, Rolland F. SnRK1 activation, signaling, and networking for energy homeostasis. Curr Opin Plant Biol 2019:51:29–36

Crozet P, MargalhaL, Confraria A, Rodrigues A, Martinho C, Adamo M, EliasCA, Baena-González E. Mechanisms of regulation of SNF1/AMPK/SnRK1 protein kinases. Front. Plant Sci 2014:5(190): doi 10.3389/fpls.2014.00190

Czedik-Eysenberg A, Arrivault, Lohse M, Feil R, Krohn N, Encke B, Szecowka M, Nunes-Nesi A, Fernie AR, Lunn JE, et al. The interplay between carbon availability and growth in different zones of the growing maize leaf. Plant Physiol 2016:172:943–967

Dai ZW, Léon C, Feil R, Lunn JE, Delrot S, Gomès E. Metabolic profiling reveals coordinated switches in primary carbohydrate metabolism in grape berry (*Vitis vinifera* L.), a non-climacteric fleshy fruit. J Exp Bot 2013:64:1345–1355

David DC, Lee S-K, Bruderer E, Abt M, Fischer-Stettler M, Tschopp M.A, Solhaug E, Sanchez K, Zeeman SC. BETA-AMYLASE9 is a plastidial non-enzymatic regulator of leaf starch degradation. Plant Physiol 2021:188:191–207 DOI:10.1093/plphys/kiab468

Dalchau N, Baek SJ, Briggs HM, Robertson FC, Dodd AN, Gardner MJ, Stancombe MA, Haydon MJ, Stan G-B, Goncalves JM, et al. The circadian oscillator gene GIGANTEA mediates a long-term response of the *Arabidopsis thaliana* circadian clock to sucrose. Proc Natl Acad Sci USA 2011:108:5104–5109

Debast S, Nunes-Nesi A, Hajirezaei MR, Hofmann J, Sonnewald U, Fernie AR, Börnke F. Altering trehalose-6-phosphate content in transgenic potato tubers affects tuber growth and alters responsiveness to hormones during sprouting. Plant Physiol 2011:156:1754–1771

de Haro LA, Arellano SM, Novák O, Feil R, Dumón AD, Mattio MF, Tarkowská D, Llauger G, Strnad M, Lunn JE, Pearce S, Figueroa CM, Del Vas M. Mal de Río Cuarto virus infection causes hormone imbalance and sugar accumulation in wheat leaves. BMC Plant Biol 2019:19(1):112 doi: 10.1186/s12870-019-1709-y

Delatte TL, Sedijani P, Kondou Y, Matsui M, de Jong GJ, Somsen GW, Wiese-Klinkenberg A, Primavesi LF, Paul MJ, Schluepmann H. Growth arrest by trehalose-6-phosphate: an astonishing case of primary metabolite control over growth by way of the SnRK1 signaling pathway. Plant Physiol 2011:157:160– 174

Delorge I, Figueroa CM, Feil R, Lunn JE, Van Dijck P. Trehalose-6-phosphate synthase 1 is not the only active TPS in *Arabidopsis thaliana*. Biochem J 2015:466:283–290

Doehlemann G, Wahl R, Horst RJ, Voll LM, Usadel B, Poree F, Stitt M, Pons-Kühnemann J, Sonnewald U, Kahmann R, Kämper J. Reprogramming a maize plant: transcriptional and metabolic changes induced by the fungal biotroph *Ustilago maydis*. Plant J. 2008:56(2):181–195 doi: 10.1111/j.1365-313X.2008.03590.x.

Dombrecht B, Xue GP, Sprague SJ, Kirkegaard JA, Ross JJ, Reid JB, Fitt GP, Sewelam N, Schenk PM, Manners JM, Kazan K.. MYC2 differentially modulates diverse jasmonate-dependent functions in Arabidopsis. Plant Cell 2007:19:2225–2245

dos Anjos L, Pandey PK, Moraes TA, Feil R, Lunn JE, Stitt M. Feedback regulation by trehalose 6-phosphate slows down starch mobilization below the rate that would exhaust starch reserves at dawn in Arabidopsis leaves. Plant Direct 2018:2:e00078

Dröge-Laser W, Weiste C. The C/S1 bZIP network: A regulatory hub orchestrating plant energy homeostasis. Trends Plant Sci 2018:23:36–49 10.1016/j.tplants.2018.02.003

Dröge-Laser W, Snoek BJ, Snel B, Weiste C. The Arabidopsis bZIP transcription factor family — an update. Curr Opin Plant Biol 2018:45:36–49

Eastmond PJ, Van Dijken AJ, Spielman M, Kerr A, Tissier AF, et al. Trehalose-6-phosphate synthase 1, which catalyses the first step in trehalose synthesis, is essential for Arabidopsis embryo maturation. Plant J. 2002:29:225–35 2002

Emanuelle S, Hossain MI, Moller IE, Pedersen HL, van de Meene AM, Doblin MS, Koay A, Oakhill JS, Scott JW, Willats WG, et al. SnRK1 from *Arabidopsis thaliana* is an atypical AMPK. Plant J 2015:82: 183–192

Fedosejevs ET, Feil F, Lunn JE, Plaxton WC. The signal metabolite trehalose-6-phosphate inhibits the sucrolytic activity of sucrose synthase from developing castor beans. FEBS Letts 2018:592: 2525–2532

Fernández-Calvo P, Chini A, Fernández-Barbero G, Chico J-M, Gimenez-Ibanez S, Geerinck J, Eeckhout D, Schweizer F, Godoy M, Franco-Zorrilla JM, et al. The Arabidopsis bHLH transcription factors MYC3 and MYC4 are targets of JAZ repressors and act additively with MYC2 in the activation of jasmonate responses. Plant Cell 2011:23:701–715

Fichtner F, Barbier FF, Feil R, Watanabe M, Annunziata MG, Stitt M, Beveridge CA, Lunn JE. Trehalose 6-phosphate is involved in triggering axillary bud outgrowth in garden pea (*Pisum sativum* L.). Plant J 2017:92: 611–23

Fichtner F, Olas JJ, Feil R, Watanabe M, Krause U, Hoefgen R, Stitt M, Lunn JE. Functional features of TREHALOSE-6-PHOSPHATE SYNTHASE1, an essential enzyme in Arabidopsis. Plant Cell 2020:32:1949– 72

Fichtner F, Barbier FF, Annunziata MG, Feil R, Olas JJ, Mueller-Roeber B, Stitt M, Beveridge A, Lunn JE. Regulation of shoot branching in Arabidopsis by trehalose 6-phosphate. New Phytol 2021:229: 2135–2151

Fichtner F, Lunn JE. The Role of trehalose 6-phosphate (Tre6P) in plant metabolism and development. Annu Rev Plant Biol 2021:72: 737–760

Figueroa CM, Feil R, Ishihara H, Watanabe M, Kölling K, Krause U, Höhne M, Encke B, Plaxton WC, Zeeman SC, et al. Trehalose 6-phosphate coordinates organic and amino acid metabolism with carbon availability. Plant J 2016:85: 410–442

Figueroa CM, Lunn JE. A tale of two sugars: trehalose 6-phosphate and sucrose. Plant Physiol 2016:172:7–27

Flis A, Sulpice R, Abel C, Ivakov A, Liput M, Millar AJ, Stitt M. Photoperiod-dependent changes in the phase of core clock transcripts, and in global transcriptional outputs at dusk and dawn in Arabidopsis. Plant Cell Environ 2016:39:1955–1981

Fu Y, Lim S, Urano D, Tunc-Ozdemir M, Phan NG, Elston TC, Jones AM. Reciprocal encoding of signal intensity and duration in a glucose-sensing circuit. Cell 2014:156(5):1084–1095 doi: 10.1016/j.cell.2014.01.013

Forzani C, Duarte GT, Van Leene J, Clément G, Huguet S, Paysant-Le-Roux C, Mercier R, De Jaeger G, Leprince A-S, Meyer C. Mutations of the AtYAK1 kinase suppress TOR deficiency in Arabidopsis. Cell Reports 2019:27:3696–3708

Frank A, Matiolli CC, Viana JC, Hearn TJ, Kusakina J, Belbin FE, Wells Newman D, Yochikawa A, Cano-Ramirez DL, Chembath A et al. Circadian entrainment in Arabidopsis by the sugar-responsive transcription factor bZIP63. Current Biol 2018:28:2597–2606

Gangappa SN, Botto JF. The multifaceted roles of HY5 in plant growth and development. Mol Plant 2016:9:1353–1365.

Gibon Y, Vigeolas H, Tiessen A, Geigenberger P, Stitt M. Sensitive and high throughput metabolite assays for inorganic pyrophosphate, ADPGlc, nucleotide phosphates, and glycolytic intermediates based on a novel enzymic cycling system. Plant J 2002:30:221–235

Gibon Y, Bläsing OE, Palacios-Rojas N, Pankovic D, Hendriks JHM, Fisahn J, Höhne M, Günther M, Stitt M. Adjustment of diurnal starch turnover to short days: depletion of sugar during the night leads to a temporary inhibition of carbohydrate utilization, accumulation of sugars and post-translational activation of ADP-glucose pyrophosphorylase in the following light period. Plant J 2004:39:847–862

Glab N, Oury C, Guérinier T, Domenichini S, Crozet P, Thomas M, Vidal J, Hodges M. The impact of *Arabidopsis thaliana* SNF1-related-kinase 1 (SnRK1)-activating kinase 1 (SnAK1) and SnAK2 on SnRK1 phosphorylation status: characterization of a SnAK double mutant. Plant J. 2017:89(5):1031–1041

Goddijn OJM, van Dun K. Trehalose metabolism in plants. Trends Plant Sci. 1999:4(8):315–319

Gómez LD, Gilday A, Feil R, Lunn JE, Graham IA. AtTPS1-mediated trehalose 6-phosphate synthesis is essential for embryogenic and vegetative growth and responsiveness to ABA in germinating seeds and stomatal guard cells. Plant J 2010:64:1–13

Griffiths CA, Sagar R, Geng Y, Primavesi LF, Patel MK, Passarelli MK, Gilmore IS, Steven RT, Bunch J, Paul MJ, et al. Chemical intervention in plant sugar signaling increases yield and resilience. Nature 2016:540:574–578

Gutierrez-Beltran E, Crespo JL. Compartmentalization, a key mechanism controlling the multitasking role of the SnRK1 complex. J Exp Bot 2022:73:7055–7067

Hanson J, Hansson M, Wiese A, Hendricks MMWB, Smeekens S. The sucrose regulated transcription factor bZIP11 affects amino acid metabolism by regulating the expression of ASPARAGINE SYNTHETASE1 and PROLINE DEHYDROGENASE2. Plant J 2008:53:935–994

Harthill JE, Meek SEM, Morrice N, Peggie MW, Borch J, Wong BHC, Mackintosh C. Phosphorylation and 14-3-3 binding of Arabidopsis trehalose-phosphate synthase 5 in response to 2-deoxyglucose. Plant J. 2006:47(2):211–223

Haydon MJ, Mielczarek O, Robertson FC, Hubbard KE, Webb AAR. Photosynthetic entrainment of the Arabidopsis thaliana circadian clock. Nature 2103:502:689–692

Haydon MJ, Mielczarek O, Frank A, Román Á, Webb AA. Sucrose and ethylene signaling interact to modulate the circadian clock. Plant Physiol 2017:175:947–958

Henry C, Bledsoe SW, Siekman A, Kollman A, Waters BM, Feil R, Stitt M, Lagramini LM. The trehalose pathway in maize: conservation and gene regulation in response to the diurnal cycle and extended darkness. J Exp Bot 2014:65:5959–5973

Henry C, Bledsoe SW, Griffiths CA, Kollman A, Paul MJ, Sakr S, Lagrimini LM. Differential role for trehalose metabolism in salt-stressed maize. Plant Physiol 2015:169:1072–1089

Hulsmans S, Rodriguez M, de Coninck B, Rolland F. The SnRK1 energy sensor in plant biotic interactions. Trends Plant Sci 2016:21: 648–661

Hwang G, Kim S, Cho J-Y, Paik I, Kim J-I, Oh E. Trehalose-6-phosphate signaling regulates thermoresponsive hypocotyl growth in *Arabidopsis thaliana*. EMBO Rep. 2019:20(10):47828

Ishihara H, Moraes T, Pyl E-T, Schulze WX, Obata T, Scheffel A, Fernie AR, Sulpice R, and Stitt M. Growth rate correlates negatively with protein turnover in Arabidopsis accessions. Plant J 2017:91: 416–429

Ishihara H, Alseekh S, Feil R, Perera P, George GM, Niedźwiecki P, Arrivault S, Zeeman SC, Fernie AR, Lunn JE, et al. Rising rates of starch degradation with time in the light and trehalose 6-phosphate levels optimize carbon availability and thereby buffer growth. Plant Physiol 2022:189:1976–2000

Jamsheer, KM, Laxmi A. Expression of Arabidopsis FCS-Like Zinc finger genes is differentially regulated by sugars, cellular energy level, and abiotic stress. Front Plant Sci 2015:6:746 10.3389/fpls.2015.00746

Jamsheer KM, Mannully CT, Laxmi A. Comprehensive evolutionary and expression analysis of FCS-Like Zinc finger gene family yields insights into their origin, expansion and divergence. PLOS One 2015:10: e0134328

Jamsheer LM, Sharma M, Laxmi A. FCS-like zinc finger 6 and 10 repress SnRK1 signaling in Arabidopsis. Plant J. 2018a:94:232–245

Jamsheer KM, Shukla BN, Jindal S, Gopan N, Mannully CT, Laxmi A. The FCS-like zinc finger scaffold of the kinase SnRK1 is formed by the coordinated actions of the FLZ domain and intrinsically disordered regions. J Biol Chem 2018b:293:13134–13150

Jamsheer KM, Jindal S, Laxmi A. A negative feedback loop of TOR signaling balances growth and stress-response trade-offs in plants. Cell Reports 2022:39:110631

Johansson M, Staiger D. Time to flower: interplay between photoperiod and the circadian clock. J Exp Bot 2014:66(3):719–730

Jossier M, Bouly JP, Meimoun P, Arjmand A, Lessard P, Hawley S, Hardie GD, Thomas M. SnRK1 (SNF1-related kinase 1) has a central role in sugar and ABA signaling in *Arabidopsis thaliana*. Plant J 2009:59:316–328

Juntawong P, Bailey-Serres J. Dynamic light regulation of translation status in *Arabidopsis thaliana*. Front Plant Sci 2012:3:66 doi: 10.3389/fpls.2012.00066.

Juntawong P, Girke T, Bazin J, Baily-Serres J. Translational dynamics revealed by genome-wide profiling of ribosome footprints in Arabidopsis. Proc Natl Acad Sci USA 2014:111:E203–212

Kaewthai N, Gendre D, Eklöf JM, Ibatullin FM, Ezcurra I, Bhalerao RP, Brumer H. Group III-A XTH genes of Arabidopsis encode predominant xyloglucan endohydrolases that are dispensable for normal growth. Plant Physiol 2013:161(1):440–54 doi: 10.1104/pp.112.207308.

Klein H, Gallagher J, Demesa-Arevalo E, Juarez MJA, Heeney M, Feil R, Lunn JE, Xiao Y, Chuck G, Whipple C, et al. Recruitment of an ancient branching program to suppress carpel development in maize flowers. Proc Natl Acad Sci USA 2022:119: e2115871119

Krapp, A, Hofmann B, Schäfer C, Stitt M. Regulation of the expression of rbcS and other photosynthetic genes by carbohydrates: a mechanism for the “sink-regulation” of photosynthesis? Plan J. 1993:3:817–828

Krapp A. Stitt M. An evaluation of direct and indirect mechanism for the “sink”-regulation of photosynthesis in spinach: changes in gas exchange, carbohydrates, metabolites, enzyme activities and steady state transcript levels after cold girdling source leaves Planta. 1995:195:313–323

Kretzschmar T, Pelayo MAF, Trijatmiko KR, Gabunada LF, Alam R, Jimenez R, Mendioro MS, Slamet-Loedin IH, Sreenivasulu N, Bailey-Serres J, et al. A trehalose-6-phosphate phosphatase enhances anaerobic germination tolerance in rice. Nat Plants. 2015:1(9):15124

Lastdrager J, Hanson J, Smeekens S. Sugar signals and the control of plant growth and development. J Exp Bot 2014:65:799–807

Leyman B, van Dijck P, Thevelein JM. An unexpected plethora of trehalose biosynthesis genes in *Arabidopsis thaliana*. Trends Plant Sci. 2001:6(11):510–513

Li L, Liu KH, Sheen J. Dynamic nutrient signaling networks in plants. Annu Rev Cell Dev Biol 2021:37: 341–367 doi: 10.1146/annurev-cellbio-010521-015047

Liao CY, Pu YT, Nolan TM, Montes C, Guo HQ, Walley JW, Yin YH, Bassham DC. Brassinosteroids modulate autophagy through phosphorylation of RAPTOR1B by the GSK3-like kinase BIN2 in Arabidopsis. Autophagy 2022:19(4): 1293–1310 DOI 10.1080/15548627.2022.2124501

Lohse M, Bolger A, Nagel A, Fernie AR, Lunn JE, Stitt M, Usadel B. RobiNA: A user-friendly, integrated software solution for RNA-Seq based transcriptomics. Nucl Acids Re. 2012:40: W622–W627. DOI: 10.1093/nar/gks540

Lunn JE. Gene families and evolution of trehalose metabolism in plants. Funct Plant Biol 2007:34(6):550–563 doi: 10.1071/FP06315

Lunn JE, Feil R, Hendriks JHM, Gibon Y, Morcuende R, Osuna D, Scheible WR, Carillo P, Hajirezaei MR, Stitt M. Sugar-induced increases in trehalose 6-phosphate are correlated with redox activation of ADPglucose pyrophosphorylase and higher rates of starch synthesis in *Arabidopsis thaliana*. Biochem J 2006:148:139–148

Lunn JE, Delorge I, Mar C, van Dijck P, Stitt M. Trehalose metabolism in plants. Plant J 2014:4:544– 567

Ma J, Hanssen M, Lundgren K, Hernández L, Delatte T, Ehlert A, Liu CM, Schluepmann H, Dröge-Laser W, Moritz T, et al. The sucrose-regulated Arabidopsis transcription factor bZIP11 reprograms metabolism and regulates trehalose metabolism. New Phytol 2011:191:733–745

Mair A, Pedrotti L, Wurzinger B, Anrather D, Simeunovic A, Weiste C, Valerio C, Dietrich K, Kirchler T, Nägele T, et al. SnRK1-triggered switch of bZIP63 dimerization mediates the low-energy response in plants. ELife 2015:4:05828

Martínez-Barajas E, Delatte T, Schluepmann H, de Jong GJ, Somsen GW, Nunes C, Primavesi LF, Coello P, Mitchell RA, Paul MJ. Wheat grain development is characterized by remarkable trehalose 6-phosphate accumulation pregrain filling: tissue distribution and relationship to SNF1-related protein kinase1 activity. Plant Physiol 2011:156:373–381

Martins MCM, Hejazi M, Fettke J, Steup M, Feil R, Krause U, Arrivault S, Vosloh D, Figueroa CM, Ivakov A, et al. Feedback inhibition of starch degradation in arabidopsis leaves mediated by trehalose 6-phosphate. Plant Physiol 2013:163:1142–1163

Meitzel T, Radchuk R, McAdam EL, Thormählen I, Feil R, et al. Trehalose 6-phosphate promotes seed filling by activating auxin biosynthesis. New Phytol. 2021:229:1553–65

Meng Y, Zhang N, Li J, Shen X, Sheen J, Xiong Y. TOR kinase, a GPS in the complex nutrient and hormonal signaling networks to guide plant growth and development. J Exp Bot 2022:73(20):7041–7054 doi: 10.1093/jxb/erac282

Mitreiter S, Gigolashvili T. Regulation of glucosinolate biosynthesis. J Exp Bot 2021:72:70–91

Moraes TA, Mengin V, Annunziata MG, Encke B, Krohn N, Höhne M, Stitt M. Response of the circadian clock and diel starch turnover to one day of low light or low CO2. Plant Physiol 2019:179: 1457–1478

Moreau M, Azzopardi M, Clément G, Dobrenel T, Marchive C, Renne C, Martin-Magniete M-L, Taconnat L, Renou J-P, Robaglia C, et al. Mutations in the Arabidopsis homolog of LST8/GβL, a partner of the target of rapamycin kinase, impair plant growth, flowering, and metabolic adaptation to long days. Plant Cell 2012:24:463–81

Nelson CJ, Millar AH. Protein turnover in plant biology. Nature Plants 2015:115017

Nietzsche M, Landgraf R, Tohge T, Börnke F. A protein-protein interaction network linking the energy-sensor kinase SnRK1 to multiple signaling pathways in *Arabidopsis thaliana*. Curr Opin Plant Biol 2016:5:36–44

Nietzsche M, Schießl I, Börnke F, Griffiths CA. The complex becomes more complex: protein-protein interactions of SnRK1 with DUF581 family proteins provide a framework for cell- and stimulus type-specific SnRK1 signaling in plants. Front Plant Sci 2014:5:1–13

Nuccio ML, Wu J, Mowers R, Zhou HP, Meghji M, Primavesi LF, Paul MJ, Chen X, Gao Y, Haque E, et al. Expression of trehalose-6-phosphate phosphatase in maize ears improves yield in well-watered and drought conditions. Nat Biotech 2015:33:862–869

Nukarinen E, Nägele T, Pedrotti L, Wurzinger B, Baena-González E, Dröge-Laser W, Weckwerth W. Quantitative phosphoproteomics reveals the role of the AMPK plant ortholog SnRK1 as a metabolic master regulator under energy deprivation. Sci Rep 2016:6:31697

Nunes C, O’Hara LE, Primavesi LF, Delatte TL, Schluepmann H, Somsen GW, Silva AB, Fevereiro PS, Wingler A, Paul MJ. The trehalose 6-phosphate/SnRK1 signaling pathway primes growth recovery following relief of sink limitation. Plant Physiol 2013a:162: 1720–1732

Nunes C, Primavesi LF, Patel MK, Martinez-Barajas E, Powers SJ, Sagar R, Fevereiro PS, Davis BG, Paul MJ. Inhibition of SnRK1 by metabolites: Tissue-dependent effects and cooperative inhibition by glucose 1-phosphate in combination with trehalose 6-phosphate. Plant Physiol Biochem 2013b:63: 89–98

Osuna D, Usadel B, Morcuende R, Scheible W-R, Gibon Y, Bläsing O-E, Thimm O, Höhne M, Günter M, Udvardi MK et al. Temporal responses of transcripts, enzyme activities and metabolites after adding sucrose to carbon-deprived Arabidopsis seedlings. Plant J 2007:49:463–491

Oszvald M, Primavesi LF, Grif A, Cohn J, Basu S, Nuccio ML, Paul MJ. Trehalose 6-phosphate regulates photosynthesis and assimilate partitioning in reproductive tissue. Plant Physiol 2018:176:2623–2638

Pal SK. Liput M. Piques M, Ishihara H, Martins MCM, Sulpice R, van Dongen J, Yadav UP, Lunn JE, Usadel B, Schulze WX, Stitt M. Diurnal changes of polysome loading track sucrose content in the rosette of wildtype Arabidopsis and the starchless *pgm* mutant. Plant Physiol. 2013:162:1246–1265

Paul MJ, Jhurreea D, Zhang Y, Primavesi LF, Delatte T, Schluepmann H, Wingler A. Upregulation of biosynthetic processes associated with growth by trehalose 6-phosphate. Plant Signal Behavior 2010:5:386–392

Paul MJ, Watson A, Griffiths CA. Linking fundamental science to crop improvement through understanding source and sink traits and their integration for yield enhancement. J Exp Bot 2020:71: 2270–2280

Pedrotti L, Weiste C, Nägele T, Wolf E, Lorenzin F, Dietrich K, Mair A, Weckwerth W, Teige M, Baena-González E, Dröge-Laser W. Snf1-RELATED KINASE 1-controlled C/S 1-bZIP signaling activates alternative mitochondrial metabolic pathways to ensure plant survival in extended darkness. Plant Cell 2018:30:495–509

Peixoto B, Moraes TA, Mengin V, Margalha L, Vicente R, Feil R, Höhne M, Sousa AGG, Lilue J, Stitt M, et al. Impact of the SnRK1 protein kinase on sucrose homeostasis and the transcriptome during the diel cycle. Plant Physiol 2021:187:1357–1373

Peixoto B, Baena-González E. Management of plant central metabolism by SnRK1 protein kinases. J Exp Bot 2022:73(20):7068–7082 doi: 10.1093/jxb/erac261.

Polge C, Jossier M, Crozet P, Gissot L, Thomas M. β-Subunits of the SnRK1 complexes share a common ancestral function together with expression and function specificities; physical interaction with nitrate reductase specifically occurs via AKINβ1-subunit. Plant Physiol 2008:148:1570–1582

Ponnu J, Wahl V, Schmid M. Trehalose-6-phosphate: Connecting plant metabolism and development. Front Plant Sci 2011:2:70

Ponnu J, Schlereth A, Zacharaki V, Dzialo MA, Abel C, Feil R, Schmid M, Wahl V. 2020. The trehalose 6-phosphate pathway impacts vegetative phase change in Arabidopsis thaliana. Plant J. 2020104(3):768–780. doi: 10.1111/tpj.14965

Ramon M, Rolland F, Thevelein JM, Van Dijck P, Leyman B. ABI4 mediates the effects of exogenous trehalose on Arabidopsis growth and starch breakdown. Plant Mol. Biol.2007:63:195–206

Ramon M, De Smet I, Vandesteene L, Naudts M, Leyman B, Van Dijck P, Rolland F, Beeckman T, Thevelein JM. Extensive expression regulation and lack of heterologous enzymatic activity of the Class II trehalose metabolism proteins from Arabidopsis thaliana. Plant Cell Environ 2009:32:1015–1032 doi: 10.1111/j.1365-3040.2009.01985.x

Ramon M, Dang TVT, Broeckx T, Hulsmans S, Crepin N, Sheen J, Rolland F. Default activation and nuclear translocation of the plant cellular energy sensor SnRK1 regulate metabolic stress responses and development. Plant Cell 2019:31:1614–1632

Randall RS. The plant AlcR-pAlcA ethanol-inducible system displays gross growth artefacts independently of downstream pAlcA-regulated inducible constructs. Sci Rep 2021:11:1–5

Rahmani F, Hummel M, Schuurmans J, Wiese-Klinenberg A, Smeekens S, Hanson J. Sucrose control of translation mediated by an upstream open reading frame-encoded peptide. Plant Physiol. 2009:150:1356–1367

Robinson MD, Oshlack A. A scaling normalization method for differential expression analysis of RNA-seq data. Genome Biol. 2010:11(3):25 doi: 10.1186/gb-2010-11-3-r25.

Rolland F, Baena-Gonzalez E, Sheen J. Sugar sensing and signaling in plants: conserved and novel mechanisms. Annu Rev Plant Biol 2006:57:675–709

Ryabova LA, Robaglia C, Meyer C. Target of Rapamycin kinase: central regulatory hub for plant growth and metabolism. J Exp Bot 2019:70:2211–2216

Sabatini DM. Twenty-five years of mTOR: Uncovering the link from nutrients to growth. Proc Natl Acad Sci USA 2018:114: doi/10.1073/pnas.1716173114

Sánchez-Villarreal A, Davis AM, Davis SJ. AKIN10 activity as a cellular link between metabolism and circadian-clock entrainment in Arabidopsis thaliana. Plant Signal Behav. 2018:13(3):e1411448

Satoh-Nagasawa N, Nagasawa N, Malcomber S, Sakai H, Jackson D. A trehalose metabolic enzyme controls inflorescence architecture in maize. Nature 2006:441:227–230

Scarpin MR, Leiboff S, Brunkard JO. Parallel global profiling of plant TOR dynamics reveals a conserved role for LARP1 in translation. eLife 2020;9:e58795. 10.7554/eLife.58795

Scarpin MR, Simmons CH, Brunkard J. Translating across kingdoms: target of rapamycin promotes protein synthesis through conserved and divergent pathways in plants. J Exp Bot 2022:73(20):7016–7025 doi: 10.1093/jxb/erac267

Schluepmann H, Pellny T, van Dijken A, Smeekens S, Paul M. Trehalose 6-phosphate is indispensable for carbohydrate utilization and growth in *Arabidopsis thaliana*. Proc Natl Acad Sci. USA 2003:100: 6849–6854

Schwacke R, Ponce GY, Krause K, Arsova B, Hallab A, Bolger AM, Gruden K, Leran S, Stitt M, Bolger ME, Usadel B. Mapman4: a refined protein classification and annotation framework applicable to multi omics data analysis. Mol Plant 2019:12:879–892

Schweizer F, Fernandez-Calvo P, Zander M, Diez-Diaz M Fonseca S, Glauser G, Lewsey MG, Ecker JR, Solano R, Reymond P. Arabidopsis basic helix-loop-helix transcription factors MYC2, MYC3, and MYC4 regulate glucosinolate biosynthesis, insect performance, and feeding behavior. Plant Cell 2013:25: 3117–3132

Sheen J. Metabolic repression of transcription in higher plants. Plant Cell 1989:2:1027–1038

Shin J, Sánchez-Villarreal A, Davis AM, Du SX, Berendzen KW, Koncz C, Ding Z, Li C, Davis SJ. The metabolic sensor AKIN10 modulates the Arabidopsis circadian clock in a light-dependent manner. Plant Cell Environ 2017:40:997–1008

Simon NLS, Jelena K, Fernandez-Lopez A, Chembath A, Belbin FE, Dodd AN. The Energy-Signaling Hub SnRK1 Is Important for Sucrose-Induced Hypocotyl Elongation. Plant Physiol. 2018:176:1299–1310 DOI10.1104/pp.17.01395

Shor E, Paik I, Kangisser S, Green R, Huq E. PHYTOCHROME INTERACTING FACTORS mediate metabolic control of the circadian system in Arabidopsis. New Phytol 2017:215:217–228

Shor E, Potavskaya R, Kurtz A, Paik I, Huq E, Green R. PIF-mediated sucrose regulation of the circadian oscillator is light quality and temperature dependent. Gene 2018:9(12):628 doi:10.3390/genes9120628

Son O, Kim S, Cheon CI. Identification of nucleosome assembly protein 1 (NAP1) as an interacting partner of plant ribosomal protein S6 (RPS6) and a positive regulator of rDNA transcription. Biochem Biophy Res Commun 2015:465:200–205

Stitt M, Lilley RMcC, Gerhardt R, Heldt HW. Determination of metabolite levels in specific cells and subcellular compartments of plant leaves. Methods in Enzymology. 1989:174:518–552

Stitt M, Krapp A. The molecular physiological basis for the interaction between elevated carbon dioxide and nutrients. Plant Cell Environ. 1999:22:583–622

Stitt M, Müller C, Matt P, Gibon Y, Carillo P, Morcuende R, Scheible W-R, Krapp A. Steps towards an integrated view of nitrogen metabolism. J Exp Bot 2002:370:959–970

Szklarczyk D, Gable AL, Nastou KC, Lyon D, Kirsch R, Pyysalo S, Doncheva NT, Legeay M, Fang T, Bork P, Jensen LJ, von Mering C. The STRING database in 2021: customizable protein–protein networks, and functional characterization of user-uploaded gene/measurement sets. Nucl Acids Res 2021:49: 10800, 10.1093/nar/gkab835

Sulpice R, Flis A, Ivakov AA, Apelt F, Krohn N, Encke B, Abel C, Feil R, Lunn JE, Stitt M, et al. Arabidopsis coordinates the diurnal regulation of carbon allocation and growth across a wide range of photoperiods. Mol Plant 2014:7:137–155

Takeshi I. What is going on with the hormonal control of flowering in plants? Plant J 2021:105:431– 445

Thimm O, Bläsing O, Gibon Y, Nagel A, Meyer S, Krüger P, Selbig J, Müller LA, Rhee SY, Stitt M. MapMan: a user-driven tool to display genomics datasets onto diagrams of metabolic pathways and other biological processes. Plant J 2004:37:914–939

Tian L, Xie Z, Lu C, Hao X, Wu S, Huan Y, Li D, Chen L. The trehalose-6-phosphate synthase TPS5 negatively regulates ABA signaling in *Arabidopsis thaliana*. Plant Cell Rep 2019:38:869–882

Usadel B, Nagel A, Steinhauser D, Gibon Y, Bläsing OE, Redestig H, Sreenivasulu N, Krall L, Hannah MA, Fernie AR, Stitt M. PageMan an interactive ontology tool to generate, display and annotate overview graphs for profiling experiments. BMC Bioinformatics 2006:7:535

Usadel B, Bläsing OE, Gibon Y, Retzlaff K, Höhne M, Günther M, Stitt M. Global transcript levels respond to small changes of the carbon status during progressive exhaustion of carbohydrates in Arabidopsis rosettes. Plant Physiol 2008:146:1834–1861

Van Leene Y, Eeckhout D, Gadeyne A, Matthijs C, Han C, De Winne N, Persiau G, Van De Slijke E, Persyn F, Mertens T, et al. Mapping of the plant SnRK1 kinase signaling network reveals a key regulatory role for the class II T6P synthase-like proteins. Nature Plants 2022:8:1241–1265

Vandesteene L, López-Galvis L, Vanneste K, Feil R, Maere S, Lammens W, Rolland F, Lunn JE, Avonce N, Beeckman T, Van Dijck P. Expansive evolution of the *TREHALOSE-6-PHOSPHATE PHOSPHATASE* gene family in Arabidopsis. Plant Physiol. 2012:160(2):884–896 doi: 10.1104/pp.112.201400

Vandesteene L, Ramon M, Le K, van Dijck P, Rolland F. A single active trehalose-6-P synthase (TPS) and a family of putative regulatory TPS-like proteins in Arabidopsis. Mol Plant. 2010:3(2):406–419

Viana AJ., Matiolli CC, Newman DW, Vieira JGP, Duarte GT. Martins MCM, Gilbault E, Hotta CT, Caldana C, Vincentz M. The sugar-responsive circadian clock regulator bZIP63 modulates plant growth. New Phytol 2021:231:1875–1889

Vidal EA, Alvarez JM, Araus V, Riveras E, Brooks MD, Krouk G, Ruffel S, Lejay L, Crawford NM, Coruzzi GM, Guriérrez RA. Nitrate in 2020: Thirty years from transport to signaling networks. Plant Cell 2020:32:2094–2119 doi: 10.1105/tpc.19.00748.

Vincentz M, Moureaux T, Leydecker M-T, Vaucheret H, Caboche M. Regulation of nitrate and nitrite reductase expression in Nicotiana plumbaginifolia leaves by nitrogen and carbon metabolites. Plant J. 1993:3(2):315–324

von Schaewen A, Stitt M, Schmidt R, Sonnewald U, Willmitzer L. Expression of a yeast-derived invertase in the cell wall of tobacco and Arabidopsis plants leads to accumulation of carbohydrate, inhibition of photosynthesis and strongly influences growth and phenotype of transgenic tobacco plants. EMBO J 1991:9:3033–3044

Wahl V, Ponnu J, Schlereth A, Arrivault S, Langenecker T, Franke A, Feil R, Lunn JE, Stitt M, Schmid M. Regulation of flowering by trehalose-6-phosphate signaling in *Arabidopsis thaliana*. Science 2013:339:704–707

Wang P, Zhao Y, Li Z, Hsu C.C, Liu X, Fu L, Hou Y-J, Du Y, Xi S, Zhang C, et al. Reciprocal regulation of the TOR kinase and ABA receptor balances plant growth and stress response. Mol. Cell 2018a:69:100– 112

Wang R, Guegler K, LaBrie ST, Crawford NM. Genomic analysis of a nutrient response in Arabidopsis reveals diverse expression patterns and novel metabolic and potential regulatory genes induced by nitrate. Plant Cell 2000:12(8):1491–1509 doi: 10.1105/tpc.12.8.1491.

Wang H, Han C, Wang JG, Chu X, Shi W, Yao L, Chen J, Hao W, Deng Z, Fan M, Bai MY. Regulatory functions of cellular energy sensor SnRK1 for nitrate signalling through NLP7 repression. Nat Plants 2022:8:1094–1107 doi: 10.1038/s41477-022-01236-5

Wang Y, Wang L, Micallef BJ, Tetlow IJ, Mullen RT, Feil R, Lunn JE, Emes MJ. AKINβ1, a subunit of SnRK1, regulates organic acid metabolism and acts as a global modulator of genes involved in carbon, lipid, and nitrogen metabolism. J Exp Bot 2020:71:1010–1028

Wang Z, Wang XM, Xie B, Hong ZL, Yang QC. Arabidopsis NUCLEOSTEMIN-LIKE 1 (NSN1) regulates cell cycling potentially by cooperating with nucleosome assembly protein AtNAP1;1. BMC Plant Biol 2018b:18:99 doi 10.1186/s12870-018-1289-2

Warner JR. The economics of ribosome biosynthesis in yeast. Trends Biochem Sci: 1999: 24:437–440

Wiese A, Elzinger N, Wobbes B, Smeekens S. A conserved upstream open reading frame mediates sucrose-induced repression of translation. Plant Cell 2004:16:1717–1729

Williams SP, Rangarajan P, Donahue JL, Hess JE, Gillaspy GE. Regulation of Sucrose non-Fermenting Related Kinase 1 genes in *Arabidopsis thaliana*. Front Plant Sci 2014:5:324

Wu Y, Shi L, Li L, Fu L, Liu Y, Xiong Y, Sheen J. Integration of nutrient, energy, light, and hormone signaling via TOR in plants. J Exp Bot 2019:70:2227–2238

Xie Q, Wang P, Liu X, Yuan L, Wang L, Zhang C, Li W, Xing H, Zhi L, Yue Z, et al. LNK1 and LNK2 are transcriptional coactivators in the Arabidopsis circadian oscillator. Plant Cell 2014:26:2843–2857

Xue X, Wang J, Shukla D, Cheung LS, Chen L-Q. When SWEETs turn tweens: updates and perspectives. Annu Rev. Plant Biology 2022:73:379–403

Yadav UP, Ivakov A, Feil R, Duan GY, Walther D, Giavaliaco P, Piques M, Carillo P, Hubberton HM, Stitt M, Lunn JE. The sucrose–trehalose 6-phosphate (Tre6P) nexus: specificity and mechanisms of sucrose signaling by Tre6P. J Exp Bot 2014:65:1051–1068

Zacharaki V, Ponnu J, Crepin N, Langenecker T, Hagmann J, Skorzinski N, Musialak-Lange M, Wahl V, Rolland F, Schmid M. Impaired KIN10 function restores developmental defects in the Arabidopsis *trehalose 6-phosphate synthase1* (*tps1*) mutant. New Phytol 2022:235:220–233

Zhai Z, Keereetaweep J, Liu H, Feil R, Lunn JE, Shanklin J. Trehalose 6-phosphate positively regulates fatty acid synthesis by stabilizing WRINKLED1. Plant Cell 2018:30:2616–2627

Zhang ZP, Deng Y, Song X, Miao M. Trehalose-6-phosphate and SNF1-related protein kinase 1 are involved in the first-fruit inhibition of cucumber. J Plant Physiol 2015:177:110–120

Zhang Z, Zhu J-Y, Roh J, Marchive C, Kim S-K, Meyer C, Sun Y, Wang W Wang Z-Y. TOR signaling promotes accumulation of BZR1 to balance growth with carbon availability in Arabidopsis. Current Biol 2016:26:1854–1860

Zhang ZZ, Sun Y, Jiang J, Wang WF, Wang, ZY. Sugar inhibits brassinosteroid signaling by enhancing BIN2 phosphorylation of BZR1. PLOS Genet 2021:17:e1009540

