## Supplementary material for "Direct and indirect responses of the Arabidopsis transcriptome to an induced increase in trehalose 6-phosphate": Combined Supplemental Figures and Tables

| **Supplemental Figures** | **Page** |
|---|---|
| Supplemental Figure S1 | 2 |
| Supplemental Figure S2 | 3 |
| Supplemental Figure S3 | 4 |
| Supplemental Figure S4 | 6 |
| Supplemental Figure S5 | 8 |
| Supplemental Figure S6 | 10 |
| Supplemental Figure S7 | 13 |
| Supplemental Figure S8 | 22 |
| Supplemental Figure S9 | 23 |
| Supplemental Figure S10 | 24 |
| Supplemental Figure S11 | 25 |
| Supplemental Figure S12 | 26 |
| Supplemental Figure S13 | 28 |
| Supplemental Figure S14 | 29 |
| Supplemental Figure S15 | 33 |
| Supplemental Figure S16 | 35 |
| Supplemental Figure S17 | 36 |
| Supplemental Figure S18 | 37 |
| Supplemental Figure S19 | 38 |
| Supplemental Figure S20 | 43 |
| Supplemental Figure S21 | 47 |
| Supplemental Figure S22 | 49 |
| Supplemental Figure S23 | 50 |
| Supplemental Figure S24 | 51 |
| Supplemental Figure S25 | 53 |
| Supplemental Figure S26 | 54 |
| **Supplemental Tables** | |
| Supplemental Table S1 | 55 |
| Supplemental Table S2 | 56 |
| Supplemental Table S3 | 57 |
| Supplemental Table S4 | 58 |
| Supplemental Table S5 | 59 |
| Supplemental Table S6 | 60 |

**Supplemental Figure S1. Comparison of changes in transcript abundance 12 h after induction of TPS in two independent iTPS lines, 29.2 and 31.3.**

iTPS lines 29.2 and 31.3 and the alcR control were grown in an equinoctial (12 h light/12 h dark) light regime at 160 µmol quanta $m^{-2}$-$s^{-1}$ for 28 days, and then either sprayed with 2% ethanol or water at dawn and harvested 12 h later at dusk (ED treatment), or sprayed with 2% ethanol or water at dusk and harvested 12 h later at dawn (EN treatment). Transcriptome analysis was performed using ATH1 arrays and data are supplied in Supplemental Dataset S1. The change in transcript abundance after ethanol treatment of an iTPS line was calculated compared to the water-sprayed iTPS line. There were only minor changes in transcript abundance due to ethanol addition to alcR controls and this was corrected as explained in Materials and Methods. The results are the mean of 3 samples (each sample consisted of 5-10 whole rosettes).

(A) Venn diagrams. The numbers represent transcripts that passed the filter. specified in the display.

(B) Reproducibility of response between *iTPS29.2* and *iTPS31.3* at dusk after inducing at dawn; scatter plots are provided for all transcripts and after filtering (FDR < 0.05).

(C) Reproducibility of response between *iTPS29.2* and i*TPS31.3* at dawn after inducing at dusk.

A.

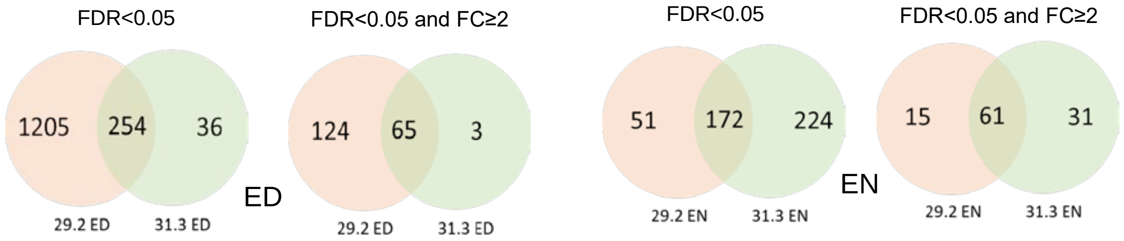

B.

ED treatment

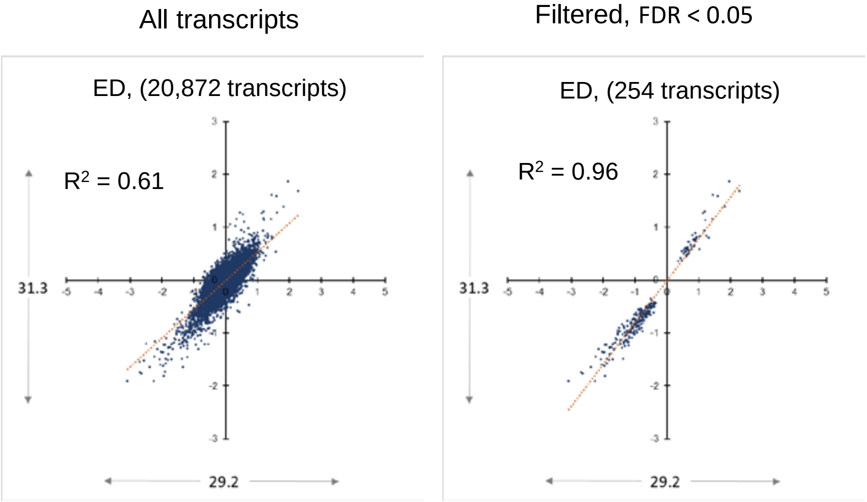

C.

EN treatment

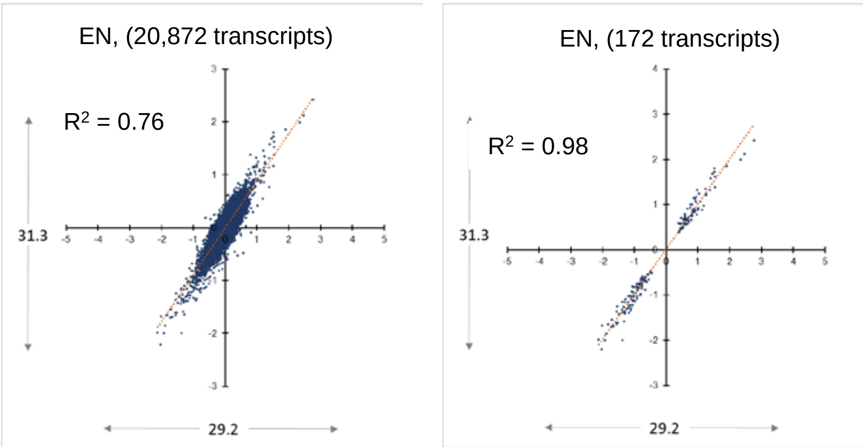

**Supplemental Figure S2**. **Estimation of the carbon response factor, CRF**. The carbon response factor (CRF) summarizes the response to an increase in sugars across a set of nine treatments. A positive and negative CRF value represents an increase or decrease in transcript abundance in response to elevated sugar, respectively. The CRF was calculated using the response of a given for each transcript in nine treatments that included four treatments in which sugars were supplied to C-starved seedlings, and five treatments that changed endogenous sucrose and reducing sugar levels in rosettes of plants growing in an equinoctial light-dark cycle at 160 µmol quanta $m^{-2}$-$s^{-1}$, similar to the conditions used for the iTPS experiments. Especially in diel cycles, transcript abundance is strongly influenced by circadian clock- or light-signaling (Bläsing et al., 2005; Usadel et al., 2008). The treatments were chosen to minimize the impact of circadian clock- or light signaling in transcript abundance.

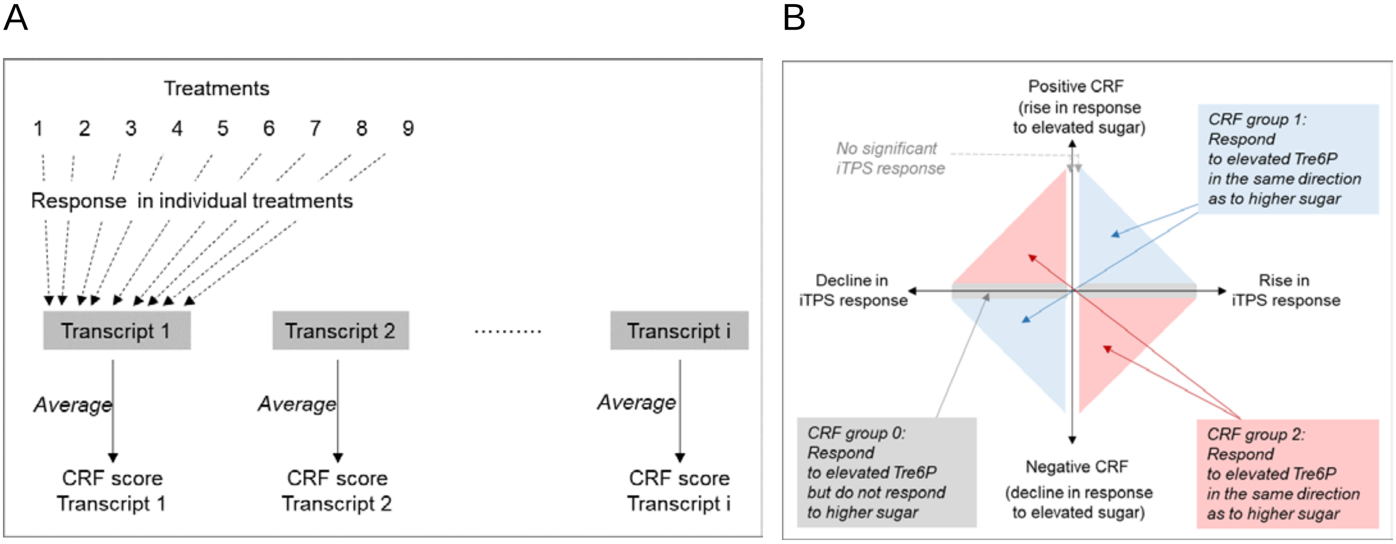

(A) Illustration of the calculation: (1-4) rosettes of *pgm* mutant plants grown at equinoctial light regime compared to wild-type Col-0 at ZT8, ZT12 (ED), ZT20 and ZT24 (EN), respectively (Gibon et al., 2004; Blasing et al., 2005); (5-8) whole seedlings in liquid culture grown under low (50 µmol quanta $m^{-2}$-$s^{-1}$) continuous light: (5) full nutrition compared 2 days C-starved seedlings (Osuna et al., 2007); (6) 30 min after adding 15mM sucrose to C-starved seedlings compared to full nutrition (Osuna et al., 2007); (7) 3-h after adding 15mM sucrose to C-starved seedlings compared to full nutrition (Osuna et al., 2007); (8) 30 min after adding 100 mM glucose to C-starved seedlings compared to full nutrition (Blasing et al., 2005); (9) Rosettes of vegetatively growing plants illuminated for 4h after dawn at 350 compared to 50 ppm $CO_2$ (Blasing et al., 2005). $Log_2$ FC values (filtered as greater or smaller than $+log_2 0.1$ or $-log_2 0.1$, respectively) were averaged across the nine treatments taking into account the treatment effect on C status, hence treatments 3-4 were assigned a reverse sign as they led to a decrease in C. The low filter was used to minimize the number of genes that were classified as C-unresponsive (see Supplemental Figure S3A-B and Methods for further details).

B) Use of the CRF in combination with the observed change in transcript abundance in iTPS to assign genes to groups. Group 1 ($G_1$) denotes transcripts where the iTPS response and CRF are qualitatively the same and, by inference, the iTPS response may be a direct response to elevated Tre6P, group 2 ($G_2$) denotes transcripts where the iTPS response and CRF are qualitatively opposed and by inference the iTPS response is unlikely to be a direct response to elevated Tre6P but is likely to be a independent response to lower sucrose or related metabolites, and group 0 ($G_0$) denotes transcripts that respond in the iTPS response but cannot be scored because they do not show a consistent response to changes in sugars.

**Supplemental Figure S3. Comparison of the carbon response factor (CRF) and iTPS at dusk and dawn.** The analysis was performed with the transcript abundance dataset for line 29.2 from Supplemental Figure S1. In the ED treatment, TPS was induced at dawn and plants were harvested at the end of the light period, and in the EN treatment TPS was induced at dusk and plants were harvested at the end of the night.

The Carbon response factor (CRF) summarizes the response of a given transcript to a change in sugars across a set of treatments, with an increasingly positive sign indicating the average increase in abundance and an increasingly negative sign denoting the average decrease in abundance, whilst a value around zero indicates that transcript abundance does not respond to sugar status (for derivation see Supplemental Figure S2, Supplemental Dataset S2 and main text).

(A) Total iTPS.29.2 response in the ED treatment (FC $log_2$) compared to the CRF.

(B) Total iTPS29.2 response in the EN treatment (FC $log_2$) compared to the CRF.

(C) Deconvolution of the iTPS response in the ED treatment, based on the CRF assigned to each transcript. Briefly, transcripts assigned to group 1 ($G_1$) respond to elevation of Tre6P in the same direction as their response to an increase of sugar supply in a panel of treatments (as defined by the CRF); transcripts assigned to group 2 ($G_2$) respond to elevation of Tre6P in the opposite direction to their response to an increase of sugar supply in the panel of treatments; transcripts assigned to group 0 ($G_0$) do not show a notable response to a change in sugar (average response cut off: $Log_2FC$ 0.1). For further explanation of the treatments and procedure used to estimate the CF CRF values for each transcript, see Supplemental Figure S2

(D) Deconvolution of the iTPS response in the EN treatment (same procedure as for panel C).

(E) Comparison of the shared deconvoluted responses in the ED and EN iTPS responses at dusk and dawn. Genes were assigned to CRF groups $G_1$, $G_2$ and $G_0$ as described above and in Supplemental Figure S2. Genes assigned to the same CRF group at ED and EN were then regressed against each other.

In all panels, the number in brackets gives the number of transcripts in the analyzed subset.

A.

Complete iTPS response at ED compared to CRF

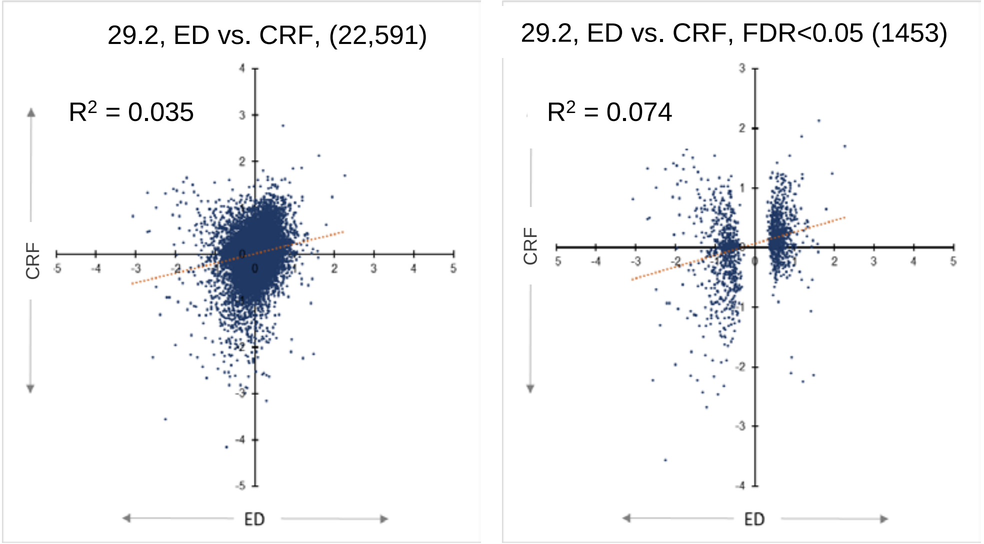

B.

Complete iTPS response at EN compared to CRF

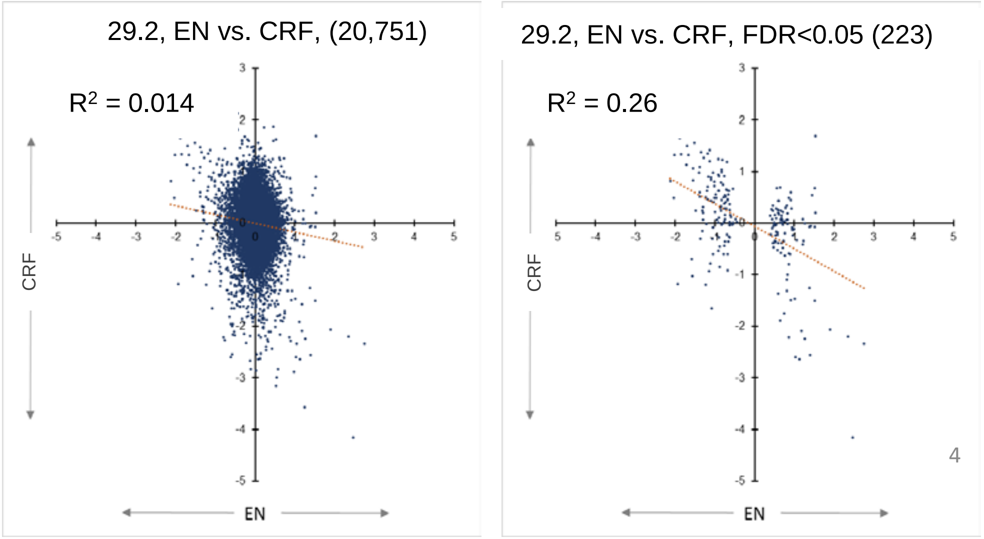

Figure S3 (continued)

#### C. iTPS response at ED compared to CRF after separation into $G_1$, $G_2$, $G_0$

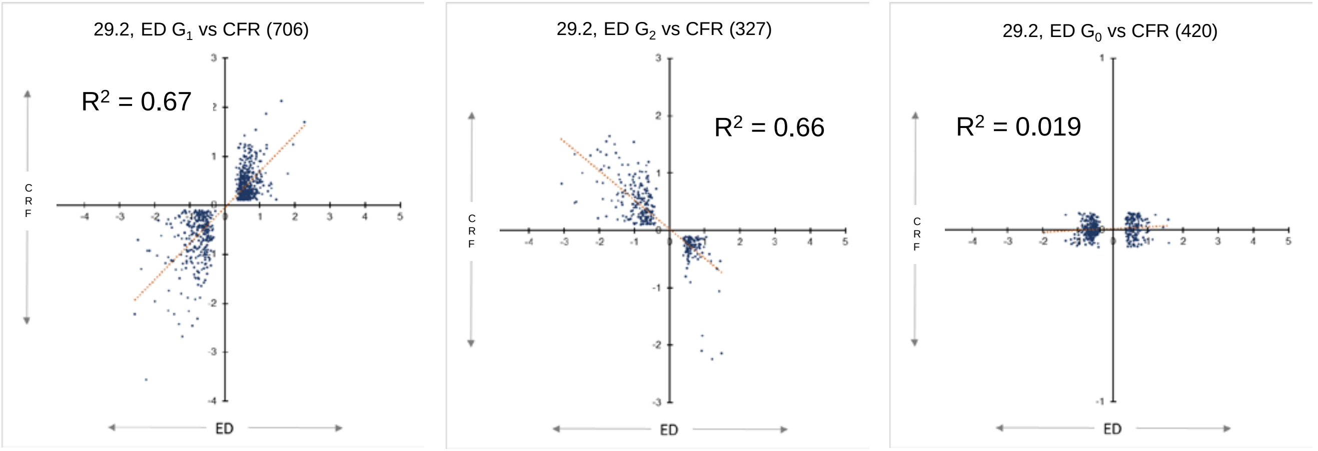

#### D. iTPS response at EN compared to CRF after separation into $G_1$, $G_2$, $G_0$

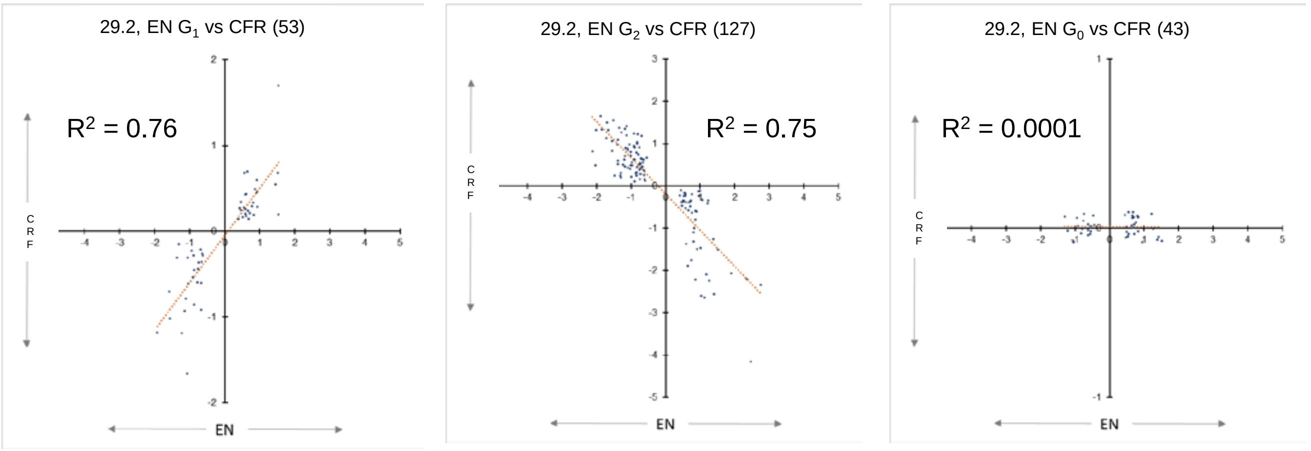

#### E. Comparison of iTPS responses at ED and EN

Genes assigned to $G_1$ | Genes assigned to $G_2$ | Genes assigned to $G_0$

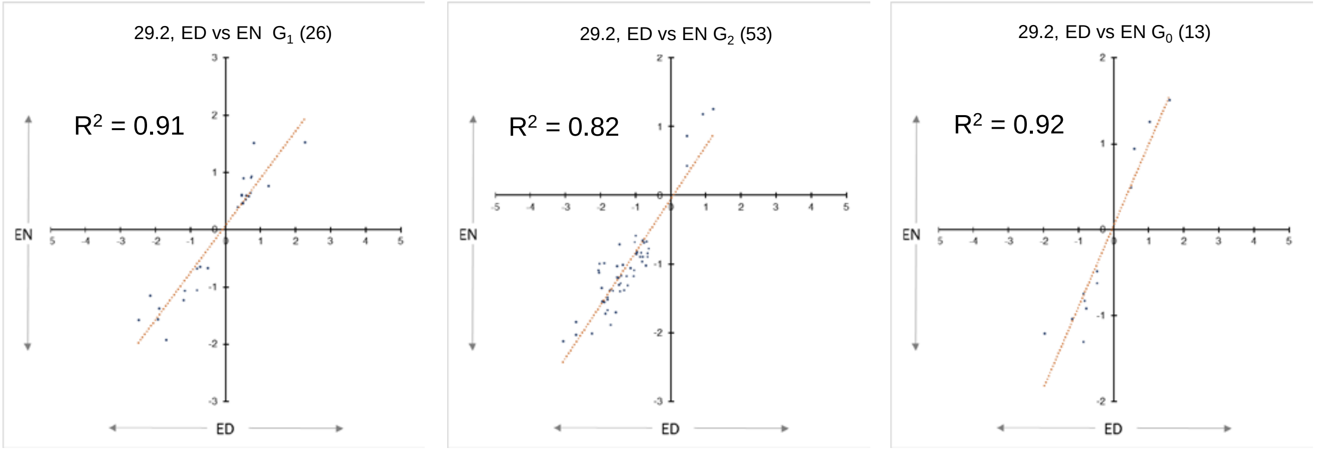

**Supplemental Figure S4. Changes of further metabolites after induction in iTPS29.2 for 4 h or 6 h in the light (supplemental to Figure 1)**. The data are for the experiment shown in Figure 1 and used for RNAseq analyses. The data are provided in Supplemental Dataset S3.

(A) Bacterial TPS protein detected by western blots of water controls and ethanol-sprayed line 29.2

(B) Regression plot of Tre6P against sucrose. The expected positive relationship is seen for the three control treatments (water and ethanol-sprayed alcR, water-sprayed iTPS 29.2. In ethanol-sprayed iTPS29.1, compared to controls and pre-spray ITPS 29.2, there is no marked change in the relationship between Tre6P and sucrose at 2-h post-spraying, and a negative relationship at 4-h and 6-h post-spraying.

(C) Further metabolites: Glc6P, Fru6P, mannose 6-P, PEP, pyruvate, shikimate, malate, fumarate, citrate, aconitate, 2-oxoglutarate. The results are plotted as mean ± S.D. (n = 4) (each replicate contained 4-5 whole rosettes). Significant changes of ethanol-sprayed iTPS 29.2 compared to the three control treatments (water and ethanol-sprayed alcR, water-sprayed iTPS 29.2) are indicated by asterisks (one-way ANOVA, Holm-Sidak, with *, **, *** correspond to P < 0.05, < 0.01, < 0.001).

The original data are provided in SI Dataset S3.

A

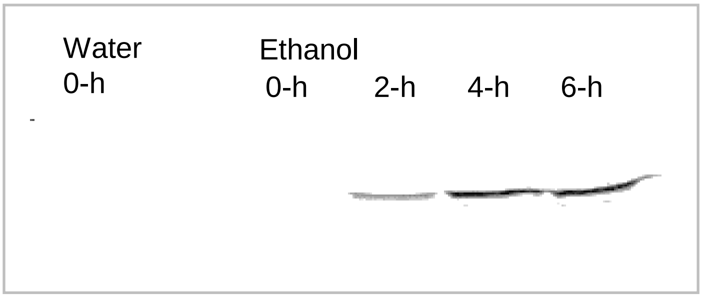

B

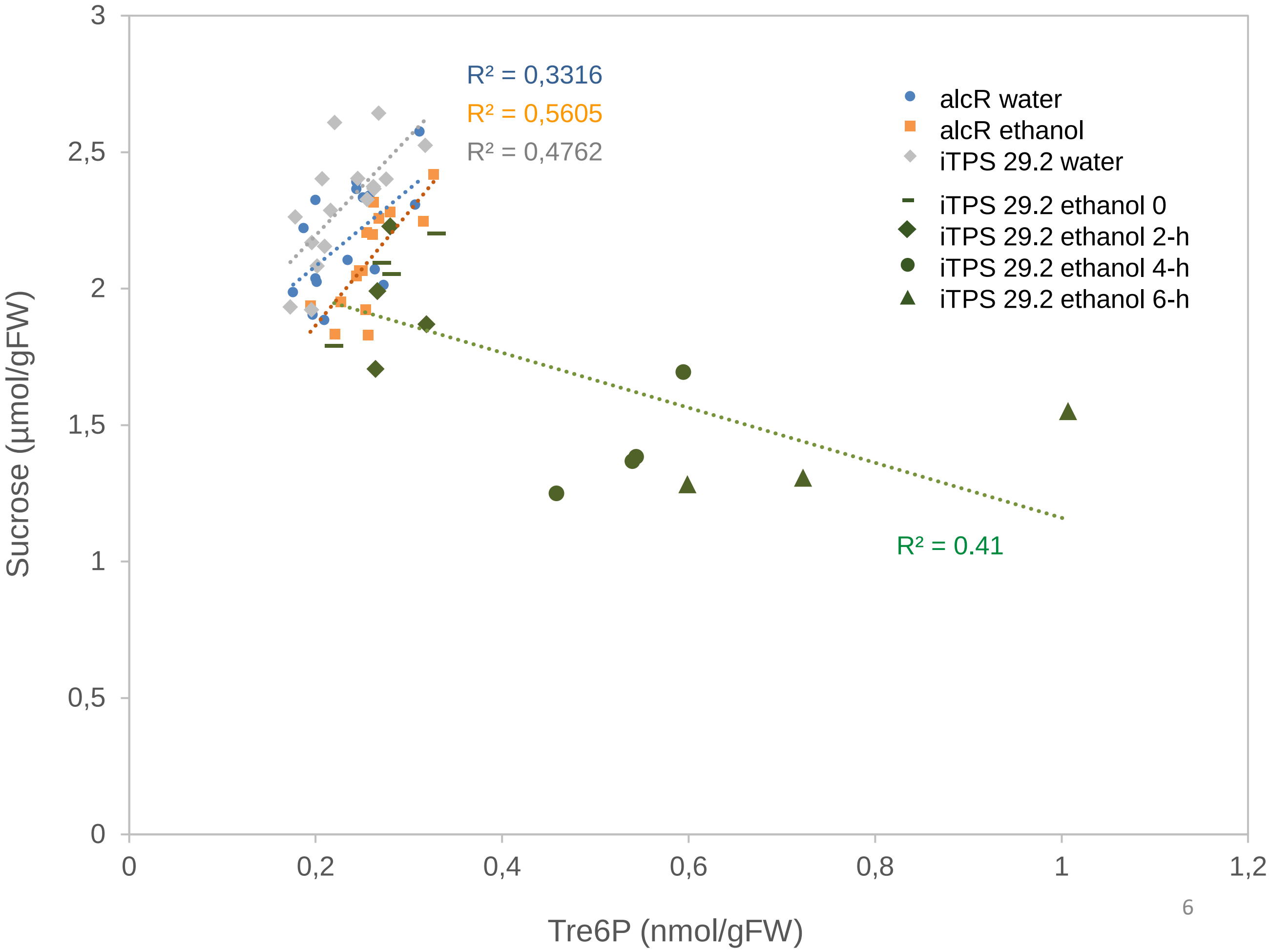

**Supplemental Figure S4. continued.**

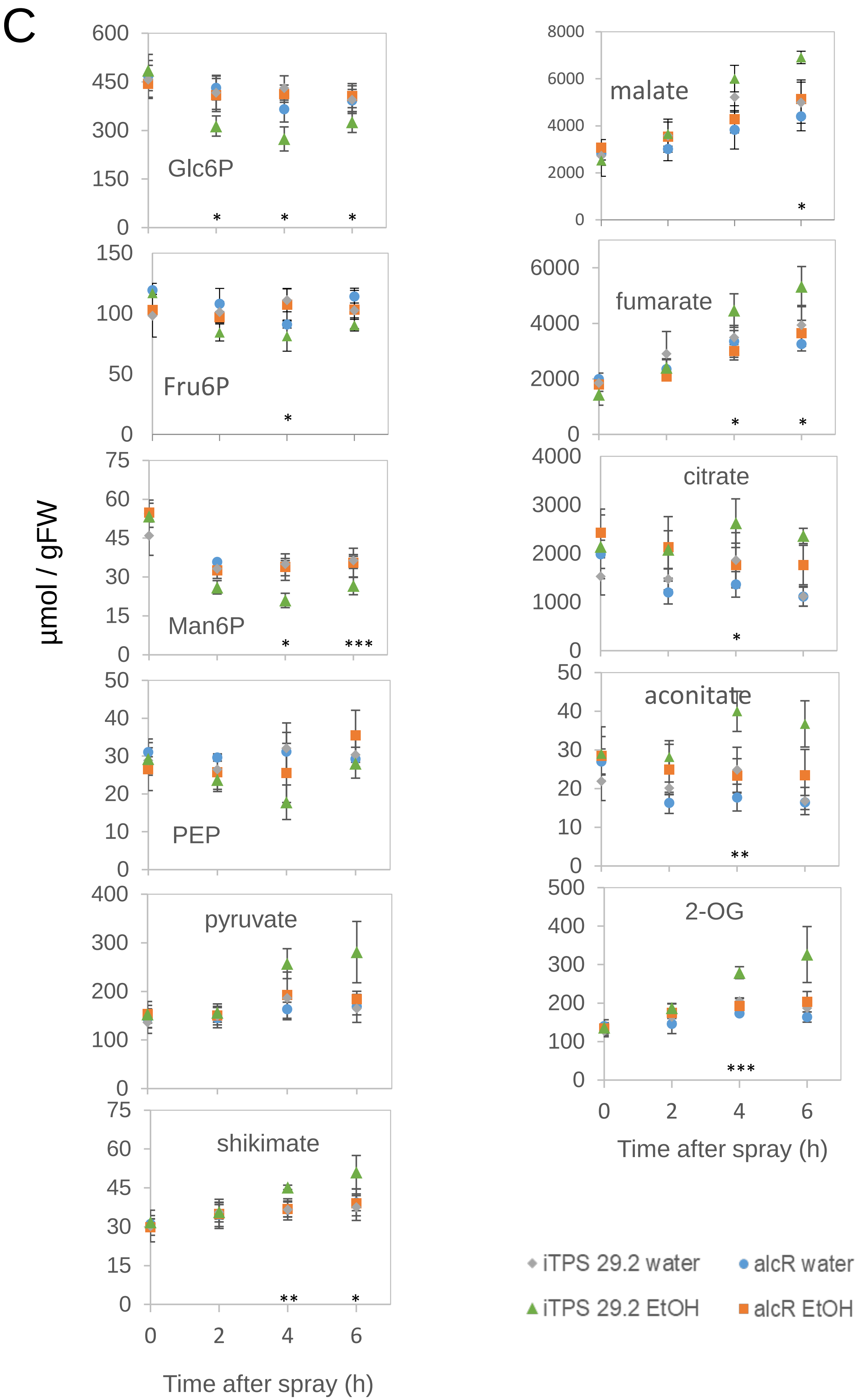

**Supplemental Figure S5. Global analysis of the response of transcript abundance at 4 h and 6 h after induction in *iTPS29.1* (supplemental to Figure 2).**

(A) Comparison of the response at 4 h and 6h; plots are shown for all transcripts, for all transcript whose responses was significant at FDR <0.05, and for all transcripts whose response was significant at FDR<0.05 and a >2-fold change (FC2, $\log_2$ scale).

(B) Plots of the response against the CRF; plots are shown separately for the response at 4 h (Figure 2D) and at 6h (this panel) for which plots are shown from left to right for all transcripts, for all transcript whose responses was significant at FDR <0.05, and for all transcripts whose response was significant at FDR<0.05 and a >2-fold change ($\log_2$ scale). For derivation of the CRF see Supplemental Figure S2.

(C) Reproducibility of the response 4-h or 6-h post-induction in the light compared with to that in the ED and EN treatments (see Supplemental Figure S1). Note that transcript abundance was assessed using different technologies for the 4h and 6h treatments (RNA seq) and the ED and EN treatments (ATH1) with the latter being technically less sensitive for low abundance genes (next page)

In all panels, the number in brackets gives the number of transcripts in the analyzed subset.

A

Overall iTPS response versus CRF

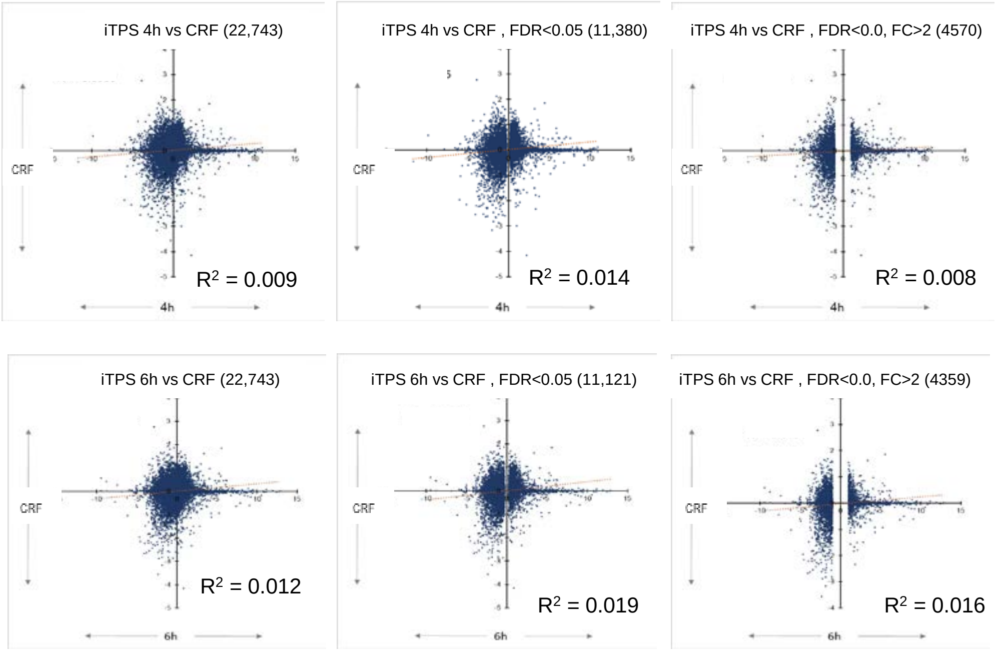

B

iTPS response at 6-h after assignment to $G_1$, $G_2$ and $G_0$ versus CRF

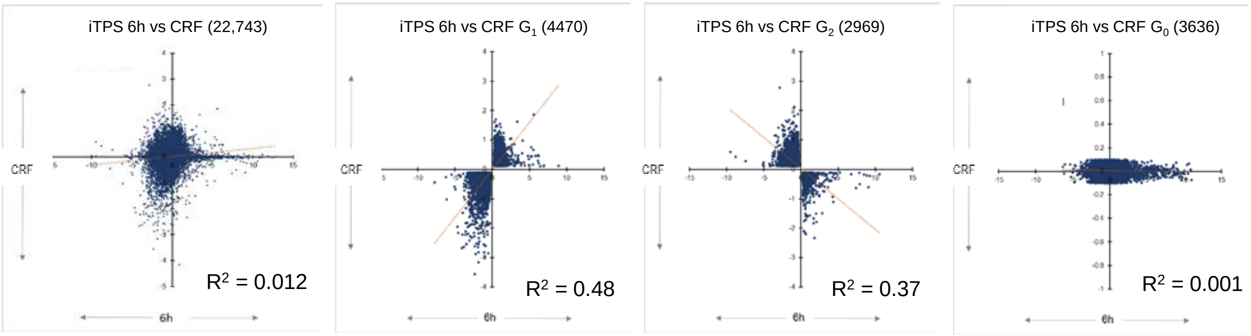

Panel C is on the next page

### Figure S5 (continued)

(C ) Reproducibility of the response 4-h or 6-h post-induction in the light (Figure 2) compared with to that in the ED and EN treatments (see Supplemental Figure S1). Note that transcript abundance was assessed using different technologies for the 4h and 6h treatments (RNA seq) and the ED and EN treatments (ATH1) with the latter being technically less sensitive for low abundance genes

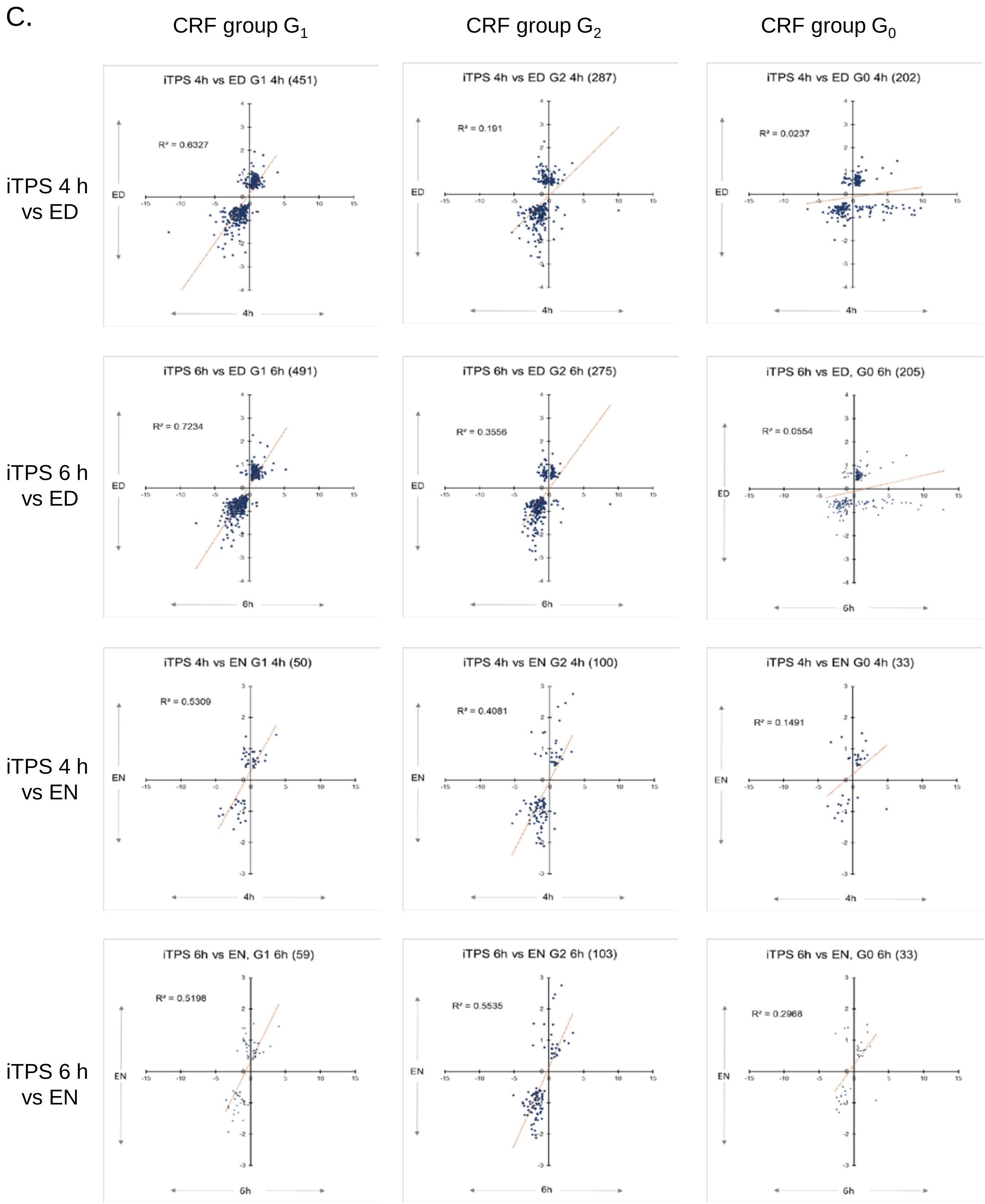

**Supplemental Figure S6. changes in expression of top 10-100 G0 genes in response to various treatments, by which CRF was determined.**

(A) Response of transcript abundance in each of the nine treatments. The plots show, for genes assigned to $G_0$, the response of transcript abundance for the 100 and 10 genes whose transcripts showed the largest increase in the iTPS response, and the 100 and 10 genes whose transcripts showed the largest decrease in the iTPS response. The analysis used the shared response between the iTPS 4h and 6h treatments.

(B) Top ten genes in the iTPS $G_0$ group UP and the iTPS $G_O$ group DOWN and their response in the nine treatments used to estimate the CRF. (see next page)

(C) Impact of transcript abundance on assignment to CRF group $G_0$. (see over-nest page)

The treatments used were (see also Supplemental Figure S2):

- p8-d8, p12-d12, d20-p20, d24-p24: pgm mutant grown at equinoctial light regime compared to Col-0 at ZT8, ZT12 (ED), ZT20 and ZT24 (EN) (Gibon et al., 2004; Blasing et al., 2005);
- FN-St: full nutrition compared 2 days C-starved seedlings (Osuna et al., 2007);
- 30mSUC: 30 min after adding 15mM sucrose to C-starved seedlings compared to full nutrition (Osuna et al., 2007);
- 3hSUC: 3-h after adding 15mM sucrose to C-starved seedlings compared to full nutrition (Osuna et al., 2007);
- 30mGlc: 30 min after adding 100 mM glucose to C-starved seedlings compared to full nutrition (Blasing et al., 2005);
- 350-50$CO_2$: rosettes illuminated for 4 h at 350 compared to 50 ppm (Blasing et al., 2005).

A

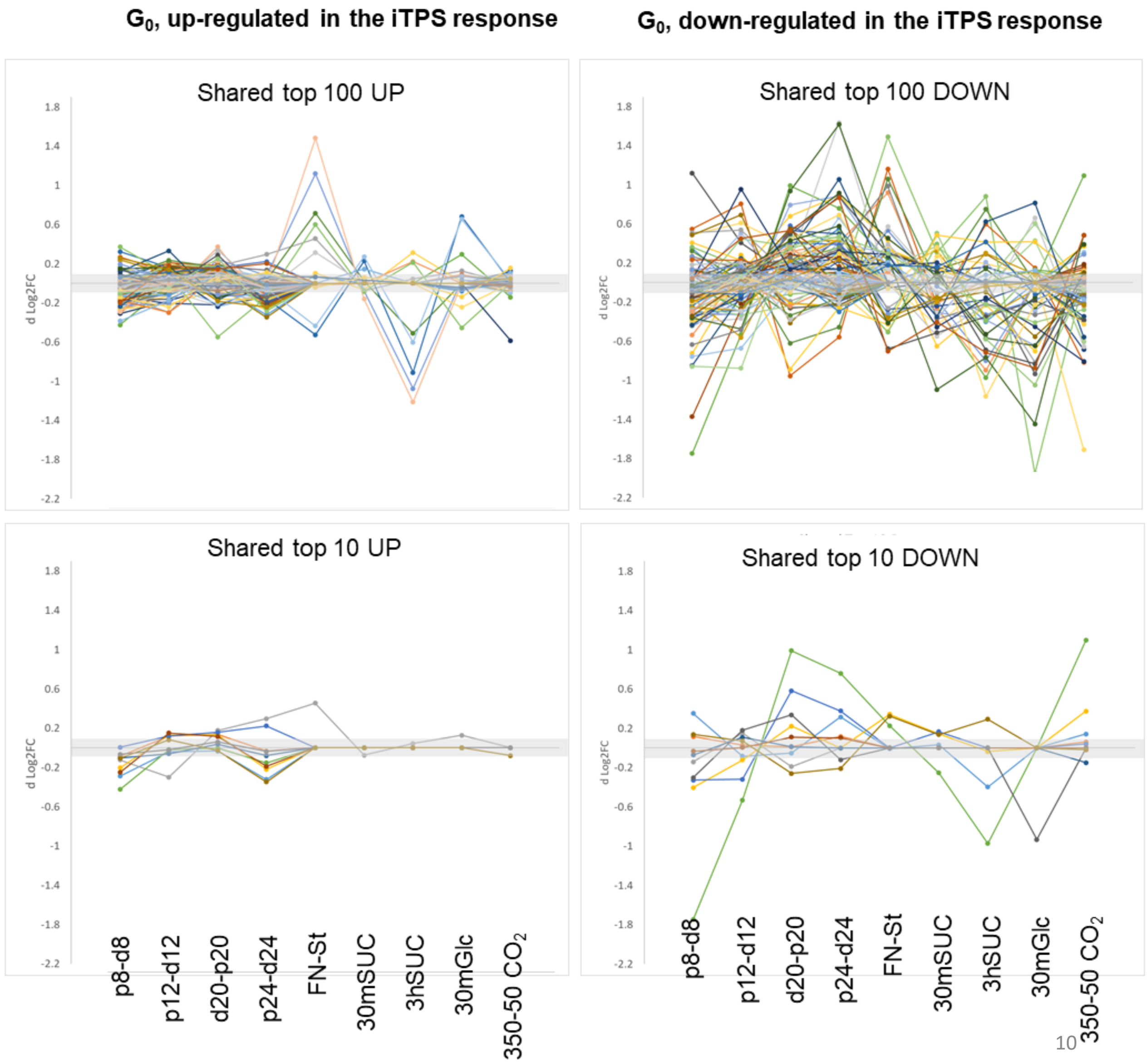

**Figure S6: changes in expression of selected $G_0$ genes in response to various treatments by which the CRF)was determined and abundance of transcripts of genes assigned to $G_0$ (continued)**

(B) The 10 top genes In the iTPS $G_0$ group UP and the iTPS $G_O$ group DOWN and their response in the nine treatments used to estimate the CRF. The analysis shows the responses in the nine treatments used to estimate the CRF (columns 1-9 from the left) and the 4h and 6h iTPS responses (the 2 right-hand columns). The transcripts are ordered based on the response at 4 h.

B

top 10 SHARED UP in CRF group $G_0$ (iTPS 4-h and 6-h)

| ID / treatment | p8-d8 | p12-d12 | p20-d20 | p24-d24 | FN - St | 30mSUC - St | 3hSUC - St | 30mGlc -St | 350 - 50 $CO_2$ | $Log_2FC$ iTPS 4h | $Log_2FC$ iTPS 6h |
|---|---|---|---|---|---|---|---|---|---|---|---|
| At3g62710 | -0.29 | -0.04 | -0.03 | -0.32 | 0.00 | 0.00 | 0.00 | 0.00 | 0.00 | 10.97 | 8.07 |
| At5g19270 | -0.09 | 0.12 | 0.14 | -0.03 | 0.00 | 0.00 | 0.00 | 0.00 | 0.00 | 10.85 | 12.85 |
| At3g14530 | -0.12 | -0.30 | 0.18 | 0.30 | 0.46 | -0.08 | 0.04 | 0.13 | 0.00 | 10.60 | 9.51 |
| At3g13390 | -0.20 | 0.13 | 0.13 | -0.22 | 0.00 | 0.00 | 0.00 | 0.00 | 0.00 | 10.52 | 9.30 |
| At5g67010 | 0.01 | 0.12 | 0.16 | 0.22 | 0.00 | 0.00 | 0.00 | 0.00 | 0.00 | 10.38 | 8.11 |
| At5g62330 | -0.42 | -0.02 | 0.00 | -0.15 | 0.00 | 0.00 | 0.00 | 0.00 | 0.00 | 10.16 | 9.45 |
| At4g36600 | -0.10 | -0.06 | 0.04 | -0.08 | 0.00 | 0.00 | 0.00 | 0.00 | 0.00 | 10.00 | 4.89 |
| At3g63360 | -0.25 | 0.15 | 0.12 | -0.19 | 0.00 | 0.00 | 0.00 | 0.00 | 0.00 | 9.86 | 11.10 |
| At4g29620 | -0.07 | -0.02 | 0.06 | -0.04 | 0.00 | 0.00 | 0.00 | 0.00 | 0.00 | 9.80 | 6.89 |
| At2g45550 | -0.11 | 0.08 | -0.03 | -0.35 | 0.00 | 0.00 | 0.00 | 0.00 | -0.08 | 9.75 | 8.70 |

top 10 SHARED DOWN in CRF group $G_0$ (iTPS 4-h and 6-h)

| ID / treatment | p8-d8 | p12-d12 | p20-d20 | p24-d24 | FN - St | 30mSUC - St | 3hSUC - St | 30mGl c-St | 350 - 50 $CO_2$ | $Log_2FC$ iTPS 4h | $Log_2FC$ iTPS 6h |
|---|---|---|---|---|---|---|---|---|---|---|---|
| At3g14340 | 0.35 | -0.08 | -0.05 | 0.32 | -0.01 | 0.03 | -0.40 | 0.00 | 0.14 | -7.90 | -9.50 |
| At3g13130 | 0.12 | 0.03 | 0.01 | 0.12 | 0.00 | 0.00 | 0.00 | 0.00 | 0.06 | -6.31 | -3.36 |
| At1g49830 | -0.14 | 0.15 | -0.19 | 0.00 | 0.00 | 0.00 | 0.00 | 0.00 | 0.00 | -6.19 | -3.17 |
| At4g16515 | -0.40 | -0.12 | 0.22 | 0.00 | 0.34 | 0.14 | -0.04 | 0.00 | 0.37 | -6.08 | -5.96 |
| At5g64700 | -0.33 | -0.32 | 0.58 | 0.38 | 0.00 | 0.17 | 0.00 | 0.00 | 0.04 | -5.32 | -3.79 |
| At2g40610 | **-1.75** | -0.53 | 0.99 | 0.76 | 0.23 | -0.25 | -0.97 | 0.00 | 1.10 | -5.23 | -4.14 |
| At5g59990 | -0.07 | 0.11 | 0.01 | 0.00 | 0.00 | 0.00 | 0.00 | 0.00 | -0.15 | -5.01 | -2.87 |
| At5g06070 | -0.03 | 0.00 | 0.11 | 0.10 | 0.00 | 0.00 | 0.00 | 0.00 | 0.00 | -4.88 | -4.92 |
| At1g29090 | -0.30 | 0.18 | 0.34 | -0.12 | 0.00 | 0.00 | 0.00 | -0.93 | -0.01 | -4.77 | -5.11 |
| At1g74660 | 0.14 | 0.07 | -0.26 | -0.21 | 0.32 | 0.13 | 0.29 | 0.00 | -0.02 | -4.75 | -2.04 |

**Figure S6. Changes in expression of selected G0 genes in response to various treatments by which the CRF was determined and abundance of transcripts of genes assigned to $G_0$ (continued).**

(C) **Impact of transcript abundance on assignment to CRF groups $G_0$.** All transcripts in a given group were ordered on abundance in the 4h water-sprayed 29.2 control (blue). The plots also show the abundance in the 4h ethanol-sprayed treatment. Taking a filter of 10 counts, there is a larger proportion (of low-expressed gene in the $G_0$ transcript set (15- 20%) than the $G_1$ and $G_2$ sets (under 2%) but the majority of transcript assigned to $G_0$ are present at abundances similar to those in the $G_1$ and $G_2$ sets. The analysis was performed with genes where the iTPS was significant at FDR<0.05.
Note that the x-axes are scaled to have the same value for all genes in set and are, per gene, more expanded for the $G_2$ and $G_0$ plots than the $G_1$ plot

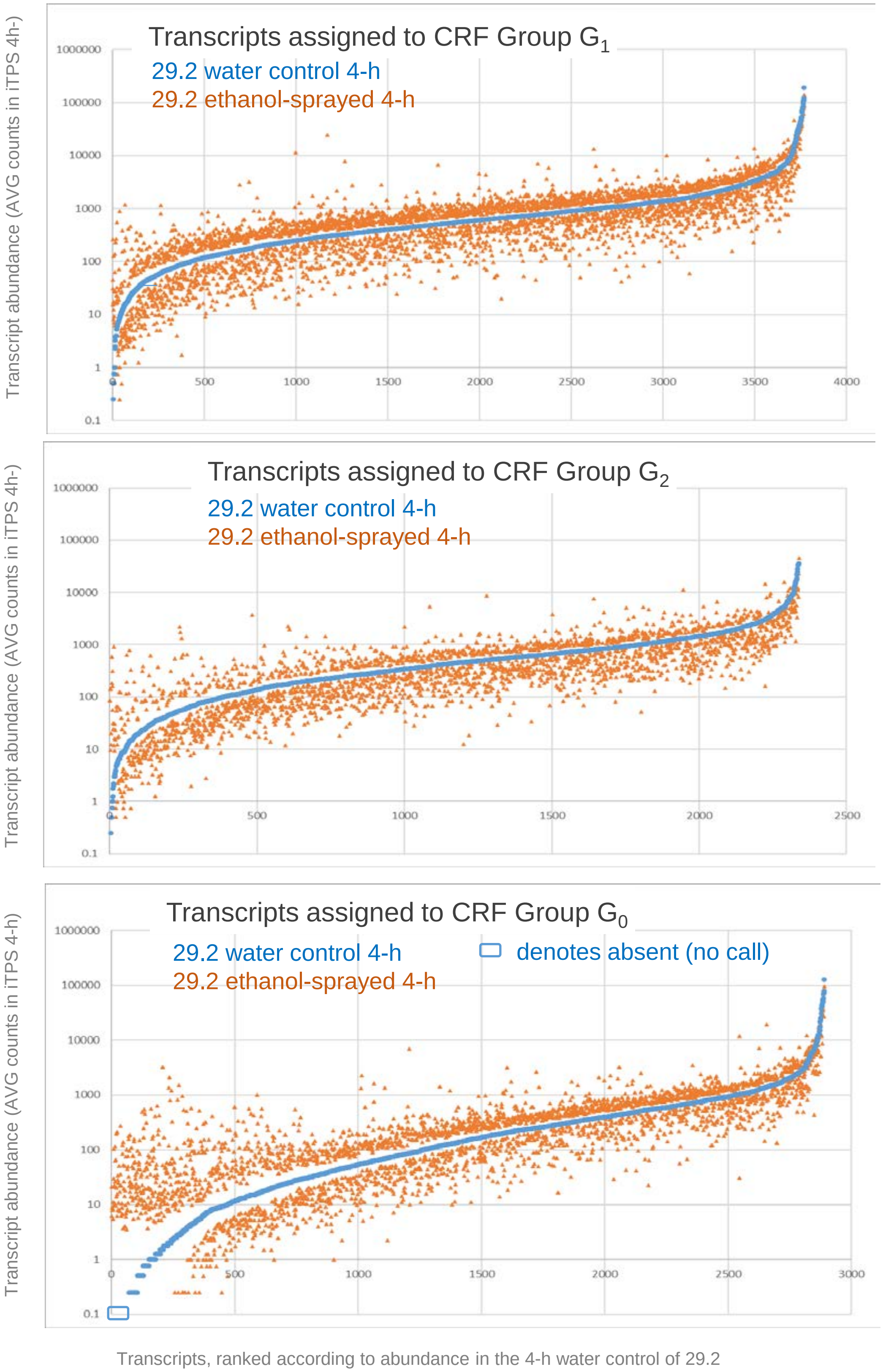

**Supplemental Figure S7. Visualization of selected responses in metabolism using PageMan displays (supplemental to Figure 3).** (A) Photosynthesis, (B) Gluconeogenesis/glyoxylate cycle, (C) N metabolism, (D) Nucleotide metabolism, (E) Specialized metabolism, (F) Protein, (G) Cell wall. The analysis was conducted using PageMan (Usadel et al., 2006) and MapMan software (version 3.6.0RC1; https://mapman.gabipd.org/; Ath_AGI_LOCUS_TAIR10_Aug2012). The analysis was performed as in Figure 3. The analysis was performed separately for the sets of genes that were assigned to the CRF groups $G_1$, $G_2$ and $G_0$ (see Supplemental Figure S2) and for the responses at 4 h and 6 h after spraying. The CFR groups are shown from left to right in the block in which the 4h and 6h response is displayed. The analyses used the $log_2FC$ values for all genes in a given category. These were filtered (FDR <0.05, FC ≥0.2; all values that did not pass the filter were set to zero) and all individual $log_2FC$ values (including the zero values) were thern averaged for all genes in that category. The average $log_2FC$ values for each subBIN are displayed as a heat map (for scale see insert.)

#### A. Photosynthesis

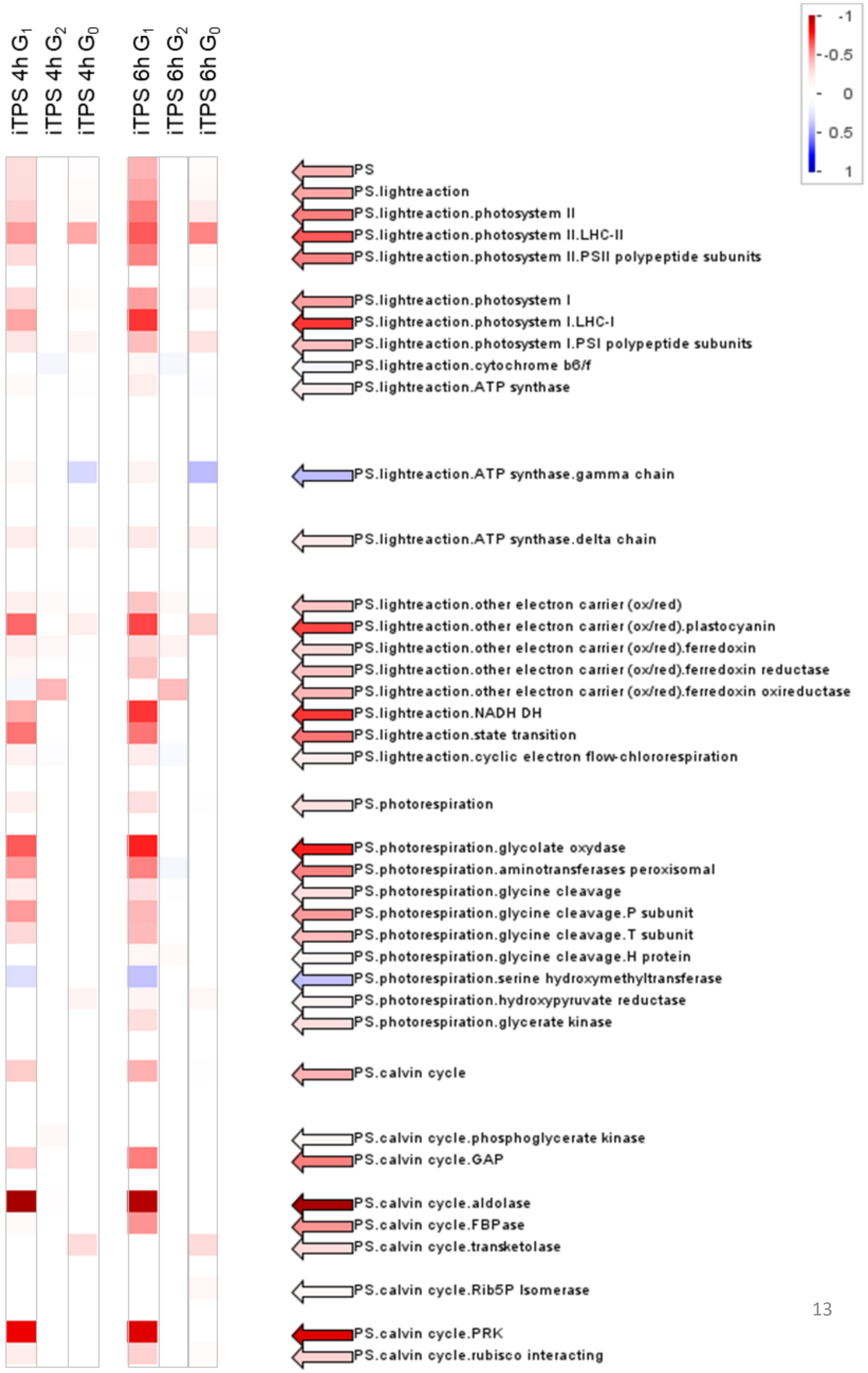

B. Gluconeogenesis/glyoxylate cycle

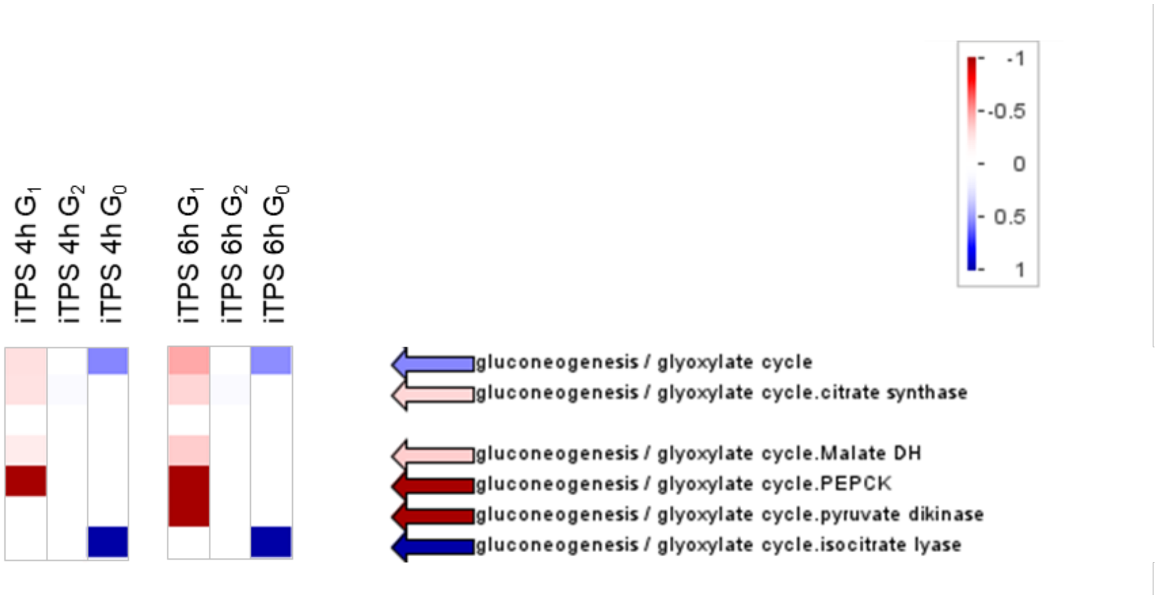

C. N metabolism

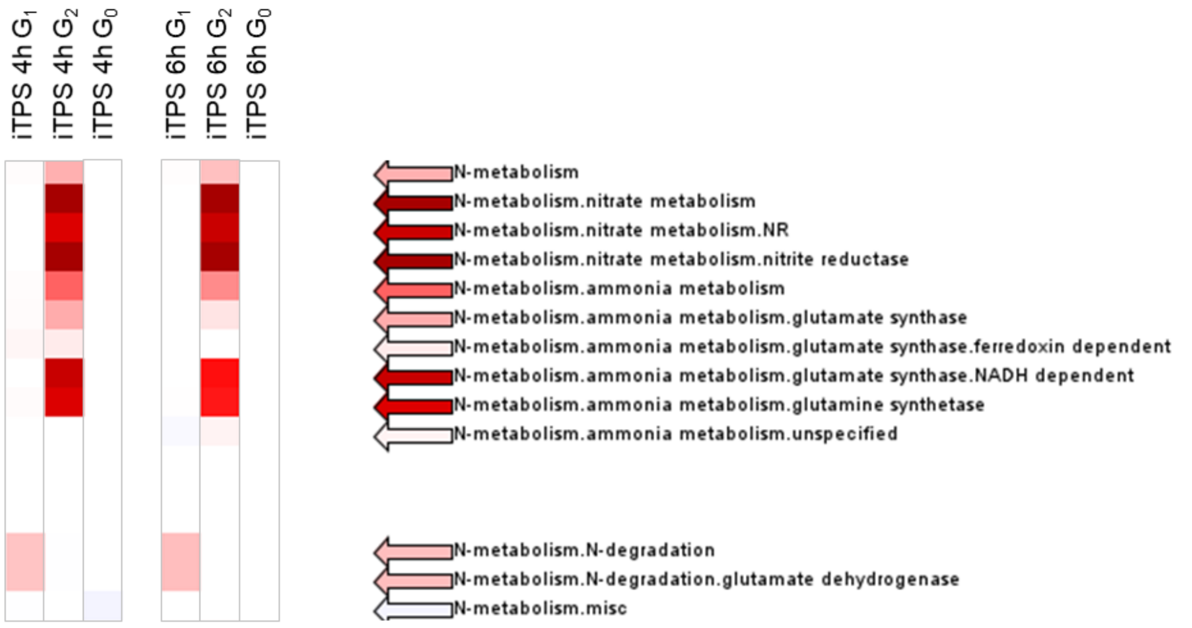

**Supplemental Figure S7, continued**

#### D. Nucleotide metabolism

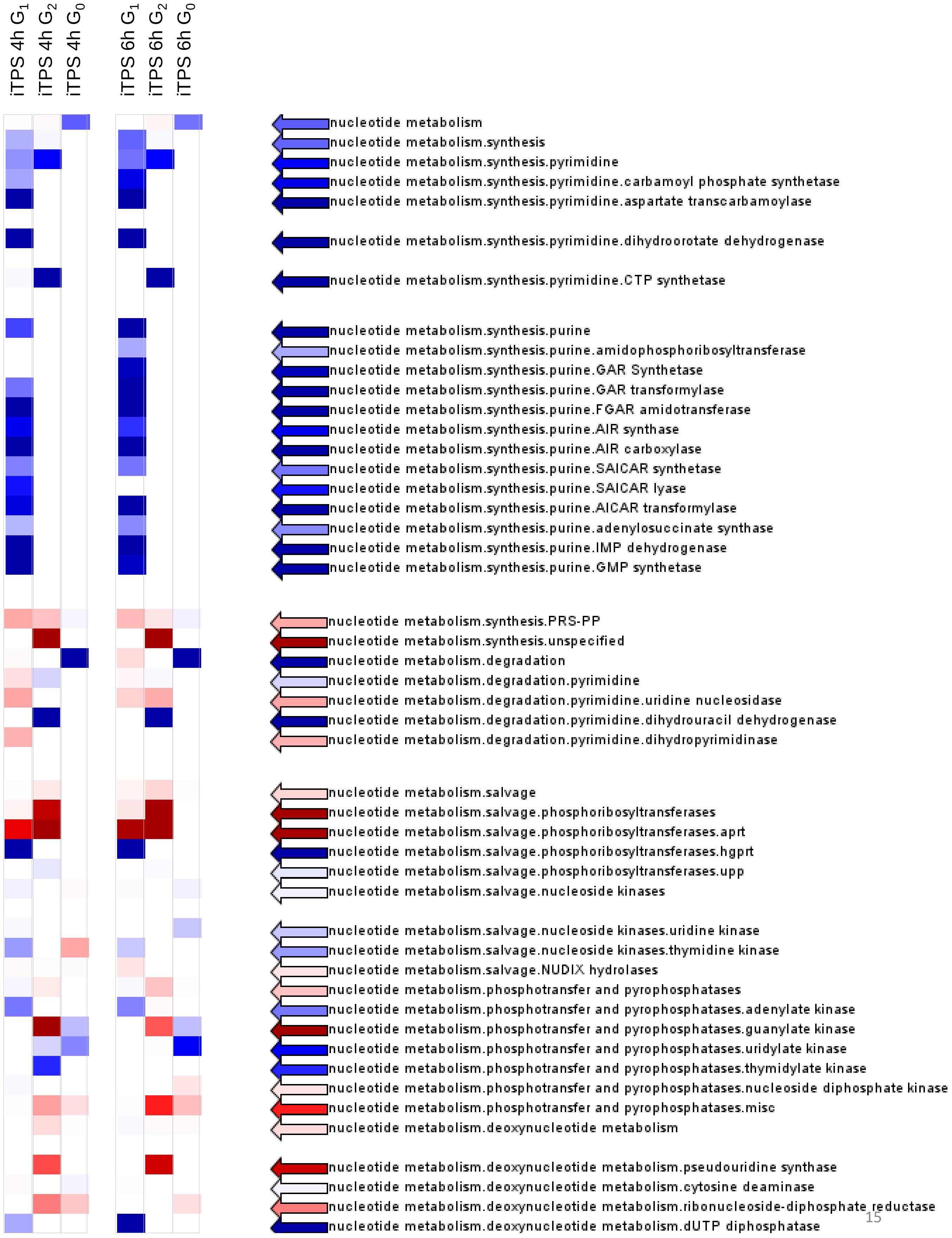

**Supplemental Figure S7, continued**

E. Specialized metabolism
(continued on next pages)

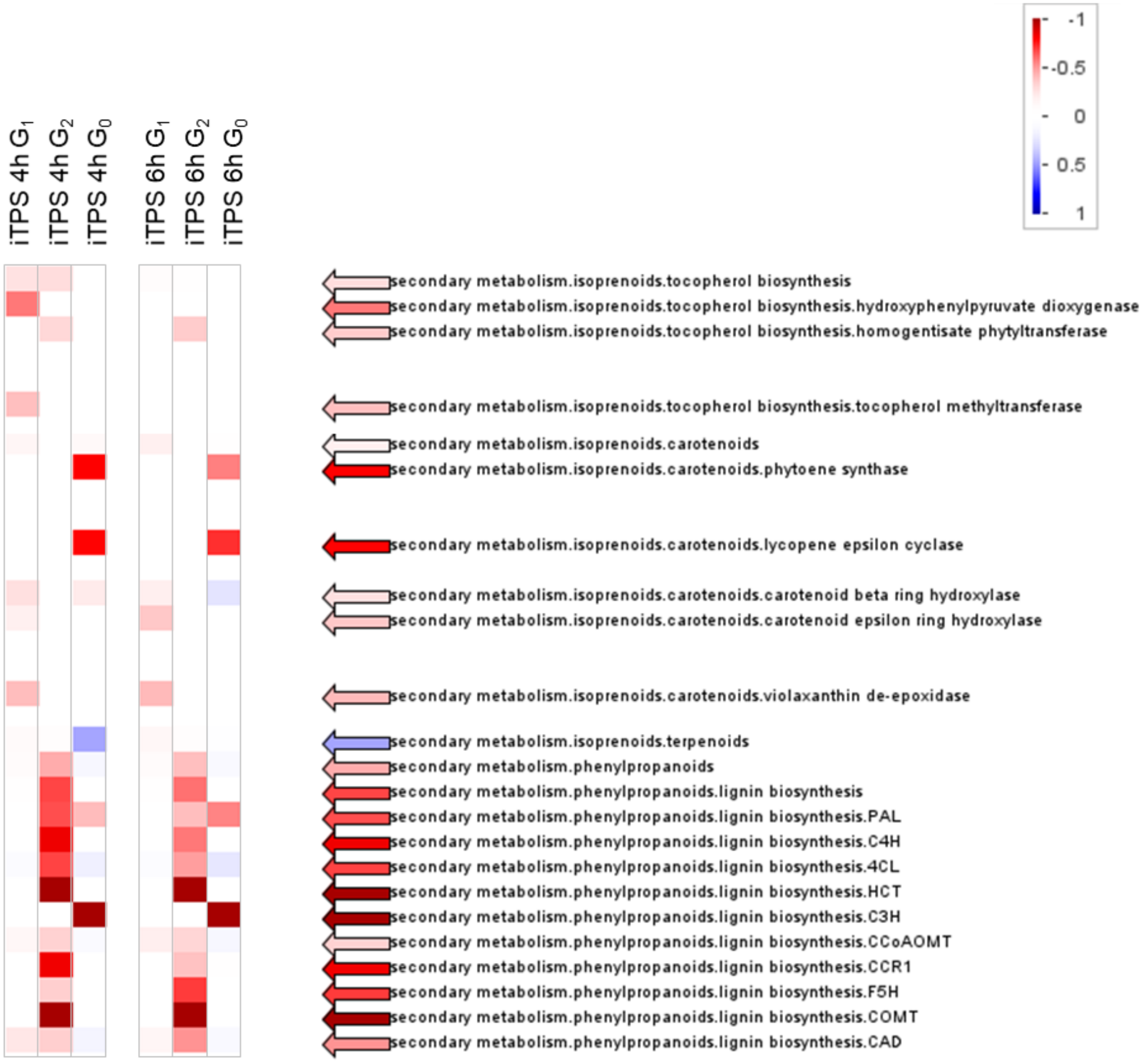

**Supplemental Figure S7, continued**

E. Specialized metabolism, continued
(mainly S-containing secondary metabolites)

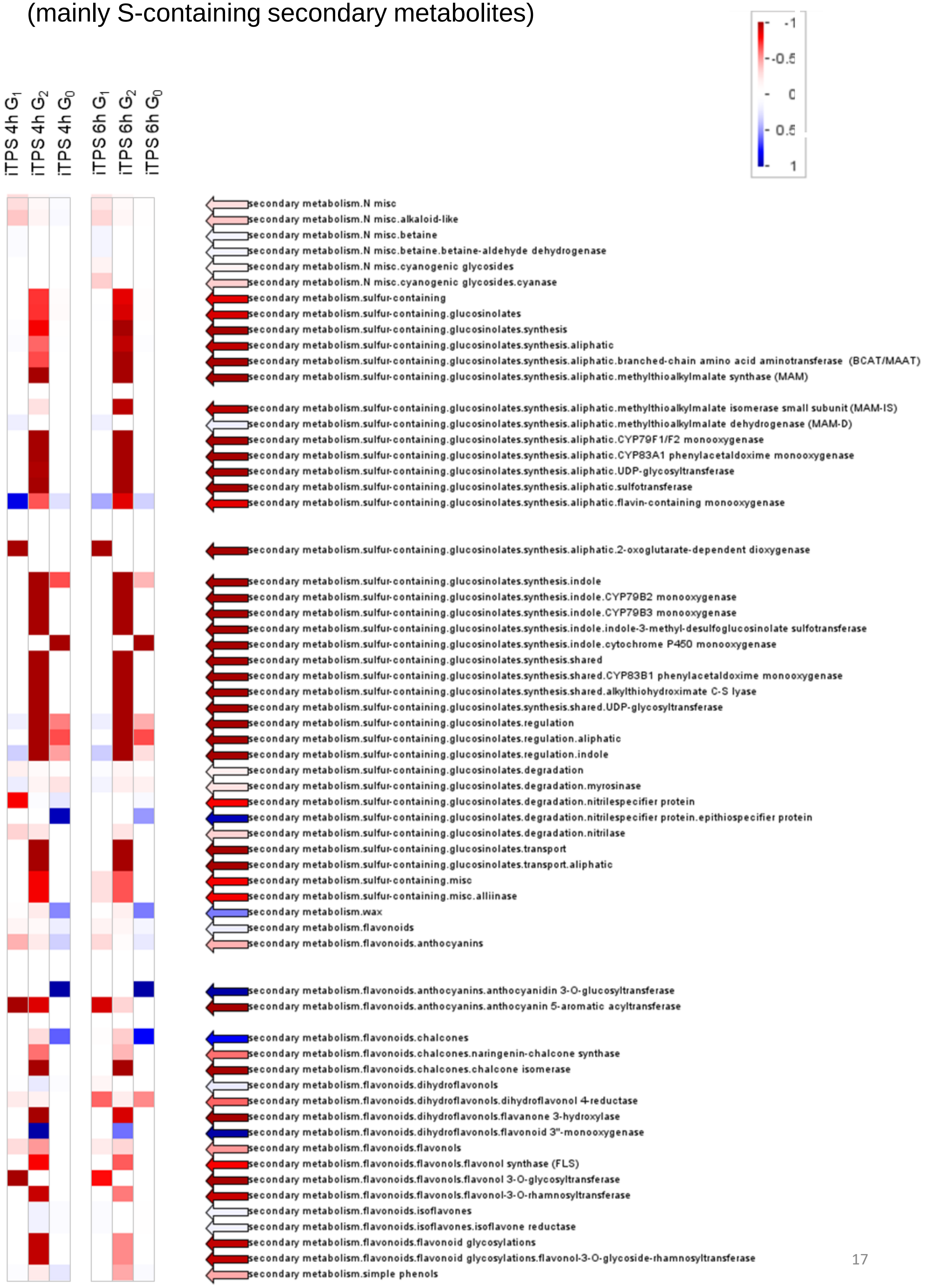

Supplemental Figure S7, continued

#### F. Protein, Chloroplast and mitochondrial ribosomal proteins

Genes annotated as prokaryotic-type ribosomal proteins.misc. (i.e., not assigned to the plastid or mitochondria) are omitted

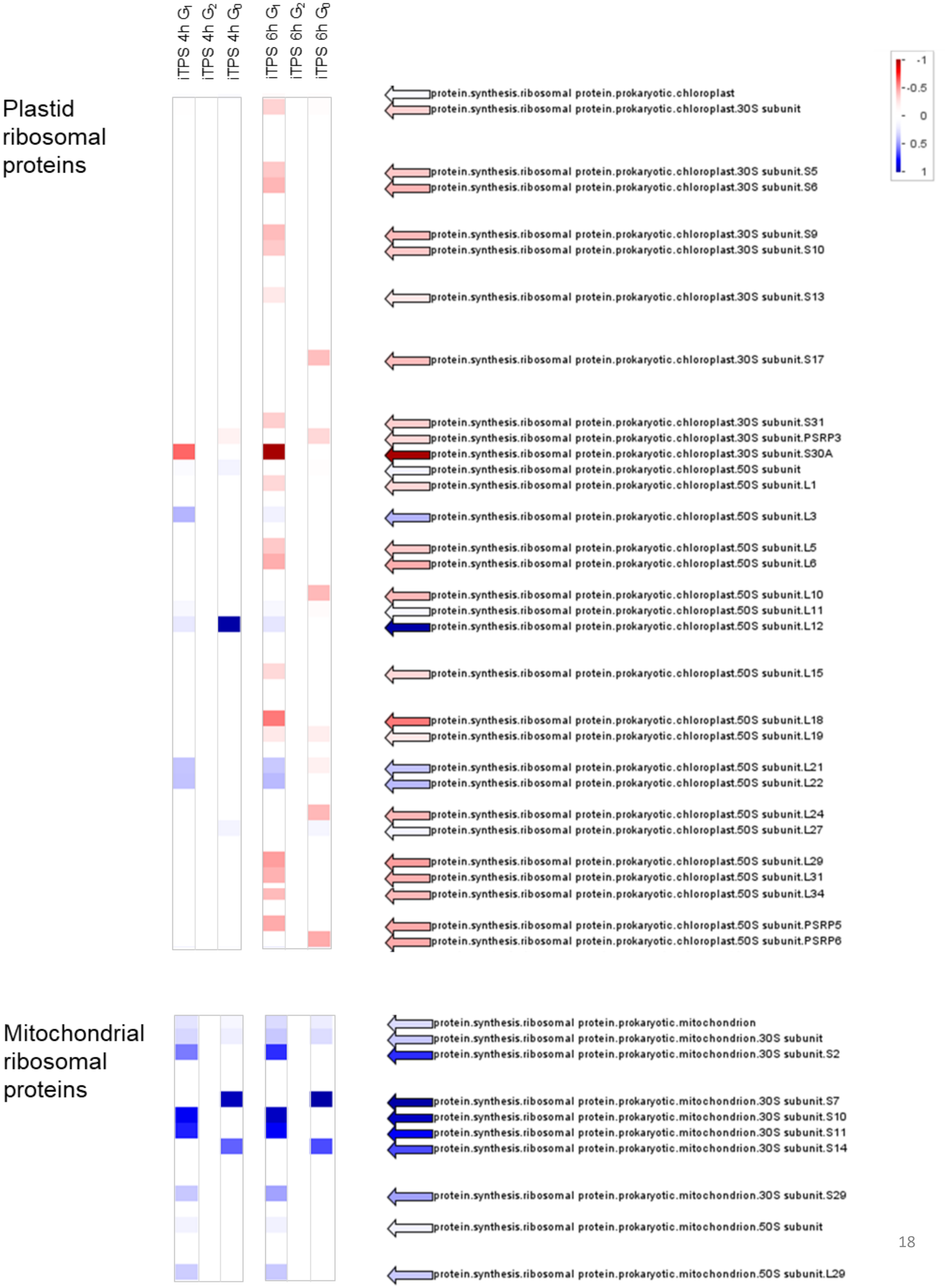

**Supplemental Figure S7, continued**

### F. Protein Continued

Eukaryotic (cytosolic) ribosomal proteins

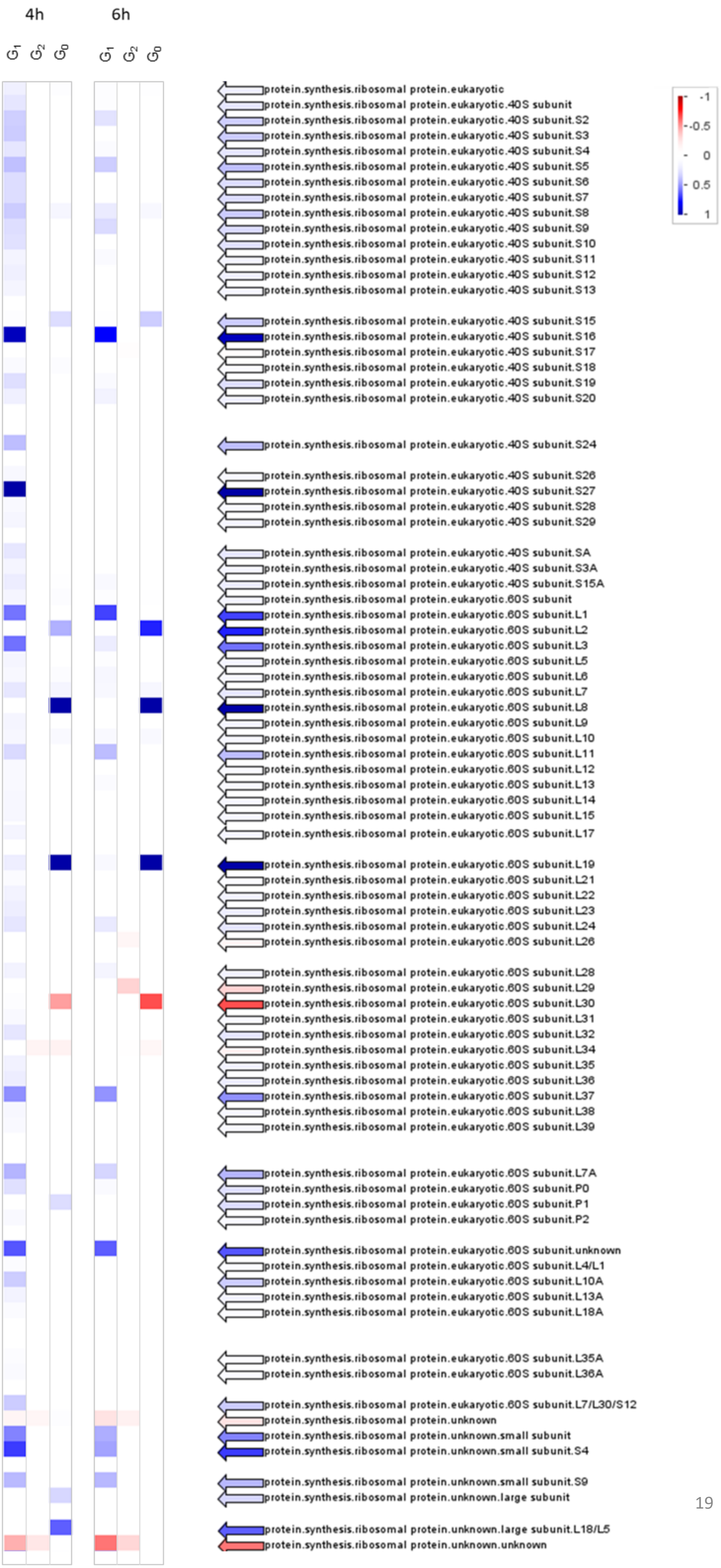

**Supplemental Figure S7, continued**

F.  Protein Continued

Protein
targeting

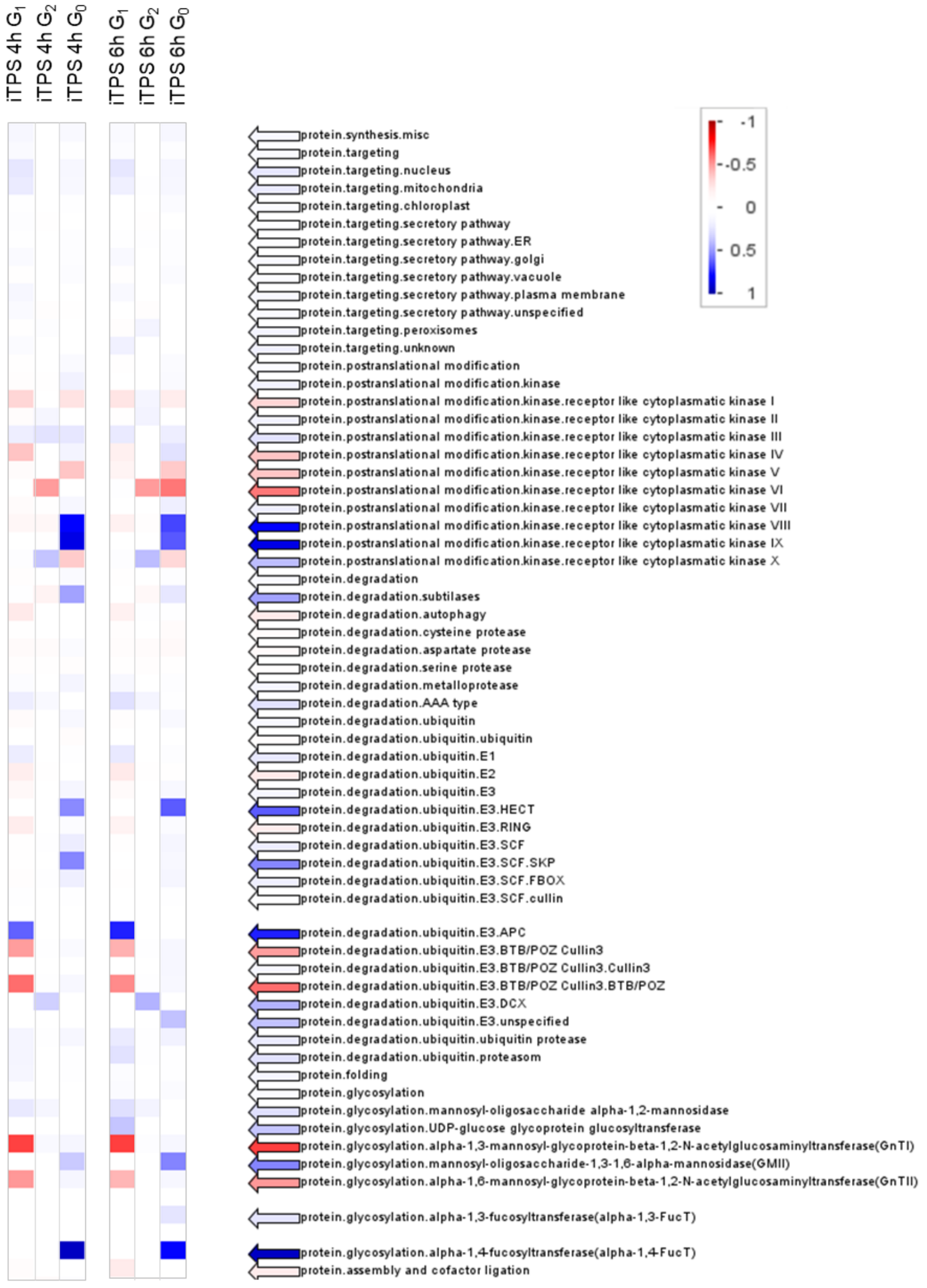

**Supplemental Figure S7, continued**

#### G. Cell Wall

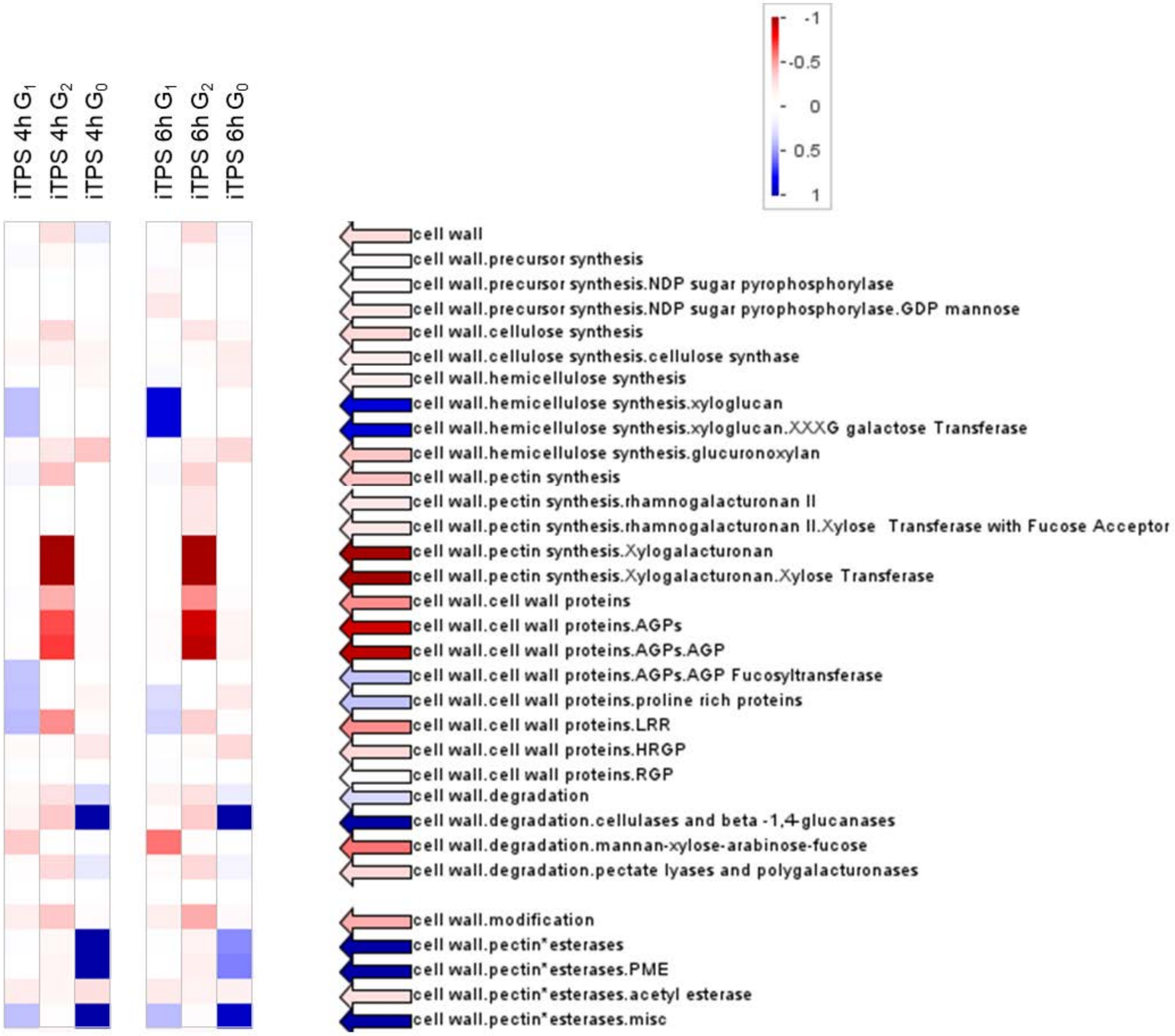

**Supplemental Figure S8. Responses of transcripts encoding proteins involved in sucrose transport and metabolism.**

Responses are shown separately for the 4-h and 6-h iTPS response, assignment to CRF groups is denoted by shading ($G_1$ orange, $G_2$ blue, $G_0$ white, no significant change white), numbers represent $log_2$ FC values and the direction of change is depicted by font color (blue for increase and red for decrease).

Information about the expression and function of SWEET family members is from Xue et al. (2022) and Braun (2022)

| | | iTPS response 4-h | 6-h 6h | Clade | Substrate, Expression, Function, |
|---|---|---|---|---|---|
| AT1G21460 | SWEET1 | -1.3 | -0.6 | I | hexoses, embryo, ABA control of germination |
| AT3G14770 | SWEET2 | +0.3 | +0.3 | I | hexoses, roots, vacuolar storage of hexoses |
| AT5G53190 | SWEET3 | | | I | hexoses, |
| AT3G28007 | SWEET4 | -0.8 | -1.4 | II | hexoses leaf veins, prevents chlorosis |
| AT5G62850 | SWEET5 | -1.1 | -0.9 | II | hexoses, galactose inhibition of pollen germination |
| AT1G66770 | SWEET6 | | | II | hexoses, |
| AT4G10850 | SWEET7 | | | II | hexoses, |
| AT5G40260 | SWEET8 | | | II | hexoses, tapetum, pollen development |
| AT2G39060 | SWEET9 | | | III | sucrose, flower, sucrose release in nectar formation |
| AT5G50790 | SWEET10 | | | III | sucrose, mature vascular tissue and shoot apex in flowering |
| AT3G48740 | SWEET11 | -1.9 | -2.1 | III | sucrose, phloem parenchyma, sucrose export |
| AT5G23660 | SWEET12 | -4.5 | -3.2 | III | sucrose, phloem parenchyma, sucrose export |
| AT5G50800 | SWEET13 | -2.8 | -1.2 | III | sucrose, bundle sheath, sucrose export , also tapetum |
| AT4G25010 | SWEET14 | | | III | sucrose, anther dehiscence and dehydration, GA signalling |
| AT5G13170 | SWEET15 | -1.6 | | III | sucrose, senescing tissues, |
| AT3G16690 | SWEET16 | -1.0 | -1.6 | IV | fructose, tonoplast of leaf cells, vacuolar storage |
| AT4G15920 | SWEET17 | -2.9 | -2.7 | IV | fructose, tonoplast of root cells, vacuolar storage |
| AT1G71880 | SUC1 | -0.6 | | | |
| AT1G22710 | SUT1/SUC2 | | -0.8 | | |
| AT2G02860 | SUT2/SUC3 | +1.0 | +1.0 | | |
| AT1G09960 | SUT4/SUC4 | | +0.4 | | |
| AT1G71890 | SUC5 | -3.2 | -2.0 | | |
| AT5G43610 | SUC6 | | | | |
| AT1G66570 | SUC7 | | | | |
| AT2G14670 | SUC8 | | | | |
| AT5G06170 | SUC9 | | | | |
| AT5G20830 | SUS1 | -3.2 | -2.0 | | |
| AT5G49190 | SUS2 | +0.8 | +0.8 | | |
| AT4G02280 | SUS3 | +1.6 | +1.8 | | |
| AT3G43190 | SUS4 | -1.6 | -2.7 | | |
| AT5G37180 | SUS5 | -0.8 | -1.0 | | |
| AT1G73370 | SUS6 | | -0.4 | | |
| AT1G62660 | VINV1 | | -0.6 | | |
| AT3G13790 | CWINV1 | +0.9 | +1.1 | | |
| AT3G52600 | CWINV2 | | | | |
| AT2G36190 | CWINV4 | | | | |
| AT3G13784 | CWINV5 | +3.5 | +2.5 | | |
| AT1G56560 | A/N-InvA | | +0.6 | | |
| AT4G34860 | A/N-InvB | | +0.5 | | |
| AT3G06500 | A/N-InvC | -1.4 | -0.9 | | |
| AT1G22650 | A/N-InvD | | +0.9 | | |
| AT5G22510 | A/N-InvE | -0.4 | -0.3 | | |
| AT1G72000 | A/N-InvF | | | | |
| AT1G35580 | A/N-InvG | | -0.9 | | |
| AT3G05820 | A/N-InvH | -1.3 | | | |
| AT4G09510 | A/N-InvI | +1.1 | +1.0 | | |
| AT5G20280 | SPS1 | | +0.4 | | |
| AT5G11110 | SPS2 | | +0.5 | | |
| AT1G04920 | SPS3 | +0.9 | +0.7 | | |
| AT4G10120 | SPS4 | -2.2 | -2.0 | | |

CRF group 1
CRF group 2
CRF group 0
No significant change

**Supplemental Figure S9. Responses of selected transcripts encoding proteins involved in the regulation of floral induction.**

Responses are shown separately for the 4-h and 6-h iTPS response, assignment to CRF groups is denoted by shading ($G_1$ orange, $G_2$ blue, $G_0$ white, no significant change white), numbers represent $\log_2$ FC values and the direction of change is depicted by font color (blue for increase and red for decrease)..

| | | | iTPS response 4-h | iTPS response 6-h |
|---|---|---|---|---|
| | AT4G20370 | TSF | | |
| | AT4G35900 | FD | -1.0 | -1.4 |
| | AT1G65480 | FT | | |
| | AT5G03840 | TFL1 | -2.0 | -2.0 |
| | AT5G06850 | FTIP1 | -2.0 | -2.0 |
| | AT5G10140 | FLC | | |
| Floral meristem identity | AT4G22950 | AGL19 | | |
| Floral meristem identity | AT5G60910 | FUL | -1.0 | -0.7 |
| Floral meristem identity | AT4G24540 | AGL24 | | |
| Floral meristem identity | AT5G61850 | LFY | +0.9 | |
| Floral meristem identity | AT1G69120 | AP1 | | |
| | AT2G45660 | SOC1 | -1.5 | -1.0 |
| | AT5G15840 | CO | +0.6 | |
| | AT2G33810 | SPL3 | -1.7 | -1.9 |
| | AT1G53160 | SPL4 | -1.0 | -1.0 |
| | AT3G15270 | SPL5 | -1.0 | |
| | AT2G42200 | SPL9 | -0.8 | -0.8 |
| | AT1G27370 | SPL10 | | |
| | AT2G22540 | SVP | -1.0 | -0.7 |
| | AT2G43010 | PIF4 | -2.4 | -2.0 |
| | AT4G31610 | REM1 | | |
| | AT5G17800 | MYB56 | | |
| | AT5G65070 | MAF4 | | |
| | AT5G65080 | MAF5 | | |
| | AT5G05660 | NFXL2 | +0.2 | |
| | AT5G57360 | ZTL | +0.3 | +0.4 |
| | AT5G61380 | PRR1 | | |
| | AT1G18080 | RACK1 | | |
| | AT3G22380 | TIC | -1.3 | -0.6 |
| flowering-time inhibitors – hormonal pathway | AT3G05120 | GID1A | -0.8 | -0.5 |
| flowering-time inhibitors – hormonal pathway | AT3G63010 | GID1B | | -0.8 |
| flowering-time inhibitors – hormonal pathway | AT5G27320 | GID1C | -1.0 | -0.7 |
| flowering-time inhibitors – hormonal pathway | AT1G25560 | TEM1 | -1.6 | -1.5 |
| flowering-time inhibitors – hormonal pathway | AT1G68840 | TEM2 | -2.3 | -2.1 |
| flowering-time inhibitors – hormonal pathway | AT2G01570 | RGA1 | -0.6 | -0.8 |
| flowering-time inhibitors – hormonal pathway | AT1G14920 | RGA2 | -2.0 | -1.7 |
| flowering-time inhibitors – hormonal pathway | AT1G66350 | RGL1 | -2.5 | -2.4 |
| flowering-time inhibitors – hormonal pathway | AT3G03450 | RGL2 | | +0.4 |
| flowering-time inhibitors – hormonal pathway | AT5G17490 | RGL3 | +0.3 | +0.4 |

CRF group 1
CRF group 2
CRF group 0
No significant change

**Supplemental Figure S10. Responses of transcripts encoding proteins involved in the circadian clock.**

Responses are shown separately for the 4-h and 6-h iTPS response, assignment to CRF groups is denoted by shading ($G_1$ orange, $G_2$ blue, $G_0$ white, no significant change white), numbers represent $log_2$ FC values and the direction of change is depicted by font color (blue for increase and red for decrease).

The right-hand display summaries the diel oscillation of these genes in plants grown in similar conditions, showing the fold change (FC $log_2$) between the diel maximum and the diel minimum, and the time of the peak (given as time after dawn, in hours). Transcripts were quantified using qRT-PCR with internal standards to allow absolute quantification. The data are taken from Moraes et al. (2019).

| | | iTPS response (FC $Log_2$) | |
|---|---|---|---|
| | | 4-h | 6-h |
| AT1G01060 | LHY | | +0.7 |
| AT2G46830 | CCA1 | | +0.6 |
| AT2G39180 | CCR2 | | -0.6 |
| AT2G46790 | PRR9 | -0.3 | -0.4 |
| AT5G02810 | PRR7 | | |
| AT5G24470 | PRR5 | -1.3 | -1.8 |
| AT5G61380 | TOC1 | | |
| AT1G22770 | GI | -0.6 | -0.7 |
| AT2G25930 | ELF3 | -0.4 | -0.5 |
| AT2G40080 | ELF4 | -1.9 | -1.5 |
| AT3G46640 | LUX | -0.7 | -1.2 |

| | Diel oscillation | |
|---|---|---|
| | FC $Log_2$ | Peak time h after dawn |
| LHYC | 9.7 | 23.7 |
| CCA1 | 9.4 | 23.9 |
| PRR9 | 11.0 | 4.2 |
| PRR7 | 7.4 | 5.9 |
| PRR5 | 7.9 | 7.9 |
| TOC1 | 3.9 | 10.9 |
| GI | 8.2 | 7.6 |
| ELF3 | 2.9 | 16.3 |
| ELF4 | 8.9 | 10.3 |
| LUX | 4.7 | 9.7 |

CRF group 1
CRF group 2
CRF group 0
No significant change

**Supplemental Figure S11. Enrichment analyses using Gene Ontology.** (A) List of enriched and underrepresented GO terms. The analysis was performed by the Gene Ontology tool (http://geneontology.org/). Entries were restricted to transcripts showed a significant response (FDR <0.05) and showed a FC≥2. A more extensive analysis with transcripts that showed a significant change in abundance (FDR < 0.05) irrespective of the size of the FC is provided in Supplemental Dataset S5 .

| CRF group | enrichment UP | enrichment DOWN |
|---|---|---|
| $G_1$<br>Up- 307<br>Down-840 | • mitochondrial RNA modification<br>• mitochondrial gene expression<br>• RNA splicing<br>• ribonucleoprotein complex biogenesis | • negative regulation of cytokinin-activated signaling pathway<br>• trehalose metabolism in response to stress<br>• abaxial cell fate specification<br>• phototropism<br>• entrainment of circadian clock<br>• water transport<br>• response to nitrate<br>• response to far red light<br>• response to light intensity<br>• ethylene-activated signaling pathway<br>• response to red light<br>• response to nematode<br>• response to gibberellin<br>• regulation of stomatal movement<br>• regulation of seedling development<br>• response to auxin<br>• response to chitin<br>• response to water deprivation<br>• response to jasmonic acid<br>• response to abscisic acid<br>• cellular response to oxygen-containing compound<br>• developmental growth<br>• organic substance catabolic process<br>• protein transport<br>• intracellular transport<br>• mRNA processing<br>• DNA repair<br>• translation<br>• meiotic cell cycle process |
| $G_2$<br>Up –180<br>Down - 756 | • regulation of seed dormancy process<br>• hyperosmotic salinity response<br>• response to salicylic acid<br>• response to water deprivation<br>• response to abscisic acid<br>• cellular response to hormone stimulus<br>• signal transduction | • arsenite transport<br>• adenine salvage<br>• indoleacetic acid biosynthetic process<br>• guard mother cell differentiation<br>• nitrate assimilation<br>• regulation of glucosinolate biosynthetic process<br>• syncytium formation<br>• L-phenylalanine biosynthetic process<br>• cutin biosynthetic process<br>• wax biosynthetic process<br>• plant-type cell wall loosening<br>• oligopeptide transport<br>• brassinosteroid biosynthetic process<br>• auxin polar transport<br>• response to nitrate<br>• lateral root formation<br>• amino acid transmembrane transport<br>• flavonoid metabolic process<br>• fatty acid biosynthetic process<br>• response to gibberellin<br>• jasmonic acid mediated signaling pathway<br>• developmental growth<br>• regulation of DNA-templated transcription<br>• protein modification process<br>• translation<br>• rRNA processing<br>• establishment of protein localization to organelle |
| $G_0$<br>Up – 710<br>Down - 599<br><br>*under-represented processes | • cytidine deamination<br>• geranylgeranyl diphosphate biosynthetic process<br>• regulation of pollen tube growth<br>• pectin catabolic process<br>• RNA modification<br>• protein phosphorylation<br>• gene expression<br>• response to light stimulus<br>• cellular nitrogen compound biosynthetic process | • xylem vessel member cell differentiation<br>• formation of plant organ boundary<br>• asymmetric cell division<br>• cotyledon development<br>• cell wall modification<br>• response to cytokinin<br>• response to auxin<br>• glucosinolate metabolic process<br>• response to red or far red light<br>• meristem development<br>• positive regulation of DNA-templated transcription<br>• response to light intensity<br>• developmental growth<br>• response to abscisic acid<br>• root development<br>• response to wounding<br>• translation<br>• ribosome biogenesis<br>• ncRNA processing |

**Supplemental Figure S12. Comparison of the changes in transcript abundance an induced increase in Tre6P, and in Arabidopsis lines that constitutively express bacterial TPS.**(supplemental to Figure 6 and Supplemental Table S5. The response to constitutive overexpression of TPS (termed oeTPS) is from Zhang et al. (2009) was analyzed using a CATMA array; responsive transcripts were defined using a filter of FDR>0.05 and FC≥2.

(A) Regression plots of oeTPS against the iTPS response of the corresponding gene, either without filtering the iTPS data or after filtering the iTPS data sets with FDR <0.05 or with FDR<0.05 and FC≥2 (left, middle and right side, respectively). Plots are shown separately for the 4h (top row) and the 6h (bottom row) iTPS responses. The number in brackets in the panel heading gives the number of transcripts in the subset plotted in the panel.

(B) Regression plots of oeTPS (Zhang et al., 2009) against the iTPS response after filtering (FDR<0.05) and assigning genes to the CRF groups $G_1$, $G_2$ or $G_0$.

(C) Comparative analysis of the response of genes involved in light–signaling. Shown on next page.

(D) Gene ontology analysis of a robust set of Tre6P-regulated genes that show significant iTPS responses, were assigned to CRF group $G_1$ and were scored as responsive to oeTPS by Zhang et al. (2009) and showed the same qualitative response as in our iTPS dataset. Shown on next page.

A oeTPS versus iTPS, All shared genes, unfiltered or filtered on FDR or FDR plus FC

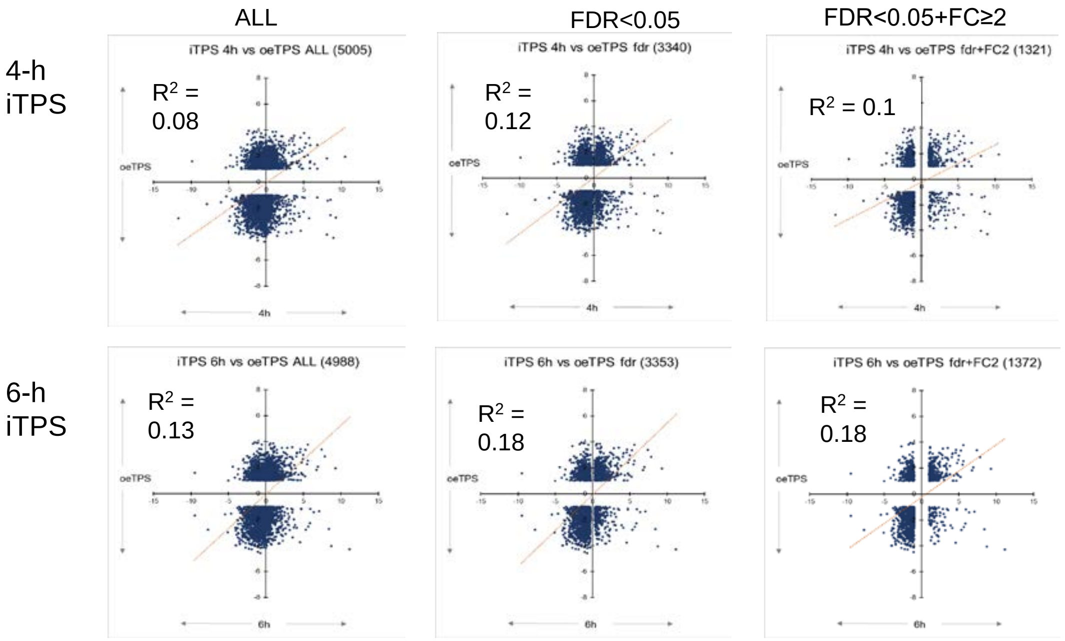

B. oeTPS versus iTPS, after assigning DEGs to $G_1$, $G_2$ and $G_O$

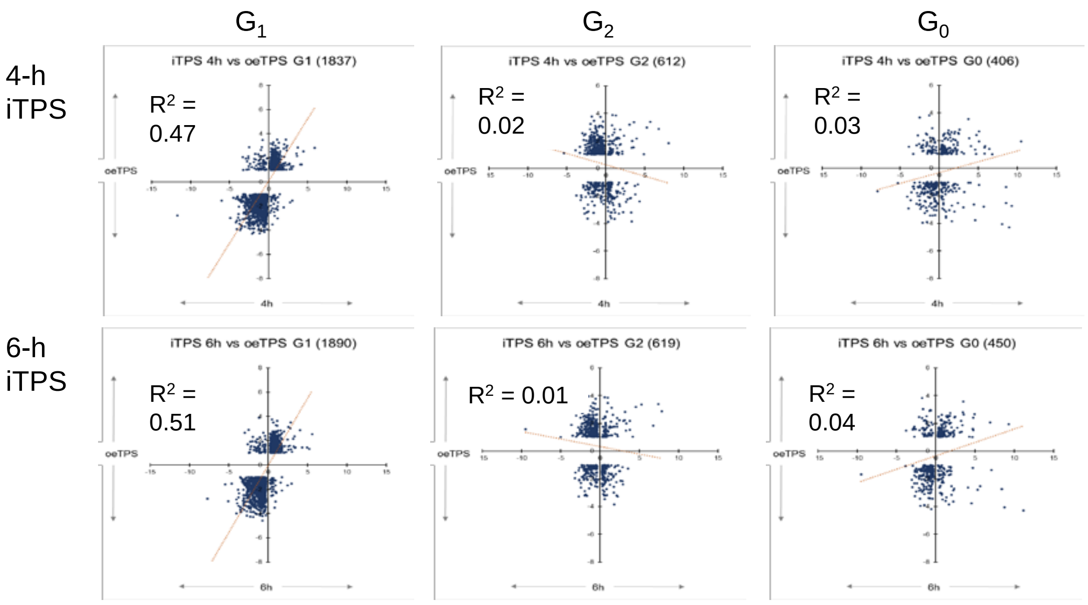

### Supplemental Figure S12 (continued)

Comparison of the response of genes involved in light–signaling to iTPS and oeTPS. The upper part of the display investigates genes highlighted as involved in light signaling and being repressed by constitutive overexpression of TPS (oeTPS) by Paul et al. (2010) (changes are given for the ratio to control as in Paul *et al* (2010) and the $\log_2$ FC as in Zhang et al. (2009). These genes show qualitatively similar responses in the iTPS response, and are almost all assigned to CRF $G_1$ (orange shaded) implying they are regulated by Tre6P-signaling. Four highlighted genes were absent from the iTPS dataset, and the absence of a CRF assignment denotes a non-significant iTPS response. The lower Table section investigates genes involved in light signaling that showed a strong iTPS response ($G_1$ or $G_0$) but were not called in the Zhang et al. (2009) data set, or genes that were strongly induced in iTPS and based on extracted data were also induced in the oeTPS dataset from Zhang et al.(2009). Data for *HY5* was also included. The insert (top right-hand corner) is a scatter plot of the oeTPS versus iTPS for the entire gene set. . .

**C**

| | oeTPS response | | Qualitative agreement of oeTPS and iTPS | iTPS response | | | | CFR | Transient SnRK1α1 | Ontology | |
|---|---|---|---|---|---|---|---|---|---|---|---|
| | Table 1, Paul et al. (2020) Ratio to control, direction of change UP DOWN | **Zhang et al 2009 oeTPS FC $\log_2$** | 1= same 0 = opposite na = not available | 4-h FC $\log_2$ | 6-h FC $\log_2$ | 4h Crf score | 6h Crf score | | FC $\log_2$ | **BINNAME_1** | **DESCRIPTION** |
| At1g49130 | 0.08 | | na | | | | | | | | |
| At1g75540 StH2 | 0.10 | -3.30 | 1 | -2.05 | -1.69 | 1 | 1 | -0.26 | 0.51 | RNA.regulation of transcript | (at1g75540):zinc finger (B-box type |
| At3g19850 | 0.10 | -3.34 | 1 | -4.19 | -3.17 | 1 | 1 | -0.31 | 0.97 | signalling.light | (at3g19850):phototropic-responsiv |
| At5g52250 | 0.12 | -3.06 | 1 | -2.30 | -1.54 | 1 | 1 | -0.95 | 3.11 | development.unspecified | (at5g52250):transducin family prot |
| At3g26740 ccL | 0.12 | -3.07 | 1 | -3.34 | -3.03 | 1 | 1 | -1.41 | 3.02 | signalling.light | (at3g26740):CCL (CCR-LIKE) |
| At3g02380 cOL2 | 0.12 | -3.38 | 1 | -1.83 | -0.36 | 1 | 1 | -0.64 | 2.47 | development.unspecified | (at3g02380):COL2 (CONSTANS-LIKE 2); zinc ion binding |
| At1g18330 ePr1 | 0.12 | -3.10 | 1 | -1.17 | -1.31 | 1 | 1 | -1.45 | 0.8 | RNA.regulation of transcript | (at1g18330):EARLY-PHYTOCHROME-RESPONSIVE1 |
| At5g37260 cir1 | 0.13 | -2.92 | 1 | -1.58 | -1.69 | 1 | 1 | -1.32 | 2.54 | RNA.regulation of transcript | (at5g37260):myb family transcription factor CIR/RVE2 |
| At3g45780 nPH1 | 0.14 | -2.86 | 1 | -2.61 | -2.30 | 1 | 1 | -0.35 | 1.86 | signalling.light | (at3g45780):PHOT1 (phototropin 1); kinase |
| At5g17300 | 0.15 | -2.74 | 1 | -1.09 | -0.39 | 1 | 1 | -1.86 | 2.88 | RNA.regulation of transcript | (at5g17300):myb family transcription factor |
| At3g15570 | 0.16 | -2.64 | 1 | -1.56 | -0.94 | 1 | 1 | -0.14 | 0.25 | signalling.light | (at3g15570):phototropic-responsive NPH3 family protein |
| At1g14280 | 0.17 | -2.48 | 1 | -1.63 | -1.40 | 1 | 1 | -0.26 | -0.2 | signalling.light | (at1g14280):PKS2 (PHYTOCHROME KINASE SUBSTRATE 2) |
| At3g09150 | 0.21 | -2.27 | 1 | -0.85 | -0.83 | 2 | 2 | 0.10 | 0.18 | PS.lightreaction.other electr | (at3g09150):HY2 (ELONGATED HYPOCOTYL 2); |
| At5g66560 | 0.22 | -2.15 | 1 | -0.69 | -0.62 | 2 | 2 | 0.18 | -0.17 | signalling.light | (at5g66560):phototropic-responsive NPH3 family protein |
| At1g67900 | 0.25 | | na | | | | | | | | |
| At5g67385 | 0.26 | | na | | | | | | | | |
| At2g43010 PiF4 | 0.27 | -1.7 | 1 | -2.43 | -1.98 | 1 | 1 | -0.31 | 0.13 | signalling.light | (at2g43010):PIF4 (PHYTOCHROME INTERACTING FACTOR 4) |
| At3g50840 | 0.27 | | na | | | | | | | | |
| At3g22104 | 0.27 | -1.87 | 1 | -0.92 | -0.60 | 1 | 1 | -0.10 | -1 | signalling.light | (at3g22104):phototropic-responsive NPH3 protein-related |
| At2g02950 | 0.28 | -1.82 | 1 | -2.99 | -2.65 | 1 | 1 | -0.80 | -0.23 | signalling.light | (at2g02950):PKS1 (PHYTOCHROME KINASE SUBSTRATE 1) |
| At4g31820 enP | 0.29 | -1.78 | 1 | -1.53 | -1.31 | 1 | 1 | -0.72 | -0.06 | signalling.light | (at4g31820):phototropic-responsive NPH3 family protein |
| At1g18810 | 0.37 | -1.70 | 1 | -3.48 | -3.00 | 1 | 1 | -0.80 | 2.53 | signalling.light | (at1g18810):phytochrome kinase substrate-related |
| At4g37590 nPY5 | 0.42 | -1.24 | 1 | -0.57 | -0.57 | 1 | 1 | -0.24 | 1.42 | signalling.light | (at4g37590):phototropic-responsive NPH3 family protein |
| | | | | **Strong in iTPS but not called in Zhang et al. 2010** | | | | | | | |
| | | 0 | na | -1.93 | -1.47 | 1 | 1 | -0.17 | 0.81 | signalling.light | (at2g40080):ELF4 (EARLY FLOWERING 4) |
| | | 0 | na | -1.54 | -0.55 | 1 | 1 | -0.29 | 0.04 | signalling.light | (at1g52770):phototropic-responsive NPH3 family protein |
| | | 0 | na | -3.57 | -3.61 | 0 | 0 | 0.01 | 0.235 | signalling.light | (at5g10250):phototropic-responsive protein, putative |
| | | 0 | na | -3.24 | -2.85 | 0 | 0 | 0.05 | -0.145 | signalling.light | (at3g08660):phototropic-responsive protein, putative |
| | | 0 | na | -2.75 | -3.51 | 0 | 0 | -0.04 | -0.55 | signalling.light | (at5g04190):PKS4 (PHYTOCHROME KINASE SUBSTRATE 4) |
| | | 0 | na | 2.13 | 0.72 | 0 | 0 | 0.09 | -0.395 | signalling.light | (at2g47860):phototropic-responsive NPH3 family protein |
| | | 0 | na | 4.21 | 3.71 | 0 | 0 | 0.05 | 0.03 | signalling.light | (at2g23050):phototropic-responsive NPH3 family protein |
| | | | | **Respond in Zhang et al. (2009) but not in Table in Paul et al. 2010** | | | | | | | |
| | | 1.34 | 1 | 0.59 | 0.42 | 1 | 1 | 0.35 | -0.425 | signalling.light.COP9 signal( | (at5g14250):COP13 (CONSTITUTIVE PHOTOMORPHOGENIC 13) |
| | | 1.41 | 1 | 0.61 | 0.64 | 1 | 1 | 0.42 | -0.575 | signalling.light.COP9 signal( | (at5g42970):COP8 (CONSTITUTIVE PHOTOMORPHOGENIC 8) |
| | | 1.53 | 1 | 0.91 | 0.96 | 1 | 1 | 0.48 | -1.085 | signalling.light.COP9 signal( | (at3g61140):FUS6 (FUSCA 6) |
| | | 1.49 | na | 0.02 | -0.11 | | | 0.27 | -0.6 | signalling.light.COP9 signal( | (at1g71230):AJH2 (COP9-signalosome 5B); protein binding |
| | | 1.16 | 1 | 0.17 | 0.30 | | 1 | 0.13 | -0.365 | signalling.light.COP9 signal( | (at1g22920):AJH1 (COP9-signalosome 5A) |
| HY5 | | -1.09 | 1 | -1.02 | -0.99 | 1 | **1** | -0.67 | -0.41 | RNA.regulation of transcript | (at5g11260):HY5 (ELONGATED HYPOCOTYL 5); |

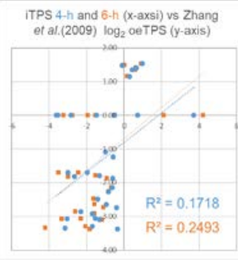

**D** Gene ontology analysis of a robust set of Tre6P-regulated genes that show significant iTPS responses, were assigned to CRF group $G_1$ and were also scored as responsive to oeTPS by Zhang et al. (2009).

Shared up

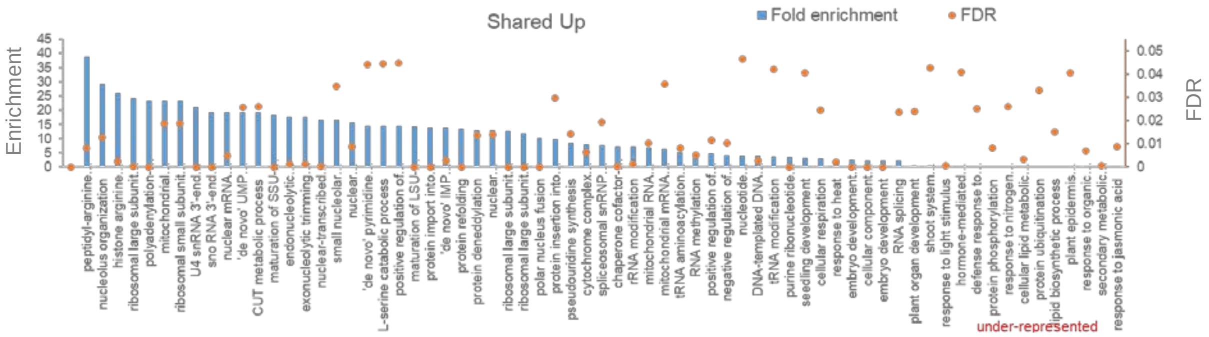

Shared down

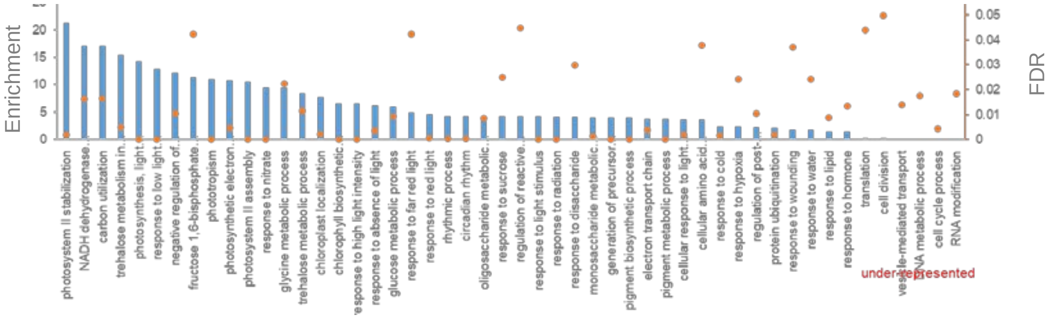

**Supplemental Figure S13. Responses of transcripts encoding proteins involved in Tre6P metabolism.** Responses are shown separately for the 4-h and 6-h iTPS response, assignment to CRF groups is denoted by shading ($G_1$ orange, $G_2$ blue, $G_0$ white, no significant change white), numbers represent $\log_2$ FC values and the direction of change is depicted by font color (blue for increase and red for decrease). A blank denotes that the transcript did not show a significant change (FDR<0.05). The panel shows the response of 11 trehalose phosphate synthase (TPS) and 10 trehalose phosphate phosphatase (TPP) gene family members. Note that TPS1 is responsible for Tre6P synthesis, TPS2-4 are expressed only in seeds and are in part inactive, and TPS5-11 are so-called TPS class II proteins that have no catalytic activity (see Introduction).

| | | iTPS response | | |
|---|---|---|---|---|
| | | 4-h | 6-h | 6h |
| AT1G78580 | TPS1 | -1.1 | -1.2 | |
| AT1G16980 | TPS2 | | | |
| AT1G17000 | TPS3 | | | |
| AT4G27550 | TPS4 | | | |
| AT4G17770 | TPS5 | -0.9 | -0.6 | |
| AT1G68020 | TPS6 | -0.4 | -0.4 | |
| AT1G06410 | TPS7 | | | |
| AT1G70290 | TPS8 | -1.5 | -1.1 | |
| AT1G23870 | TPS9 | -1.3 | -1.4 | |
| AT1G60140 | TPS10 | -1.2 | -1.0 | |
| AT2G18700 | TPS11 | -1.9 | -1.8 | |
| AT5G51460 | TPPA | -2.0 | -1.6 | |
| AT1G78090 | TPPB | -3.0 | -2.4 | |
| AT1G22210 | TPPC | | | |
| AT1G35910 | TPPD | -1.4 | -1.2 | |
| AT2G22190 | TPPE | -3.0 | -2.3 | |
| AT4G12430 | TPPF | | | |
| AT4G22590 | TPPG | +0.8 | +0.9 | |
| AT4G39770 | TPPH | -3.9 | -2.5 | |
| AT5G10100 | TPPI | | | |
| AT5G65140 | TPPJ | +0.7 | +0.5 | |

CRF group 1
CRF group 2
CRF group 0
No significant change

**Supplemental Figure S14. Responses of transcripts encoding SnRK1 subunits, or implicated as downstream transcriptional targets of SnRK1 signaling.**

(A) Responses of transcripts of genes encoding for SnRK1 structural subunits including catalytic subunits (SnRL1α1, SnRK1α2) and regulatory subunits (snf4 βγ, SnRK1β1, SnRK1β2, SnRK1β3).

(B) Responses of transcript for genes reported to be top 30 downstream targets of SnRK1 signaling (based on transient overexpression of SnRL1α1 in protoplasts, Baena-Gonzales *et al.*, 2007)

In panels A and B, responses are shown separately for the 4-h and 6-h iTPS response, assignment to CRF groups is denoted by shading ($G_1$ orange, $G_2$ blue, $G_0$ white, no significant change white), numbers represent $log_2$ FC values and the direction of change is depicted by font color (blue for increase and red for decrease).

(C) Comparison of the response to transient overexpression of SnRL1α1 in protoplast (tSnRL1α1 response) and the iTPS response of transcripts assigned to CRF groups $G_2$ and $G_0$, shown exemplarily for the iTPS 6h response The response for transcripts assigned to $G_1$ is shown in Figure 8B). (Supplemental to Figure 8 and to Supplemental Table S6).

See next pages for panels D-J

A.

| | | iTPS response 4-h | iTPS response 6-h |
|---|---|---|---|
| AT3G01090 | SnRK1 α1 | +0.2 | +0.3 |
| AT3G29160 | SnRK1 α2 | +0.2 | +0.5 |
| AT1G09020 | snf4 βγ | +0.5 | +0.6 |
| AT5G21170 | SnRK1 β1 | -0.9 | -1.1 |
| AT4G16360 | SnRK1 β2 | +0.3 | +0.3 |
| AT2g28060 | SnRK1 β3 | +0.3 | |

CRF group 1
CRF group 2
CRF group 0
No significant change

B.

| | | | iTPS response 4-h | iTPS response 6-h |
|---|---|---|---|---|
| Induced by transient overexpression of KIN10 | AT3G06850 | DIN3 | | |
| | AT3G47340 | DIN6 | +2.2 | +0.8 |
| | AT5G20250 | DIN10 | -1.0 | -1.8 |
| | AT3G15450 | SEN5 | | -1.0 |
| | AT5G28770 | bZIP63 | -1.5 | -1.6 |
| | AT2G33830 | AXP | -1.5 | -4.1 |
| | AT4G35770 | DIN1 | | -2.3 |
| | AT5G49450 | bZIP1 | -1.6 | -1.9 |
| | AT5G18630 | LIP | -1.3 | -0.8 |
| | AT4G15530 | PPDK | | -1.2 |
| | AT1G12240 | VIn2 | -2.4 | -2.8 |
| | AT1G70290 | TPS8 | -1.5 | -1.1 |
| | AT1G12780 | UDG | -1.9 | -2.0 |
| | AT5G22920 | PGPD14 | -1.2 | -2.3 |
| repressed by transient overexpression of KIN10 | AT4G34590 | bZIP11 | -0.4 | |
| | AT1G26770 | EXP10 | +0.6 | +1.6 |
| | AT3G50660 | DWF4 | -2.4 | -1.8 |
| | AT1G19640 | JMT | +1.3 | +0.5 |
| | AT1G58360 | AAP1 | +0.4 | +0.4 |
| | AT2G36910 | MDR1 | -0.7 | |
| | AT2G20610 | SUR1 | -1.3 | -1.4 |
| | AT3G09820 | ADK1 | | |
| | AT3G44750 | HDT1 | +0.6 | +0.5 |
| | AT5G60690 | IFL1 | -0.9 | -0.6 |
| | AT3G28910 | MYB30 | -1.7 | -1.2 |
| | AT1G56650 | MYB75 | +0.7 | +1.5 |
| | AT3G21500 | DPS | | |
| | AT3G28900 | L34 | | |
| | AT1G15250 | L37 | +1.3 | +1.6 |

C.

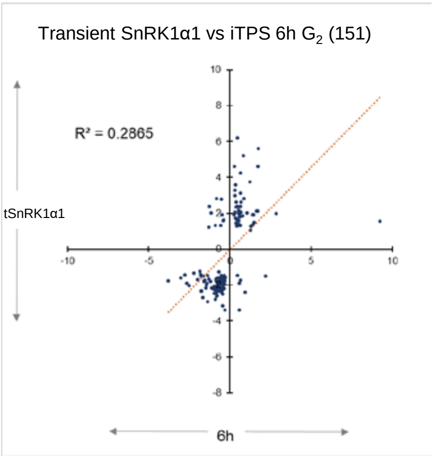

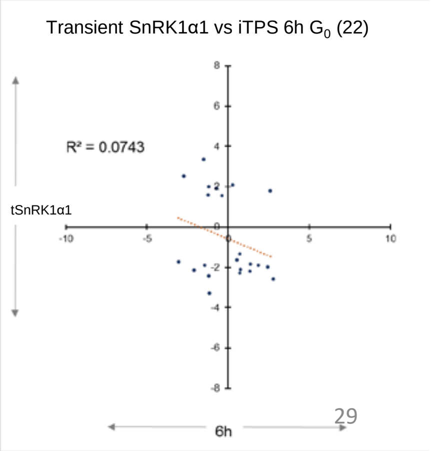

**Supplemental Figure S14. Responses of transcripts encoding SnRK1 subunits, or implicated as downstream transcriptional targets of SnRK1 signaling (continued)**

(C) Further comparison of the tSnRK1α1 response and the iTPS response. The analysis uses the top 100 (50 most strongly upregulated, 50 most strongly down-regulated) transcripts in the tSnRK1α1 dataset. The left-hand panels show the regression between the published tSnRK1α1 response and the4-h (top) and 6-h (bottom) iTPS responses for all 100 transcripts. Of the 100 transcripts in the tSnRK1α1 dataset, 52 and 72 were retrieved in the 4-h and 6-h iTPS responses, respectively, after filtering at FDR<0.05. Regressions of the published tSnRK1α1 response against the iTPS responses of these transcripts are shown in the middle panels). The 100 transcripts were also filtered based on their iTPS response (FDR>0.05, $log_2FC \geq 0.2$) and compared with the CRF (Supplemental Figure S2) to assign transcript to the GRF group $G_1$ (i.e.., transcripts whose iTPS response qualitatively similar to their CRF, and is probably due to a direct effect of elevated Tre6P). A total of 52 and 49 transcripts in the filtered tSnRL1α1 response were assigned to $G_1$ in the 4-h and 6-h iTPS datasets, respectively. Regression plots against CRF are shown in the right hand panels..

(D) Regression plots of CRF against the tSnRK1a1 responses for genes assigned (following the procedure in panel C) to CRF groups $G_2$ and $G_0$. More information about these analyses and correlations between the tSnRL1α1 response and the iTPS response of transcripts assigned to CRF groups $G_2$ and $G_O$ is provided in Supplemental Table S6.

C

4-h iTPS

6-h iTPS

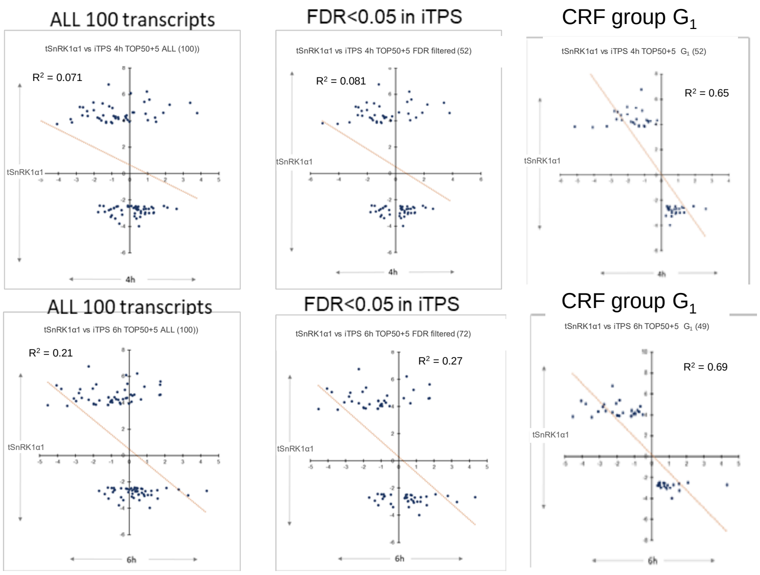

D

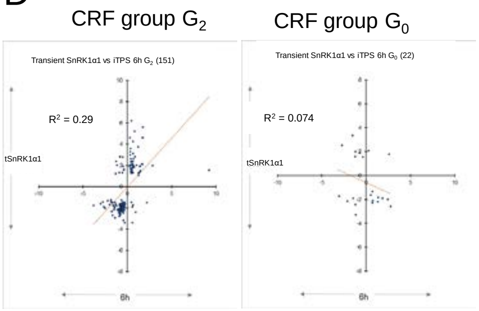

(E) The 1001 genes that were listed by Baena-Gonzalez et al. (2007) as downstream target of SnRK1 were divided into two sets to separate genes that were induced and repressed in the tSnRL1α1 response. In each set, the qualitative response in iTPS was determined, after filtering the iTPS response with an FDR<0.05 and FC<2 filter to allow comparison of two comparably filtered data sets. The table lists, for genes that were induced and repressed in the tSnRL1α1 response, the numbers of transcripts that increased and decreased in the iTPS response. The analysis was done separately for the 4-h and 6-h iTPS responses. The display also shows (in red font) for tSnRL1α1-induced and -repressed genes, what percentage showed a qualitatively opposite response in the tSnRL1α1 response and the iTPS response. A qualitatively opposite response is consistent with a mechanism in which SnRK1 signaling is inhibited by Tre6P. A qualitatively similar response is inconsistent with this mechanism, and consistent with a mechanism in which sugars or other factors regulate SnRK1 activity.

E

| | | **4-h iTPS response** | | **6-h iTPS response** | |
|---|---|---|---|---|---|
| Gene set (tSnRK1α1, iTPS filter) | | in tSnRK1a1 UP gene set | In tSnRK1α1 DOWN set | In tSnRK1α1 UP gene set | In tSnRK1α1 DOWN set |
| ALL FDR < 0.05 + FC2 | UP in iTPS | **32** | 45 | **19** | 65 |
| | DOWN in iTPS | 193 | **58** | 197 | **35** |
| | **Qualitatively opposite response** | **85.8%** | **43.7%** | **83.6%** | **65%** |

**Supplemental Figure S14. Responses of transcripts encoding SnRK1 subunits, or implicated as downstream transcriptional targets of SnRK1 signaling (continued)**

(F-H) Comparison of the iTPS response and the tSnRL1α1with the for three sets of genes related to metabolism, growth and signaling. These gene sets showed a iTPS response with a large CRF $G_1$ component indicating that many of them they are responses to elevated Tre6P. The analysis asks if the observed response in iTPS may be due to inhibition of SnRK1 by Tre6P. (G) Photosynthesis, (H) ribosomal proteins and ribosome assembly,(J) light signaling. In the following three panels, the left-hand display shows the response for all transcript assigned to the category, and the right-hand panel shows the response of those assigned to CRF group $G_1$.

F Comparison of the response of genes involved in **photosynthesis** to iTPS and to tSnRK1α1.

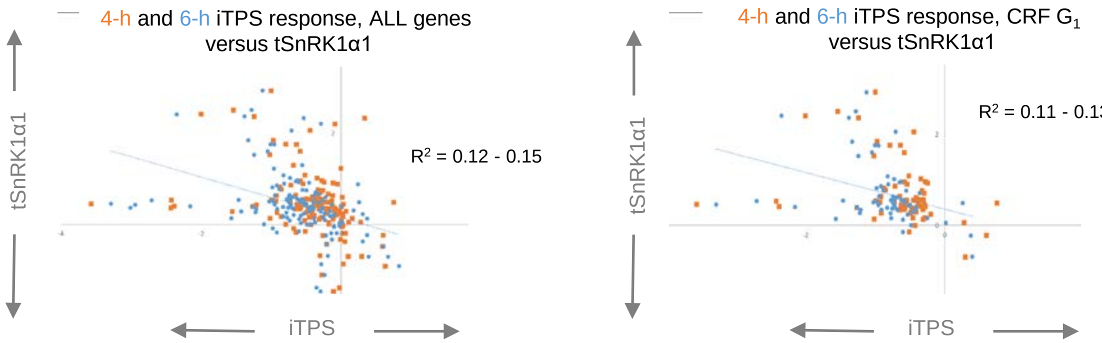

G Comparison of the response of genes involved in **light signaling** to iTPS and to tSnRK1α1. Genes are taken from Supplemental Figure S12D..

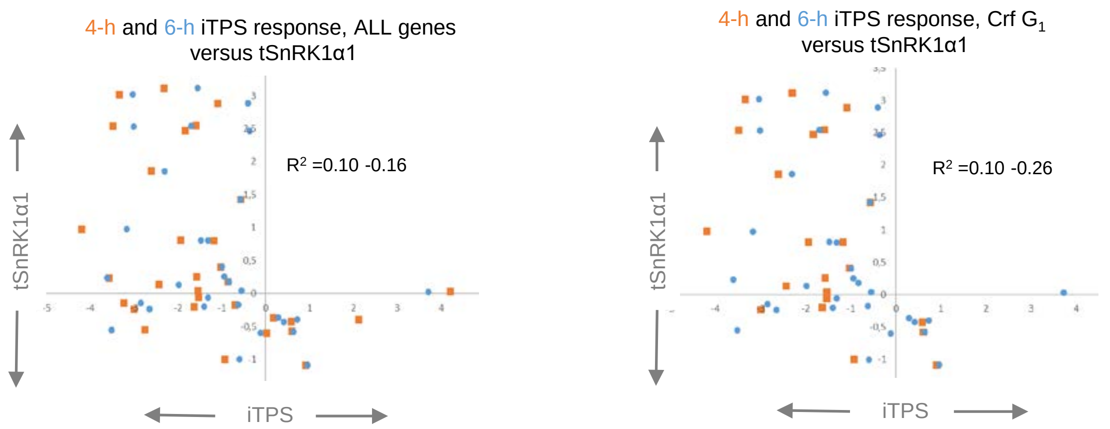

H Comparison of the response of genes encoding **cytosolic ribosomal proteins** to iTPS and to tSnRK1α1. For the responses of genes involved in ribosome assembly see Figure 8C.

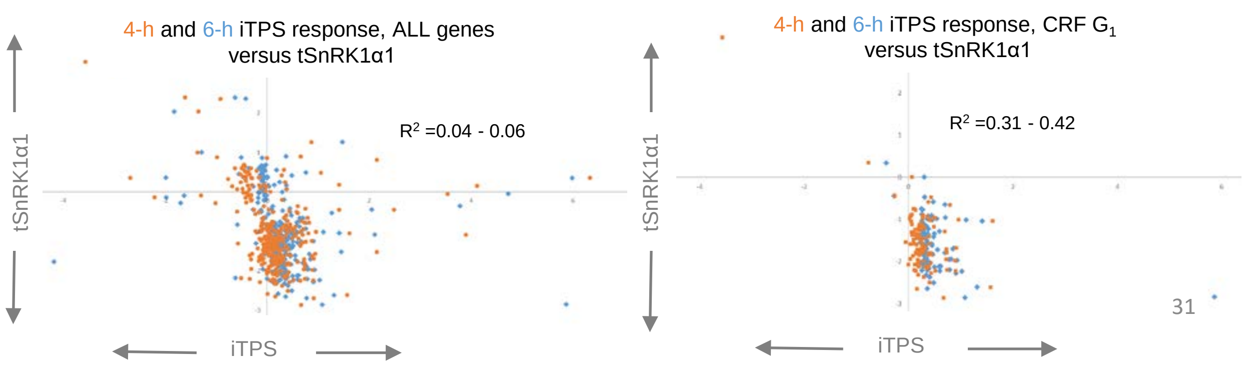

**Supplemental Figure S14. Responses of transcripts encoding SnRK1 subunits, or implicated as downstream transcriptional targets of SnRK1 signaling (continued)**

I Comparison of the tSnRK1α1 response and the CRF of genes annotated as ribosome assembly factors. Most of these genes are induced by sugar and, as reported for a subset by Baena-González et al. (2007), are repressed by tSnRK1a1. The responses of these genes to iTPS and oeTPS are shown in Figure 8C.

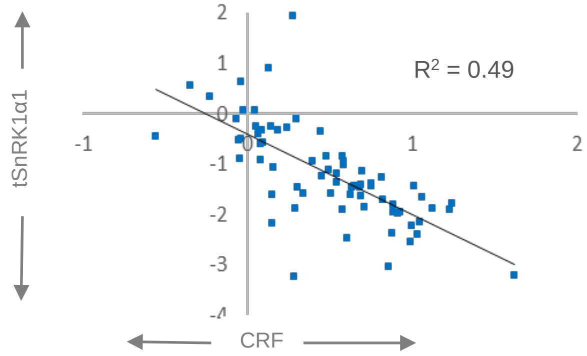

J

| Gene | | iTPS response | | | | | | CRF | Other responses | | | | Direction |
|---|---|---|---|---|---|---|---|---|---|---|---|---|---|
| Name | Code | iTPS 4-h adj.p | iTPS 6-h adj.p | 29.2 4-h log2 | 29.2 6-h log2 | iTPS 4-h CRF score | iTPS 6-h CRF score | | oeTPS log2 | SnRK1 TARGET GENES Log2 | tSnRK1a1 log 2 | oe bZIP11 log2 | iTPS versus tSnRK1α1 |
| *TPS1* | At1g78580 | **0,00** | **0,00** | **-1,07** | **-1,19** | **2** | **2** | 0,48 | 0,00 | | -0,70 | | |
| *TPS2* | At1g16980 | 0,36 | 0,55 | -2,45 | -1,76 | | | 0,01 | 0,00 | | 0,36 | | |
| *TPS3* | At1g17000 | | | | | | | 0,04 | 0,00 | | 0,65 | | |
| *TPS4* | At4g27550 | 0,37 | 0,15 | -1,25 | -3,62 | | | 0,07 | 0,00 | | 0,64 | | |
| *TPS5* | At4g17770 | 0,00 | 0,00 | -0,92 | -0,57 | 2 | 2 | 1,24 | 3,00 | | -1,79 | | Same |
| *TPS6* | At1g68020 | **0,00** | **0,00** | **-0,37** | **-0,39** | **1** | **1** | -0,62 | -1,70 | | 0,60 | | OPP |
| *TPS7* | At1g06410 | 0,59 | 0,83 | -0,08 | -0,04 | | | -0,10 | 0,00 | | 0,31 | | |
| *TPS8* | At1g70290 | **0,00** | **0,00** | **-1,54** | **-1,08** | **1** | **1** | -2,30 | -2,33 | 2,94 | 2,94 | | OPP |
| *TPS9* | At1g23870 | **0,00** | **0,00** | **-1,30** | **-1,35** | **1** | **1** | -1,77 | -2,49 | 3,32 | 3,32 | | OPP |
| *TPS10* | At1g60140 | **0,00** | **0,00** | **-1,17** | **-1,03** | **1** | **1** | -1,63 | -2,27 | 1,79 | 1,79 | | OPP |
| *TPS11* | At2g18700 | **0,00** | **0,00** | **-1,88** | **-1,76** | **1** | **1** | -2,46 | -3,44 | 2,21 | 2,21 | | OPP |
| *TPPA* | At5g51460 | **0,00** | **0,00** | **-1,98** | **-1,56** | **2** | **2** | 0,51 | 2,87 | | -0,22 | | |
| *TPPB* | At1g78090 | **0,00** | **0,00** | **-2,99** | **-2,43** | **2** | **2** | 0,37 | 0,00 | | -0,01 | | |
| *TPPC* | At1g22210 | 0,56 | 0,63 | -1,72 | -0,87 | | | 0,06 | 0,00 | | -0,49 | | |
| *TPPD* | At1g35910 | **0,00** | **0,00** | **-1,41** | **-1,16** | **0** | **0** | 0,06 | 0,00 | | 0,07 | | |
| *TPPE* | At2g22190 | **0,00** | **0,00** | **-3,04** | **-2,27** | **2** | **2** | 0,67 | 0,00 | | -0,27 | | |
| *TPPF* | At4g12430 | 0,38 | 0,54 | 0,16 | 0,12 | | | -0,09 | -4,03 | | -0,17 | 1,26 | |
| *TPPG* | At4g22590 | **0,00** | **0,00** | **0,83** | **0,94** | **0** | **0** | 0,09 | -1,21 | | -1,21 | 1,77 | |
| *TPPH* | At4g39770 | **0,00** | **0,00** | **-3,93** | **-2,48** | **0** | **0** | 0,02 | -2,91 | | -0,29 | | |
| *TPPJ* | At5g65140 | **0,00** | **0,00** | **0,74** | **0,52** | **0** | **0** | 0,07 | 0,00 | | 0,26 | | |
| *TPPI* | At5g10100 | 0,56 | 0,87 | -0,39 | 0,11 | | | 0,06 | 0,00 | | -0,05 | | |
| *TRE* | At4g24040 | **0,00** | **0,00** | **-2,14** | **-2,30** | **1** | **1** | -1,04 | -2,44 | | 1,31 | 3,43 | OPP |
| *AKIN10* | At3g01090 | **0,02** | **0,04** | **0,24** | **0,29** | **2** | **2** | -0,16 | 0,00 | 4,63 | 4,63 | | |
| *AKIN11* | At3g29160 | **0,01** | **0,00** | **0,25** | **0,48** | **2** | **2** | -0,35 | 0,00 | | 0,39 | 1,79 | |
| *SNF4/βγ* | At1g09020 | **0,00** | **0,00** | **0,48** | **0,58** | **1** | **1** | 0,11 | 0,00 | | 0,12 | | |
| *SnRK1β1* | At5g21170 | **0,00** | **0,00** | **-0,92** | **-1,07** | **1** | **1** | -2,19 | -2,25 | 3,23 | 3,23 | | OPP |
| *SnRK1β2* | At4g16360 | **0,01** | **0,01** | **0,28** | **0,34** | **1** | **1** | 0,18 | 1,10 | | -0,52 | | OPP |

Comparison of the iTPS response with the tSnRL1α1 response for TPSs, Class II TPSs, TPPs and TRE (upper part of panel) and for SnRK1 subunits (lower part of panel). The display shows, from left to right,

- the gene
- the iTPS response: p-value, $log_2$ fold change and CRF assignment (genes assigned to CRF group $G_1$ are shaded in light orange). Significant changes in the iTPS response are highlighted in bold font.
- the CRF value (green shading, see Supplemental Figure S2).
- the response to constitutive overexpression of bacterial TPS (oeTPS, from Zhang et al., 2009)
- the response to transient overexpression of SnRK1α1 (t SnRK1α1, two columns, the first showing changes called significant and the second then $log_2$ FC for all, from Baena-Gonzalez et al., 2007).
- the response to constitutive overexpression of bZIP11 (oebZIP11, from Ma et al., 2011).

The right-hand column compares, for genes assigned to CRF group $G_1$ in the iTPS response, whether the direction of change in the iTPS response an the tSnRK1α1 response was in the same direction ('same') or in opposite directions ('OPP' , shaded orange). An opposite response is consistent with it being due to Tre6P-inhibition of SnRK1.

**Supplemental Figure S15. TOR subunits, higher levels regulators of ribosome biogenesis and other TOR targets including ABA receptors**.

This Supplemental Figure investigates the possible interaction of Tre6P in regulation of TOR expression and TOR signaling. See Supplemental Figure S14K for a description of the column headings.

(A) TOR subunits.

(B-E) Proteins identified as targets of TOR phosphorylation (Scarpin et al., 2020, Meng et al., 2022) divided into (B) protein kinases that are direct targets of TOR, (C) further genes implicated in ribosome biogenesis, (D) the family of ABA receptors that were highlighted as TOR phosphorylation targets in the analysis of Meng et al. (2022) and (E) further proteins with diverse functions, the upper block is from Meng et al. (2022), the middle block from Scarpin et al. (2020) and the lower block from Liao et al. (2022). The displays show the response in iTPS at 4h and 6h, assignment of the iTPS response to CRF groups, the CRF value (defined as in Supplemental Figure S1) and the response to constitutive overexpression of TPS (oeTPS, Zhang et al., 2009) and transient overexpression of SnRL1α1 in protoplasts (tSnRL1α1, Baena-Gonzalez et al., 2007).

Significant changes in the iTPS response are shown with bold font. Assignment to CRF groups is performed only for significant responses and only CRF1 assignments are shaded. No entry for oeTPS or tSnRL1α1 indicates the response did not pass the filter in the publication..

A

| **Gene** | | **iTPS response** | | | | | | **CRF** | **Other responses** | |
|---|---|---|---|---|---|---|---|---|---|---|
| Name | Code | 4-h adj. p | 6-h adj. p | **4-h FC $log_2$** | **6-h FC $log_2$** | 4-h CRF score | 6-h CRF score | | oeTPS FC $log_2$ | tSnRK1α1 FC $Log_2$ |
| TOR | At1g50030 | 0,75 | 0,55 | -0,06 | 0,15 | | | -0,17 | 0,00 | 0,51 |
| RAPTOR2 | At5g01770 | **0,00** | 0,36 | **-0,45** | 0,19 | 2 | | 0,18 | 0,00 | -0,24 |
| RAPTOR1 | At3g08850 | **0,01** | **0,02** | **0,37** | **0,45** | 2 | 2 | -0,14 | 0,00 | 0,44 |
| LST1 / 2 | At3g18140/ At | na | na | na | na | | | -0,16 | na | na |

B

| **Gene** | | **iTPS response** | | | | | | **CRF** | **Other responses** | |
|---|---|---|---|---|---|---|---|---|---|---|
| Name | Code | 4-h adj. p | 6-h adj. p | **4-h FC $log_2$** | **6-h FC $log_2$** | 4-h CRF score | 6-h CRF score | | oeTPS FC $log_2$ | tSnRK1α1 FC $Log_2$ |
| S6K1 | At3g08720 | 0,11 | **0,06** | 0,19 | **0,33** | | 2 | -0,34 | 1,16 | 0,56 |
| S6K2 | At3g08730 | 0,24 | 0,31 | -0,16 | -0,15 | | | -0,04 | | -0,51 |
| LARP1a | At5g21160 | **0,00** | **0,00** | **0,43** | **0,48** | 2 | 2 | -0,44 | | 1,08 |
| LARP1b | At5g66100 | 0,36 | **0,00** | 0,10 | **0,36** | | 1 | 0,20 | | |
| LARP1c | At4g35890 | **0,02** | **0,00** | **0,23** | **0,60** | 1 | 1 | 0,33 | | |
| YAK1 | At5g35980 | 0,11 | 0,01 | 0,21 | **0,44** | | 0 | -0,05 | | |

C

| **Gene** | | **iTPS response** | | | | | | **CRF** | **Other responses** | |
|---|---|---|---|---|---|---|---|---|---|---|
| Name | Code | 4-h adj. p | 6-h adj. p | **4-h FC $log_2$** | **6-h FC $log_2$** | 4-h CRFs core | 6-h CRF score | | oeTPS FC $log_2$ | tSnRK1α1 FC $Log_2$ |
| NAP1;1 | At4g26110 | **0,00** | **0,06** | **0,34** | 0,26 | 1 | | 0,46 | | -1,37 |
| RPS6 | At4g31700 | **0,01** | 0,36 | **0,26** | 0,15 | 1 | | 0,62 | 1,52 | -2,23 |

**Supplemental Figure S15. TOR subunits, higher levels regulators of ribosome biogenesis and other TOR targets including ABA receptors**. Continued .

## D

The family of ABA receptors that were highlighted as TOR phosphorylation targets in the data set of Meng *et al*. (2022).

| Gene | | iTPS response | | | | | | CRF | Other responses | |
|---|---|---|---|---|---|---|---|---|---|---|
| Name | Code | 4-h adj.p | 6-h adj.p | 4-h FC $log_2$ | 6-h FC $log_2$ | 4-h Crf score | 6-h Crf score | | oeOTSA FC $log_2$ | tSnRK1α1 FC $Log_2$ |
| **PYL1** | At5g46790 | **0.00** | **0.00** | **-2.26** | **-1.67** | **2** | **2** | 0.30 | 0.00 | 0.33 |
| **PYL2** | At2g26040 | **0.00** | **0.00** | **-0.89** | **-1.45** | **0** | **0** | 0.08 | 0.00 | **1.03** |
| **PYL3** | At1g73000 | 0.20 | 0.62 | -1.99 | 1.41 | | | -0.03 | 0.00 | **0.97** |
| **PYL4** | At2g38310 | **0.00** | **0.00** | **-3.39** | **-2.79** | **2** | **2** | 0.11 | 0.00 | -0.32 |
| **PYL5** | At5g05440 | **0.00** | **0.00** | **-4.11** | **-3.56** | **1** | **1** | -0.53 | 0.00 | **1.92** |
| **PYL6** | At2g40330 | **0.00** | **0.00** | **-3.94** | **-3.84** | **0** | **0** | 0.04 | 0.00 | 0.51 |
| **PYL7** | At4g01026 | **0.00** | **0.00** | **-1.49** | **-1.26** | **1** | **1** | -0.95 | **-3.24** | **2.57** |
| **PYL8** | At5g53160 | **0.00** | **0.00** | **-1.61** | **-1.39** | **1** | **1** | -0.73 | **-2.55** | **0.58** |
| **PYR9** | At1g01360 | **0.01** | **0.01** | **-0.38** | **-0.49** | **1** | **1** | -0.60 | **-1.35** | **0.93** |
| **PYR1** | At4g17870 | **0.00** | **0.00** | **-1.36** | **-1.96** | **0** | **0** | -0.05 | 0.00 | 0.44 |

## E

Further proteins that were highlighted as downstream TOR phosphorylation targets in the data sets of Meng *et al.* (2022) and Brunkold *et al*. (2020), as well as the autophagysome assembly factor ATG13a that is considered a target of TOR (Liao *et al*., 2022)

| Gene | | iTPS response | | | | | | CRF | Other responses | | Annotation |
|---|---|---|---|---|---|---|---|---|---|---|---|
| Name | Code | 4-h adj.p | 6-h adj.p | 4-h FC $log_2$ | 6-h FC $log_2$ | 4-h CRF score | 6-h CRFscore | | oeTPS FC $log_2$ | tSnRK1α1 FC $Log_2$ | |
| E2Fa | At2g36010 | **0,03** | 0,38 | **0,32** | 0,15 | 1 | | 0,41 | 0,00 | 0,32 | E2F TRANSCRIPTION FACTOR-3; cell cycle, components of cyclin D/retinoblastoma/E2F pathway |
| E2Fb | At5g22220 | 0,31 | 0,31 | 0,17 | 0,20 | | | -0,24 | 0,00 | 1,36 | E2F1; transcription factor, component of cyclin D/retinoblastoma/E2F pathway. RBR1 proteins |
| ATG1B | At3g53930 | 0,66 | 0,34 | -0,07 | -0,16 | | | -0,24 | 0,00 | 0,73 | Protein kinase superfamily protein |
| ATG13 | At3g49590 | **0,01** | **0,00** | **0,26** | **0,48** | 2 | 2 | -0,59 | 0,00 | 2,40 | Autophagy protein |
| EIN2 | At5g03280 | 0,08 | **0,00** | 0,26 | **0,66** | | 0 | 0,00 | 0,00 | -0,08 | ETHYLENE INSENSITIVE 2; transporter, ethylene signal transduction acting downstream of CTR1 |
| PIN2 | At5g57090 | **0,00** | **0,03** | **1,89** | **0,98** | 1 | 1 | 0,10 | 1,61 | 0,74 | EIR1 (ETHYLENE INSENSITIVE ROOT 1); auxin:hydrogen symporter/ transporter |
| S40-7 | At3g15040 | 0,32 | 0,11 | -0,13 | -0,24 | | | -0,21 | 0,00 | 0,47 | Senescence regulator |
| hyp. prot. | At3g50370 | 0,24 | 0,09 | 0,19 | 0,36 | | | 0,02 | 0,00 | -0,22 | similar to cupin family protein [Arabidopsis thaliana] (TAIR:AT2G18540.1) |
| eIF2B-δ1 | At2g44070 | 0,10 | **0,02** | -2,43 | **3,24** | | 1 | 0,26 | na | na | eukaryotic translation initiation factor 2B family protein / eIF-2B family protein |
| TAP46 | At5g53000 | **0,00** | **0,02** | **0,37** | **0,30** | 1 | 1 | 0,18 | 0,00 | 0,34 | PP2A-associated protein with a possible function in the chilling response |
| AML5 | At1g29400 | 0,20 | **0,00** | -0,13 | **0,51** | | 2 | -0,50 | 0,00 | 0,26 | member of the *mei2-like* gene family, encoding RNA -binding proteins |
| eS6b | At5g10360 | **0,00** | 0,10 | **0,33** | 0,25 | 1 | | 0,19 | #N/A | -1,72 | EMB3010 \| EMB3010 (embryo defective 3010); structural constituent of ribosome \| chr5 |
| eIF4B1 | At3g26400 | **0,02** | 0,17 | **0,25** | 0,16 | 1 | | 0,17 | #N/A | #N/A | EIF4B1 \| EIF4B1; translation initiation factor \| chr3 |
| eIF4B2 | At1g13020 | **0,05** | **0,01** | **0,20** | **0,36** | 1 | 1 | 0,31 | #N/A | #N/A | eIF4B, mRNA unwinding factor |
| eIF3m | At3g0220( | **0,00** | **0,00** | **0,58** | **0,488** | 2 | 2 | -0,13 | #N/A | #N/A | eIF3 mRNA-to-PIC binding complex.eIF3m component |
| VLN2 | At2g41740 | **0,00** | 0,24 | **-0,34** | -0,19 | 2 | | 0,12 | #N/A | #N/A | VILLIN 2; actin binding \| chr2 |
| VLN3 | At3g57410 | **0,00** | **0,00** | **-0,75** | **-0,78** | 1 | 1 | -0,25 | #N/A | #N/A | VILLIN 3; actin binding \| chr3 |
| TOPLESS | At1g15750 | **0,00** | **0,00** | **-1,14** | **-0,90** | 1 | 1 | -0,25 | #N/A | #N/A | TOPLESS, transcription repressor \| chr1 |
| CBE1 | At4g01290 | 0,25 | **0,04** | 0,16 | **0,37** | | 9 | 0,05 | #N/A | #N/A | CONSERVED BINDING OF EIF4E |
| PUX5 | At4g15410 | 0,06 | 0,14 | 0,24 | 0,21 | | | 0,36 | #N/A | #N/A | plant ubiquitin regulatory X domain-containing protein |
| ATG13a | At3g49590 | **0,01** | **0,00** | **0,26** | **0,48** | 2 | 2 | -0,59 | #N/A | 2,4 | Vesicle trafficking.autophagosome formation, ATG13 accessory component |

**Supplemental Figure S16. FCS-Like Zinc finger (FLZ) family proteins.**

The annotation is taken from Jamsheer et al. (2015). All of these proteins are annotated in TAIR11 as 'SnRK1 metabolic regulator system. FLZ SnRK1-interacting factors'.

They have been reported to respond differently to sugar addition and other treatments like hormone addition and environmental stress (Nietzsche et al., 2014; Jamsheer et al., 2015). The responsiveness to sugar as reported by Jamsheer and Laxmi (2015) is given in the left-hand column, with genes allocated to three sets that that were defined in this publication: set 1 (repressed by >10fold), set 2 (repressed by 2-10-fold) or set 3 (inconsistent responses or are induced). FLZ6 (like FLZ10) was subsequently found to be starvation-induced (Jamsheer *et a*l., 2018).

The display shows the 4-h and 6-h iTPS response, the assignment of the iTPS response to CRF groups, the CRF value (defined as in Supplemental Figure 1 and the response to constitutive overexpression of TPS (oeTPS, from Zhang et al., 2009) ) and transient overexpression of SnRK1α1 in protoplasts (tSnRL1α1, from Baena-Gonzalez et al., 2007). Significant changes in the iTPS response are shown with bold font. Assignment to CRF groups is performed only for significant responses and only CRF $G_1$ assignments are shaded.
No entry for oeTPS or tSnRL1α1 indicates that the response did not pass the filter in the original publication.

Significant changes in the iTPS response are shown with bold font. Assignment to Crf groups is performed only for significant responses, and only CRF $G_1$ assignments are shaded

Many FLZ family members are significantly repressed in the iTPS response including four of five in set 1 (FLZ1, FLZ5, FLZ8, FLZ14), two of four in set 2 (FLZ3, FLZ15), and four of five in set 3 (FLZ6, FLZ9, FLZ13, FLZ17) of Jamsheer and Laxmi (2015). One was induced in set 2 (FLZ10) of Jamsheer and Laxmi (2015) However, internal analysis in the iTPS data set and especially comparison of the iTPS response with the Crf indicated that most of the responses were indirect (Crf groups 2 or 0) and therefore unlikely to be due to Tre6P signalling. Those assigned to a possible Tre6P-response ((CRF group G1) were FLZ9, FLZ13 and FLZ19, all of which had been previously classed as non-sugar responsive by Jamsheer and Laxmi (2015).

Of the sugar-responsive FLZs, experimental evidence has been presented for an inhibitory effect on SnRK1 for FLZ3 (Bortlik et al., 2022), FLZ8 (Jamsheer et al., 2022) and FLZ6 and FLZ10 (Jamsheer et al., 2018). All of these are inhibited in the iTPS response but all are assigned in CRF groups $G_2$, indicating that the response is indirect, probably due to the decrease in sugars.

FLZ4 (At4g65040), FLZ7 (At4g39795) and FLZ18 (At1g53903) were not present in the filtered iTPS response dataset, and FLZ16 was not scored in the iTPS response.

| Grouping in Jamsheer and Laxmi (2015) | Gene | | iTPS response | | | | | | CRF | Other responses | | Ontology and annnotation |
|---|---|---|---|---|---|---|---|---|---|---|---|---|
| | Name | Code | 4-h adj.p | 6-h adj.p | **4-h FC $log_2$** | **6-h FC $log_2$** | 4-h CRF score | 6-h CRF score | | oeTPS FC $log_2$ | tSnRK1α1 FC $Log_2$ | |
| 1 | FLZ1 | At5g47060 | **0,00** | 0,91 | **-0,55** | 0,02 | **2** | | 1,23 | **3,10** | #N/A | Multi-process regulation.SnRK1 metabolic regulator system.FLZ SnRK1-interacting factor |
| 1 | FLZ2 | At4g17670 | 0,14 | 0,81 | -0,27 | -0,04 | | | 1,19 | #N/A | #N/A | Multi-process regulation.SnRK1 metabolic regulator system.FLZ SnRK1-interacting factor |
| 1 | FLZ5 | At1g22160 | **0,00** | **0,00** | **-1,16** | **-1,20** | **2** | **2** | 0,21 | #N/A | #N/A | Multi-process regulation.SnRK1 metabolic regulator system.FLZ SnRK1-interacting factor |
| 1 | FLZ8 | At3g22550 | **0,00** | **0,00** | **-1,50** | **-0,75** | **2** | **2** | 0,71 | #N/A | #N/A | Multi-process regulation.SnRK1 metabolic regulator system.FLZ SnRK1-interacting factor |
| 1 | FLZ14 | At5g20700 | **0,00** | **0,00** | **-1,16** | **-0,90** | **2** | **2** | 0,84 | #N/A | -2,78 | Multi-process regulation.SnRK1 metabolic regulator system.FLZ SnRK1-interacting factor |
| 2 | FLZ3 | At2g44670 | **0,00** | **0,00** | **-1,30** | **-0,64** | **2** | **2** | 0,29 | #N/A | #N/A | Multi-process regulation.SnRK1 metabolic regulator system.FLZ SnRK1-interacting factor |
| 2 | FLZ10 | At5g11460 | **0,00** | **0,00** | **0,47** | **0,96** | **0** | **0** | 0,09 | #N/A | #N/A | Multi-process regulation.SnRK1 metabolic regulator system.FLZ SnRK1-interacting factor |
| 2 | FLZ11 | At2g25690 | 0,19 | 0,17 | 0,23 | 0,26 | | | 0,23 | #N/A | #N/A | Multi-process regulation.SnRK1 metabolic regulator system.FLZ SnRK1-interacting factor |
| 2 | FLZ15 | At5g49120 | **0,00** | **0,00** | **-2,01** | **-1,29** | **0** | **0** | 0,03 | #N/A | #N/A | Multi-process regulation.SnRK1 metabolic regulator system.FLZ SnRK1-interacting factor |
| 2 | FLZ16 | At3g63230 | #N/A | #N/A | #N/A | #N/A | | | 0,01 | #N/A | #N/A | Multi-process regulation.SnRK1 metabolic regulator system.FLZ SnRK1-interacting factor |
| 3 | FLZ6 | At1g78020 | **0,00** | **0,00** | **-1,53** | **-0,72** | **2** | **2** | 0,28 | #N/A | #N/A | Multi-process regulation.SnRK1 metabolic regulator system.FLZ SnRK1-interacting factor |
| 3 | FLZ9 | At3g63210 | **0,00** | **0,00** | **-3,52** | **-3,00** | 1 | 1 | -0,53 | #N/A | #N/A | Multi-process regulation.SnRK1 metabolic regulator system.FLZ SnRK1-interacting factor |
| 3 | FLZ12 | At1g19200 | 0,94 | 0,18 | -0,03 | -0,35 | | | -0,01 | #N/A | #N/A | Multi-process regulation.SnRK1 metabolic regulator system.FLZ SnRK1-interacting factor |
| 3 | FLZ13 | At1g74940 | 0,59 | **0,00** | -0,09 | **-0,40** | | **1** | -0,17 | #N/A | #N/A | Multi-process regulation.SnRK1 metabolic regulator system.FLZ SnRK1-interacting factor |
| 3 | FLZ17 | At1g53885 | **0,01** | **0,00** | **-0,93** | **-1,39** | **1** | **1** | -1,49 | #N/A | #N/A | Multi-process regulation.SnRK1 metabolic regulator system.FLZ SnRK1-interacting factor |

**Supplemental Figure S17. Brassinosteroid signalling**.

(A) Genes involved in brassinosteroid synthesis and signaling, based on the MapMan ontology.

(B) Set of brassinosteroid-regulated genes, taken from Wang *et al*. (2016)

The displays show the 4-h and 6-h iTPS response, assignment of the iTPS response to CRF groups, the CRF value (defined as in Supplemental Figure S2) and the response to constitutive overexpression of TPS (oeTPS, from Zhang *et al*., 2009) and transient overexpression of SnRL1α1 in protoplasts (tSnRL1α1, from Baena-Gonzalez *et al*., 2007). Significant changes in the iTPS response are shown with bold font. Assignment to CRF groups is performed only for significant responses and only CRF $G_1$ assignments are shaded. No entry for oeTPS or tSnRL1α1 indicates that the response did not pass the filter in the original publication.

A

| Gene | | iTPS response | | | | | | CRF | Other responses | | Annotation |
|---|---|---|---|---|---|---|---|---|---|---|---|
| Name | Code | 4-h adj.p | 6-h adj.p | **4-h FC log$_2$** | **6-h FC log$_2$** | 4-h CrRFscore | 6-h CRF score | | oeTPS FC log$_2$ | tSnRK1α1 FC Log$_2$ | |
| BR6OX1 | At5g38970 | **0,12** | **0,02** | **-3,83** | **-5,58** | | 9 | 0,04 | | -0,10 | BRASSINOSTEROID-6-OXIDASE; oxygen binding |
| | At3g50750 | **0,00** | **0,00** | **-3,27** | **-2,37** | 1 | 1 | -0,33 | -1,56 | 0,92 | brassinosteroid signalling positive regulator-related |
| BEE2 | At4g36540 | **0,00** | **0,00** | **-3,05** | **-2,66** | 1 | 1 | -0,82 | -2,14 | 0,94 | BR ENHANCED EXPRESSION 2; DNA binding / transcription factor |
| ROT1 | At4g36380 | **0,00** | **0,00** | **-2,88** | **-1,63** | 2 | 2 | 0,10 | | 0,47 | ROTUNDIFOLIA 3; oxygen binding / steroid hydroxylase |
| BEE3 | At1g73830 | **0,00** | **0,00** | **-2,44** | **-1,76** | 1 | 1 | -0,43 | | -0,51 | BR ENHANCED EXPRESSION 3; DNA binding / transcription factor |
| CYP90D1 | At3g13730 | **0,00** | **0,00** | **-2,39** | **-2,51** | 1 | 1 | -0,15 | | -0,38 | CYTOCHROME P450, FAMILY 90, SUBFAMILY D, POLYPEPTIDE 1; oxidoreductase, |
| DWF4 | At3g50660 | **0,00** | **0,00** | **-2,38** | **-1,77** | 2 | 2 | 0,38 | 1,06 | -2,32 | DWARF 4 |
| BR6OX2 | At3g30180 | **0,00** | **0,00** | **-2,32** | **-1,14** | 2 | 2 | 0,39 | | -0,98 | BRASSINOSTEROID-6-OXIDASE 2; monooxygenase/ oxygen binding |
| CYP72C1 | At1g17060 | **0,00** | **0,00** | **-2,28** | **-1,96** | 1 | 1 | -0,14 | 0,00 | -1,68 | cytochrome P450, family 72, subfamily C, polypeptide 1; oxygen binding |
| BRH1 | At3g61460 | **0,00** | **0,00** | **-2,25** | **-1,71** | 0 | 0 | -0,05 | 0,00 | -0,42 | BRASSINOSTEROID-RESPONSIVE RING-H2; protein binding / zinc ion binding |
| CPD | At5g05690 | **0,00** | **0,00** | **-2,19** | **-1,48** | 1 | 1 | -0,87 | -1,71 | -0,07 | CABBAGE 3; oxygen binding |
| SMT3 | At1g76090 | **0,00** | **0,00** | **-2,00** | **-2,31** | 1 | 1 | -0,20 | | 0,34 | S-adenosyl-methionine-sterol-C-methyltransferase 3) |
| BRS1 | At4g30610 | **0,00** | **0,00** | **-1,93** | **-2,05** | 2 | 2 | 0,26 | 0,00 | -0,21 | BRI1 SUPPRESSOR 1 |
| SQP1,2 | At5g24160 | **0,00** | **0,00** | **-1,72** | **-2,04** | 1 | 1 | -1,07 | -1,16 | 1,43 | squalene monooxygenase 1,2 / squalene epoxidase 1,2 (SQP1,2) |
| | At5g08130 | **0,00** | **0,00** | **-1,58** | **-1,07** | 2 | 2 | 0,29 | | -1,12 | bHLH family protein, BRZ signaling, interacts with BES1 to bind E box sequences (CANNTG) |
| SMT2 | At1g20330 | **0,00** | **0,00** | **-1,44** | **-1,18** | 9 | 9 | 0,01 | | -1,35 | STEROL METHYLTRANSFERASE 2 |
| BRL2 | At2g01950 | **0,00** | **0,00** | **-1,33** | **-1,24** | 1 | 1 | -0,22 | | 0,07 | BRI1-LIKE 2; ATP binding / protein serine/threonine kinase |
| BEE1 | At1g18400 | **0,00** | **0,00** | **-1,29** | **-1,69** | 2 | 2 | 0,16 | | -0,18 | BR ENHANCED EXPRESSION 1; transcription factor |
| SMT1 | At5g13710 | **0,00** | **0,00** | **-1,06** | **-1,87** | 1 | 1 | -0,34 | | -0,60 | STEROL METHYLTRANSFERASE 1 |
| BIN2 | At4g18710 | **0,00** | **0,00** | **-1,02** | **-0,69** | 1 | 1 | -0,30 | | 0,35 | BRASSINOSTEROID-INSENSITIVE 2; kinase |
| | At4g18890 | **0,00** | **0,00** | **-0,70** | **-0,59** | 2 | 2 | 0,30 | | -0,82 | brassinosteroid signalling positive regulator-related |
| DWF1 | At3g19820 | **0,00** | **0,00** | **-0,55** | **-0,87** | 0 | 0 | 0,03 | | -1,08 | DIMINUTO 1; catalytic |
| BRI1 | At4g39400 | **0,00** | **0,21** | **-0,52** | **-0,24** | 2 | | 0,22 | | -0,28 | BRASSINOSTEROID INSENSITIVE 1; kinase |
| | At2g22830 | **0,00** | **0,00** | **-0,49** | **-0,45** | 2 | 2 | 0,14 | | 0,55 | qualene monooxygenase, putative |
| BRZ1 | At1g75080 | **0,00** | 0,10 | **-0,49** | -0,24 | 2 | | 0,10 | | -0,17 | BRASSINAZOLE-RESISTANT; transcription regulatora, (BRI1-EMS-SUPPRESSOR 1) |
| DET2 | At2g38050 | **0,00** | **0,00** | **-0,45** | **-0,63** | 0 | 0 | -0,09 | | -0,19 | DE-ETIOLATED 2 |
| CAS1 | At2g07050 | **0,03** | **0,07** | **-0,29** | **-0,28** | 0 | | 0,05 | -1,78 | -0,06 | (CYCLOARTENOL SYNTHASE 1 |
| CYP51G1 | At1g11680 | **0,01** | **0,05** | **-0,26** | **-0,29** | 2 | 2 | 0,17 | | -0,49 | CYTOCHROME P450 51; oxygen binding |
| DWF5 | At1g50430 | **0,06** | 0,92 | **-0,22** | -0,02 | | | -0,09 | | 0,10 | DWARF 5; sterol delta7 reductase |
| STE1 | At3g02580 | 0,33 | **0,00** | -0,14 | **0,30** | | 1 | 0,34 | | -0,93 | STEROL 1; C-5 sterol desaturase |
| | At4g36780 | 0,22 | **0,04** | -0,14 | **0,33** | | 2 | -0,37 | | -0,58 | brassinosteroid signalling positive regulator-related |
| HYD1 | At1g20050 | 0,31 | 0,91 | -0,11 | -0,02 | | | 0,20 | | -1,04 | Hydra 1 |
| FK | At3g52940 | 0,48 | 0,94 | -0,09 | 0,01 | | | 0,56 | | -1,07 | FACKEL; delta14-sterol reductase |
| IMK2 | At3g51740 | 0,53 | 0,04 | -0,08 | -0,31 | | 0 | 0,08 | | 0,89 | INFLORESCENCE MERISTEM RECEPTOR-LIKE KINASE 2 |
| | At1g78700 | 0,77 | 0,16 | -0,05 | -0,21 | | | -0,08 | | 0,00 | brassinosteroid signalling positive regulator-related |
| | At2g16530 | 0,89 | **0,00** | -0,03 | **0,78** | | 0 | -0,09 | | 0,11 | 3-oxo-5-alpha-steroid 4-dehydrogenase family protein |
| BAK1 | At4g33430 | 0,86 | **0,00** | 0,02 | **0,47** | | 0 | -0,06 | | -0,05 | BRI1-ASSOCIATED RECEPTOR KINASE; kinase |
| SQP1 | At5g24150 | 0,53 | **0,00** | 0,07 | **0,66** | | 2 | -0,33 | -1,29 | 0,05 | squalene monooxygenase 1 |
| | At4g37760 | 0,40 | **0,00** | 0,09 | **0,42** | | 2 | -0,61 | | 1,30 | squalene monooxygenase, putative / squalene epoxidase, putative |
| | At5g16010 | 0,19 | **0,00** | 0,19 | **0,44** | | 2 | -0,29 | -1,25 | 0,37 | 3-oxo-5-alpha-steroid 4-dehydrogenase family protein |
| BAS1 | At2g26710 | 0,22 | **0,00** | 0,26 | **-0,76** | | 1 | -0,23 | | 0,09 | PHYB ACTIVATION TAGGED SUPPRESSOR 1; oxygen binding |
| CP1 | At5g50375 | **0,00** | **0,00** | **0,35** | **0,55** | 1 | 1 | 0,31 | | 0,40 | CYCLOPROPYL ISOMERASE |
| | At1g74360 | **0,02** | **0,00** | **0,43** | **0,98** | 1 | 1 | 0,13 | 1,39 | -0,34 | leucine-rich repeat transmembrane protein kinase, putative |
| ATTOP6B | At3g20780 | **0,00** | **0,00** | **0,49** | **0,55** | 1 | 1 | 0,28 | | -1,68 | BRASSINOSTEROID INSENSITIVE 3, ROOT HAIRLESS 3 |
| | At3g02590 | 0,73 | 0,94 | 0,62 | -0,14 | | | 0,06 | | -0,18 | delta 7-sterol-C5-desaturase, putative |
| XF1 | At1g58440 | **0,00** | **0,00** | **0,88** | **0,97** | 1 | 1 | 0,28 | | -0,73 | XF1; oxidoreductase |
| SQP2 | At5g24140 | 0,08 | 0,91 | 1,20 | -0,12 | | | -0,34 | | 0,89 | Squalene monooxygenase 2; oxidoreductase |
| ST | At2g03760 | **0,00** | **0,00** | **1,42** | **2,47** | 2 | 2 | -0,92 | 2,21 | 1,14 | steroid sulfotransferase |
| BRL3 | At3g13380 | **0,00** | **0,00** | **1,99** | **2,48** | 1 | 1 | 0,18 | 2,08 | -0,02 | BRI1-LIKE 3; protein binding / protein kinase |

B

| Gene | | iTPS response | | | | | | CRF | Other responses | | Annotation |
|---|---|---|---|---|---|---|---|---|---|---|---|
| Name | Code | 4-h adj.p | 6-h adj.p | **4-h FC log$_2$** | **6-h FC log$_2$** | 4-h CRF score | 6-h CRFf score | | oeTPS FC log$_2$ | tSnRK1α1 FC Log$_2$ | |
| EXPA1 | At1g69530 | **0,00** | **0,00** | **-4,10** | **-3,07** | 0 | 0 | 0,00 | -2,07 | -1,71 | EXPANSIN A1 |
| EXPA3 | At2g37640 | **0,00** | **0,00** | **-1,18** | **-2,07** | 2 | 2 | 0,18 | 0,00 | 2,17 | ARABIDOPSIS THALIANA EXPANSIN A3 |
| EXPA4 | At2g39700 | **0,00** | **0,00** | **-1,98** | **-1,60** | 2 | 2 | 0,92 | 0,00 | -0,14 | EXPANSIN A4 |
| EXPA6 | At2g28950 | **0,00** | **0,00** | **-2,04** | **-2,06** | 2 | 2 | 0,70 | -1,03 | -0,45 | EXPANSIN A6 |
| EXPA8 | At2g40610 | **0,00** | **0,00** | **-5,23** | **-4,14** | 0 | 0 | -0,05 | 0,00 | 0,00 | EXPANSIN A8 |
| EXPA9 | At5g02260 | **0,00** | **0,00** | **-1,17** | **-1,16** | 2 | 2 | 0,31 | 0,00 | 0,37 | (at5g02260):ATEXPA9 (ARABIDOPSIS THALIANA EXPANSIN A9) |
| EXPA11 | At1g20190 | **0,00** | **0,00** | **-3,71** | **-3,67** | 0 | 0 | -0,06 | 0,00 | 3,19 | EXPANSIN A11 |
| EXPA14 | At5g56320 | **0,00** | **0,00** | **-1,94** | **-2,25** | 0 | 0 | 0,05 | 0,00 | -1,34 | EXPANSIN A14 |
| TIP2/TIP2; | At4g17340 | **0,00** | **0,00** | **-2,16** | **-1,84** | 1 | 1 | -0,14 | 1,22 | -0,26 | DELTA-TIP2/TIP2;2 (tonoplast intrinsic protein 2;2); water channel |
| KCS | At1g01120 | **0,00** | **0,00** | **-2,83** | **-2,27** | 2 | 2 | 0,64 | -3,29 | 0,27 | KETOACYL-COA SYNTHASE 1; acyltransferase |
| XTH18 | At4g30280 | **0,93** | **0,00** | 0,05 | **-1,59** | | 0 | -0,02 | 3,03 | 0,45 | XYLOGLUCAN ENDOTRANSGLUCOSYLASE/HYDROLASE 18 |
| XTH19 | At4g30290 | **0,00** | **0,04** | **-2,00** | **-0,95** | 2 | 2 | 0,52 | 2,85 | -0,95 | XYLOGLUCAN ENDOTRANSGLUCOSYLASE/HYDROLASE 19 |

**Supplemental Figure S18. Cell wall modification**

(A) EXPANSINs, (B) XYLOGLUCAN ENDOTRANSGLUCOSYLASEs. The displays show the iTPS response at 4h and 6h, assignment of the iTPS response to CRF groups, the CRF value (defined as in Supplemental Figure S2) and the response to constitutive overexpression of TPS (oeTPS, from Zhang et al., 2009) and transient overexpression of SnRL1α1 in protoplasts (tSnRL1α1 response, from Baena-Gonzalez *et al*., 2007). Significant changes in the iTPS response are shown with bold font. Assignment to CRF groups is performed only for significant responses and only CRF1 assignments are shaded. No entry for oeTPS or tSnRL1α1 indicates that the response did not pass the filter in the original publication.

A

| Gene | iTPS response | | | | | | CRF | Other responses | | Annotation |
|---|---|---|---|---|---|---|---|---|---|---|
| Code | 4-h adj.p | 6-h adj.p | **4-h FC $log_2$** | **6-h FC $log_2$** | 4-h CRF score | 6-h CRF score | | oeTPS FC $log_2$ | tSnRK1α1 FC $Log_2$ | |
| At2g40610 | **0,00** | **0,00** | **-5,23** | **-4,14** | 9 | 9 | -0,05 | 0,00 | 0,00 | (at2g40610):ATEXPA8 (ARABIDOPSIS THALIANA EXPANSIN A8) |
| At1g69530 | **0,00** | **0,00** | **-4,10** | **-3,07** | 9 | 9 | 0,00 | -2,07 | -1,71 | (at1g69530):ATEXPA1 (ARABIDOPSIS THALIANA EXPANSIN A1) |
| At1g20190 | **0,00** | **0,00** | **-3,71** | **-3,67** | 9 | 9 | -0,06 | 0,00 | 3,19 | (at1g20190):ATEXPA11 (ARABIDOPSIS THALIANA EXPANSIN A11) |
| At2g20750 | **0,00** | **0,00** | **-3,31** | **-2,35** | 2 | 2 | 0,51 | 0,00 | -0,52 | (at2g20750):ATEXPB1 (ARABIDOPSIS THALIANA EXPANSIN B1) |
| At4g01630 | **0,00** | **0,04** | **-2,49** | **-1,71** | 2 | 2 | 0,16 | 2,01 | -0,07 | (at4g01630):ATEXPA17 (ARABIDOPSIS THALIANA EXPANSIN A17) |
| At2g28950 | **0,00** | **0,00** | **-2,04** | **-2,06** | 2 | 2 | 0,70 | -1,03 | -0,45 | (at2g28950):ATEXPA6 (ARABIDOPSIS THALIANA EXPANSIN A6) |
| At2g39700 | **0,00** | **0,00** | **-1,98** | **-1,60** | 2 | 2 | 0,92 | 0,00 | -0,14 | (at2g39700):ATEXPA4 (ARABIDOPSIS THALIANA EXPANSIN A4) |
| At3g29030 | **0,00** | **0,00** | **-1,97** | **-2,16** | 2 | 2 | 0,99 | 0,00 | 0,60 | (at3g29030):ATEXPA5 (ARABIDOPSIS THALIANA EXPANSIN A5) |
| At5g56320 | **0,00** | **0,00** | **-1,94** | **-2,25** | 0 | 0 | 0,05 | 0,00 | -1,34 | (at5g56320):ATEXPA14 (ARABIDOPSIS THALIANA EXPANSIN A14) |
| At3g45960 | **0,02** | **0,00** | **-1,83** | **-2,59** | 0 | 0 | -0,02 | 0,00 | 1,64 | (at3g45960):ATEXLA3 (ARABIDOPSIS THALIANA EXPANSIN-LIKE A3) |
| At4g17030 | **0,00** | **0,00** | **-1,82** | **-2,47** | 2 | 2 | 0,20 | 0,00 | -0,84 | (at4g17030):ATEXLB1 (ARABIDOPSIS THALIANA EXPANSIN-LIKE B1) |
| At3g15370 | 0,56 | #N/A | -1,72 | #N/A | | | 0,03 | 0,00 | 0,75 | (at3g15370):ATEXPA12 (ARABIDOPSIS THALIANA EXPANSIN 12) |
| At3g45970 | **0,00** | **0,00** | **-1,62** | **-2,36** | 1 | 1 | -0,50 | 0,00 | 3,45 | (at3g45970):ATEXLA1 (ARABIDOPSIS THALIANA EXPANSIN-LIKE A1) |
| At2g03090 | **0,00** | **0,00** | **-1,62** | **-1,43** | 2 | 2 | 0,74 | 1,28 | -0,02 | (at2g03090):ATEXPA15 (ARABIDOPSIS THALIANA EXPANSIN A15) |
| At2g18660 | **0,00** | **0,00** | **-1,62** | **-0,68** | 0 | 0 | -0,02 | 2,96 | 0,64 | (at2g18660):expansin family protein (EXPR3) |
| At2g37640 | **0,00** | **0,00** | **-1,18** | **-2,07** | 2 | 2 | 0,18 | 0,00 | 2,17 | (at2g37640):ATEXPA3 (ARABIDOPSIS THALIANA EXPANSIN A3) |
| At5g02260 | **0,00** | **0,00** | **-1,17** | **-1,16** | 2 | 2 | 0,31 | 0,00 | 0,37 | (at5g02260):ATEXPA9 (ARABIDOPSIS THALIANA EXPANSIN A9) |
| At4g28250 | **0,00** | **0,00** | **-0,98** | **-1,07** | 2 | 2 | 1,03 | 0,00 | -0,76 | (at4g28250):ATEXPB3 (ARABIDOPSIS THALIANA EXPANSIN B3) |
| At3g03220 | **0,13** | **0,01** | -0,23 | **-0,36** | | 2 | 0,52 | 0,00 | -0,85 | (at3g03220):ATEXPA13 (ARABIDOPSIS THALIANA EXPANSIN A13) |
| At3g55500 | **0,48** | **0,48** | -0,17 | -0,21 | | | 0,11 | 0,00 | 0,14 | (at3g55500):ATEXPA16 (ARABIDOPSIS THALIANA EXPANSIN A16) |
| At1g26770 | **0,00** | **0,00** | **0,60** | **1,58** | 1 | 1 | 0,67 | 0,00 | -3,27 | (at1g26770):ATEXPA10 (ARABIDOPSIS THALIANA EXPANSIN A10) |
| At4g38400 | **0,00** | 0,12 | **0,63** | -0,30 | 2 | | -0,17 | 1,32 | 0,09 | (at4g38400):ATEXLA2 (ARABIDOPSIS THALIANA EXPANSIN-LIKE A2) |
| At5g39260 | 0,09 | 0,35 | 0,97 | 0,79 | | | 0,04 | 0,00 | 0,06 | (at5g39260):ATEXPA21 (ARABIDOPSIS THALIANA EXPANSIN A21) |
| At5g05290 | 0,30 | 0,17 | 1,09 | 1,08 | | | -0,03 | 0,00 | -0,19 | (at5g05290):ATEXPA2 (ARABIDOPSIS THALIANA EXPANSIN A2) |
| At1g65680 | 0,53 | 0,95 | 1,11 | -0,14 | | | -0,03 | 0,00 | -0,07 | (at1g65680):ATEXPB2 (ARABIDOPSIS THALIANA EXPANSIN B2) |
| At4g30380 | 0,09 | 0,20 | 2,55 | 1,98 | | | 0,02 | 0,00 | 0,07 | (at4g30380):expansin-related |
| At4g38210 | **0,00** | **0,00** | **4,59** | **4,65** | 0 | 0 | 0,01 | 0,00 | -2,08 | (at4g38210):ATEXPA20 (ARABIDOPSIS THALIANA EXPANSIN A20) |
| At1g12560 | **0,00** | **0,00** | **8,06** | **9,03** | 0 | 0 | -0,01 | 0,00 | 0,35 | (at1g12560):ATEXPA7 (ARABIDOPSIS THALIANA EXPANSIN A7) |
| At1g62980 | #N/A | 0,75 | #N/A | 0,60 | | | 0,11 | 0,00 | -0,90 | (at1g62980):ATEXPA18 (ARABIDOPSIS THALIANA EXPANSIN A18) |
| At2g45110 | #N/A | #N/A | #N/A | #N/A | | | 0,03 | 0,00 | -0,12 | at2g45110: ATEXPB4 (ARABIDOPSIS THALIANA EXPANSIN B4) |
| At3g60570 | #N/A | #N/A | #N/A | #N/A | | | -0,01 | 0,00 | 0,25 | (at3g60570):ATEXPB5 (ARABIDOPSIS THALIANA EXPANSIN B5) |
| At5g39270 | #N/A | #N/A | #N/A | #N/A | | | -0,04 | 0,00 | 0,11 | at5g39270: ATEXPA22 (ARABIDOPSIS THALIANA EXPANSIN A22) |
| At5g39280 | #N/A | #N/A | #N/A | #N/A | | | 0,00 | 0,00 | 0,09 | at5g39280: ATEXPA23 (ARABIDOPSIS THALIANA EXPANSIN A23) |
| At5g39310 | #N/A | #N/A | #N/A | #N/A | | | -0,03 | -2,54 | 0,86 | ARABIDOPSIS THALIANA EXPANSIN A24 |

B

| Gene | iTPS response | | | | | | CRF | Other responses | | Annotation |
|---|---|---|---|---|---|---|---|---|---|---|
| Code | 4-h adj.p | 6-h adj.p | **4-h FC $log_2$** | **6-h FC $log_2$** | 4-h CRF score | 6-h CRF score | | oeTPS FC $log_2$ | tSnRK1α1 FC $Log_2$ | |
| At5g65730 | 0,00 | 0,00 | **-3,04** | **-3,03** | 1 | 1 | -1,12 | 1,55 | 0,66 | (at5g65730):xyloglucan:xyloglucosyl transferase, putative |
| At1g10550 | 0,00 | 0,00 | **-2,69** | **-3,87** | 2 | 2 | 0,24 | 0,00 | 0,86 | (at1g10550):XTH33 (xyloglucan:xyloglucosyl transferase 33); |
| At1g32170 | 0,00 | 0,00 | **-2,67** | **-1,98** | 1 | 1 | -0,70 | 0,00 | 2,60 | (at1g32170):XTR4 (XYLOGLUCAN ENDOTRANSGLYCOSYLASE 4); |
| At5g57560 | 0,00 | 0,00 | **-1,81** | **-2,26** | 1 | 1 | -1,65 | 1,07 | 4,24 | (at5g57560):TCH4 (TOUCH 4); hydrolase, acting on glycosyl bonds |
| At1g11545 | 0,00 | 0,00 | **-1,80** | **-3,04** | 2 | 2 | 0,80 | 0,00 | 0,08 | (at1g11545):xyloglucan:xyloglucosyl transferase, putative |
| At3g25050 | 0,56 | #N/A | -1,72 | #N/A | | | 0,04 | 0,00 | 0,40 | (at3g25050):XTH3 (XYLOGLUCAN ENDOTRANSGLUCOSYLASE/HYDROLASE 3)s |
| At4g25820 | 0,56 | #N/A | -1,72 | #N/A | | | -0,04 | 3,42 | 2,07 | (at4g25820):XTR9 (XYLOGLUCAN ENDOTRANSGLYCOSYLASE 9); |
| At5g57550 | 0,00 | 0,00 | **-1,70** | **-1,99** | 1 | 1 | -1,66 | 1,03 | 1,59 | (at5g57550):XTR3 (XYLOGLUCAN ENDOTRANSGLYCOSYLASE 3); hydrolase, |
| At3g45970 | 0,00 | 0,00 | **-1,62** | **-2,36** | 1 | 1 | -0,50 | 0,00 | 3,45 | (at3g45970):ATEXLA1 (ARABIDOPSIS THALIANA EXPANSIN-LIKE A1) |
| At4g03210 | 0,00 | 0,00 | **-1,58** | **-2,15** | 0 | 0 | 0,07 | -1,32 | 0,06 | (at4g03210):XTH9 (XYLOGLUCAN ENDOTRANSGLUCOSYLASE/HYDROLASE 9); s |
| At4g25810 | 0,00 | 0,00 | **-1,33** | **-1,93** | 1 | 1 | -0,36 | 2,31 | 1,30 | (at4g25810):XTR6 (XYLOGLUCAN ENDOTRANSGLYCOSYLASE 6); hydrolase |
| At3g23730 | 0,00 | 0,00 | **-1,25** | **-1,59** | 2 | 2 | 0,30 | 0,00 | 0,44 | (at3g23730):xyloglucan:xyloglucosyl transferase, putative /e |
| At1g14720 | 0,00 | 0,00 | **-1,20** | **-1,14** | 1 | 1 | -0,14 | 0,00 | -1,14 | (at1g14720):XTR2 (XYLOGLUCAN ENDOTRANSGLYCOSYLASE RELATED 2); |
| At2g01850 | 0,00 | 0,00 | **-1,03** | **-1,06** | 1 | 1 | -0,74 | 0,00 | 0,73 | (at2g01850):EXGT-A3 (endo-xyloglucan transferase A3); hydrolase, |
| At4g37800 | 0,00 | 0,00 | **-1,03** | **-1,64** | 2 | 2 | 0,82 | 0,00 | -0,60 | (at4g37800):xyloglucan:xyloglucosyl transferase, putative |
| At5g48070 | 0,64 | 0,06 | -0,85 | -4,55 | | | 0,32 | 0,00 | 0,57 | (at5g48070):ATXTH20 (XYLOGLUCAN ENDOTRANSGLUCOSYLASE/HYDROLASE 20); |
| At1g65310 | 0,22 | 0,00 | -0,61 | **-2,26** | | 2 | 0,21 | 2,75 | 1,21 | (at1g65310):ATXTH17 (XYLOGLUCAN ENDOTRANSGLUCOSYLASE/HYDROLASE 17); |
| At4g18990 | 0,33 | 0,31 | -0,58 | -0,65 | | | -0,09 | 0,00 | 0,11 | (at4g18990):xyloglucan:xyloglucosyl transferase, putative / |
| At3g48580 | 0,34 | 0,04 | -0,45 | **-1,18** | | 0 | 0,02 | 0,00 | 0,44 | (at3g48580):xyloglucan:xyloglucosyl transferase, putative / |
| At2g14620 | 0,17 | 0,12 | -0,39 | -0,38 | | | -0,08 | 0,00 | 0,74 | (at2g14620):xyloglucan:xyloglucosyl transferase, putative / |
| At2g06850 | 0,01 | 0,00 | **-0,36** | **-1,69** | 1 | 1 | -0,15 | 0,00 | 0,26 | (at2g06850):EXGT-A1 (ENDO-XYLOGLUCAN TRANSFERASE); |
| At4g14130 | 0,77 | 0,00 | -0,13 | **-1,64** | | 1 | -0,57 | -1,18 | 0,09 | (at4g14130):XTR7 (XYLOGLUCAN ENDOTRANSGLYCOSYLASE 7); |
| At4g30280 | 0,93 | 0,00 | 0,05 | **-1,59** | | 0 | -0,02 | 3,03 | 0,45 | (at4g30280):ATXTH18/(XYLOGLUCAN ENDOTRANSGLUCOSYLASE/HYDROLASE 18) |
| At2g36870 | 0,00 | 0,00 | **0,33** | **0,44** | 1 | 1 | 0,60 | 0,00 | 0,14 | (at2g36870):xyloglucan:xyloglucosyl transferase, putative / |
| At5g13870 | 0,00 | 0,00 | **1,95** | **1,63** | 2 | 2 | -0,16 | 0,00 | 0,99 | (at5g13870):EXGT-A4 (ENDOXYLOGLUCAN TRANSFERASE A4); |
| At2g45110 | #N/A | #N/A | #N/A | #N/A | | | 0,03 | 0,00 | -0,12 | at2g45110: ATEXPB4 (ARABIDOPSIS THALIANA EXPANSIN B4) |
| At4g13080 | #N/A | #N/A | #N/A | #N/A | | | 0,00 | 0,00 | 0,30 | (at4g13080):xyloglucan:xyloglucosyl transferase, putative / |
| At4g13090 | #N/A | #N/A | #N/A | #N/A | | | -0,01 | 2,10 | 0,07 | (at4g13090):xyloglucan:xyloglucosyl transferase, putative / |
| At4g28850 | #N/A | #N/A | #N/A | #N/A | | | 0,02 | 0,00 | 0,11 | (at4g28850):xyloglucan:xyloglucosyl transferase, putative / |
| At5g57530 | #N/A | 0,62 | #N/A | 1,41 | | | 0,26 | 0,00 | 1,80 | (at5g57530):xyloglucan:xyloglucosyl transferase, putative / |
| At5g57540 | #N/A | #N/A | #N/A | #N/A | | | 0,28 | -2,65 | -0,78 | (at5g57540):xyloglucan:xyloglucosyl transferase, putative / |

**Supplemental Figure S19. Comparison of iTPS with constitutive overexpression of zBIP11.**

Sucrose acts at an uORF to inhibit translation of bZIP11, and falling sucrose allows synthesis of this $S_1$ type bZIP protein that then interacts with C-type bZIPs to transcriptionally activate starvation responses and inhibit growth (Hanson et al., 2009; Ma et al., 2011, Dröge-Laser and Weiste, 2018).

Ma *et al.* (2011) reported compared to wild-type Arabidopsis, 232 transcripts that showed a significant change and passed a FC filter of $\log_2$ >1 in lines with constitutive overexpression of *bZIP11* (oebZIP11). This published response to oebZIP11 was compared with the iTPS response.

Panels A-C are below, Panels D-G are on the following pages

A

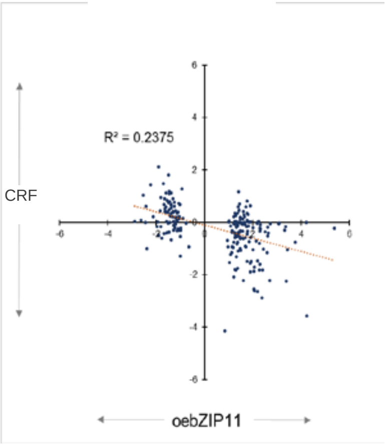

**Relationship between the oebZIP11 response and the carbon response factor (CRF)** (estimated for each gene from nine treatments in which sugar levels were changed (Supplemental Figure S2).

B

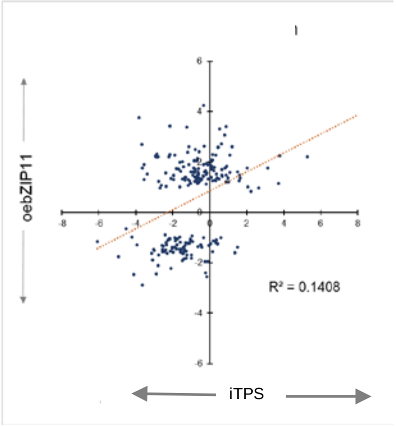

**Relationship between the oebZIP11 response and the overall iTPS response.** The iTPS data set was not filtered, in order to show not only the response of transcripts that changed but also of any that did not change. The iTPS response is the average of the 4-h and 6-h post-induction responses.

C **Relationship between oebZIP11 and the iTPS response after deconvolution to assign transcripts to CRF groups $G_1$, $G_2$ and $G_0$ (from left to right).** The iTPS response is the average of the 4-h and 6-h post-induction response. The number in brackets is the number of genes in each subset

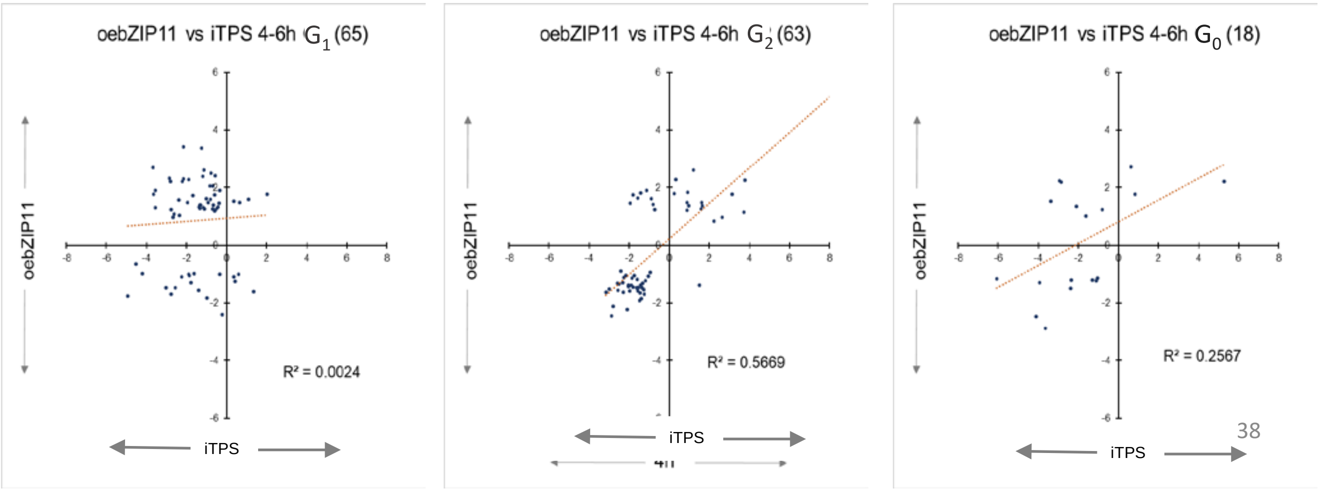

**Supplemental Figure S19. Comparison of iTPS with constitutive overexpression of zBIP11 (continued)**

D **oebZIP11-responsive genes: comparison of the iTPS response and the response to constitutive oeTPS** (Zhang et al., 2009). The data set of Zhang et al. (2009) was filtered by the authors (FDR<0.05, FC < $\log_2 1$) and contained 94 of the 232 genes that Ma et al. (2009) identified as oebZIP11-responsive. The upper panel shows the relationship between iTPS and oeTPS for all 94 shared genes, the middle panel shows the relationship between iTPS and oeTPS for the 43 genes that were assigned to CRF $G_1$ in the iTPS response, and the lower panel shows the relationship between iTPS and oeTPS for the 30 genes that were assigned to CRF $G_2$ in the iTPS response. The iTPS response is the average of the 4-h and 6-h post-induction iTPS responses.

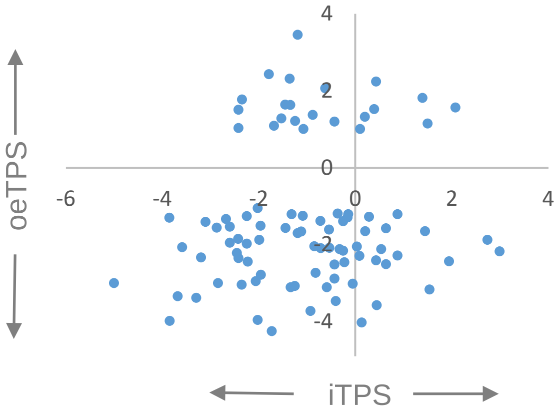

All genes in the constitutive oeTPS response

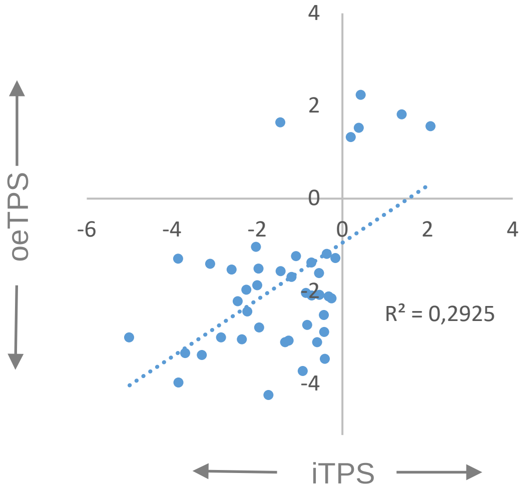

Genes assigned to CRF group $G_1$ in the iTPS response

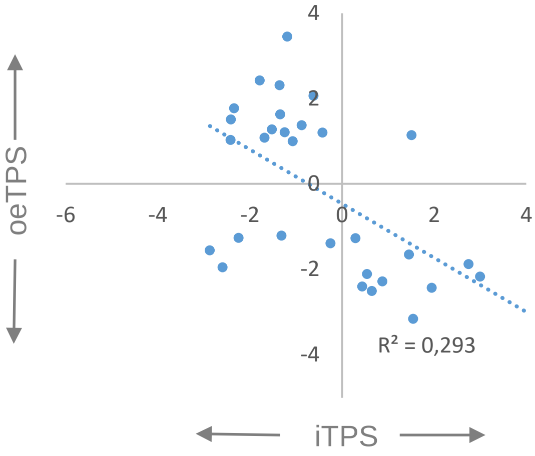

Genes assigned to CRF group $G_2$ in the iTPS response

#### Supplemental Figure S19. Comparison of iTPS with constitutive overexpression of zBIP11 (continued)

(E) **List of oebZIP1 responsive genes that are included in genes assigned to CRF group $G_1$ in the iTPS response.** The genes are separated into those whose transcripts show a qualitatively similar response in oebZIP11 and iTPS (upper part, blue shaded) and those that show an opposite response (lower part, pink-shaded). The display showsthe 4-h and 6-h iTPS responses, assignment of the iTPS response to CRF groups, the CRF value (defined as in Supplemental Figure S2), the response to constitutive overexpression of TPS (oeTPS, from Zhang et al., 2009) and to transient overexpression of tSnRL1α1 in protoplasts (tSnRL1α1, from Baena-Gonzalez et al., 2007) and to oebZIP11. Significant changes in the iTPS response are shown with bold font. Assignment to CRF groups is performed only for significant responses and only CRF $G_1$ assignments are shaded. No entry for oeTPS or tSnRL1α1 indicates that the response did not pass the filter in the original publication

E

| Gene | | iTPS response | | | | | | CRF | Other responses | | | | Ontology and annnotation |
|---|---|---|---|---|---|---|---|---|---|---|---|---|---|
| Name | Code | 4-h adj.p | 6-h adj.p | 4-h FC $log_2$ | 6-h FC $log_2$ | 4-h CRF score | 6-h CRF score | | oeTPS FC $log_2$ | tSnRK1α1 FC $Log_2$ | oebZIP11 | direction bZIP11 compared to iTPS | |
| GAS6 | At1g74670 | 0,00 | 0,00 | -4,92 | -5,07 | 1 | 1 | -0,60 | -3,00 | | **-1,76** | same | GA-STIMULATED 6 Cell wall protein downstream of RGL2, integrating GA, ABA, and Glc signaling |
| SWEET12 | At5g23660 | 0,00 | 0,00 | -4,53 | -3,16 | 1 | 1 | -0,93 | -1,30 | | **-0,66** | same | SWEET 12, sucrose expoert from leaf |
| NPH3 protein | At3g19850 | 0,00 | 0,00 | -4,19 | -3,17 | 1 | 1 | -0,31 | -3,34 | | **-0,98** | same | Phototropic-responsive **NPH3** family protein |
| WRKY17 | At5g24570 | 0,00 | 0,00 | -3,02 | -2,80 | 1 | 1 | -0,36 | | | **-1,47** | same | WRKY DNA-BINDING PROTEIN 17 |
| HSP20-like | At4g21870 | 0,00 | 0,00 | -2,76 | -2,92 | 1 | 1 | -0,42 | -3,00 | | **-1,68** | same | HEAT SHOCKr PROTEIN20 like |
| NPF6.2 | At2g26690 | 0,00 | 0,00 | -2,55 | -2,63 | 1 | 1 | -0,38 | -1,54 | | **-1,47** | same | NRT1/PTR anion transporter, Major facilitator superfamily protein |
| | At5g62280 | 0,00 | 0,00 | -1,76 | -2,56 | 1 | 1 | -0,12 | | | **-1,29** | same | DUF1442 family protein |
| SAUR14 | At4g38840 | 0,00 | 0,00 | -2,25 | -1,73 | 1 | 1 | -0,25 | -1,88 | | **-1,09** | same | SMALL AUXIN UPREGULATED RNA 14 |
| NAC2/ORESARA1 | At5g39610 | 0,00 | 0,01 | -1,89 | -1,16 | 1 | 1 | -1,30 | | | **-1,01** | same | NAC20 TF, positively regulates leaf senescence, also abiotic stress responses, affects primary root |
| TN13 | At3g04210 | 0,00 | 0,00 | -1,86 | -1,04 | 1 | 1 | -0,30 | 1,64 | | **-1,05** | same | TIR-NBS protein, contributes to RPS5-triggered immunity. |
| ARR9 | At3g57040 | 0,00 | 0,00 | -1,40 | -1,49 | 1 | 1 | -0,38 | -1,57 | 1,54 | **-1,56** | same | Response regulator ARR9, cytokin signalling |
| Zn Finger | At5g60710 | 0,00 | 0,00 | -1,63 | -1,26 | 1 | 1 | -0,23 | | | **-0,98** | same | Zinc finger (C3HC4-type RING finger) family protein, UBQ-ligase E3 |
| JAZ1 | At1g72450 | 0,00 | 0,00 | -0,97 | -0,75 | 1 | 1 | -0,16 | | | **-1,82** | same | jasmonate-zim-domain protein 6, TIFY DOMAIN PROTEIN 11B |
| | At5g25840 | 0,00 | 0,00 | -0,34 | -0,87 | 1 | 1 | -0,54 | | | **-1,00** | same | DUF1677 family protein |
| FLZ13 | At1g74940 | 0,59 | 0,00 | -0,09 | -0,40 | | 1 | -0,17 | | | **-1,96** | same | |
| ALDH6B2 | At2g14170 | 0,04 | 0,04 | -0,21 | -0,28 | 1 | 1 | -1,00 | -2,16 | 2,05 | **-2,39** | same | Methylmalonate-semialdehyde dehydrogenase |
| | At2g39130 | 0,00 | 0,47 | 0,34 | 0,11 | 1 | | 0,34 | | | **1,26** | same | Transmembrane amino acid transporter family protein; C |
| | At2g14835 | 0,00 | 0,01 | 0,36 | 0,34 | 1 | 1 | 0,14 | | | **1,53** | same | |
| | At5g17760 | 0,73 | 0,00 | 0,09 | 0,78 | | 1 | 0,58 | 2,23 | -1,98 | **1,72** | same | P-loop containing nucleoside triphosphate hydrolases superfamily protein |
| APC2 | At2g04660 | 0,00 | 0,00 | 0,66 | 0,80 | 1 | 1 | 0,20 | | | **1,47** | same | ANAPHASE-PROMOTING COMPLEX/CYCLOSOME 2, cell cycle |
| ALKBH10A | At2g48080 | 0,01 | 0,00 | 1,09 | 1,11 | 1 | 1 | 0,14 | | | **1,58** | same | 2OG-Fe (II) oxygenase family protein; mRNA demethylation |
| SAUR30 | At5g53590 | 0,00 | 0,00 | 2,02 | 2,13 | 1 | 1 | 0,35 | 1,56 | 1,95 | **1,76** | same | SMALL AUXIN UPREGULATED RNA 30 |
| BT2 | At3g48360 | 0,00 | 0,00 | -3,66 | -4,02 | 1 | 1 | -2,22 | -3,98 | 1,97 | **2,70** | opposite | BT2 is an essential component of the TAC1-mediated telomerase activation pathway. |
| | At2g25200 | 0,00 | 0,00 | -3,54 | -3,43 | 1 | 1 | -1,82 | | 2,20 | **1,31** | opposite | |
| BXL1 | At5g49360 | 0,00 | 0,00 | -2,80 | -3,78 | 1 | 1 | -2,64 | -3,38 | 5,06 | **2,21** | opposite | BETA-XYLOSIDASE 1, rhamnogalacturonan I.modification and degradation. |
| | At3g26510 | 0,00 | 0,00 | -3,55 | -2,65 | 1 | 1 | -1,57 | -1,41 | 2,71 | **1,89** | opposite | Octicosapeptide/Phox/Bem1p family protein; |
| NAC2 | At3g15510 | 0,00 | 0,00 | -3,65 | -2,27 | 1 | 1 | -0,46 | | | **1,75** | opposite | NAC transcription factor |
| | At1g62510 | 0,00 | 0,00 | -2,62 | -2,78 | 1 | 1 | -1,22 | | 3,74 | **1,09** | opposite | Expressed in root cortex |
| GRXS6 | At3g62930 | 0,00 | 0,00 | -2,39 | -2,50 | 1 | 1 | -0,61 | -2,22 | | **1,57** | opposite | CC-type glutaredoxin (ROXY) family, interacts wth TF TGA2 and suppress ORA59 promoter activity |
| | At5g11070 | 0,00 | 0,00 | -2,34 | -2,36 | 1 | 1 | -1,53 | -3,04 | | **1,03** | opposite | hypothetical protein |
| XTH30 | At1g32170 | 0,00 | 0,00 | -2,67 | -1,98 | 1 | 1 | -0,70 | | 2,60 | **0,96** | opposite | XYLOGLUCAN ENDOTRANSGLUCOSYLASE/HYDROLASE 30, cell wall modification |
| PRX47 | At4g33420 | 0,00 | 0,00 | -2,77 | -1,80 | 1 | 1 | -0,37 | | | **1,23** | opposite | Peroxidase superfamily protein |
| BBE18 | At4g20820 | 0,00 | 0,00 | -2,84 | -1,65 | 1 | 1 | -0,24 | -1,97 | | **2,32** | opposite | FAD-binding Berberine family protein |
| TRE1 | At4g24040 | 0,00 | 0,00 | -2,14 | -2,30 | 1 | 1 | -1,04 | -2,44 | | **3,43** | opposite | TREHALASE 1 |
| PLL12 | At5g04310 | 0,00 | 0,00 | -1,88 | -2,17 | 1 | 1 | -0,23 | -1,05 | | **2,28** | opposite | PECTATE LYASE LIKE12, cell wall modification |
| OTU1 | At2g28120 | 0,00 | 0,00 | -1,96 | -1,96 | 1 | 1 | -1,00 | -1,51 | | **1,47** | opposite | OVARIAN TUMOR DOMAIN (OTU)-CONTAINING DUB (DEUBIQUITILATING ENZYME) 1 |
| | At5g08350 | 0,00 | 0,00 | -2,22 | -1,68 | 1 | 1 | -1,48 | -2,79 | | **2,21** | opposite | |
| DOF1.8 | At1g64620 | 0,00 | 0,00 | -2,16 | -1,57 | 1 | 1 | -0,10 | | | **2,30** | opposite | DOF TF which regulates vascular cell differentiation and lignin biosynthesis |
| RZPF34, | At5g22920 | 0,00 | 0,00 | -1,19 | -2,26 | 1 | 1 | -2,88 | -4,25 | 6,76 | **2,38** | opposite | CHY ZINC-FINGER AND RING PROTEIN 1, |
| HB-12 | At3g61890 | 0,00 | 0,00 | -1,68 | -1,64 | 1 | 1 | -0,93 | | 2,66 | **1,72** | opposite | HOMEOBOX 12 |
| COR47 | At1g20440 | 0,00 | 0,00 | -1,31 | -1,78 | 1 | 1 | -0,51 | | | **1,38** | opposite | COLD-REGULATED 47 |
| | At4g27657 | 0,00 | 0,00 | -1,24 | -1,63 | 1 | 1 | -0,17 | | | **3,37** | opposite | hypotheticl protein |
| CHX17 | At4g23700 | 0,00 | 0,00 | -1,47 | -1,39 | | 1 | -0,18 | | | **1,32** | opposite | CATION/H+ EXCHANGER 17, |
| SCPL31, | At1g11080 | 0,00 | 0,00 | -1,36 | -1,35 | 1 | 1 | -0,50 | | | **1,33** | opposite | SERINE CARBOXYPEPTIDASE-LIKE 31 |
| PPDK | At4g15530 | 0,00 | 0,00 | -1,48 | -1,23 | | 1 | -1,06 | | 2,24 | **2,00** | opposite | PYRUVATE ORTHOPHOSPHATE DIKINASE |
| SEN1 | At4g35770 | 0,22 | 0,00 | -0,33 | -2,34 | | 1 | -3,56 | -3,11 | 3,76 | **4,25** | opposite | Senescence-associated gene that is strongly induced by phosphate starvation |
| BTB/POZ | At2g30600 | 0,00 | 0,00 | -1,33 | -1,17 | 1 | 1 | -1,81 | -3,07 | 3,85 | **1,27** | opposite | BTB/POZ domain-containing proteins |
| | At4g18340 | 0,00 | 0,00 | -1,32 | -1,06 | 1 | 1 | -1,08 | -1,70 | | **1,38** | opposite | Glycosyl hydrolase superfamily protein; |
| | At1g12080 | 0,00 | 0,00 | -0,80 | -1,37 | 1 | 1 | -0,33 | -1,25 | | **1,58** | opposite | Vacuolar calcium-binding protein-like protein; |
| NPY3 | At5g67440 | 0,00 | 0,00 | -1,15 | -0,77 | 1 | 1 | -0,52 | | | **1,30** | opposite | NAKED PINS IN YUC MUTANTS Family (1-5) involved in auxin-mediated organogenesis. |
| PP2-A13 | At3g61060 | 0,00 | 0,00 | -0,70 | -1,15 | 1 | 1 | -2,56 | -3,73 | 2,99 | **2,06** | opposite | PHLOEM PROTEIN 2-A13, PP2-A13 |
| NAC047 | At3g04070 | 0,00 | 0,00 | -1,11 | -0,60 | 1 | 1 | -0,59 | | | **1,26** | opposite | NAC DOMAIN CONTAINING PROTEIN 47, SPEEDY HYPONASTIC GROWTH |
| UGT88A1 | At3g16520 | 0,00 | 0,00 | -1,14 | -0,56 | 1 | 1 | -0,19 | -2,04 | | **2,61** | opposite | UDP-GLUCOSYL TRANSFERASE 88A1 |
| XMPP | At2g32150 | 0,00 | 0,00 | -1,01 | -0,63 | 1 | 1 | -2,21 | -2,74 | 2,75 | **1,60** | opposite | xanthosine monophosphate (XMP) phosphatase. Initial step in purine nucleotide catabolism |
| HMP37 | At4g27590 | 0,03 | 0,98 | -1,62 | 0,02 | 1 | | -0,20 | | | **1,99** | opposite | HEAVY METAL ASSOCIATED PROTEIN 37 Heavy metal transport/detoxification superfamily |
| HAT2 | At5g47370 | 0,00 | 0,00 | -0,60 | -0,98 | 1 | 1 | -0,16 | | | **1,75** | opposite | Homeobox-leucine zipper genes induced by auxin, but not by other phytohormones. |
| CHX20 | At3g53720 | 0,00 | 0,00 | -0,70 | -0,85 | 1 | 1 | -0,41 | | | **1,26** | opposite | Member of putative Na+/H+ antiporter family. Omoregulation through $K^+$ flux, possibly pH modulation |
| SDP1 | At5g04040 | 0,00 | 0,00 | -0,94 | -0,51 | 1 | 1 | -1,01 | -1,39 | 2,49 | **1,48** | opposite | SUGAR-DEPENDENT1 Triacylglycerol lipase involved in storage lipid breakdown in seed germination |
| BAM9 | At5g18670 | 0,00 | 0,00 | -0,84 | -0,59 | 1 | 1 | -1,87 | -2,10 | 3,59 | **2,06** | opposite | BETA-AMYLASE 9, positive regulation of starch breakdown |
| PDH2 | At5g38710 | 0,02 | 0,01 | -0,65 | -0,76 | 1 | | -0,76 | | | **1,36** | opposite | Methylenetetrahydrofolate reductase family protein, prolone catabolism |
| GOLS | At1g56600 | 0,04 | 0,51 | -1,03 | -0,31 | 1 | | -0,31 | | | **1,14** | opposite | galactinol synthase |
| | At4g30490 | 0,00 | 0,00 | -0,79 | -0,47 | 1 | 1 | -0,37 | | | **2,50** | opposite | AFG1-like ATPase family protein |
| | At1g21680 | 0,19 | 0,00 | -0,20 | -0,97 | | 1 | -1,49 | -3,11 | | **1,95** | opposite | DPP6 N-terminal domain-like protein |
| CMCU | At5g66650 | 0,01 | 0,09 | -0,65 | -0,48 | 1 | | -0,72 | | 3,35 | **2,14** | opposite | CHLOROPLAST-LOCALIZED MITOCHONDRIAL CALCIUM UNIPORTER, |
| | At3g47000 | 0,00 | 0,00 | -0,66 | -0,43 | 1 | 1 | -0,51 | -1,61 | | **1,40** | opposite | Glycosyl hydrolase family protein; |
| ZIF2 | At2g48020 | 0,00 | 0,00 | -0,58 | -0,48 | 1 | 1 | -0,76 | -2,08 | | **1,22** | opposite | ZINC-INDUCED FACILITATOR transport protein |
| SIP2 | At2g30360 | 0,00 | 0,11 | -0,68 | -0,37 | 1 | | -0,54 | | | **1,45** | opposite | CBL-interacting protein kinase. Regulates $H^+$ transporter.Also ABI5, phosphorylation induced by ABA, |
| | At3g04010 | 0,00 | 0,68 | -1,15 | 0,11 | 1 | | -0,15 | | | **1,07** | opposite | O-Glycosyl hydrolases family 17 protein |
| ACX2 | At5g65110 | 0,00 | 0,01 | -0,59 | -0,35 | 1 | 1 | -0,97 | | | **1,20** | opposite | ACYL-COA OXIDASE 2 |
| DIN4 | At3g13450 | 0,03 | 0,00 | -0,34 | -0,52 | 1 | 1 | -1,89 | -2,52 | 4,02 | **1,90** | opposite | branched chain alpha-keto acid dehydrogenase E1 beta |
| ACR9 | At2g39570 | 0,00 | 0,03 | -0,57 | -0,28 | 1 | 1 | -1,83 | -2,89 | 1,98 | **2,40** | opposite | ACT DOMAIN REPEATS 9. ACT domain is an amino acid-binding site in feedback-regulation of amino acid metabolic enzymes |
| | At2g36310 | 0,00 | 0,00 | -0,48 | -0,35 | 1 | 1 | -0,48 | | | **1,29** | opposite | Cytoplasmic nucleoside hydrolase, Inosine/uridine-preferring nucleoside hydrolase |
| ACHT5 | At5g61440 | 0,67 | 0,00 | -0,09 | -0,72 | | 1 | -1,31 | -3,47 | | **1,32** | opposite | Encodes a member of the thioredoxin family protein, chloroplast located |
| SHL | At4g39100 | 0,00 | 0,02 | -0,43 | -0,30 | 1 | 1 | -0,57 | -1,19 | | **1,33** | opposite | SHORT LIFE. Plant-specific histone reader capable of recognizing both H3K27me3 and H3K4me3 |
| XER | At2g04240 | 0,02 | 0,04 | -0,37 | -0,34 | 1 | 1 | -0,39 | | | **1,46** | opposite | XERICO |
| LNK2 | At3g54500 | 0,00 | 0,62 | -0,55 | -0,08 | 1 | | -0,82 | -2,12 | | **1,34** | opposite | LINK2 Member of a small family. Along with LNK1 integrates light signaling and circadian clock. |
| | At1g63840 | 0,00 | 0,49 | -0,66 | 0,14 | 1 | | -0,38 | | | **1,15** | opposite | RING/U-box superfamily protein |
| RMR2 | At1g71980 | 0,00 | 0,54 | -0,31 | -0,10 | 1 | | -0,40 | | | **2,64** | opposite | Secretory pathway protein localized to the trans-golgi network. family of vacuolar sorting receptors. |
| | At3g43430 | 0,02 | 0,25 | -0,59 | 0,28 | 1 | | -0,35 | -1,29 | | **1,55** | opposite | RING-H2-type E3 ligase |
| LSH6 | At1g07090 | 0,79 | 0,01 | 0,04 | 0,36 | | 1 | 0,60 | 1,32 | | **-1,25** | opposite | LIGHT SENSITIVE HYPOCOTYLS 6, |
| TWN2 | At1g14610 | 0,00 | 0,01 | 0,37 | 0,41 | 1 | 1 | 0,39 | 1,52 | | **-1,10** | opposite | VALYL TRNA SYNTHETASE |
| TBL27 | At1g70230 | 0,00 | 0,00 | 0,43 | 0,47 | 1 | 1 | 0,76 | | -1,84 | **-1,25** | opposite | ALTERED XYLOGLUCAN 4, AXY4, TRICHOME BIREFRINGENCE-LIKE 27 |
| EXPA3 | At1g26770 | 0,00 | 0,00 | 0,60 | 1,58 | 1 | 1 | 0,67 | | -3,27 | **-1,01** | opposite | EXPANSIN A3 |
| HSP70-4 | At3g12580 | 0,00 | 0,00 | 1,36 | 1,43 | 1 | 1 | 0,58 | 1,81 | | **-1,59** | opposite | HEAT SHOCK PROTEIN 70-4 |

**Supplemental Figure S19. Comparison of iTPS with constitutive overexpression of zBIP11 (continued)**

F **Transcripts that are present in the oebZIP11 gene set of Ma et al. (2009) and are assigned to the iTPS Crf group $G_1$: comparison of the response to oebZIP11 with the responses to iTPS or to constitutive oeTPS (from Zhang et al., 2009)**

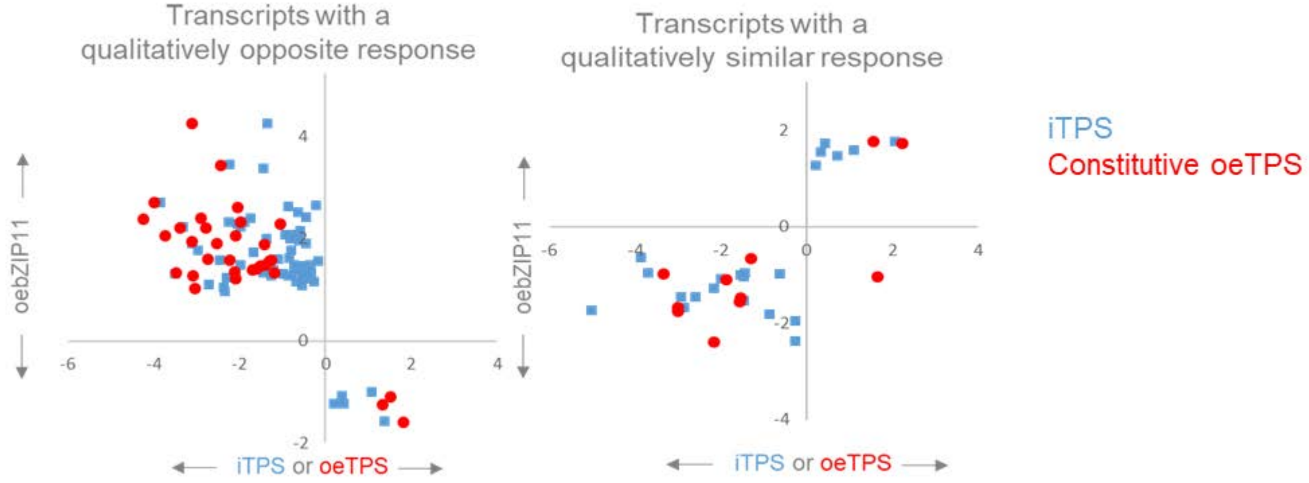

G **Transcripts that are present in the oebZIP11 gene set of Ma et al. (2009): Comparison of their iTPS and their tSnRK1α1 response. (from Baena-Gonález et al. 2007).** This is done from left to right) or all transcripts and for transcripts assigned to CRF group $G_1$ and group $G_2$

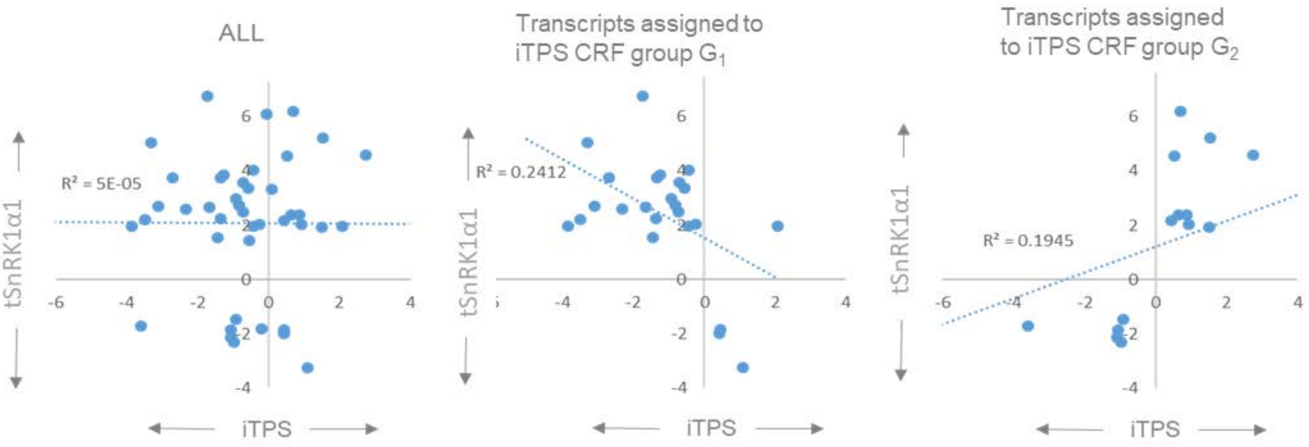

H **Transcripts that are present in the oebZIP11 gene set of Ma et al. (2009): comparison of their tSnRK1α1 response and their oebZIP11 response**. This is done from left to right) or all transcripts and for transcripts assigned to CRF group $G_1$ and group $G_2$

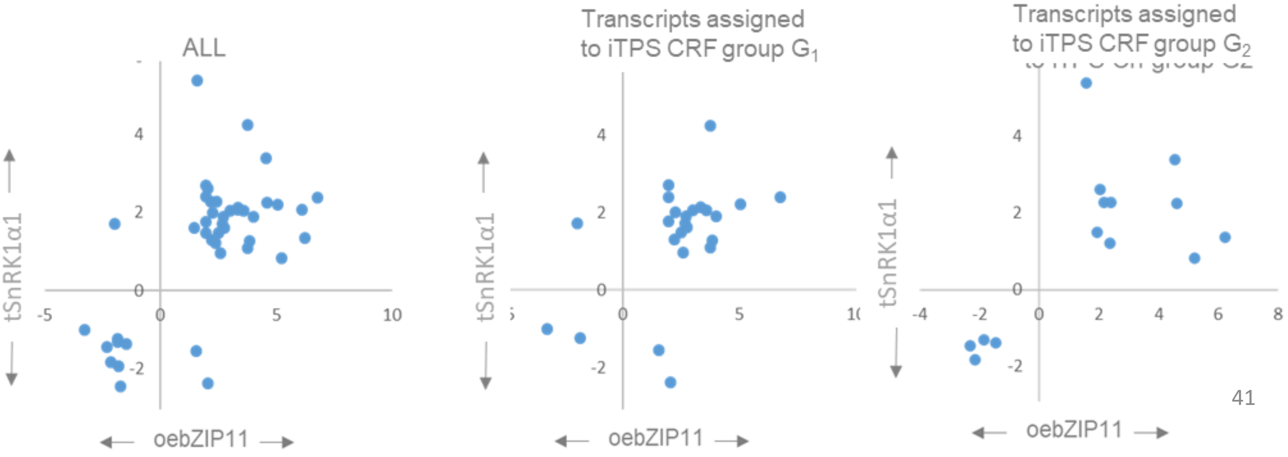

J **Response of trehalose-metabolism related genes.** The display shows the 4-h and 6-h iTPS response, assignment of the iTPS response to CRF groups, the CRF value (defined as in Supplemental Figure S12), the response to constitutive overexpression of TPS (oeTPS, from Zhang et al., 2009) and to transient overexpression of tnRL1α1 in protoplasts (tSnRL1α1, from Baena-Gonzalez et al., 2007) and to oebZIP11. Significant changes in the iTPS response are shown with bold font. Assignment to CRF groups is performed only for significant responses and only CRF $G_1$ assignments are shaded. No entry for oeTPS or tSnRL1α1indicates that the response did not pass the filter in the publication.

| **Gene** | | **iTPS response** | | | | | | **CRF** | **Other responses** | | | | **Comment** |
|---|---|---|---|---|---|---|---|---|---|---|---|---|---|
| Name | Code | 4-h adj.p | 6-h adj.p | **4-h FC log$_2$** | **6-h FC log$_2$** | 4-h CRF score | 6-h CRFscore | | oeTPS FC log$_2$ | tSnRKα1 FC Log$_2$ | **oe bZIP11** | **direction, bZIP11 compared to iTPS** | |
| TRE1 | At4g24040 | 0,00 | 0,00 | -2,14 | -2,30 | 1 | 1 | -1,04 | -2,44 | | **3,43** | opposite | responses in iTPS and constituve oeTPS are in same direction |
| TPPG | At4g22590 | 0,00 | 0,00 | 0,83 | 0,94 | 0 | 0 | 0,09 | -1,21 | | **1,77** | same | responses in iTPS and constituve oeTPS in opposed directions |
| TPPF | At4g12430 | 0,38 | 0,54 | 0,16 | 0,12 | | | -0,09 | -4,03 | | **1,26** | none | no response in iTPS and large decrease in oeTPS |

**Supplemental Figure S20. Transcription factors that respond to an induced increase in Tre6P levels.**

(A) Examples of TFs that showed a strong response and were assigned to CRF G group G1.

(B-E) Transcription factor listed according to their family: (B) bZIPS, (C) WRKYs, (D) AP2/ERF, ED) MYBs. These panels show the 4-h and 6-h iTPS response. All changes were significant (FDR<0.05) and a filter was set for FC<$\log_2$1 for at least one time point. Responses are listed separately for different CRF groups. For comparison, the published responses to constitutive overexpression of TPS in Arabidopsis is provided (oeTPS, from Zhang et al., 2009; a missing value means that the data is not available).

A

| ID | iTPS 4h | iTPS 6h | Name | Description |
|---|---|---|---|---|
| At5g57520 | -4.6 | -2.7 | ZFP2 | negative regulation of floral organ abscission |
| At5g59780 | -4.3 | -3.5 | MYB59 | cellular response to potassium ion. In roots it is involved in K+/NO3- transport and expression of the NPF7.3 transporter |
| At3g46130 | -4.2 | -3.3 | MYB48 | regulate flavonol biosynthesis primarily in cotyledons |
| At2g18300 | -4.1 | -3.1 | HBI1 | bHLH protein involved in positive regulation of cell elongation and proliferation and, negative control of plant immunity. BR signaling. |
| At1g69490 | -3.8 | -1.9 | NAP | It is expressed in floral primordia and upregulated by AP3 and PI |
| At5g61590 | -3.7 | -3.9 | DEWAX | ERF107. preferentially expressed in the epidermis and induced by darkness and negatively regulates cuticular wax AND anthocyanins biosynthesis |
| At3g15510 | -3.7 | -2.3 | ANAC2 | putative regulator of phloem parenchyma wall ingrowth deposition |
| At3g29035 | -3.2 | -2.4 | ANAC059 | contains CCT domain (CONSTANS like) |
| At3g54990 | -3.2 | -2.0 | SMZ | Encodes a AP2 domain transcription factor that can repress flowering. SMZ and its paralogous gene, SNARCHZAPFEN (SNZ), share a signature with partial complementarity to the miR172 microRNA, whose precursor is induced upon flowering |
| At4g00050 | -3.2 | -2.8 | PIF8 | Encodes a phytochrome interacting factor that inhibits phytochrome A-mediated far-red light responses and binds to promoter regions of AT2G46970 and AT3G62090. |
| At5g15830 | -3.1 | -2.3 | bZIP3 | bZIP3) is a novel sugar-responsive transcription factor in Arabidopsis plants. The expression of bZIP3 was rapidly repressed by sugar. Genetic analysis indicated that bZIP3 expression was modulated by the SNF1-RELATED KINASE 1 (SnRK1) pathway |
| At5g56860 | -3.1 | -2.2 | GATA21 | GATA, nitrate-inducible, carbon metabolism-involved |
| At2g20180 | -2.7 | -2.4 | PIF1 | Inhibited by light, gibberellin biosynthesis. key negative regulator of phytochrome-mediated seed germination and acts by inhibiting chlorophyll biosynthesis, light-mediated suppression of hypocotyl elongation and far-red light-mediated suppression of seed germination, and promoting negative gravitropism in hypocotyls |
| At1g79700 | -2.7 | -2.4 | WRI4 | specifically controls cuticular wax biosynthesis. functions to activate transcription of genes involved fatty acid biosynthesis during seed and flower development as well as stem wax biosynthesis |
| At4g17460 | -2.7 | -2.7 | HAT1 | regulates meristematic activity |
| At1g66230 | -2.6 | -2.4 | MYB20 | directly activates lignin biosynthesis genes and phenylalanine biosynthesis genes during secondary wall formation. down-regulates the expression of PP2Cs, the negative regulator of ABA signaling, and enhances salt tolerance |
| At1g71030 | -2.2 | -1.5 | MYBL2 | anthocyanin-containing compound biosynthetic process |
| At1g25550 | -2.1 | -2.0 | HHO3 | Transcriptional repressors that functions with other NIGT genes as an important hub in the nutrient signaling network associated with the acquisition and use of nitrogen and phosphorus. |
| At5g65210 | -1.9 | -1.3 | TGA1 | important regulatory factors of the nitrate response in Arabidopsis roots via NRT2.1 and NRT2.2 |
| At5g37260 | -1.6 | -1.7 | CIR1 | RVE2, Involved in circadian regulation |
| At5g49450 | -1.6 | -1.9 | bZIP1 | A positive regulator of plant tolerance to salt, osmotic and drought stresses, crucial transcriptional regulators in Pro, Asn, and branched-chain amino acid metabolism. Expression is repressed by sugars, mediated by HEX. |
| At5g28770 | -1.5 | -1.6 | bZIP63 | Phosphorylated by SnRK1. composes a regulatory interface between the metabolic and circadian control of starch breakdown to optimize C usage and plant growth. Hexokinase 1 is required for glucose-induced repression of bZIP63 |
| At4g01120 | -1.5 | -1.6 | bZIP54 | glucosinolate metabolic process, response to blue light |
| At2g18160 | -1.5 | -1.5 | bZIP2 | GBF5, FLORAL TRANSITION AT THE MERISTEM3 |
| At1g79430 | -1.2 | -1.4 | APL | required for several aspects of phloem development in the root: (1) the specific divisions organizing the phloem pole, (2) sieve element differentiation and (3) the expression of a companion-specific gene |
| At1g18330 | -1.2 | -1.3 | EPR1 | RVE7, EARLY-PHYTOCHROME-RESPONSIVE1 |
| At5g11260 | -1.0 | -1.0 | HY5 | photomorphogenesis (through UVR8 signaling pathway) |
| At1g56650 | 0.7 | 1.5 | PAP1 | involved in anthocyanin metabolism and radical scavenging, Essential for the sucrose-mediated expression of the dihydroflavonol reductase gene |
| At1g62150 | 1.5 | 1.4 | mTERF20 | Mitochondrial transcription termination factor familiy member, active in chloroplasts |
| At1g17460 | 1.7 | 1.4 | TRFL3 | TRFL3, negative regulation of gene expression |

**Supplemental Figure S20. Transcription factors that respond to an induced increase in Tre6P levels (continued).**

#### B. bZIPs

$G_1$

| ID | iTPS 4h | iTPS 6h | oeTPS | Name |
|---|---|---|---|---|
| At5g15830 | -3.13 | -2.31 | n.d | bZIP3 |
| At5g65210 | -1.91 | -1.28 | n.d | TGA1 |
| At2g22850 | -1.76 | -1.97 | -2.16 | bZIP6 |
| At3g51960 | -1.75 | -2.12 | n.d | bZIP24 |
| At5g49450 | -1.55 | -1.88 | -3.35 | bZIP1 |
| At5g28770 | -1.53 | -1.65 | -2.86 | bZIP63 |
| At4g01120 | -1.52 | -1.64 | n.d | bZIP54 |
| At2g18160 | -1.49 | -1.45 | -2.09 | bZIP2 |
| At1g13600 | -1.29 | -1.72 | n.d | bZIP58 |
| At5g24800 | -1.18 | -0.67 | n.d | bZIP9 |
| At5g11260 | -1.02 | -0.99 | -1.09 | HY5 |
| At2g40950 | 0.73 | 1.12 | N.D | bZIP17 |

$G_2$

| ID | iTPS 4h | iTPS 6h | oeTPS | Name |
|---|---|---|---|---|
| At2g42380 | -3.78 | -3.19 | -1.97 | bZIP34 |
| At3g58120 | -3.22 | -3.10 | n.d | bZIP61 |
| At5g10030 | -2.42 | -1.93 | -1.26 | OBF4 |
| At1g06850 | -1.46 | -1.44 | n.d | bZIP52 |
| At1g68640 | -1.03 | -1.39 | n.d | TGA8 |
| At4g34590 | -0.42 | 0.16 | 1.68 | bZIP11 |
| At1g77920 | -0.76 | -0.44 | n.d | TGA7 |
| At1g75390 | -0.49 | -0.55 | n.d | bZIP44 |
| At2g31370 | -0.30 | -0.24 | n.d | bZIP59 |
| At5g06960 | 0.38 | 0.57 | n.d | OBF5 |
| At3g10800 | 0.69 | 0.92 | n.d | bZIP28 |
| At2g40620 | 1.65 | 1.28 | n.d | bZIP18 |
| At2g41070 | 1.70 | 1.40 | -1.03 | bZIP12 |

$G_0$

| ID | iTPS 4h | iTPS 6h | oeTPS | Name |
|---|---|---|---|---|
| At3g30530 | -2.24 | -0.12 | n.d | bZIP42 |
| At2g17770 | -1.51 | -1.72 | n.d | bZIP27 |
| At2g36270 | -1.09 | -1.45 | n.d | ABI5 |
| At4g35900 | -0.99 | -1.45 | -1.14 | bZIP14 (FD) |
| At4g38900 | -0.39 | -0.24 | n.d | bZIP29 |
| At5g06950 | 0.43 | 0.30 | n.d | TGA2 |
| At3g56850 | 0.61 | 0.50 | n.d | DBPF3 |

#### C. WRKY

$G_1$

| ID | iTPS 4h | iTPS 6h | oeTPS | Name |
|---|---|---|---|---|
| At5g07100 | -2.93 | -2.96 | n.d | WRKY26 |
| At4g01720 | -2.75 | -2.00 | n.d | WRKY47 |
| At2g03340 | -1.59 | -1.14 | n.d | WRKY3 |
| At4g01250 | -1.28 | -0.99 | n.d | WRKY22 |
| At3g58710 | -0.75 | -1.03 | 1.08 | WRKY69 |
| At2g46400 | 0.65 | 0.70 | 2.04 | WRKY46 |
| At5g22570 | 1.80 | 1.83 | n.d | WRKY38 |

$G_2$

| ID | iTPS 4h | iTPS 6h | oeTPS | Name |
|---|---|---|---|---|
| At1g29860 | -2.82 | -0.62 | -1.39 | WRKY71 |
| At5g52830 | -2.59 | -2.02 | n.d | WRKY27 |
| At4g23810 | -1.35 | -0.50 | n.d | WRKY53 |
| At2g24570 | -1.15 | -0.76 | 1.16 | WRKY17 |
| At2g25000 | -0.73 | -0.53 | n.d | WRKY60 |
| At4g12020 | 0.34 | 0.59 | n.d | WRKY19 |
| At3g01970 | 0.58 | 0.79 | n.d | WRKY45 |
| At2g40750 | 0.62 | 0.72 | n.d | WRKY54 |
| At3g56400 | 1.36 | 1.29 | n.d | WRKY70 |
| At4g31800 | 1.41 | 0.88 | n.d | WRKY18 |

$G_0$

| ID | iTPS 4h | iTPS 6h | oeTPS | Name |
|---|---|---|---|---|
| At4g39410 | -3.46 | -2.54 | n.d | WRKY13 |
| At2g34830 | -2.51 | -1.77 | n.d | WRKY35 |
| At2g44745 | -2.21 | -1.99 | n.d | WRKY12 |
| At5g28650 | -1.90 | -2.12 | n.d | WRKY74 |
| At4g24240 | -1.38 | -1.18 | n.d | WRKY7 |
| At1g29280 | -1.35 | -2.52 | n.d | WRKY65 |
| At2g40740 | -0.89 | -0.35 | n.d | WRKY55 |
| At1g30650 | -0.86 | -0.71 | n.d | WRKY14 |
| At1g69310 | -0.81 | -0.44 | n.d | WRKY57 |
| At4g30935 | 0.66 | 0.78 | n.d | WRKY32 |
| At2g21900 | 1.59 | 0.72 | n.d | WRKY59 |
| At4g26440 | 6.02 | 4.97 | n.d | WRKY34 |

down
up

**Supplemental Figure S20. Transcription factors that respond to an induced increase in Tre6P levels Continued**

D. AP2/ERF

$G_1$

| ID | iTPS 4h | iTPS 6h | oeTPS | Name |
|---|---|---|---|---|
| At1g06160 | -4.45 | -3.57 | n.d | ERF59 |
| At5g61590 | -3.72 | -3.95 | -3.17 | DEWAX |
| At2g20880 | -3.54 | -1.92 | n.d | ERF53 |
| At3g54990 | -3.21 | -1.96 | -1.15 | SMZ |
| At1g79700 | -2.68 | -2.41 | -1.49 | WRI4 |
| At3g16770 | -2.53 | -2.60 | n.d | ERF72 |
| At4g28140 | -2.12 | -0.51 | n.d | ERF54 |
| At3g60490 | -1.85 | -1.30 | n.d | ERF35 |
| At1g46768 | -1.81 | -1.98 | n.d | RAP2.1 |
| At5g67190 | -1.68 | -1.54 | n.d | DEAR2 |
| At5g47220 | -1.56 | -1.27 | n.d | ERF2 |
| At4g39780 | -1.50 | -0.72 | n.d | ERF60 |
| At4g17490 | -1.24 | -1.05 | n.d | ERF6 |
| At1g72360 | -1.12 | -0.71 | n.d | ERF73 |
| At1g36060 | -1.06 | -0.21 | n.d | WIND3 |
| At5g61600 | -0.99 | -1.43 | n.d | ERF104 |
| At1g22190 | -0.97 | -0.63 | n.d | WIND2 |
| At4g17500 | -0.85 | -0.38 | n.d | ERF1 |
| At3g50260 | -0.68 | -0.48 | n.d | DEAR1 |
| At1g53910 | -0.56 | -0.58 | n.d | ERF74 |
| At1g50640 | -0.53 | -0.41 | n.d | ERF3 |
| At1g25470 | -0.34 | -0.39 | n.d | CRF12 |
| At5g22770 | 0.34 | 0.54 | n.d | AP2A1 |
| At2g31230 | 0.40 | 0.62 | n.d | ERF15 |
| At2g23340 | 0.42 | 0.19 | n.d | DEAR3 |
| At4g23750 | 0.80 | 0.63 | n.d | CRF2 |

$G_2$

| ID | 29.2 4h | 29.2 6h | oeOTSA | name |
|---|---|---|---|---|
| At5g07580 | -2.81 | -1.68 | -1.26 | DEWAX2 |
| At2g39250 | -2.55 | -1.97 | n.d | SNZ |
| At5g44210 | -1.81 | -1.50 | n.d | ERF9 |
| At4g37750 | -1.56 | -1.69 | n.d | DRG |
| At4g32800 | -1.55 | -1.27 | n.d | ERF43 |
| At4g36920 | -1.43 | -1.07 | n.d | FLO2 |
| At5g53290 | -0.68 | -0.52 | 1.66 | CRF3 |
| At1g16060 | -0.61 | -1.14 | n.d | WRI3 |
| At5g60120 | -0.60 | -0.53 | n.d | EAT2 |
| At1g68550 | -0.56 | -0.31 | 1.19 | CRF10 |
| At2g44940 | -0.53 | -0.31 | n.d | ERF34 |
| At2g28550 | -0.41 | -0.30 | n.d | EAT1 |
| At4g36900 | 1.14 | 1.21 | n.d | DEAR4 |
| At5g13330 | 1.18 | 2.43 | n.d | related to AP2 6l |
| At2g38340 | 1.31 | 1.00 | 3.86 | DREB19 |
| At3g11020 | 1.94 | 0.73 | n.d | DREB2 |
| At1g43160 | 2.92 | 4.33 | n.d | related to AP2 6 |

$G_0$

| ID | 29.2 4h | 29.2 6h | oeOTSA | name |
|---|---|---|---|---|
| At4g25490 | -3.28 | -2.37 | n.d | CBF1 |
| At5g25390 | -3.25 | -2.79 | n.d | SHN3 |
| At5g25190 | -3.19 | -2.65 | -1.49 | ESE3 |
| At1g33760 | -3.18 | -2.27 | n.d | ERF022 |
| At4g25470 | -2.60 | -2.96 | n.d | CBF2 |
| At2g22200 | -2.42 | -2.33 | n.d | T26C19.14 |
| At5g57390 | -2.33 | -1.81 | n.d | AIL5 |
| At1g77200 | -2.11 | -0.81 | n.d | T14N5.6 |
| At4g11140 | -1.79 | -2.21 | n.d | CRF1 |
| At4g25480 | -1.75 | -0.95 | n.d | DREB1A |
| At4g16750 | -1.54 | -1.68 | n.d | ERF39 |
| At1g78080 | -1.32 | -0.62 | n.d | WIND1 |
| At5g47230 | -1.11 | -0.42 | 1.16 | ERF5 |
| At5g65510 | -1.11 | -1.51 | -2.58 | AIL7 |
| At3g54320 | -0.98 | -0.85 | 1.17 | WRI1 |
| At3g23230 | -0.94 | -0.77 | n.d | TDR1 |
| At5g67180 | -0.75 | -1.19 | -1.41 | TOE3 |
| At3g59990 | 0.35 | 0.40 | n.d | MAP2B |
| At3g15210 | 0.99 | 1.04 | n.d | ERF4 |
| At1g74930 | 1.76 | 1.17 | -1.69 | ORA47 |
| At5g25810 | 3.04 | 2.72 | n.d | TNY |
| At1g71450 | 3.23 | 3.33 | n.d | FUF1 |
| At2g47520 | 3.69 | 3.64 | n.d | ERF71 |
| At1g12630 | 4.51 | 4.53 | n.d | T12C24.16 |
| At5g52020 | 4.95 | 4.78 | n.d | MSG15.10 |
| At1g72570 | 5.95 | 5.86 | n.d | F28P22.24 |
| At5g67000 | 6.20 | 3.54 | n.d | K8A10.7 |
| At1g12610 | 8.86 | 6.79 | n.d | DDF1 |
| At1g63030 | 9.22 | 8.64 | n.d | DDF2 |

down
up

**Supplemental Figure S20. Transcription factors that respond to an induced increase in Tre6P levels Continued.**

#### E. MYBs

### $G_1$

| ID | iTPS 4h | iTPS 6h | oeTPS | Name |
|---|---|---|---|---|
| At5g59780 | -4.33 | -3.52 | n.d | MYB59 |
| At3g46130 | -4.18 | -3.33 | -1.40 | ATMYB48 |
| At5g53200 | -3.51 | -2.35 | n.d | TRY |
| At1g66230 | -2.63 | -2.40 | -2.66 | MYB20 |
| At2g30420 | -2.37 | -1.98 | n.d | ETC2 |
| At1g22640 | -2.35 | -2.55 | n.d | ATMYB3 |
| At5g57620 | -2.31 | -2.65 | n.d | MYB36 |
| At4g37260 | -2.28 | -2.68 | n.d | MYB73 |
| At1g71030 | -2.15 | -1.53 | -1.36 | ATMYBL2 |
| At1g19510 | -2.13 | -3.18 | n.d | ATRL5 |
| At1g25550 | -2.12 | -1.98 | n.d | HHO3 |
| At5g44190 | -2.00 | -1.29 | n.d | GLK2 |
| At4g09460 | -1.94 | -1.85 | -2.23 | AtMYB6 |
| At2g23290 | -1.75 | -1.83 | n.d | AtMYB70 |
| At5g42630 | -1.72 | -2.07 | n.d | KAN4 |
| At2g47190 | -1.58 | -2.11 | n.d | ATMYB2 |
| At5g37260 | -1.58 | -1.69 | -2.92 | CIR1 |
| At5g15310 | -1.54 | -1.41 | n.d | ATMYB16 |
| At1g70000 | -1.47 | -0.42 | n.d | MYBD |
| At5g08520 | -1.46 | -1.18 | n.d | MYBS2 |
| At1g75250 | -1.41 | -1.91 | n.d | ATRL6 |
| At1g79430 | -1.24 | -1.38 | n.d | APL |
| At3g06490 | -1.23 | -3.04 | n.d | MYB108 |
| At1g49010 | -1.21 | -1.17 | n.d | MYBS1 |
| At1g18330 | -1.17 | -1.31 | -3.10 | EPR1 |
| At3g47600 | -1.10 | -0.83 | n.d | MYB94 |
| At5g17300 | -1.09 | -0.39 | -2.74 | RVE1 |
| At5g06800 | -1.06 | -1.76 | n.d | MPH15.16 |
| At5g59430 | -1.04 | -0.52 | n.d | ATTRP1 |
| At2g46410 | -1.03 | -0.54 | n.d | CPC |
| At1g19000 | -1.02 | -0.89 | -1.08 | F14D16.15 |
| At1g74840 | -0.99 | -1.13 | -1.01 | F25A4.19 |
| At3g12730 | -0.98 | -1.63 | n.d | MBK21.11 |
| At1g56650 | 0.72 | 1.54 | n.d | PAP1 |
| At3g55730 | 1.52 | 1.56 | n.d | MYB109 |
| At1g17460 | 1.69 | 1.44 | n.d | TRFL3 |

down
up

### $G_2$

| ID | iTPS 4h | iTPS 6h | oeTPS | Name |
|---|---|---|---|---|
| At5g60890 | -4.33 | -3.52 | n.d | MYB34 |
| At2g38090 | -3.11 | -2.74 | n.d | F16M14.2 |
| At2g16720 | -2.92 | -2.24 | n.d | MYB7 |
| At5g07690 | -2.86 | -3.05 | n.d | MYB29 |
| At1g13300 | -2.82 | -2.64 | n.d | HRS1 |
| At5g65790 | -2.61 | -1.99 | n.d | MYB68 |
| At1g79180 | -2.60 | -1.16 | n.d | MYB63 |
| At5g05790 | -2.59 | -2.38 | n.d | MJJ3.20 |
| At4g38620 | -2.57 | -1.98 | n.d | MYB4 |
| At5g16600 | -2.39 | -1.82 | n.d | MYB43 |
| At2g36890 | -2.36 | -0.73 | n.d | MYB38 |
| At5g58900 | -2.17 | -1.90 | 1.58 | DIV1 |
| At3g48920 | -2.13 | -0.69 | n.d | MYB45 |
| At2g02820 | -1.93 | -1.85 | n.d | MYB88 |
| At4g17695 | -1.81 | -0.99 | n.d | KAN3 |
| At5g61420 | -1.74 | -1.86 | n.d | MYB28 |
| At4g12350 | -1.65 | -1.42 | n.d | MYB42 |
| At2g03500 | -1.51 | -1.38 | n.d | EFM |
| At1g14350 | -1.49 | -1.63 | n.d | MYB124 |
| At2g37630 | -1.43 | -1.27 | -1.36 | MYB91 |
| At4g22680 | -1.14 | -0.70 | n.d | MYB85 |
| At5g49330 | -0.98 | -0.26 | 2.06 | MYB111 |
| At1g58220 | 0.93 | 1.09 | n.d | DRMY1 |
| At4g01280 | 1.06 | 1.06 | n.d | RVE5 |
| At1g01520 | 1.12 | 1.65 | n.d | REV3 |

### $G_0$

| ID | iTPS 4h | iTPS 6h | oeTPS | Name |
|---|---|---|---|---|
| At1g18710 | -2.84 | -2.11 | n.d | MYB47 |
| At1g74080 | -2.60 | -0.85 | n.d | MYB122 |
| At3g12820 | -2.16 | -3.15 | n.d | MYB10 |
| At1g63910 | -2.08 | -1.30 | n.d | MYB103 |
| At3g50060 | -2.05 | -1.69 | n.d | MYB77 |
| At1g66390 | -2.04 | -1.62 | 1.68 | MYB90 |
| At5g07700 | -2.03 | -2.03 | n.d | MYB76 |
| At1g66370 | -1.98 | -1.08 | n.d | MYB113 |
| At1g18570 | -1.97 | -1.16 | 2.01 | MYB51 |
| At5g10280 | -1.73 | -2.05 | n.d | MYB92 |
| At3g28910 | -1.68 | -1.18 | n.d | MYB30 |
| At5g67300 | -1.65 | -1.46 | n.d | MYBR1 |
| At3g60110 | -1.62 | -1.66 | n.d | T2O9.90 |
| At4g01680 | -1.60 | -1.66 | n.d | MYB55 |
| At2g02060 | -1.52 | -1.52 | n.d | F14H20.13 |
| At5g23000 | -1.51 | -2.16 | n.d | MYB37 |
| At5g14750 | -1.49 | -1.45 | n.d | MYB66 |
| At1g06180 | -1.48 | -0.94 | n.d | MYB13 |
| At2g40260 | -1.35 | -1.15 | n.d | T7M7.13 |
| At1g66380 | -1.28 | -1.54 | n.d | MYB114 |
| At2g47460 | -1.26 | -2.29 | n.d | MYB12 |
| At3g04030 | -1.13 | -1.27 | n.d | MYR2 |
| At5g58340 | 0.83 | 1.04 | n.d | MCK7.21 |
| At5g40350 | 1.69 | 0.35 | n.d | MYB24 |
| At3g28470 | 1.74 | 2.63 | n.d | MYB35 |
| At5g16770 | 2.39 | 0.94 | n.d | AtMYB9 |
| At2g42150 | 3.65 | 3.46 | n.d | T24P15.6 |
| At3g02940 | 4.91 | 2.04 | n.d | MYB107 |
| At5g55020 | 5.94 | 4.01 | n.d | MYB120 |
| At5g61620 | 7.91 | 6.86 | n.d | K11J9.15 |

**Supplemental Figure S21. Analysis of transcription factors assigned to CRF group $G_1$.**

(A Enrichment analysis using Gene Ontology terms. Blue denotes fold enrichment and orange the FDR value. The nine categories at the bottom of the display were underrepresented.

(B) STRING analysis (see next slide)

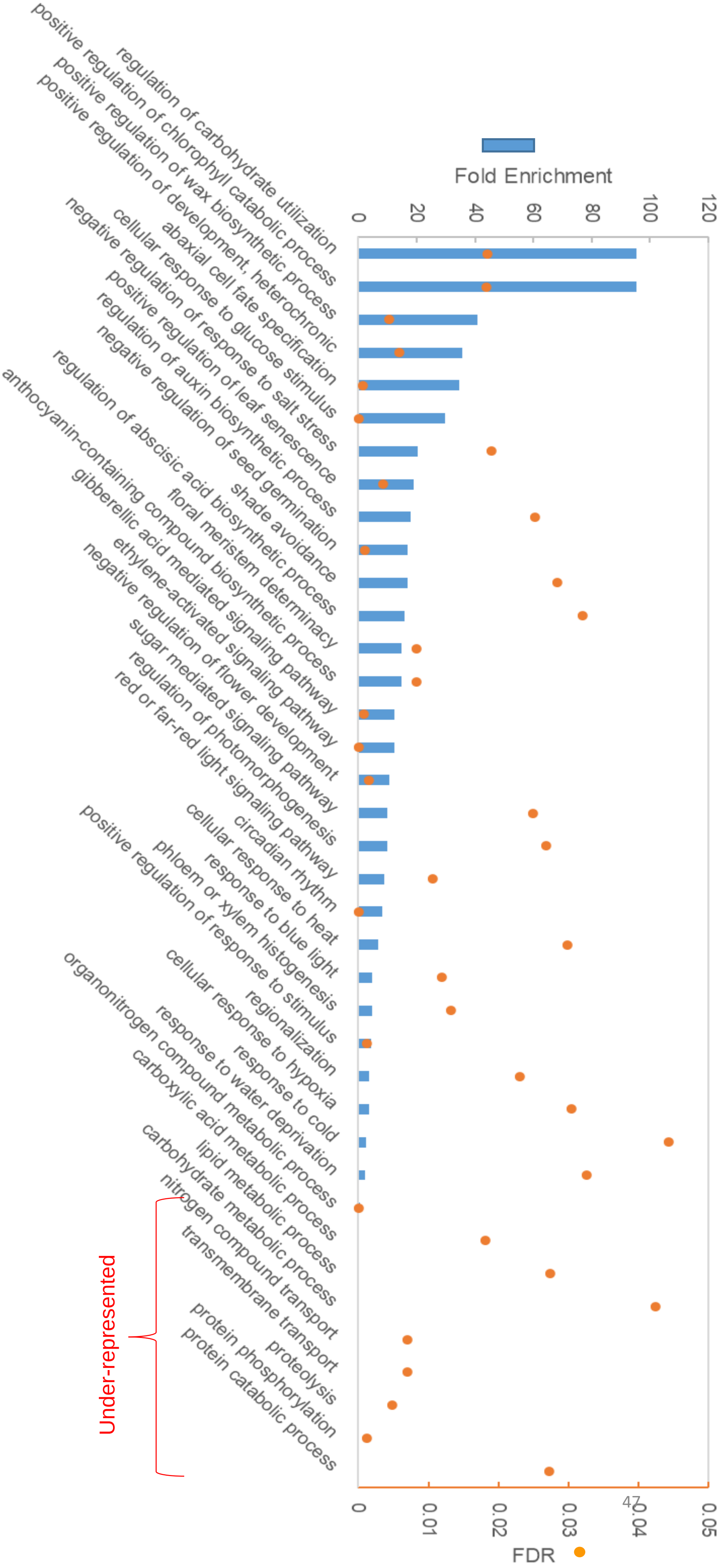

**Supplemental Figure S21. Analysis of transcription factors assigned to Crf group $G_1$ (continued) .**

(B) STRING analysis. The tool utilizes available datasets, published work, automated text mining and computational predictions from various organisms to score the likelihood of association between proteins, taking into account physical (direct) and functional (indirect) interactions (Szklarczyk et al., 2021). We used the more relaxed filter for expression (FDR < 0.05 and log2FC ≥ 0.2; 290 TFs in total) and set a high interaction confidence score filter (interaction score ≥ 0.7). The thickness of the line connecting two nodes indicates the strength of data support, with thicker lines representing higher interaction score

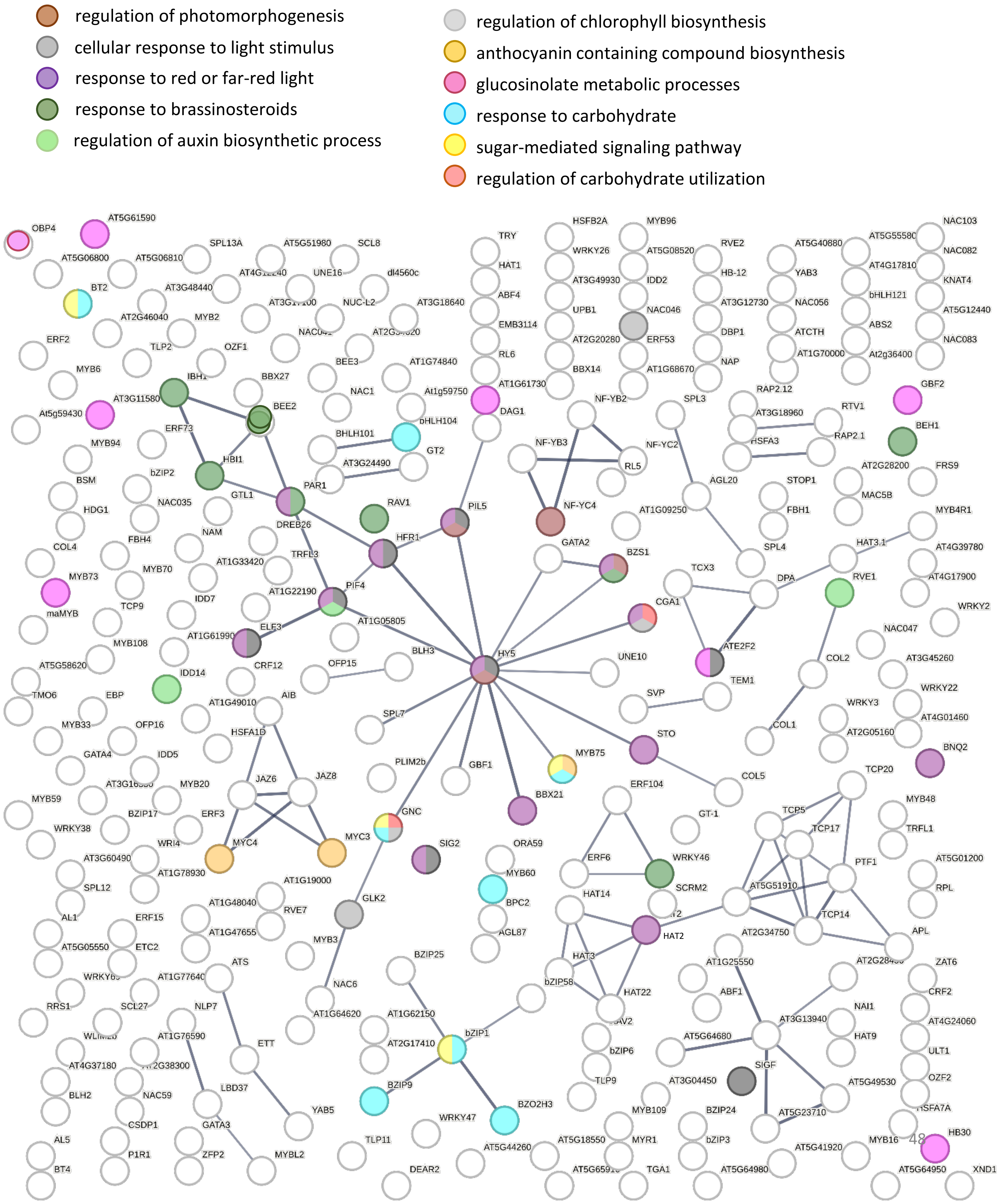

**Supplemental Figure S22. C and $S_1$ bZIP transcription factors**

(A) (A) Response of members of the C and $S_1$ bZIP families. The displayshows 4-h and 6-h iTPS response, assignment of the iTPS response to CRF groups, the CRF value (defined as in Supplemental Figure S2) and the response to constitutive overexpression of TPS (oeTPS, from Zhang et al., 2009) and transient overexpression of SnRL1α1 in protoplasts (tSnRL1α1 response, from Baena-Gonzalez et al., 2007). Significant changes in the iTPS response are shown with bold font. Assignment to CRF groups is performed only for significant responses and only CRF $G_1$ assignments are shaded. No entry for oeTPS or tSnRL1α1 indicates that the response did not pass the filter in the original publication.

(B) Relationship between the iTPS response and the tSnRK1α1 response, shown for transcripts assigned to CRF group $G_1$, $G_2$ and $G_0$ (from left to right). The display shows the response of transcript abundance4-h and 6-h post-induction, assignment of the iTPS response to CRF groups, the CRF value (defined as in Supplemental Figure S2) and the response of transcript abundance to constitutive overexpression of TPS (oeTPS, from Zhang et al., 2009) and transient overexpression of SnRL1α1 in protoplasts (tSnRL1α1,from Baena-Gonzalez et al., 2007). Significant changes in the iTPS response are shown with bold font. Assignment to CRF groups is performed only for significant responses and only CRF $G_1$ assignments are shaded. No entry for oeTPS or tSnRL1α1 indicates that the response did not pass the filter in the publication.

A

| **Gene** | | **iTPS response** | | | | | | **CRF** | **Other responses** | | |
|---|---|---|---|---|---|---|---|---|---|---|---|
| Name | Code | 4-h adj.p | 6-h adj.p | 4-h FC $log_2$ | 6-h FC $log_2$ | 4-h CRF score | 6-h CRF score | | oeTPS FC $log_2$ | tSnRK1α1 FC $Log_2$ | tSnRK1α1 compared to iTPS |
| bZIP9 | At5g24800 | **0.00** | **0.00** | **-1.18** | **-0.67** | **1** | **1** | -1.54 | | -1.32 | Same |
| bZIP10 | At4g02640 | 0.80 | 0.70 | -0.03 | 0.06 | | | 0.01 | | -0.60 | No iTPS response |
| bZIP25 | At3g54620 | **0.00** | **0.00** | **-0.59** | **-0.41** | **1** | **1** | -0.21 | | 0.38 | **Reciprocal** |
| bZIP63 | At5g28770 | **0.00** | **0.00** | **-1.53** | **-1.65** | **1** | **1** | -1.68 | 1.35 | 2.87 | **Reciprocal** |
| bZIP1 | At5g49450 | **0.00** | **0.00** | **-1.55** | **-1.88** | **1** | **1** | -2.64 | 1.46 | 1.57 | **Reciprocal** |
| bZIP2 | At2g18160 | **0.00** | **0.00** | **-1.49** | **-1.45** | **1** | **1** | -0.49 | | | Same |
| bZIP11 | At4g34590 | **0.00** | 0.27 | **-0.42** | 0.16 | 2 | | 1.16 | 1.68 | -3.11 | Inconsistent |
| bZIP44 | At1g75390 | **0.03** | **0.02** | **-0.49** | **-0.55** | 2 | 2 | 0.34 | | -0.03 | No oe SnRK1 response |
| bZIP53 | At3g62420 | 0.52 | 0.95 | 0.07 | 0.01 | | | -0.58 | | 0.44 | No iTPS response |

B

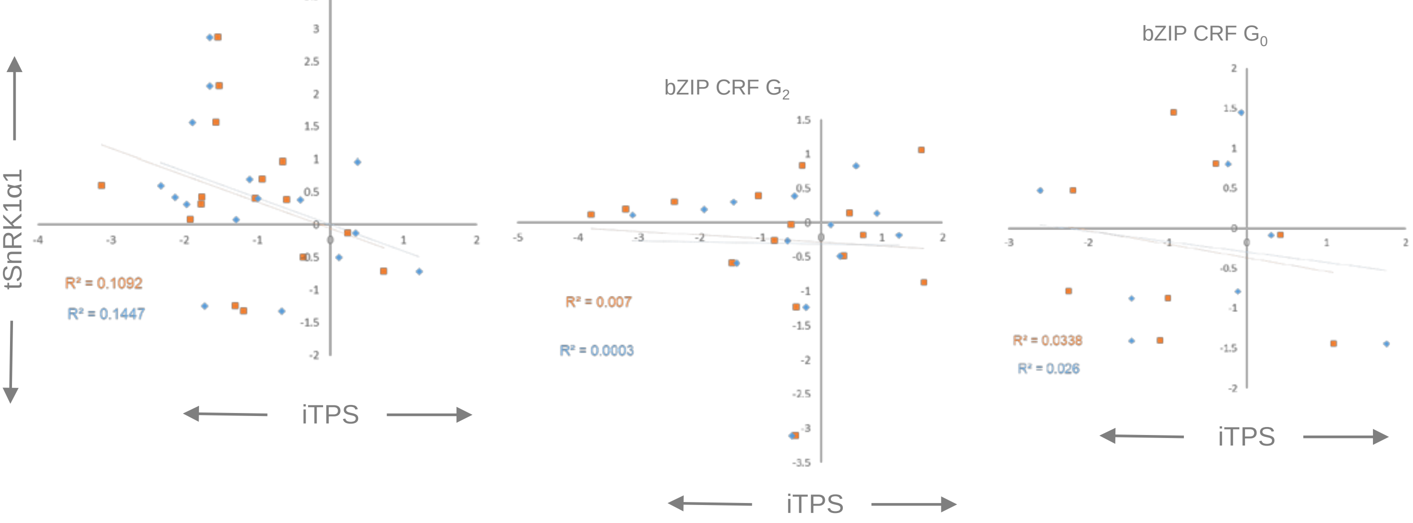

iTPS on x-axis, tSnRK1α1 on y-axis

**Supplemental Figure S23: TFs that were assigned to Crf group G1 in the iTPS response: comparison of their iTPS response and in their tSnRK1α1 response.** Responses are shown for genes in different TF families (see displays) focusing on TF families with at least 5 TFs that showed a $G_1$ type response.

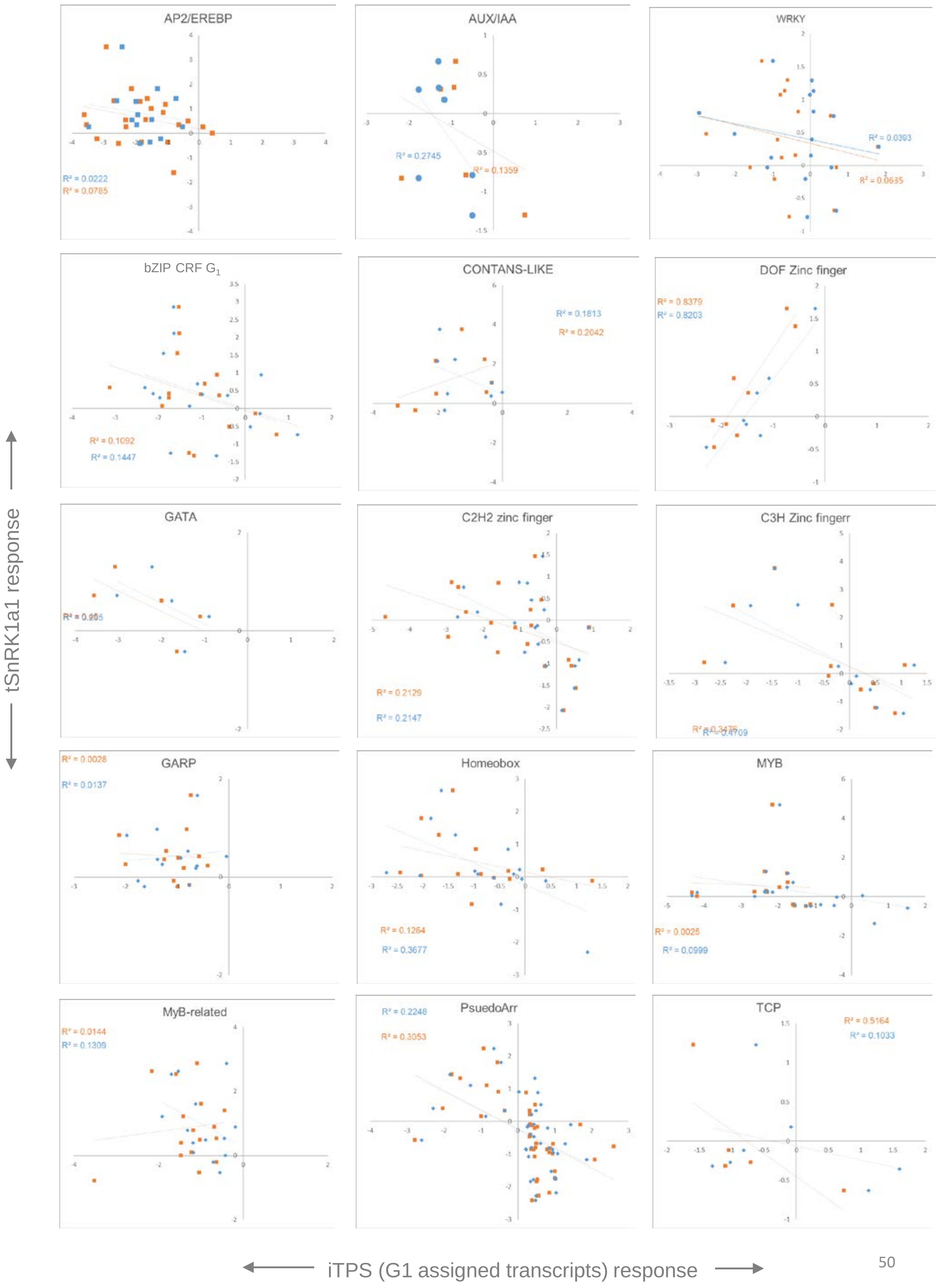

**Supplemental Figure S24. Analysis of transcription factors assigned to CRF group $G_2$.**

(A) Enrichment analysis using Gene Ontology terms. . Blue denotes fold enrichment and orange the FDR value. The nine categories at the bottom of the display were underrepresented.

(B) STRING analysis, see next slide

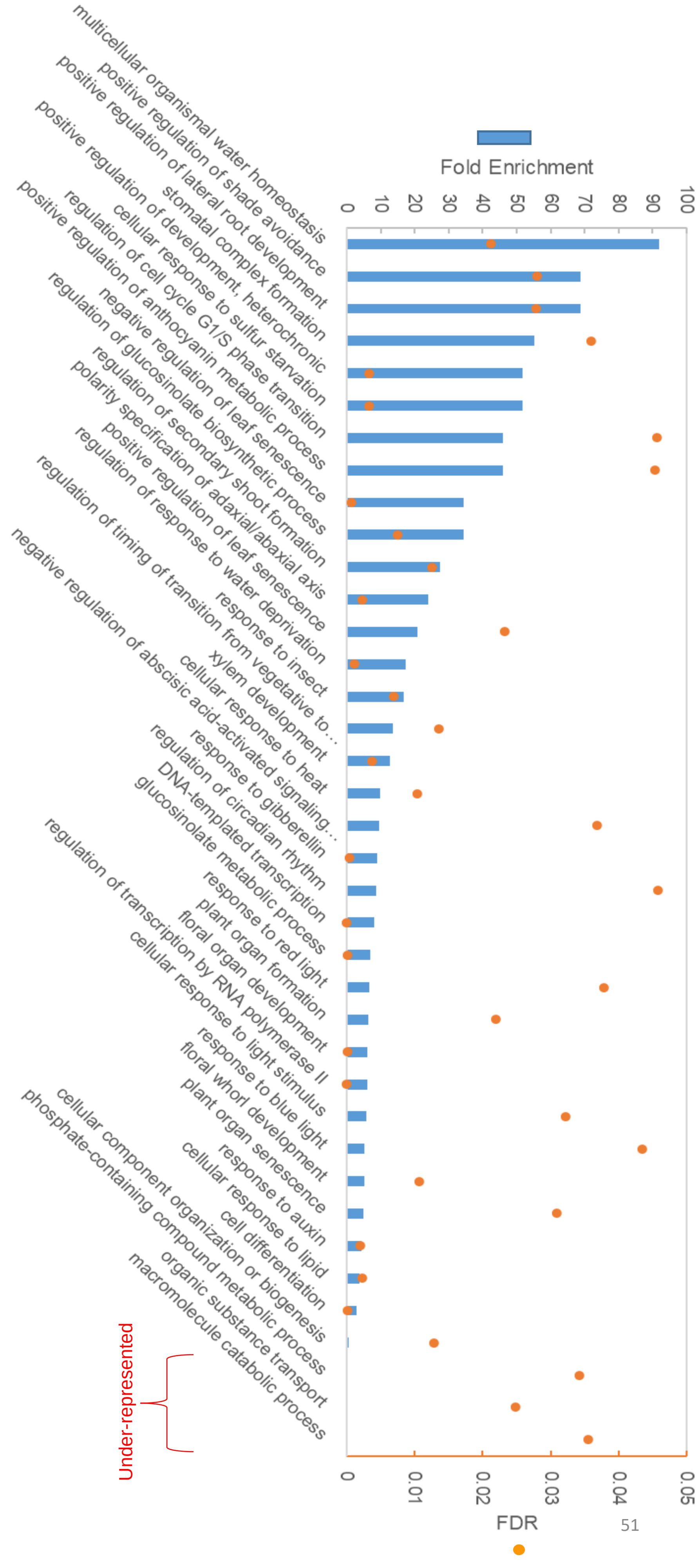

**Supplemental Figure S24. Analysis of transcription factors assigned to Crf group $G_2$ (continued)**

(B) STRING analysis. The tool utilizes available datasets, published work, automated text mining and computational predictions from various organisms to score the likelihood of association between proteins, taking into account physical (direct) and functional (indirect) interactions (Szklarczyk et al., 2021). We used the more relaxed filter for expression (FDR < 0.05 and log2FC ≥ 0.2; 780 TFs in total) and set a high interaction confidence score filter (interaction score ≥ 0.7). The thickness of the line connecting two nodes indicates the strength of data support, with thicker lines representing higher interaction score

- Response to nitrogen compound
- positive regulation of nitrogen compound metabolic processes
- Cellular response to sulfur starvation
- regulation of secondary metabolite metabolic processes
- secondary metabolic processes
- regulation of anthocyanin metabolic processes
- Positive regulation of anthocyanin metabolic process
- Regulation of flavonoid biosynthetic process
- Regulation of glucosinolate biosynthetic process

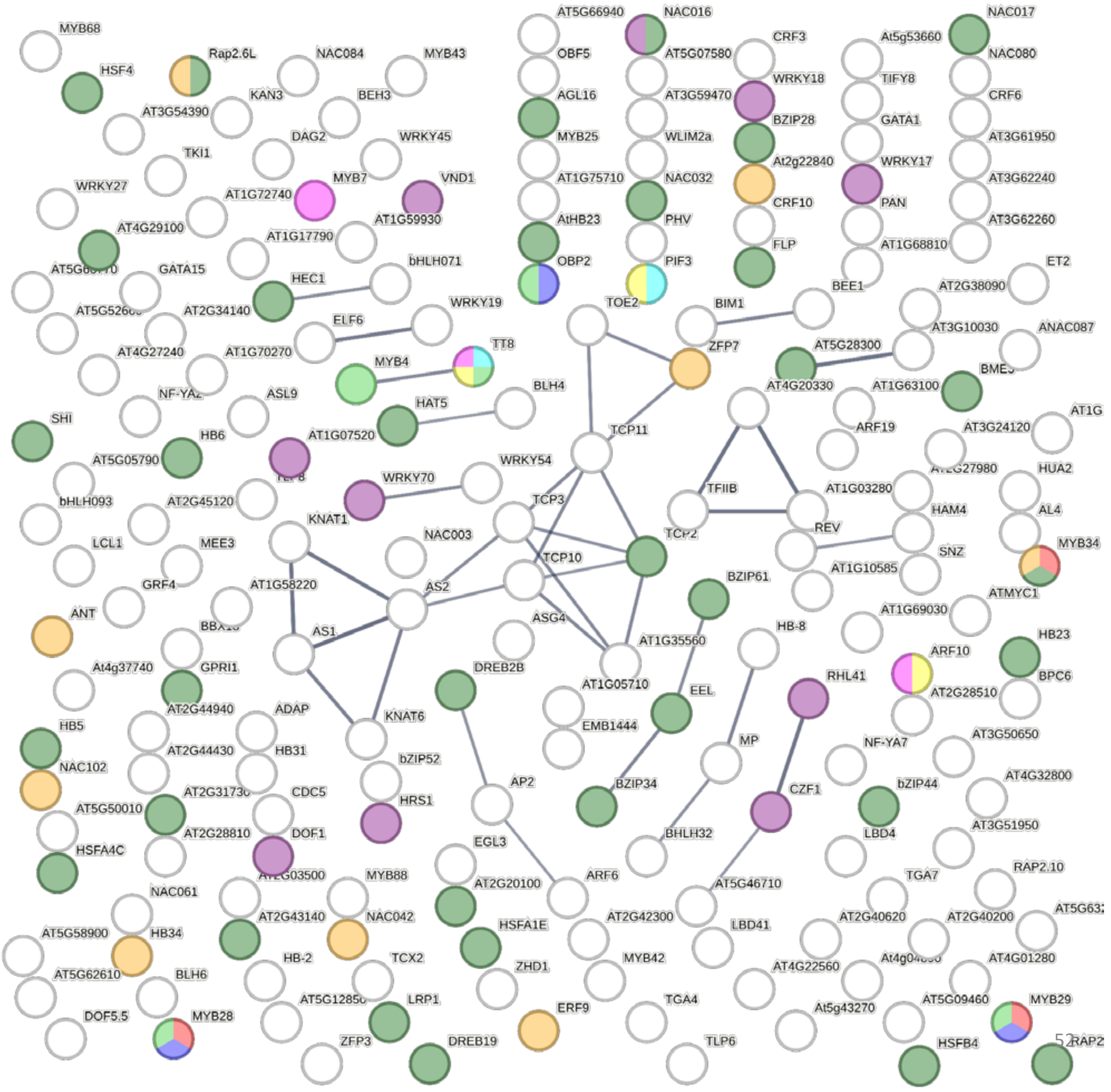

#### Supplemental Figure S25 Glucosinolate metabolism and its transcriptional regulation

(A) Genes involved in glucosinolate biosynthesis

(B) Transcription factors involved in the regulation of glucosinolate biosynthesis

The displays show the4-h and 6-h iTPS response, assignment of the iTPS response to CRF groups, the CRF value (defined as in Supplemental Figure S2) and the response to constitutive overexpression of TPS (oeTPS, from Zhang et al., 2009) and transient overexpression oftSnRL1α1 in protoplasts (tSnRL1α1, from Baena-Gonzalez et al., 2007). Significant changes in the iTPS response are shown with bold font. Assignment to CRF groups is performed only for significant responses and only CRF $G_1$ assignments are shaded. No entry for oeTPS or tSnRL1α1 indicates that the response did not pass the filter in the original publication.

A

| Gene | iTPS response | | | | | | CRF | Other responses | | Annotation |
|---|---|---|---|---|---|---|---|---|---|---|
| Code | 4-h adj.p | 6-h adj.p | **4-h FC $log_2$** | **6-h FC $log_2$** | 4-h CRF score | 6-h CRF score | | oeTPS FC $log_2$ | tSnRK1α1 FC $Log_2$ | |
| At4g39950 | 0,00 | 0,00 | **-2,98** | **-3,02** | 2 | 2 | 1,41 | 2,57 | -1,59 | CYP79B2 (cytochrome P450, family 79, subfamily B, polypeptide 2); oxygen binding |
| At5g57220 | 0,00 | 0,00 | **-2,69** | **-1,72** | 0 | 0 | -0,02 | 2,32 | 0,26 | CYP81F2 (cytochrome P450, family 81, subfamily F, polypeptide 2); oxygen binding |
| At2g22330 | 0,00 | 0,00 | **-2,59** | **-2,78** | 2 | 2 | 0,50 | | 0,85 | CYP79B3 (cytochrome P450, family 79, subfamily B, polypeptide 3); oxygen binding |
| At1g65880 | 0,11 | 0,06 | **-2,02** | **-4,55** | | | 0,05 | | -1,53 | AMP-dependent synthetase and ligase family protein |
| At1g65860 | 0,00 | 0,00 | **-1,91** | **-2,35** | 2 | 2 | 1,32 | | -1,06 | flavin-containing monooxygenase family protein / FMO family protein |
| At1g18590 | 0,00 | 0,00 | **-1,79** | **-2,53** | 2 | 2 | 1,51 | | -0,82 | sulfotransferase family protein |
| At4g03050 | 0,56 | 0,55 | **-1,72** | **-1,76** | | | 0,00 | | -0,99 | AOP3 (2-oxoglutarate?dependent dioxygenase 3); oxidoreductase, a |
| At5g23020 | 0,00 | 0,00 | **-1,58** | **-1,66** | 2 | 2 | 1,37 | | -0,17 | MAM-L (METHYLTHIOALKYMALATE SYNTHASE-LIKE); 2-isopropylmalate synthase |
| At1g62560 | 0,00 | 0,00 | **-1,40** | **-2,02** | 2 | 2 | 1,65 | | -0,59 | flavin-containing monooxygenase family protein / FMO family protein |
| At5g23010 | 0,00 | 0,00 | **-1,32** | **-1,92** | 2 | 2 | 0,48 | | 0,70 | MAM1 (2-isopropylmalate synthase 3); 2-isopropylmalate synthase |
| At2g20610 | 0,00 | 0,00 | **-1,26** | **-1,43** | 2 | 2 | 0,79 | 1,85 | -2,01 | UR1 (SUPERROOT 1); transaminase |
| At4g13770 | 0,00 | 0,00 | **-1,23** | **-1,63** | 2 | 2 | 0,83 | 0,00 | 0,57 | CYP83A1 (CYTOCHROME P450 83A1); oxygen binding |
| At1g74100 | 0,00 | 0,00 | **-1,00** | **-1,03** | 2 | 2 | 0,63 | 2,03 | -1,06 | sulfotransferase family protein |
| At1g74090 | 0,00 | 0,00 | **-0,96** | **-1,65** | 2 | 2 | 0,60 | | -0,32 | sulfotransferase family protein |
| At2g43100 | 0,00 | 0,00 | **-0,55** | **-0,91** | 2 | 2 | 1,35 | | -0,71 | aconitase C-terminal domain-containing protein |
| At3g58990 | 0,09 | 0,00 | **-0,17** | **-1,01** | | 2 | 1,56 | 1,41 | -0,08 | aconitase C-terminal domain-containing protein |
| At4g31500 | 0,00 | 0,00 | **-0,14** | **-0,86** | 2 | 2 | 0,66 | 2,28 | -1,69 | CYP83B1 (CYTOCHROME P450 MONOOXYGENASE 83B1); oxygen binding |
| At4g13430 | 0,37 | 0,36 | **-0,09** | **-0,15** | | | 0,39 | | -0,25 | aconitase family protein / aconitate hydratase family protein |
| At4g12030 | 0,00 | 0,00 | **-0,07** | **-0,82** | 2 | 2 | 1,07 | | 0,07 | ile acid:sodium symporter family protein |
| At1g80560 | 0,63 | 0,28 | **0,05** | **-0,16** | | | 0,29 | | -1,30 | 3-isopropylmalate dehydrogenase, chloroplast, putative |

B

| Gene | | iTPS response | | | | | | CRF | Other responses | | Annotation |
|---|---|---|---|---|---|---|---|---|---|---|---|
| Name | Code | 4-h adj.p | 6-h adj.p | **4-h FC $log_2$** | **6-h FC $log_2$** | 4-h CRF score | 6-h CRF score | | oeTPS FC $log_2$ | tSnRK1α1 FC $Log_2$ | |
| MYB29 | At5g07690 | 0,00 | 0,00 | **-2,86** | **-3,05** | 2 | 2 | 0,97 | | -0,30 | myb domain protein 29); regulates aliphatic GLS biosynthesis |
| MYB28 | At5g61420 | 0,00 | 0,00 | **-1,74** | **-1,86** | 2 | 2 | 1,00 | | -1,31 | myb domain protein 28; regulates aliphatic GLS biosynthesis |
| MYB76 | At5g07700 | 0,00 | 0,00 | **-2,03** | **-2,03** | 0 | 0 | -0,05 | | -0,39 | myb domain protein 76; regulates aliphatic GLS biosynthesis |
| MYB34 | At5g60890 | 0,00 | 0,00 | **-4,33** | **-3,52** | 2 | 2 | 1,15 | | -0,16 | ATR1/MYB34 (ALTERED TRYPTOPHAN REGULATION); regulates indolic GLS synthesis |
| MYB122 | At1g74080 | 0,00 | 0,20 | **-2,60** | **-0,85** | 0 | | 0,01 | | -1,65 | myb domain protein 122); regulates indolic GLS synthesis |
| MYB51 | At1g18570 | 0,00 | 0,00 | **-1,97** | **-1,16** | 0 | 0 | -0,10 | 2,01 | -1,63 | myb domain protein 51; regulates indolic GLS synthesis |
| MYC2 | At1g32640 | 0,00 | 0,00 | **-1,57** | **-1,82** | 1 | 1 | -0,56 | | -0,26 | ATMYC2 (JASMONATE INSENSITIVE 1); DNA binding / transcription factor |
| MYC3 | At5g46760 | 0,00 | 0,04 | **-0,66** | **-0,30** | 1 | 1 | -0,39 | | 1,34 | JAZ-interacting transcription factor |
| MYC4 | At4g17880 | 0,00 | 0,00 | **0,64** | **0,75** | 1 | 1 | 0,32 | | | JAZ-interacting transcription factor thatwith MYC2 and MYC3 activatse JA-responses. |
| MYC5 | At5g46830 | 0,02 | 0,24 | **-0,93** | **-0,44** | 0 | | 0,05 | | -0,89 | (at5g46830):basic helix-loop-helix (bHLH) family protein |
| OBP2 | At1g07640 | 0,00 | 0,00 | **-1,00** | **-1,01** | 2 | 2 | 0,23 | | 0,64 | OBF BINDING PROTEIN 2; upstream regulator of indolic GLS |
| SR1 | At2g22300 | 0,23 | 0,94 | **-0,16** | **0,02** | | | -0,54 | | 0,56 | (at2g22300):ethylene-responsive calmodulin-binding protein, putative (SR1) |
| WRKY33 | At2g38470 | 0,37 | 0,00 | **-0,16** | **0,63** | | 2 | -0,56 | 1,48 | -0,12 | WRKY DNA-binding protein 33; directly regulates indolic GSL biosynthesis |

**Supplemental Figure S26. Proposed interplay between Tre6P-mediated signaling and other C-signaling pathways in wild-type plants.**

The proposed scheme is derived from Figures 5 and 7. It is based on difference between the observed overall response to induction of TPS, in which Tre6P rises but sucrose and many related other metabolites decline, and the known response in wild-type plants to increased C availability, in which Tre6P, sucrose and related metabolites rise (see Figure 1, Supplemental Figure S4, also Martins et al., 2013 and Figueroa et al., 2016, Avidan et al., 2023). For transcripts assigned to CFR group $G_1$, the response to rising C availability in wild-type plants is predicted to qualitatively similar to their iTPS response. in contrast, for transcripts assigned to CFR group $G_2$, the response to rising C availability in wild-type plants is predicted to be qualitatively opposite to their iTPS response (and similar to their CRF). Analyses summarized in Figures 5 and 7 identified processes where there was a marked bias for the underlying genes to be assigned to either CRF group $G_1$ or group $G_2$. The projected scenarios were generated by reversing the direction of change for processes that were linked to CRF group $G_2$ and whose observed iTPS response is therefore likely to be driven by the decease of sucrose and related metabolites.

**A. Responses of metabolism and growth- and defense-related processes to a rising C supply in wild-type plants., and the contribution of Tre6P and other C-signaling pathways to this response**

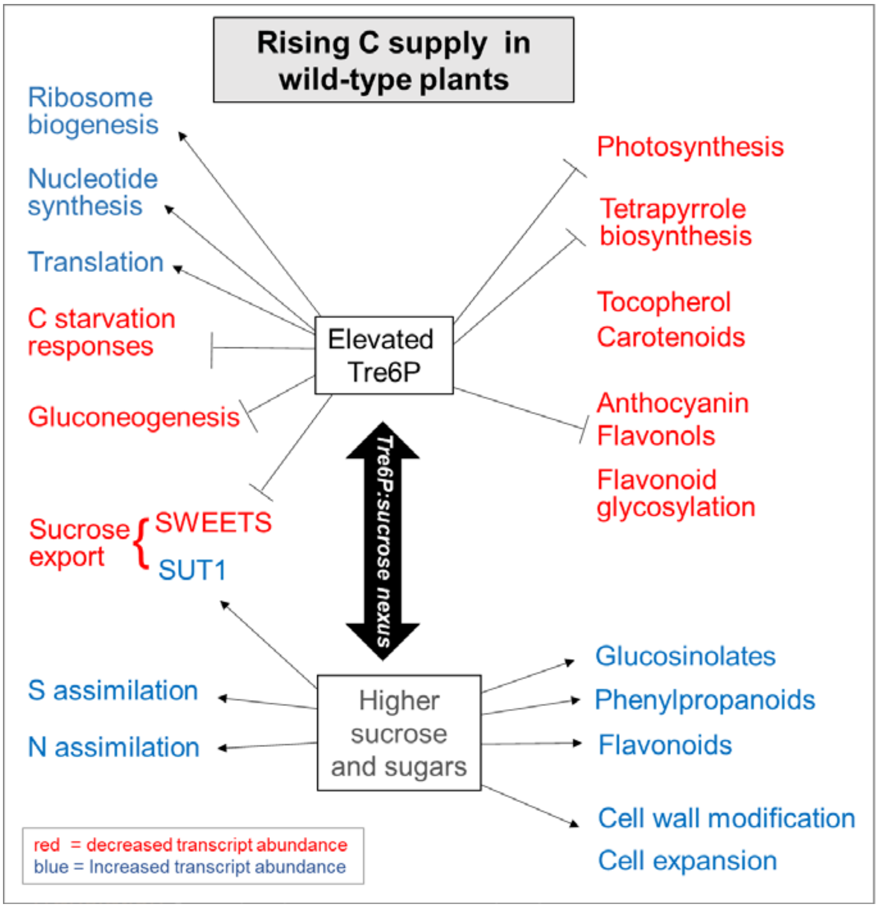

**B. Responses of genes involved in Tre6P metabolism and signaling and in the SnRK1 complex.** Genes assigned to CRF group $G_0$ are unlikely to respond to rising C availability, and if they do, not in a predicable way, and are colored black

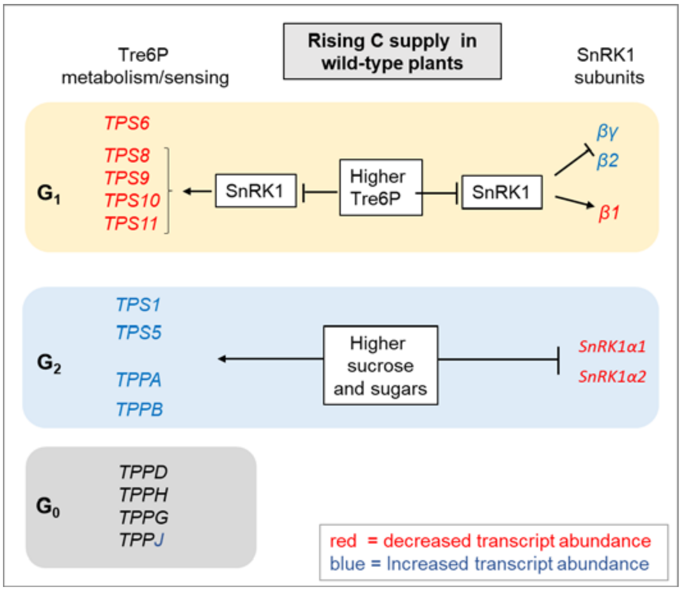

**Supplemental Table S1. List of transcriptome data sets.** The Table summarizes microarray and RNA-sequencing (RNAseq) datasets used in this study, with a description of treatments and transgenic lines with number of differentially expressed genes (DEGs) in each. The ATH1 datasets for iTPS 29.2, iTPS 31.3 lines and alcR lines are from this study. They are for treatments similar to those described in Martins *et al*., (2013) and were either sprayed at dawn and harvested 12 h later at the end of the day (ED) or were sprayed at dusk and harvested 12 h later at the end of the night (EN). These data are provided in Supplemental Dataset S1, and analyzed in Supplemental Figures S1 and S3 and Supplemental Tables S2-S4. The RNA seq data using iTPS 29.2 are for iTPS 29.2 and alcR lines from this study. They are for plants sprayed 0.5 h after dawn and harvested 4 or 6 h later, respectively. Whilst a maximum of about 33K transcripts can be detected, the number of transcripts detected across lines and treatments in our study was about 23.8K. These data are provided in Supplemental Dataset S4 and analysed in Figures 2-8, Supplemental Figures S5-S24, and Supplemental Tables S4-S6. oeTPS refers to a constitutive overexpression of bacterial TPS, published in Zhang et al., (2009). The downloaded data are analyzed in Figure 6, Supplemental Figure S12 and Supplemental Table S5. tSnRK1α1 efers to transient overexpression in protoplasts, published in Baena-Gonzales et al. (2007), respectively. The downloaded data are analyzed in Figure 8, Supplemental Figure S14 and Supplemental Table S6. oebZIP11 refers to transient over expression of bZIP11 for 2 h in 7 days old published in Hanson et al. (2008) and Ma et al. (2011). For the published datasets, data was provided only for transcript that showed a significant change, as scored in the publication. The value under 'total genes detected' refers to the number of features on the array, and is given as NN* as the actual number detected will have been lower. h – hours, d – days.

| Line | Dataset | Tissue analyzed | Age or time after induction | Total genes detected | DEGs FDR<0.05 | Up (%) | Down (%) | DEGs FDR<0.05 and FC>2 | Up (%) | Down (%) |
|---|---|---|---|---|---|---|---|---|---|---|
| iTPS 31.3 | ATH1 | whole rosette | ED | 22.7K | 290 | 62 (21.4) | 228 (78.6) | 68 | 9 (13.2) | 59 (86.8) |
| iTPS 31.3 | ATH1 | whole rosette | EN | 22.7K | 396 | 198 (50) | 198 (50) | 76 | 24 (31.6) | 52 (68.4) |
| iTPS 29.2 | ATH1 | whole rosette | ED | 22.7K | 1459 | 684 (46.9) | 775 (53.1) | 189 | 43 (22.8) | 146 (77.2) |
| iTPS 29.2 | ATH1 | whole rosette | EN | 22.7K | 223 | 104 (46.6) | 119 (53.4) | 92 | 30 (32.6) | 62 (67.4) |
| iTPS 29.2 | RNA seq | whole rosette | 4 h | 23.8K | 13400 | 6530 (48.7) | 6870 (51.3) | 5618 | 2066 (36.8) | 3552 (63.2) |
| iTPS 29.2 | RNA seq | whole rosette | 6 h | 23.8K | 13109 | 10934 (83.4) | 2175 (16.6) | 5437 | 2017 (37.1) | 3420 (62.9) |
| oeTPS | ATq3.6.2 | seedling | 7 d | 29K* | - | - | - | 5272 | | |
| tSnRK1α1 | ATH1 | leaf protoplasts | | 24K* | - | - | - | 1021 | | |
| oebZIP11 | ATH1 | seedling | 7 d, 2 h | 24K* | - | - | - | 236 | | |

**Supplemental Table S2. Changes in transcript abundance 12 h after induction of TPS, investigated at either ED or EN.**

iTPS 29.2, iTPS 31.1 and alcR were either sprayed at dawn and harvested 12 h later at the end of the day (ED) or were spayed at dusk and harvested 12 h later at the end of the night (EN).

The upper part summarizes the total number of shared DEGS (Filter: FDR<0.05 only, no fold-change cutoff between line 29.2 and line 31.3 at ED and EN), and the correlation between the response in the two lines at ED and at EN.

In the middle part, the assignment of transcript to GRF groups $G_1$, $G_2$ and $G_O$ (see Supplemental Figure S2) are shown separately for line 29.2 and line 31.3, and the correlation between the change in transcript abundance and the CRF value in the sets of transcripts assigned to $G_1$ $G_2$ and $G_0$

The lower part summarizes the % of DEGS assigned to $G_1$and $G_2$ at ED and EN.

Correlation between iTPS 29.2 and iTPS 31.3 for the response of all shared DEGs

| Line | Lines | CRF | treatment | Number of DEGs | $R^2$ | Slope direction | p value |
|---|---|---|---|---|---|---|---|
| iTPS | 29.2 vs 31.3 | - | ED | 254 | 0.9565 | + | $1.35x10^{-173}$ |
| iTPS | 29.2 vs 31.3 | - | EN | 172 | 0.9821 | + | $2.23x10^{-150}$ |
| iTPS | 29.2 vs 31.3 | $G_1$ | ED | 138 | 0.969 | + | $4.12x10^{-104}$ |
| iTPS | 29.2 vs 31.3 | $G_2$ | ED | 88 | 0.925 | + | $4.46x10^{-49}$ |
| iTPS | 29.2 vs 31.3 | $G_0$ | ED | 30 | 0.977 | + | $1.47x10^{-24}$ |
| iTPS | 29.2 vs 31.3 | $G_1$ | EN | 43 | 0.9781 | + | $1.16x10^{-35}$ |
| iTPS | 29.2 vs 31.3 | $G_2$ | EN | 108 | 0.9837 | + | $1.22x10^{-96}$ |
| iTPS | 29.2 vs 31.3 | $G_0$ | EN | 21 | 0.9895 | + | $2.99x10^{-20}$ |

Correlation between the C response factor (CRF) and the iTPS response of a given line after assigning transcripts to $G_1$, $G_2$ or $G_0$

| Line | Line | CRF | tireatment | Number of DEGs | $R^2$ | Slope direction | p value |
|---|---|---|---|---|---|---|---|
| iTPS | 29.2 | $G_1$ | ED | 706 | 0.67 | + | $5.4x10^{-172}$ |
| iTPS | 29.2 | $G_2$ | ED | 327 | 0.656 | - | $1.73x10^{-77}$ |
| iTPS | 29.2 | $G_0$ | ED | 420 | 0.019 | | 0.005 |
| iTPS | 29.2 | $G_1$ | EN | 53 | 0.76 | + | $2.13x10^{-17}$ |
| iTPS | 29.2 | $G_2$ | EN | 127 | 0.754 | - | $6.2x10^{-40}$ |
| iTPS | 29.2 | $G_0$ | EN | 43 | 0.0001 | | 0.95 |
| iTPS | 31.3 | $G_1$ | ED | 166 | 0.645 | + | $8.9510x10^{-39}$ |
| iTPS | 31.3 | $G_2$ | ED | 91 | 0.435 | - | $1.1x10^{-12}$ |
| iTPS | 31.3 | $G_0$ | ED | 33 | 0.004 | | 0.732 |
| iTPS | 31.3 | $G_1$ | EN | 114 | 0.69 | + | $2.63x10^{-30}$ |
| iTPS | 31.3 | $G_2$ | EN | 200 | 0.684 | - | $2.21x10^{-51}$ |
| iTPS | 31.3 | $G_0$ | EN | 82 | 0.035 | | 0.09 |

| | % of DEGs in $G_1$ | | % of DEGs in $G_2$ | |
|---|---|---|---|---|
| | iTPS 29.2 | iTPS 31.1 | iTPS 29.2 | iTPS 31.1 |
| ED | 48 | | 23 | |
| EN | 24 | | 57 | |

**Supplemental Table S3. Summary of the number of differentially expressed genes, their assignment to CRF groups and the $R^2$, direction and p-value of the relationship between the change in transcript abundance and the CRF value** (extends Figure 2 and Supplemental Figure S5A-B).

The Table lists the numbers of DEGs (FDR < 0.05) that were assigned to CRF groups $G_1$, $G_2$ and $G_O$ for the 4-h and the 6-h data sets, as well as for transcript that showed a shared response and alignment in both the 4-h and 6-h data sets (termed 4-6h shared). The last three columns show the $R_2$ value, the slope direction and the p-value when, for that set of genes, the change in transcript abundance is plotted against their CRF value.

| | | | | Plot of change in transcript abundance against CRF | | |
|---|---|---|---|---|---|---|
| **Line** | **CRF** | **Time after induction** | **Number of DEGs** | **$R^2$** | **Slope direction of plot** | **p value** |
| iTPS 29.2 | $G_1$ | 4 h | 4576 | 0.45 | + | 0 |
| iTPS 29.2 | $G_2$ | 4 h | 2887 | 0.35 | - | $2.11 \times 10^{-274}$ |
| iTPS 29.2 | $G_0$ | 4 h | 3573 | 0.002 | | 0.0067 |
| iTPS 29.2 | $G_1$ | 6 h | 4470 | 0.48 | + | 0 |
| iTPS 29.2 | $G_2$ | 6 h | 2969 | 0.37 | - | $2.17 \times 10^{-296}$ |
| iTPS 29.2 | $G_0$ | 6 h | 3636 | 0.0007 | | 0.11 |
| iTPS 29.2 | $G_1$ | 4-6 h shared | 3769 | 0.92 | + | 0 |
| iTPS 29.2 | $G_2$ | 4-6 h shared | 2338 | 0.90 | + | 0 |
| iTPS 29.2 | $G_0$ | 4-6 h shared | 2890 | 0.92 | + | 0 |

**Supplemental Table S4. Pairwise comparison of responses 4-h and 6-h post-induction in the light, and in the ED and the EN treatments (extends Supplemental Figure S5C).**

Pairwise comparisons of genes assigned to different CRF groups are made between all treatments. Data is plotted for transcripts that showed a significant change in abundance in at least one of the treatments (FDR < 0.05, $Log_2$ FC ≥ 0.2). The two treatments that are compared are shown in the two left-hand columns The next two columns identify which treatment was used to filter, and which CRF group is compared. Transcripts were assigned to CRF group 1 when their response to elevation of Tre6P was in the same direction as their response to an increase of sugar supply in a panel of treatments (as defined by the CRF); transcripts assigned to CRF group 2 respond to elevation of Tre6P in the opposite direction to the response to an increase of sugar supply; transcripts assigned to CRF group 0 do not show any notable response to a change in sugar in the panel of treatments (for details see Supplemental Figure S2).

| | | Filtered by: | Filtered by : | | | | |
|---|---|---|---|---|---|---|---|
| Treatment a | Treatment b | treatment a | treatment b | Number of DEGs | $R^2$ | p-value | Slope direction |
| iTPS (4h) | iTPS (ED) | $G_1$ | | 451 | 0.632 | $8.2x10^{-101}$ | + |
| iTPS (4h) | iTPS (ED) | | $G_1$ | 519 | 0.413 | | + |
| iTPS (4h) | iTPS (ED) | $G_2$ | | 287 | 0.191 | $1.1x10^{-15}$ | + |
| iTPS (4h) | iTPS (ED) | | $G_2$ | 244 | 0.403 | | + |
| iTPS (4h) | iTPS (ED) | $G_0$ | | 202 | 0.022 | 0.024 | + |
| iTPS (4h) | iTPS (ED) | | $G_0$ | 197 | 0.018 | | + |
| iTPS (6h) | iTPS (ED) | $G_1$ | | 491 | 0.723 | $1.1x10^{-138}$ | + |
| iTPS (6h) | iTPS (ED) | | $G_1$ | 528 | 0.574 | | + |
| iTPS (6h) | iTPS (ED) | $G_2$ | | 275 | 0.355 | $7.3x10^{-29}$ | + |
| iTPS (6h) | iTPS (ED) | | $G_2$ | 240 | 0.574 | | + |
| iTPS (6h) | iTPS (ED) | $G_0$ | | 205 | 0.055 | 0.00068 | + |
| iTPS (6h) | iTPS (ED) | | $G_0$ | 205 | 0.055 | | + |
| iTPS (4h) | iTPS (EN) | $G_1$ | | 50 | 0.530 | $5.4x10^{-10}$ | + |
| iTPS (4h) | iTPS (EN) | | $G_1$ | 48 | 0.315 | | + |
| iTPS (4h) | iTPS (EN) | $G_2$ | | 100 | 0.408 | $1.3x10^{-13}$ | + |
| iTPS (4h) | iTPS (EN) | | $G_2$ | 108 | 0.517 | | + |
| iTPS (4h) | iTPS (EN) | $G_0$ | | 33 | 0.149 | 0.026 | + |
| iTPS (4h) | iTPS (EN) | | $G_0$ | 34 | 0.169 | | + |
| iTPS (6h) | iTPS (EN) | $G_1$ | | 59 | 0.519 | $1.2x10^{-10}$ | + |
| iTPS (6h) | iTPS (EN) | | $G_1$ | 49 | 0.533 | | + |
| iTPS (6h) | iTPS (EN) | $G_2$ | | 103 | 0.560 | $2.2x10^{-19}$ | + |
| iTPS (6h) | iTPS (EN) | | $G_2$ | 114 | 0.573 | | + |
| iTPS (6h) | iTPS (EN) | $G_0$ | | 33 | 0.297 | 0.001046 | + |
| iTPS (6h) | iTPS (EN) | | $G_0$ | 33 | 0.297 | | + |

**Supplemental Table S5: Comparison of the responses to constitutive overexpression of TPS (oeTPS) and induced overexpression of TPS (iTPS) (supplemental to Figure 6 and Supplemental Figure S12).** The oeTPS response (from Zhang *et al.*, 2009) was filtered by the authors (FDR<0.05, FC≥2), The Tables show the number of DEGs shared DEGS with the iTPS response, after filtering and assigning genes in the iTPS response to CRF groups.

(A) Comparison of the oeTPS response with the overall iTPS response, either unfiltered or after filtering showing numbers of shared genes and p-value of the regression

(B) Comparison of the oeTPS response with the TPS response after filtering it (FDR<0,05) and assigning genes in the iTPS response to the CRF groups $G_1$, $G_2$ or $G_0$. The Table summarizes the % of genes in the oeTPS response that were assigned to a given CRF group (for details of assignment to CRF groups, see Supplemental Figure S2).

A. · Comparison of the oeTPS response with the overall iTPS response, either unfiltered or after filtering showing numbers of shared genes and p-value of the regression

| Lines | ALL | p Value | FDR < 0.05 | p Value | FDR < 0.05 + FC2 | p Value |
|---|---|---|---|---|---|---|
| 4h vs oeTPS | 5005 | $1.41x10^{-91}$ | 3340 | $8.26x10^{-95}$ | 1321 | $5.59x10^{-32}$ |
| 6h vs oeTPS | 4988 | $1.41x10^{-149}$ | 3353 | $1.1x10^{-144}$ | 1372 | $4.7x10^{-59}$ |

B. Comparison of the oeTPS response with the TPS response after filtering it (FDR<0,05) and assigning genes in the iTPS response to the CRF groups $G_1$, $G_2$ or $G_0$. The tables summaries the % of genes in the oeTPS response that were assigned to a given CRF group (for details of assignment to CRF groups, see Supplementary Figure S2)

| G1 | | DEGs assigned to $G_1$ in iTPS and are present also in oeTPS | |
|---|---|---|---|
| # shared iTPS | Comparisons | # | % |
| | 4h vs oeTPS | 1837 | 40.1 |
| | 6h vs oeTPS | 1890 | 42.3 |
| 3769 | 4-6h vs oeTPS | 1596 | 42.3 |

| G2 | | DEGs assigned to $G_2$ in iTPS and are present also in oeTPS | |
|---|---|---|---|
| # shared iTPS | Comparisons | # | % |
| | 4h vs oeTPS | 612 | 21.2 |
| | 6h vs oeTPS | 619 | 20.8 |
| 2338 | 4-6 vs oeTPS | 484 | 20.7 |

| G0 | | DEGs assigned to $G_0$ in iTPS and are present also in oeTPS | |
|---|---|---|---|
| # shared iTPS | Comparisons | # | % |
| | 4h vs oeTPS | 406 | 14.1 |
| | 6h vs oeTPS | 450 | 15.2 |
| 2890 | 4-6h vs oeTPS | 347 | 12.0 |

**Supplemental Table S6. Comparison of the tSnRK1α1 and iTPS responses (extends Figure 8 and Supplemental Figure S14).**

The Table shows the relationship between all 1005 genes from the tSnRK1α1 data set (FDR < 0.05, $Log_2$ FC ≥ 1) that were retrieved in the unfiltered iTPS data set (ALL), and the relationship for the subsets of genes from the tSnRK1α1 response that were retained in the filtered iTPS response (FDR < 0.05, $Log_2$ FC ≥ 0.2) and assigned to the CRF groups $G_1$, $G_2$ and $G_0$ (for details of assignments to CRF groups, see Supplemental Figure S2).

| Comparison | CRF in the iTPS response | Number of DEGs | $R^2$ | p Value |
|---|---|---|---|---|
| iTPS 4h vs tSnRK1α1 | ALL | 1005 | 0.140 | $7.63 \times 10^{-35}$ |
| | $G_1$ | 580 | 0.644 | $7 \times 10^{-132}$ |
| | $G_2$ | 144 | 0.415 | $2.9 \times 10^{-18}$ |
| | $G_0$ | 22 | 0.042 | 0.36 |
| iTPS 6h vs tSnRK1α1 | ALL | 1005 | 0.212 | $6.97 \times 10^{-54}$ |
| | $G_1$ | 541 | 0.677 | $1.64 \times 10^{-134}$ |
| | $G_2$ | 151 | 0.286 | $1.44 \times 10^{-12}$ |
| | $G_0$ | 22 | 0.074 | 0.22 |
