## Supplemental Text for "Direct and indirect responses of the Arabidopsis transcriptome to an induced increase in trehalose 6-phosphate"

Avidan et al.

### Supplemental Text. Analysis of selected areas of metabolism, signaling and development

| ***Subsection*** | Page |
|---|---|
| ***Initial experiment with harvest 12 hours post-induction*** | ***2*** |
| ***MapMan BINS related to metabolism*** | ***3*** |
| *Photosynthesis* | *3* |
| *Gluconeogenesis* | *3* |
| *N metabolism* | *3* |
| *Nucleotide metabolism* | *4* |
| *Specialized metabolism* | *4* |
| ***Sucrose metabolism and transport*** | ***4*** |
| ***Protein synthesis, ribosomal proteins and ribosome biogenesis*** | *5* |
| ***Cell wall synthesis and modification; cell expansion*** | *5* |
| ***Flowering induction*** | ***6*** |
| ***Circadian clock components*** | ***6*** |
| ***Light signaling*** | ***7*** |
| ***TOR signaling*** | ***9*** |
| ***Finger-Like Zinc (FLZ) family proteins*** | ***11*** |
| ***Brassinosteroid signaling*** | ***13*** |
| ***Cell wall modification*** | ***14*** |
| ***Overlap with bZIP11 signaling*** | ***14*** |
| ***Transcription factors*** | ***17*** |
| ***References*** | ***20*** |

***Initial experiment with harvest 12 hours post-induction***

In a first experiment, we investigated if the response was similar in two separate iTPS lines, iTPS29.1 and iTPS321.3. This initial experiment was carried out using 12-h induction and also served to show that shorter treatments were needed and that induction should be carried out in the light period, because indirect effects predominated, especially during the night.

iTPS lines 29.2 and 31.3 and control alcR line plants were sprayed with 2% ethanol or water at the beginning of the day and harvested 12 h later at the end of the day (ED treatment) or were sprayed at dusk and harvested 12 h later at the end of the night (EN treatment). In published data for metabolite levels in the same plant material, after ethanol induction Tre6P levels increased 3- to 4-fold at ED and 2-to 3-fold at EN compared to the control treatments (Martins et al., 2013, see also Figueroa *et al.,* 2016 for separate experiments). Transcript abundance was analyzed using the Affymetrix ATH1 array (Supplemental Dataset S1). The ethanol-sprayed iTPS line was compared to the corresponding water-sprayed iTPS control to calculate the fold-change (FC) for each transcript. These were corrected for minor responses to alcR (see Methods). We termed the resulting change in transcript abundance the 'iTPS response'. Line 29.1 showed a stronger response than line 31.3 in the ED treatment, and a similar response to line 31.3 in the EN treatment (Supplemental Figure 1A, Supplemental Table S1). For shared genes, the responses in the two lines were strongly correlated (Supplemental Figure S1A-C, Supplemental Table S2, $R^2$ = 0.96 and 0.98 for FDR<0.05 filtered data sets at ED and EN, respectively).

Further analyses were performed with the data set for line 29.1. When the iTPS response was compared with CRF values (for explanation and calculation see main test and Supplemental Figure S2) across all transcripts there was little similarity in the ED treatment and even less in EN treatment (Supplemental Figure S3A-B). Transcripts assigned to $G_1$ showed a strong positive correlation between their CRF values and iTPS response ($R^2$ = 0.67 and 0.76 at ED and EN, respectively) consistent with them responding to elevated Tre6P (Supplemental Figure S3C-D, Supplemental Table S2). Transcripts assigned to $G_2$ showed a strong negative correlation between their CRF values and iTPS response, consistent with them responding to the decrease in sugars (Supplemental Figure S3C-D, Supplemental Table S2). There was a strong correlation between the response at ED and EN of transcripts that were assigned to $G_1$, or to $G_2$, or to $G_0$ (Supplemental Figure S2E).

A larger proportion of transcripts were assigned to $G_1$ and a smaller proportion to $G_2$ at ED than at EN (Supplemental Table S2). This might reflect the larger decrease of sugars in the EN than the ED treatment. At night, the sucrose pool is strongly dependent on starch mobilization and falls because elevated Tre6P leads to a strong inhibition of starch mobilization (Martins et al., 2013; dos Anjos et al., 2018). In the light period, elevated Tre6P leads to diversion of more of the fixed C to organic acid and amino acid synthesis, but the sucrose pool is partly stabilized by photosynthetic C fixation and possibly by changes in sucrose export (Figueroa et al., 2016).

This initial experiment with harvest 12-h post induction led us to focus on the iTPS response in the light period, to harvest at earlier times post-harvest and to focus on line 29.2.

#### *MapMan BINS related to metabolism*

As highlighted in the main text and Figure 3, analysis of the iTPS response at the highest level of the MapMan ontology (BINs) revealed that genes assigned to CRF groups $G_1$, $G_2$ and $G_0$ were often associated with different functions. MapMan BINs group genes that participate in a given metabolic sector or cellular function, irrespective of whether they are involved in biosynthesis or catabolism. In some BINs, different processes are also grouped together. (e.g., 'Cell wall' includes biosynthesis of cell wall components, but also their modification and degradation; 'Protein' includes protein synthesis, protein modification and protein degradation). We therefore inspected the responses in selected BINs at higher resolution (Supplemental Figure S7) to provide further support for the separate identify of the $G_1$ response (Tre6P-mediated) and $G_2$ response (mediated by the decline in sucrose or other indirect effects). This analysis was also performed to learn if the $G_0$ response (respond to iTPS but not to changes in sugar availability) was associated with any specific functions.

**Photosynthesis**: Resolving the photosynthesis BIN into subBINS revealed that many genes encoding proteins involved in the light reactions, electron transport, photorespiration and the Calvin-Benson cycle were assigned to $G_1$ and repressed (Supplemental Figure S7A). This indicates that elevated Tre6P drives a rapid and broad transcriptional repression of photosynthesis. In the Discussion section, we argue that this provides very strong evidence that the well-known repression of photosynthetic gene expression by sugar is not just due to changes in the N status andNn-signaling but includes a strong input from C-signaling, mediated at least partly by Tre6P.

**Gluconeogenesis:** Genes involved in the glyoxylate cycle and gluconeogenesis were assigned to $G_1$ and repressed, with the exception of isocitrate lyase that was assigned to $G_0$ and induced (Supplemental Figure S7B). It is not clear why isocitrate lyase shows such a divergent response.

**N metabolism**: A more complex picture was found for N metabolism (Supplemental Figure S7C). This BIN includes genes involved in nitrate and ammonium assimilation, which are typically induced by sugar (Vincentz et al., 1993; Krapp et al., 1995; Wang et al., 2000; Curuzzi 2003; Usadel et al., 2008, Parul et al., 2016, Vidal et al., 2020, see also the CRF values in Supplemental Dataset S4). These genes were assigned to $G_2$ and repressed in the iTPS response, implying that repression is not mediated by Tre6P (o,r at least, not primarily mediated by Tre6P) but is, instead, an indirect response to low sugar.

Glutamate dehydrogenases (GDHs) are induced by low sugar (Osuna et al., 2007; Jean-Xavier et al., 2012; Cookson et al., 2016) and contribute to recycling of C skeletons from amino acids (Coruzzi, 2003; Fontaine et al., 2012). They were repressed in iTPS and assigned to $G_1$, indicating that they are repressed by Tre6P, resembling genes involved in gluconeogenesis (see above). Indeed, the major function of GDHs may be in recycling of C skeletons under low-C conditions.

**Nucleotide metabolism**: In nucleotide metabolism (Supplemental Figure S7D), genes involved in purine biosynthesis and, to a greater extent, genes involved in purine biosynthesis were mainly assigned to CRF group $G_1$ and induced, whereas genes involved in nucleotide salvage or breakdown, in equilibration of nucleotide phosphates or in synthesis of deoxy DNA precursors were assigned to various CRF groups and showed mixed responses.

**Secondary metabolism**: In the secondary metabolism BIN (Supplemental Figure S7E), genes involved in tocopherol and carotenoid biosynthesis were partly assigned to $G_1$ and repressed (resembling the repression of genes encoding photosynthesis proteins, see Supplemental Figure 7) and partly to $G_0$ and repressed (resembling the response for tetrapyrrole biosynthesis, see Figure 3).

This contrasted with genes involved in phenylpropanoid biosynthesis, flavonoid biosynthesis and, even more strikingly, glucosinolate biosynthesis that were assigned to $G_2$ and repressed, presumably as a response to the decline in sugars. The responses of transcription factors involved in the regulation of glucosinolate biosynthesis are presented in the main text, Results section 'Changes in expression of transcription factors'.

#### *Sucrose metabolism and transport*

Given that Tre6P is known to regulate sucrose metabolism (see Introduction, also Figure 1), we inspected the responses of individual genes involved in sucrose metabolism and transport (Supplemental Figure S8).

There were complex changes in expression of members of the sucrose phosphate synthase (SPS) family that is involved in sucrose synthesis, and members of the sucrose synthase (SUS) and invertase (INV) families that are involved in sucrose degradation. Responses in $G_1$ included a weak induction of

*SPS1* and stronger repression of *SPS4*, induction of *SUS1* and repression of *SUS2*, and repression of *VINV1*, *A/N-InvC*, *A/N-InvE*, and *A/N-invH*.

There were widespread changes for sucrose-$H^+$ transporter (SUT) and the sugar efflux SWEET family members (for background on their functions see Xue *et al*., 2022; Braun 2022). *SUT1/ SUT/AT1g 22710*, the sucrose/$H^+$ transporter that catalyzes active uptake of sucrose from the cell wall into companion cells in leaves, was weakly repressed at 6h. There were mixed responses for other *SUT* family members. There was a large decrease in transcript abundance for *SWEET11* and *SWEET12*, which mediate passive efflux of sucrose from the phloem parenchyma cells into the cell wall, and *SWEET13*, which mediates passive export of sucrose from bundle sheath cells into the cell wall, with *SWEET11* and *SWEET12* being assigned to $G_1$ indicating they are repressed by Tre6P-signaling, and *SWEET13* being assigned to $G_2$ indicating an indirect response. It might be noted that *SWEET12* was also reported to be induced by constitutive overexpression of bacterial TPS in Arabidopsis (Zhang et al., 2009) but to be repressed by vascular tissue-specific overexpression of TPS in Arabidopsis (Fichtner et al., 2021) and by supplying a caged Tre6P precursor to wheat shoots (Oszwald et al., 2018). Possible reasons for these diverging responses are given in the Discussion

Many other *SWEETs* were also repressed, including *SWEET16* and *SWEET17*, with the former responding in a way consistent with it being repressed by Tre6P. *SWEET16* and *SWEET17* are located on the tonoplast and implicated in vacuolar storage of fructose (Guo et al., 2013; Chardon et al., 2013; Klemens et al., 2013). Their repression might decrease recycling of fructose from the vacuole in C-replete conditions.

SWEET proteins have been implicated in the regulation of sink-source interactions by FT homologs in potato (Aberlanda *et al.*, 2019) and SWEET10 has been implicated in the induction of flowering by FLOWERING TIME (FT) in Arabidopsis (Andrés *et al.* 2022) *SWEET10* transcript abundance was not significantly altered in the iTPS response (Supplemental Figure S8)

#### *Protein synthesis, ribosomal proteins and ribosome biogenesis*

The BIN 'Protein synthesis' represents a very large set of genes with diverse functions, including not only genes for protein synthesis but also for post-translational modification and for protein degradation. To focus better on the process of protein synthesis, we inspected the subBINs for amino acid activation and the translation process, ribosomal proteins and ribosome biogenesis (Figure 4, Supplemental Figure S7F).

Genes involved in amino acid activation and translation (initiation, elongation, release) were mainly assigned to $G_1$ and induced (Supplemental Figure S7F). Genes encoding cytosolic and mitochondrial ribosomal proteins were assigned to $G_1$ and broadly induced (Figure 4A, Supplemental Figure S7F), whereas genes encoding chloroplast ribosomal proteins were assigned to $G_2$ and repressed (Figure 4A. Supplemental Figure S7F), reminiscent of the response of genes involved in photosynthesis. Genes involved in ribosome biogenesis were assigned to $G_1$ and showed a particularly strong induction (Figure 4A). Ribosome biogenesis genes generally showed a positive correlation between their iTPS response and CRF value (Figure 4B) and a broadly reciprocal response to that after transient overexpression of SnRK1 in protoplasts (Baena-González et al., 2007) (Figure 8C).

#### *Cell wall synthesis and modification; cell expansion*

We also inspected the responses of genes involved in cell wall biosynthesis, which is a major consumer of C during cell growth, and cell wall modification, which is required for expansion growth (Supplemental Figure S7G).

In the BIN 'Cell wall', many genes assigned to cellulose, hemicellulose and pectin synthesis were assigned to $G_2$ and repressed, as were several genes for various classes of cell wall proteins, especially arabinogalactan proteins (AGPs). Overall, there was a trend to repression of genes involved in cell wall synthesis but much of this response is indirect.

Some genes involved in cell wall modification were assigned to $G_1$ and even more genes to $G_2$, with both sets being repressed. This repression affected both expansins (*EXPA*s) and xyloglucan endotransglucosylases (*XTH*s) (see below, sections Brassinosteroid signaling' and 'Cell wall modification' for more analysis). Pectin esterases were assigned to $G_1$, $G_2$ and $G_0$, with an overall trend to induction. This indicates a mixed impact of an induced elevation of Tre6P on cell wall modification with some direct effects but many indirect effects, which tend to be repressive.

#### *Flowering induction*

Flowering time is regulated by a large number of pathways that provide information about photoperiod, temperature, plant maturity and metabolic status (Jin and Ahn, 2020; Quiroz et al., 2021; Izawa 2021). Tre6P has been implicated in the regulation of floral initiation (Schleupmann et al., 2003, Wahl et al., 2013, Fichtner et al., 2020), acting to promote the CONSTANS/FLOWERING TIME (CO/FT) photoperiod pathway (for background see Turck et al., 2007; Shim et al. 2017) and the miR156 maturity pathway (see also Wang, 2014). Further, the delay in the floral transition in mutants with constitutively low Tre6P levels is partly reverted in mutants with modified SnRK1 function (Zacharaki et al., 2022). We therefore inspected how genes involved in the various floral induction pathways respond to transient elevation of Tre6P (Supplemental Fig S9).

Many floral induction genes were assigned to $G_1$ and repressed by elevated Tre6P. This includes S*UPPRESSOR OF CONSTANS1* (*SOC1*), *SQUAMOSA BINDING PROTEIN LIKE* (*SPL*) *SPL3* and *SPL4*, *SHORT VEGETATIVE PHASE* (*SVP*) and *PHYTOCHROME INTERACTING FACTOR 3* (*PIF3)* and *PIF4)*, which are inhibitory genes in the temperature-dependent flowering pathway (Yin and Ahn 2021; Brightbill and Sung 2022). *PIFs* are also involved in the maturity pathway (Wang, 2014). In addition to this repression of *PIF3* and *PIF4*, *PIF5* was also repressed, although its assignment to CRF $G_2$ indicates this is not due to Tre6P-signaling.

Other genes repressed by elevated Tre6P and assigned to CRF $G_1$ included members of the *GIBBERELLIN-INSENSITIVE DWARF* family (*GID1A*, *GID1C*, and *TEMPRANILLO* family (*TEM1*, *TEM2*) that are involved in the hormonal regulation of flowering under long days (Griffiths et al., 2007; Takeshi et aI., 2021). Gibberellin signaling interacts negatively with DELLA proteins (Li et al., 2018; Takeshi ,2021; Takeshi *et al*., 2021). The DELLA protein *REPRESSOR OF GA1-3* 1 (RGA1) was assigned to $G_1$ and weakly repressed. Three further DELLAs (*RGA2/GAI* and *RGA-LIKE 1* (*RGL1*) and *RGL3*) were assigned to $G_2$, with *RGA2* and *RGL1* being strongly repressed and $RGL_3$ being weakly induced.

Many genes involved in meristem identity were repressed but assigned to $G_2$ indicating that the responses may be indirect. Several genes were assigned to $G_O$ indicating that they may be regulated by Tre6P in a manner that is cryptic in wild-type plants; this included the *FT-INTERACTING PROTEIN 1* (*FTIP1*) that is required for FT transport from the leaf to the shoot apex (Liu et a*l*., 2013) and *FD/BZIP14* that interacts with FT at the shoot apex with FT to promote flowering (Abe et a*l*., 2005). Other genes assigned to $G_0$ included the LOV-domain blue light receptor *ZEITLUPE* (*ZTL*) as well as *SPL5* and *SPL9* which, like *SPL4*, were repressed. SPL family genes are downstream of miR156 in the age floral induction pathway (Wang et al., 2014), which has previously been implicated in the regulation of flowering by Tre6P (Wahl et al., 2013)

It has previously been reported that constitutively elevated Tre6P leads to increased expression of *FLOWERING TIME* (*FT*) (Wahl et al., 2013; Fichtner et al., 2020) and accelerates flowering in long days. *FT* expression was not significantly increased in the iTPS data set but this may be because the plants were harvested early in the light period, before FT transcript rises.

#### *Circadian clock components*

Sugar-signaling can modulate the circadian clock (Haydon et a*l*., 2013; Frank et al., 2018; Webb *et al*., 2019; Viani *et al*., 2022). We Inspected the iTPS response for core components of the circadian clock (Supplemental Fig S10), to learn if these responses were in part due to Tre6P-mediated signaling.

There were widespread changes in transcript abundance of clock components including *PSEUDO RESPONSE REGULATOR 5* (*PRR5)* and all three components of the Evening Complex (*EARLY FLOWERING 3* (*ELF3*), *ELF4* and *LUX ARRHYTHMO*, all of which were assigned to $G_1$ and repressed by iTPS. *PRR7* has been reported to be induced by low C (Haydon *et al.*, 2013; Moraes *et al*., 2019), but no significant change was found in the iTPS response (see later in this section for a possible explanation) . Relatively few circadian clock genes were assigned to $G_2$, but some were assigned to $G_0$ including *GIGANTEA* (*GI*) and the clock output *CONSTANS* (*CO*) that induces *FT* in a light-dependent and clock-gated manner from about 12-14 h after dawn onwards (Mikael and Dorothee, 2014; Jae et al., 2017) to induce flowering in long days.

There was a general trend to increased abundance of transcripts (especially at 6h post-induction) for dawn clock components that peak at dawn and subsequently decline, and decreased abundance of components for day, dusk and especially evening components that are low at dawn and rise during the light period (Supplemental Figure S10). This observation is consistent with a rise in Tre6P early in the light period leading to a delay in the subsequent progression of the clock. The magnitude of these changes (FC on a $\log_2$ scale) was up to 0.7 for dawn genes, and to -0.4 to -1-9 for day, dusk and evening genes with the largest observed decrease being observed for *ELF4* transcript. This was small relative to the amplitude of the diel changes of transcript abundance in plants growing in the conditions used for the RNAseq experiment (FC $\log_2$ 8-11; Flis et al., 2019; Moraes et al., 2019). These observations nevertheless point to elevation of Tre6P at 4-6 h into the light period having a measurable impact on core clock transcripts, possibly due to a delay in clock progression leading to slower decay of dawn transcripts and slower rise of transcript that peak later in the diel cycle. Indeed, ‘Entrainment of the circadian clock’ was highlighted as an over-enriched category in the GO analysis of the iTPS CRF $G_1$ response (Supplemental Figure S11).

The absence of a significant response of *PRR7* transcript is puzzling, as this gene has been reported to be a target of low sugar signaling (Haydon et al., 2013; Moraes et al., 2019) mediated by SnRK1-dependent phosphorylation of bZIP11 (Frank et al. 2019; Viani et al., 2021). It is possible that the absence of a response of *PRR7* is because it has reached its diel peak at 4-6 h after dawn and relatively insensitive to a transient elevation of Tre6P at this time. Alternatively, the absence of a response of *PRR7* might reflect the complexity of upstream signaling with both Tre6P and sugars providing input, that in wild-type plants would change in parallel but in the iTPS response change in a reciprocal manner. For example, it has been proposed that sucrose acts via ZTL (which is repressed in the iTPS response and assigned to CRF G0) to stabilize GI protein (Dalchau et al., 2011; Haydon et al.,2017). An additional factor may be that light modulates action of SnRK1 on the clock (Shin et al., 2017).

#### *Light signaling*

Paul et al. (2010) identified 23 genes that are involved in light signaling and were upregulated in response to constitutive overexpression of bacterial TPS (oeTPS). Of these genes, 19 were present in the iTPS data set after fusing it with the set of genes for which CRF could be scored (see Supplemental Figure S2). We inspected the responses of these and further genes involved in light signaling (Supplemental Figure S12C).

Transcript abundance for all 19 genes was elevated in the iTPS response, with 17 being assigned to CRF $G_1$. This included *CCR-like* (*CCL*), *EARLY PHYTOCHROME RESPONSIVE 1* (*EPR1*), *REVEILLE2* (*CIR/RVE2*), *PHOTOTROPIN1* (*PHOT1*), P*HYTOCHROME KINASE SUBSTRATE 1* and *2* (*PKS1, PKS2*), *PHYTOCHROME INTERATING FACTOR 4* (*PIF4,* which has previously been shown to be sugar regulated during diel cycle (Moraes et al. 2019) and some phototrophic response proteins. Further genes involved in light signaling that were strongly upregulated in the iTPS data set included the clock component *ELF4* (see also above), *PKS4*, further phototropic response proteins, whilst *ELONGATED HYPOCOTYL 5* (*HY5*) was repressed. The latter is involved in transcriptional regulation of many processes including photomorphogenesis, ABA signaling and anthocyanin biosynthesis inhibition (for more details see Discussion) . Several of these including *HY5* were assigned to CRF group $G_1$. Analysis of the iTPS response also uncovered induction of several genes involved in COP9 signaling and these were also induced in oeTPS dataset of Zhang et al. (2009). Overall, this comparison reveals remarkably agreement between the response to transiently elevated Tre6P and a constitutive increase in Tre6P, and provides evidence that this response largely due to signaling down stream of Tre6P.

Overall, as suggested by Paul et al. (2010), Tre6P interacts with and inhibits light signaling. This provides a mechanism whereby light-induced morphogenesis and growth responses can be modified and tuned by carbon availability.

#### *TOR signaling*

There was an increase in transcript abundance for many genes that encode ribosome biogenesis factors and structural components of the cytosolic and mitochondrial ribosome (Figure 4, Supplemental Figure S7F). TOR is a trimer consisting of the TOR catalytic subunit, REGULATORY-ASSOCIATED PROTEIN OF TOR (RAPTOR) and LETHAL WITH SEC THIRTEEN (LST), whereby both RAPTOR and LST are encoded by two genes. TOR is known to positively control ribosome assembly in mammals, fungi and plants, acting in a broadly opposed manner to SnRK1 (Sabatini, 2017; Ryabova et al., 2018; Wu et al., 2019; Meng et al., 2022; Scarpin et al., 2020; 2022). The molecular relationship between SnRK1 and TOR is not well understood, and probably complex (see also following section). However, the observation that SnRK1α1 (the catalytic subunit of SNRK1) and RAPTOR1B proteins interact in the cytosol (Nukarinin et al., 2016) and that SnRK1α1 phosphorylates RAPTOR1B protein i*n vitro* (Nukarinen et al., 2016) indicates that the mutual interactions may include direct action of SnRK1 on TOR. Another possible connection is that ABA-activated SnRK2s can directly phosphorylate RAPTOR or release activated SnRK1 to phosphorylate RAPTOR, thereby repressing TOR signaling (Wang et al., 2018; Belda-Palazon *et al.*, 2020). We therefore asked if the iTPS response provided any evidence for an interaction between Tre6P-signaling and TOR signaling.

We first inspected the impact of iTPS on expression of TOR subunits (Supplemental Figure S15A). There was little response, apart from a weak induction of RAPTOR 1, which the assignment to CRF $G_2$ indicated was unlikely to be a response to Tre6P signaling.

TOR acts by phosphorylating direct targets like S6Kinase, YET ANOTHER KINASE 1(YAK1) and LA-RELATED PROTEIN 1 (LARP1) that in turn phosphorylate diverse downstream targets (Sabatini, 2017; Ryabova et al., 2018; Wu et al., 2019; Scarpin et al., 2020; 2022). S6Kinase and LARP1 promote growth by stimulating ribosome biogenesis, protein translation and other processes (Scarpin et al., 2022), whereas inactivation of YAK1 promotes growth (Barrada et al., 2019; Forzani et al., 2019). The LARP1 protein is involved in a TOR-LARP1-50’TOP signaling axis that is conserved in plants and animals and regulates expression of 50’TOP mRNAs, including transcripts encoding ribosome assembly factors and ribosomal proteins (Scarpin et al., 2020; Scarpin et al. 2022). The resulting increase in ribosome abundance is one of the ways in which TOR orchestrates an increase in protein synthesis and growth.

We investigated whether elevated Tre6P increases expression of S6Kinases (S6PKs), LARP1s and YAK1 (Supplemental Figure S15B). Whilst Tre6P did not consistently alter transcript abundance for S6Kinases, *S6K1* was weakly induced 6-h after induction and was also strongly induced in the constitutive oeTPS data set of Zhang et al., 2009). Transcript abundance for *LARP1* family members increased, especially at 6-h. The responses of *LARPb* and *LARP1c* were assigned to CRF $G_1$, consistent with signaling downstream of Tre6P enhancing their expression, whereas LARP1a was assigned to $G_2$. Our analysis also revealed a weak induction of *YAK1*, but this response was assigned to CRF $G_0$ indicating it may be an indirect effect.

We also inspected the response of further genes implicated in the upstream regulation of ribosome biogenesis (Supplemental Figure S15C). There was weak induction of *NUCLEOSOME ASSEMBLY PROTEIN 1* (*NAP1:1*) and *RIBOSOMAL PROTEIN S6* (*RS6)* which jointly promote transcription of rRNA (Son et al., 2015). The response was significant for *NAP1:1* at 4-h and 6-h, and for *RS6* at 4-h. RS6 was also strongly induced in the oeTPS dataset of Zhang et al. (2009). The iTPS responses were assigned to CRF group $G_1$ and reciprocal responses were seen in tSnRK1α1 data set of Baena-Gonzalez et al. (2007) indicating that Tre6P might act to enhance expression of *NAP1:1* and *RSP,* and hence potentially increase RNA transcription, by inhibiting SnRK1 activity.

We next inspected whether elevated Tre6P might promote transcription of downstream TOR targets listed in Scarpin et al. (2020) and Meng et a*l*. (2022). The *PYR*/*PYL* gene family encodes ABA receptors, which are phosphorylated and inhibited by TOR (Wang et a*l.*, 2018a). Remarkably, all eight members of the gene family were repressed in the iTPS response (Supplemental Figure 15D, significant in all cases except *PYL3*), and four were assigned to CRF group $G_1$, indicating that their repression is mediated by Tre6P signaling rather than indirect effects. Further, three of the family (*PYL7, PYL8, PYR9*) were repressed in the constitutive oeTPS data set of Zhang et al. (2009). These findings point to a concerted inhibition of ABA signaling by TOR and by Tre6P, acting post-translationally and transcriptionally, respectively. Interestingly, most of these genes were significantly induced by tSnRK1α1 *(PYL2, PYL3, PYL5, PYL7, PYL8, PYR5,* data from Baena-Gonzaléz et al., 2007) consistent with Tre6P acting via inhibition of SnRK1 to repress ABA receptors.

Inspection of other TOR phosphorylation targets shortlisted by Scarpin et al. (2020) and Meng et al. (2022) (Supplemental Figure S15E) pointed to several of them being transcriptionally regulated by Tre6P-signaling (i.e., significant iTPS response, assigned to CRF group $G_1$, including weak induction of several elongation initiation factors (*elF4B1*, *elF4B2*, e*lF2B-δ1*), of *E2F TRANSCRIPTION FACTOR-3* that is involved in cell cycle division, of *ETHYLENE INSENSIVE 2* and of the auxin transporter *PIN2*, as well as inconsistent effects at 4h and 6h on transcripts encoding further translation initiation factors, *CONSERVED BINDING of eIFE* (*CBE1*), the developmental regulator *TOPLESS* and *VILLIN* actin-binding proteins. Many of these responses were also assigned to CRF groups $G_1$. There was also a small but significant increase in expression of the autophagosome assembly factor *ATG13a*, but this was assigned to CRF $G_2$.

Overall, this analysis revealed a potential synergy between the post-translational regulation of ribosome assembly, translation and other processes by TOR, and the transcriptional regulation of these same processes by signaling downstream of Tre6P, with an especially clear synergy between TOR and Tre6P for ABA receptors.

### ***Finger-Like Zinc (FLZ) family proteins***

FLZ family proteins (Jamsheer et al., 2015) are emerging as newly identified negative regulators of SnRK1 and, possibly, as being involved in interactions between SnRK1 and TOR (Nietsch et al., 2014; 2016; Jamsheer and Lamxi, 2015; Jamsheer et al., 2015; 2018a; 2018b; 2022; Bortlik et al., 2022). We therefore inspected their transcriptional response to iTPS.

As background, the expression of FLZ family proteins is known to be differentially regulated by sugars, cellular energy level and abiotic stress (Jamsheer and Laxmi, 2015). Whilst many *FLZs* are induced by high sugars, others are unaffected or repressed. This pattern is broadly reciprocal to their response to constitutive overexpression of SnRK1α1/KIN10, indicating that SnRK1 is involved in their response to sugars (Jamsheer and Laxmi 2015). Many FLZ proteins interact with and negatively regulate SnRK1 activity (Nietsch *et al*., 2014, 2016; Jamsheer et al., 2018a,2018b, 2022) by, at least for FLZ3, interfering with phosphorylation of the T-loop of SnRK1α1 (Bortlik et al., 2022). The finding that TOR activity is attenuated in some *flz* mutants (*flz6*, *flz10*, Jamsheer et al., 2018a; *flz8*, Jamsheer et al., 2022) supports a model in which FLZ proteins promote TOR signaling in high sugar conditions by inhibiting SnRK1 (Jamsheer et al., 2018a,2018b, 2022). There may also be a reciprocal interaction between TOR and FLZ function. The finding that several FLZ proteins (FLZ3, FLZ4, FLZ5, FLZ6, FLZ7) are encoded by mRNAs with 5'TOP motifs indicates that that their translation may be promoted by the TOR-LARP1-50'TOP signaling axis (Scarpin et al., 2022). The implication is that in conditions where TOR is activated, TOR may fine-tune translation of *FLZ* mRNAs to increase FLZ protein levels, restrict SnRK1 activity and promote growth (Scarpin et al., 2022). This emerging role of FLZ proteins as possible mediators between SnRK1- and TOR-signaling led to us to inspect whether their expression was modified by elevated Tre6P (Supplemental Figure S16).

FLZ family members are ordered in Supplemental Figure S16 according to the three sets that Jamsheer and Laxmi (2015) defined based on the response after exogenous adding sugar to seedlings; set 1 shows a large induction by sugar, set 2 a weaker induction and set 3 shows no response or repression. This assignment was largely confirmed by the CRF that we calculated for each gene response (Supplemental Figure S16). Most of the *FLZ* family members in set 1 (*FLZ1, FLZ5, FLZ8, FLZ14*) and some in set 2 (*FLZ3, FLZ10, FLZ15*) were significantly repressed in the iTPS response, as well as *FLZ6* from set 3 (which was classified as sugar non-responsive by Jamsheer and Laxmi, (2015) but which was classed as sugar-responsive using the broader set of treatments that we used to calculate CRF values). The above iTPS responses were all assigned to CRF groups $G_2$ or $G_0$, indicating that they are triggered by lower sugar or other indirect effects, rather than by elevated Tre6P itself. At least some of these *FLZ* family members genes were previously reported to be repressed by transient overexpression of SnRK1α1 (*FLZ3, FLZ8* by Jamsheer and Laxmi (2015); *FLZ14* by Baena-Gonzalez et al. (2007), see Supplemental Figure S16)). The implication is that, at least in these cases, SnRK1-signaling is not modulated by Tre6P or, at least, that tre6P is not the major input that regulates SnRK1 activity. Of the remaining four FLZ family members, which are repressed by sugar, three (*FLZ9, FLZ13, FLZ17*) were also significantly repressed in the iTPS response, and these were assigned to CRF group $G_1$. At least two of them (*FLZ9, FLZ17)* were previously reported to be induced by transient overexpression of SnRK1α1 (Jamsheer and Laxmi, 2015). The implication is that SnRK1 induces these *FLZ* family members and that this is counteracted by elevated Tre6P.

Overall, this analysis adds to the evidence that expression of the *FLZ* family is highly regulated by C status and points to Tre6P inhibition of SnRK1 activity contributing to the regulation of a subset that are induced in high sugar, whereas other signaling pathways are involved in the regulation of family members whose expression increases in low C conditions.

#### *Brassinosteroid signaling*

There is emerging evidence for a connection between sugar and BR signaling (Zhang et al., 2016; 2021; Liao et al., 2022). In the dark, TOR signaling stabilizes and promotes accumulation of the brassinosteroid signaling transcription factor BRASSINAZOLE-RESISTANT 1 (BZR1), providing one mechanism by which sugar-signaling can act to balance growth with C availability, in this case to promote hypocotyl extension (Zhang et al., 2016). There is a contrasting response in the light-grown plants, where sugar acts independently of TOR to inhibit BR by stabilizing BRASSINOSTEROID-INSENSITIVE 2 KINASE (BIN2) (Zhang et al., 2021). BIN2 acts as a negative regulator in brassinosteroid signaling, attenuating dephosphorylation of BZR1. Further, as BIN2 directly phosphorylates RAPTOR1B it has been proposed (Liao *et al.*, 2022) that when brassinosteroids are absent BIN2 phosphorylates and inactivates RAPTOR1B leading to activation of the ATG13a-dependent autophagy pathway, and that when BIN2 is inhibited by brassinosteroids RAPTOR1B is less inhibited and TOR activity increases leading to decreased autophagy.

This emerging connection between sugar-signaling and brassinosteroid-sensing and -signaling prompted us to inspect the iTPS response of genes assigned to brassinosteroid synthesis and downstream responses (Supplemental Figure 17).

iTPS modified the transcript abundance of many genes involved in brassinosteroid metabolism and signaling including repression of genes in the biosynthesis pathway (*DWARF4*, *BR6OX2*, *DIMINUT*; all assigned to CRF group $G_2$ so these are probably indirect effects) and signaling components (*BIN2*, *BEE1, BEE2*, all assigned to CRF $G_1$) (Supplemental Figure S17A). A particularly striking response was seen for a subset of brassinosteroid-regulated transcripts highlighted by Zhang *et al.*, 2016) including eight

*EXPANSIN* (*EXPA*) family members and two *XYLOGLUCAN ENDOTRANSGLUCOSYLASE* (*XTH*) family members (Supplemental Figure S17A), which were all repressed and assigned to CRF groups $G_2$ or $G_0$.

### Cell wall modification

The identification of cell wall modifying proteins in the set of brassinosteroid-regulated genes prompted us to inspect the response of the complete *EXPA* and *XTH* families (Supplemental Figure S18). These were already highlighted in the MapMan (Supplemental Figure S7G) and Gene ontology (Supplemental Figure S11, 'plant-type cell wall loosening') enrichment analyses as broadly-responding gene sets in the CRF group $G_2$ response. The detailed analysis in (Supplemental Figure S18) confirms a broad repression of most of the *EXPA* family, with most responses being assigned to CRF groups $G_2$ or $G_0$ (i.e., Tre6P.independent) and a broad repression of *XTH* family members, with some of these being assigned to CRF group $G_1$ and the others to $G_2$ and $G_0$ (i.e., a mix of Tre6P-dependent and Tre6P-independent responses). It is possible that this reflects the need of these different types of cell wall modification for a carbon supply for synthesis of cell wall polymers. EXPAs allow cell wall expansion without incorporation of new cell wall polymers, whilst XTHs allow this and are also required for insertion of newly synthesized hemicellulose polymers into the cell wall.

### Overlap with bZIP11 signaling

$S_1$ and C class bZIP transcription factors play an important role in low energy signaling in plants (Dröge-Laser and Weiste (2018). The C supply regulates expression of the $S_1$ type transcription factor bZIP11 translationally, with high sucrose acting at upstream open reading frames (uORFs) to stall ribosome progression, this block being removed in low C conditions to promote translation (Wiese et al., 2004; Rahmani et al., 2009). A similar translational regulation may also apply for other $S_1$ bZIP proteins (bZIP1, bZIP2, bZIP44, bZIP53) (Juntawong *et al.*, 2014). It is also known that growth is inhibited by overexpression of some $S_1$ class bZIPs including bZIP11 (Hanson et al., 2009; Ma et al., 2011, Dröge-Laser and Weiste, 2018). Further, constitutive overexpression of *bZIP11* in Arabidopsis led to changes in transcript abundance that partly mimic the response to starvation (Ma et al., 2011). Changes included increased expression of genes involved in catabolism, genes involved in the synthesis of minor carbohydrates such as myo-inositol and raffinose, and increased expression of *TPPF* and *TPPG* which are thought to catalyze the breakdown of Tre6P to trehalose.

We therefore compared the response of transcript abundance to *bZIP11* overexpression with that after transient elevation of Tre6P (Supplemental Figure S19).

In their study of in Arabidopsis lines with constitutive overexpression of *bZIP1,* Ma et al. (2011) reported that 232 transcripts showed a significant change and passed a FC filter of $>\log_2 1$ (hereafter termed the oebZIP11 response). We first compared the oebZIP11 response with the carbon response factor (CRF) (Supplemental Figure S19A). There was a broadly negative relationship ($R^2$ = 0.24, negative slope), confirming the conclusion of Ma et al. (2011) that bZIP11 overexpression partly mimics the response to low C. When the oebZIP11 response was compared to the overall iTPS response, there was a weak positive relationship ($R^2$ = 0.14) but many transcripts showed opposing changes or did not change in the iTPS response (Supplemental Figure S19B). Deconvolution of the iTPS response into CRF groups (Supplemental Figure S19C) revealed no relationship between the bZIP11 and iTPS response for transcripts assigned to $G_1$, a strong positive relationship for transcripts assigned to $G_2$, and a weak positive relationship for transcripts assigned to $G_0$. The absence of a relationship for CRF group $G_1$ is consistent with Tre6P-signaling being largely independent of bZIP11-signaling, and the positive correlation for CRF groups $G_2$ is consistent with the decline in sucrose levels in iTPS leading to increased bZIP11-signaling. This might be in part due to lower sucrose leading to depression of the translational arrest. It might be noted that any such effects might mask overlap between the response of genes to bZIP signaling and to elevated Tre6P.

The dataset for constitutive oeTPS (Zhang et al., 2009) contained 94 of the 232 transcripts identified as bZIP11-responsive by Ma *et al.* (2009). We inspected for these transcripts how much agreement there was between their iTPS response and constitutive oeTPS response (Supplemental Figure S19D). There was little agreement for all 94 transcripts, very good qualitative agreement for the subset assigned to iTPS CRF group G1, and a very divergent behavior for the subset assigned to iTPS CRF group G2. This indicates a relatively robust response for transcripts that lie downstream of Tre6P-signaling, but not for transcripts that change due to indirect effects.

Even though the responses to iTPS (or oeTPS) and oebZIP11 were very different, and especially for transcripts assigned to iTPS CRF group $G_1$, many transcripts were shared between the iTPS and oebZIP11 data sets. When the transcripts in the oebZIP11 data set were scored for their iTPS response, (84, 79 and 28 transcripts were assigned to $G_1$, $G_2$, $G_0$, corresponding to 36, 34 and 12% of the oebZIP11 set, respectively. Only 18% of the transcript in the oebZIP11 showed non-significant changes in the iTPS response. The proportion of the total transcripts that showed significant changes in the iTPS response was higher in the subset that showed an oebZIP11 response (82%) than in the total iTPS response (55%, see Supplemental Table S1) and a strikingly high proportion of these (44%) are assigned to CRF groups $G_1$. Further, many of these shared transcripts showed a qualitatively similar response in the constitutive oeTPS data set of Zhang *et al*. (2009). This indicates that although bZIP11- and Tre6P-signaling are independent of each other, they often affect the same genes.

This led us to inspect the response of the iTPS $G_1$ subset more closely (Supplemental Figure S19E). This display separates genes whose transcripts changed in the same and in an opposite direction in the responses to iTPS and oebZIP11. Given that Tre6P acts a positive signal for sucrose, and bZIP11 is translationally activated by low sucrose, it would be expected transcripts would show qualitatively opposite responses to iTPS and in oebZIP11. This was the case for 61 (74%) of the transcripts, indicating that these genes are regulated in a mutually-reinforcing manner by Tre6P-signaling and by sucrose-modulated bZIP11-signaling (Supplemental Figure 19F). Reciprocal responses were also found for the 32 transcripts from this set that were reported by Zhang *et al.* (2009). However, 21 (26%) of the transcripts responded in a qualitatively similar manner in the iTPS and oebZIP11 responses indicating that, for these genes, Tre6P-signaling and bZIP11-signaling may counteract each other Supplemental Figure 19F). Ten of these transcripts were reported in the dataset of Zhang et al .(2009) and 9 of them responded in a qualitatively similar manner to their response to oebZIP11. Genes in this subset included GA-STIMULATED 6 (a cell wall protein downstream of RGL2 that integrates GA, ABA, and glucose signaling, Zhong et al., 2015), the sucrose effluxer SWEET12, a phototropic-responsive NPH3 family member, two SMALL AUXIN UPREGULATED RNAs (SAURs), and the transcription factors JAZ6/TIFY DOMAIN PROTEIN 11B, WRKY17 and NAC20.

We next investigated how Tre6P and SnRK1 interact in the regulation of targets that are shared with oebZIP11 (Supplemental Figure 19G-H). For the 232 genes shortlisted by Ma et al. (2009) as bZIP11 targets, there was no relationship between the response of transcripts to iTPS and to transient overexpression of SnRK1α1. However, after deconvolution into CRF groups, we found a negative relationship for transcripts assigned to CRF group $G_1$, and a positive relationship for transcripts assigned to CRF groups $G_2$ (Supplemental Figure 19G). This mirrors the relationship in the entire iTPS data set (Figure 2, Supplemental Figure S5B). The negative relationship in the subset of bZIP11-regulated transcripts that are assigned to iTPS CRF group $G_1$ is consistent with them being regulated both by bZIP11 and also by Tre6P-inhibition of SnRK1. We also compared the response of transcripts to tSnRK1α1 and oebZIP11 (Supplemental Figure 19H). The vast majority of the transcripts showed qualitatively similar responses to oebZIP11 and tSnRK1α1, both for the total set shared genes (43) as well as for the subsets that were assigned to iTPS CRF group1 (25 genes) and CRF group 2 (14 genes). This again points to their transcript abundance being regulated in parallel by bZIP11-signaling and by SnRK1-signaling.

Finally, Ma et al (2009) highlighted the response of two TPPs (*TPPG* and *TPPF)* to overexpression of bZIP11. We also noticed that trehalase (*TRE*) is also repressed in the oebZIP11 data set. The responses of their transcript ar3e examined in (Supplemental Figure 19J). Trehalase is repressed in the iTPS response and assigned to CRF group $G_1$, is also repressed by constitutive oeTPS, and is induced by oebZIP11, indicating that Tre6P- and bZIP11 act in parallel to repress TRE. A more complicated picture emerged for the two TPPs. These are induced by oebZIP11, but are either weakly induced (but assigned to CRF group $G_0$) or unaffected in iTPS, and both were repressed by constitutive oeTPS. This indicates that constitutive TPS overexpression leads to complex indirect effects on these TPPs.

### Transcription factors

We also inspected the response of transcription factors (TFs), as a subset of genes that might give independent insights into processes that are impacted in the iTPS response.

Elevated Tre6P led to changes in transcript abundance for about 780 TFs spread across all three CRF groups (290 $G_1$, 199 $G_2$, 291 $G_0$) using a relaxed filter of FDR < 0.05, $Log_2FC \geq 0.2$, and about 416 TFs (141 $G_1$, 108 $G_2$, 167 $G_0$) using a stringent filter of FDR < 0.05 and $log_2FC \geq 1$ filters and requiring the filter to be passed in both the 4-h and 6-h datasets. Supplemental Figure S20A lists examples of strongly responding TFs and Supplemental Figure SB-E lists TFs that passed a filter of FDR<0.05 and FC>2 at one or more of the two time points, broken down into TF families and, within each family, the CRF response group.

To provide an overview of the response, we first performed Gene Ontology analysis (Supplemental Figure S21A). We focused on TFs assigned to CRF group $G_1$, which are probably responding to signaling downstream of elevated Tre6P. This analysis highlighted several categories related to carbohydrate metabolism and C-signaling (regulation of carbohydrate utilization, cellular response to glucose stimulus, sugar-mediated signaling process) and biosynthetic pathways (regulation of chlorophyll catabolism, synthesis, regulation of way biosynthesis, anthocyanin-containing compound biosynthetic process) as well as signaling pathways related to light (shade avoidance, far or red light signaling pathway, response to blue light) and hormones (regulation of auxin biosynthetic process, gibberellic acid-mediated signaling process, ethylene-activated signaling pathway, abscisic acid biosynthetic process), the clock (circadian rhythm ) and development (floral meristem determination, phloem or xylem histogenesis). Many of these processes were highlighted in our preceding analyses of the complete iTPS $G_1$ response.

We next used the STRING on-line tool (https://string-db.org/) to search for functional associations between the TFs that respond to transient elevation of Tre6P. The tool utilizes available datasets, published work, automated text mining and computational predictions from various organisms to score the likelihood of association between proteins, and takes into account physical (direct) and functional (indirect) interactions (Szklarczyk et al., 2021). We again focused on transcription factors assigned to CRF group $G_1$, (Supplemental Figure S21B). Within these 290 TFs, there were significantly more interactions than expected by chance (p = 1E-16), Multiple interactions (≥5) are observed for a small number of TFs, indicating that these may be potential central hubs in the iTPS response. The most strongly linked transcription factor was HY5, followed by PIF4, HFR1, DREB26, JAZ6, JAZ8, HAT2, a set of TCPs (includingTCP14, TCP17, TCP5, TCP20 and PTF1/TCP13, APL (ALTERED PHLOEM DEVELOPMENT) and At3g13940) and bZIP1. There was also a small network of NUCLEAR FACTOR Y (NF-YC) family members. Of these, HY5, is a master regulator of thousands of genes and coordinates light, environmental and developmental signaling (Gangappa and Botto, 2016; Dröge-Laser et al., 2018). PIF4, HFR1 are also involved in light signaling, DREB26 in stress responses, JAZ6/MYC3 and JAZ8/MYC4 in jasmonic acid signaling, TCPs in morphogenesis and cell cycle regulation, HATs in regulating auxin-mediated morphogenesis the shoot and root tissues, APL1 in phloem development, NF-YCs in floral regulation and bZIP11 in responses to low energy, whereby PIF4 is also implicated in aspects of carbon signaling. In Supplemental Figure S21B, TFs assigned to selected categories in signaling and metabolism are colored. This again highlights, as expected, a major impact of elevated Tre6P on transcription factors involved responses to C supply, but also on chlorophyll biosynthesis, anthocyanin and glucosinolate biosynthesis, as well as light signaling and hormone signaling (including brassinosteroid and auxin). This recapitulates many of the processes that were highlighted in our analysis of the overall iTPS response.

We next looked in more detail at the C and $S_1$ subfamilies of bZIP TFs. C-bZIPs are translationally regulated by sucrose that represses translation at upstream open reading frames (uORFs) (see above, section ‘Overlap with bZIP11 signaling’) and heterodimerize with group $S_1$ members to control, in a redundant manner, large sets of genes involved in C and energy signaling (Dröge-Laser and Weiste, 2018). They also in interact with SnRK1-signaling. For example, phosphorylation of bZIP63 by SnRK1 alters it dimerization behaviors and triggers a transcriptional low energy response (Mair et al., 2015), and SnRK1-regulated C/S1 bZIP signaling transcriptionally activates the starvation response in prolonged darkness (Pedrotti et al., 2018).

Most C and $S_1$ family members showed significant changes in transcript abundance in the iTPS response (Supplemental Figure S22A), including *bZIP9*, *bZIP25* and *bZIP63* in the C subfamily, and *bZIP1*, *bZIP2*, and *bZIP44* in the $S_1$ subfamily and, of these, *bZIP9*, *bZIP25*, *bZIP63, bZIP1* and *bZIP2* were assigned to CRF groups $G_1$ indicating that they may be repressed by signaling downstream of Tre6P. Given the known role of SnRK1 in the post-translational regulation of these bZIPs (Dröge-Laser and Weiste, 2018), we compared their transcriptional response to iTPS response with the published response to transient SnRK1α1 overexpression (Baena-Gonzalez et al., 2007) (Supplemental Figure S22A). Reciprocal responses were found for *bZIP25*, *bZIP63, bZIP1* transcripts, consistent with the transcriptional repression in iTPS being due to inhibition of SnRK1 by Tre6P. However, qualitatively similar responses were found for *bZIP9* and *bZIP2*, whilst *bZIP10* responded to tSnRK1α1 but not to iTPS, *bZIP11* and *bZIP44* were assigned to the CRF group $G_2$ indicating that they are responding to lower sugar or other indirect effects, and *bZIP53* did not respond to iTPS. Further whilst *bZIP11* was significant repressed at 4h this effect was reverted at 6h and in the constitutive oeTPS data set of Zhang et al. (2009) it was induced. Thus, the transcriptional regulation of bZIP11 may be rather complex with Tre6P-SnRK1 signaling playing only a minor role. It is also noteworthy that *bZIP63* and *bZIP1* were induced by constitutive oeTPS, in contrast to them being repressed by iTPS, again underling the complexity of the signaling networks that regulate their expression and the likelihood that the response to constitutive overexpression of TPS may be overlaid by indirect effects. Overall, transcriptional regulation of the C/S1 bZIPs appears to be rather complicated and it is likely that the translational regulation of by sucrose (see references above) may play a more important role in their response to C than changes in transcription per se.

We next looked more widely at the response of transcripts in the bZIP family. This family consists of many further groups that are implicated in light responses, auxin signaling and transport, pathogen responses and basal resistance (Dröge-Laser et al., 2018; Dröge-Laser and Weiste, 2018). iTPS led to changes in expression of many members of this family, with some being assigned to CRF group $G_1$, and others to $G_2$ or $G_9$ (Supplemental Figure S22B). For each of these sets of genes, we compared the iTPS response with the response to tSnRK1α1. There was a weak negative correlation for genes assigned to $G_1$ ($R^2$ = 0.11 and 0.14 at 4-h and 6-h, respectively) and no relationship for gene assigned to $G_2$ or $G_0$. This is consistent with Tre6P-SnRK1 signaling contributing to the transactional regulation the bZIPs assigned to CRF group $G_1$. The weak correlation might be because Tre6P-SnRK1 signaling makes only a minor contribution for many of the genes, or may reflect limitations in comparing responses in two differing experimental systems. The absence any relation for the genes assigned to $G_2$ and $G_0$ is consistent with their expression being regulated by other indirect effects. We also expanded this analysis to investigate further TF families, focusing on genes assigned to CRF group G1 (Supplemental Figure S23). These was a general trend to many of the genes showing a qualitatively similar response to iTPS and tSnRK1α1, especially for TFs in the AP2/EREBP, C3H Zinc finger, MYB-related and pseud-ARR families.. One notable response was PAP1, a MYB TF that is involved in the sucrose-mediated induction of genes for anthocyanin and flavonoid biosynthesis (Supplemental Figure S20A).

Finally, we next inspected TFs assigned to CRF group $G_2$, which may be responding to lower sugar in the iTPS response. As already seen for all transcripts assigned to $G_2$ (Figure 3, Supplemental Figures S7 and S11), very different sets of TFs were assigned to $G_2$ compared to $G_1$. Enrichment analyses (Supplemental Figure S24A) highlighted, for example, TFs assigned to water homeostasis, water deprivation, response to sulfur starvation, regulation of glucosinolate biosynthetic process, glucosinolate metabolic process, maturity transition, negative regulation of abscisic acid signaling. Some categories were found in both CRF groups $G_2$ and $G_1$, for example, some categories related to light signaling, indicating that not only Tre6P but also other sugar signaling pathways modulate light responses. STRING analysis (Supplemental Figure S24B) highlighted many TFs involved in the response to nitrogen and in the regulation of nitrogen metabolism, sulfur starvation and specialized metabolism including anthocyanin metabolism, flavonoid biosynthesis and glucosinolate biosynthesis.

Supplemental Figure S25 provides the responses of TFs that might contribute to the widespread repression of genes involved in phenylpropanoid and, especially, glucosinolate biosynthesis (see Supplemental Figure S7E, shown for glucosinolate metabolism in more detail in Supplemental Figure S24A) where many genes were repressed and assigned to $G_2$. A set of six upstream MYB family TFs (*MYB29, MYB28, MYB34, MYB51, MYB76, MYB122*) that have been shown to coordinately induce aliphatic or indolic glucosinolate biosynthetic genes (Mitreiter and Gigolashvili, 2021) were also repressed and assigned to CRF group $G_2$, as were OBP1 and WRKY33 which have also been implicated in upstream regulation of glucosinolate biosynthesis (Mitreiter and Gigolashvili, 2021) (Supplemental Figure S24A). This is consistent with the coordinated repression of glucosinolate biosynthesis genes being an indirect effect, possibly due to low sugar repressing the MYB TFs, OBP1 and WTK33. A set of MYC TFs have been implicated in regulating jasmonic acid signaling including its impact on glucosinolate synthesis (Dombrecht et al., 2007; Schweizer *et al*., 2013, Fernandez-Caldo et al 2011). These MYC TFs act independently of the MYB TFs to regulate glucosinolate biosynthesis (Schweizer et al., 2013). Curiously, two of the MYB were repressed and one was induced and these were assigned to CRF group $G_1$, consistent with them being regulated by Tre6P. This points to the complex response of glucosinolate biosynthesis to C availability, with low sugar repressing sulfur metabolism and glucosinolate metabolism, and Tre6P modulating jasmonic acid-induced defense responses.

### References

Publications that are cited in the main text can be found in the reference list of the main text. The following lists publications that are cited only in the Supplemental text.

**Abe M, KobayashiY, Yamamoto S, Daimon Y, Araki T.** FD, a bZIP protein mediating signals from the floral pathway integrator FT at the shoot apex: Science 2005:**309**:1052–1056, doi: 10.1126/science.1115983

**Abelanda JA, Bergonzi S, Oortwijn M, Sonnewald S, Du M, Visser RG, Sonnewald S, Bachem CWB.** Source-sink regulation is mediated by interaction of an FT homolog with a SWEET protein in potato. Current Biol 2019:**29**:1178-1186

**Andrés F, Kinoshita A, Kalluri N, Fernández V, Falavigna VS, Cruz TMD, Jang S, Chiba Y, Seo M, Mettler-Altmann T, Huettel B, Coupland G.** The sugar transporter SWEET10 acts downstream of *FLOWERING LOCUS T* during floral transition of *Arabidopsis thaliana*. BMC Plant Biol 2020:**20**:1-14

**Brightbill CM, Sung S.** Temperature-mediated regulation of flowering time in *Arabidopsis thaliana*. aBIOTECH 2022:**3**:78–84

**Chardon F, Bedu M, Calenge F, Klemens PAW, Spinner L, Clement G, Chietera G, Léran S, Ferra M, Lacombe B, et al.** Leaf fructose content is controlled by the vacuolar transporter SWEET17 in *Arabidopsis*. Current Biol 2013:**23**:697-792

**Flis A, Mengin V, Ivakov AA, Mugford ST, Hubberten HM, Encke B, Krohn N, Höhne M, Feil R, Hoefgen R, et al.** Multiple circadian clock outputs regulate diel turnover of carbon and nitrogen reserves. Plant Cell Environ 2019:**42**:549–573

**Fontaine J-X, Tercé-Laforgue T, Armengaud P, Clément G, Renou J-P, Pelletier S Catterou M, Azzopardi M, Gibon Y, Lea PJ et al.** Characterization of a NADH-Dependent Glutamate Dehydrogenase Mutant of Arabidopsis Demonstrates the Key Role of this Enzyme in Root Carbon and Nitrogen Metabolism. Plant Cell 2012:**24**(10):4044-4065

**Griffiths J, Murase K, Rieu I, Zentella R, Zhang Z-L, Powers SJ, Gong F, Phillips AL, Hedden P, Sun T-P, Thomas SG.** Genetic characterization and functional analysis of the GID1 gibberellin receptors in Arabidopsis. Plant Cell 2007:**18**:3399–3414

**Guo WJ, Nagy R, Chen G H-Y, Pfruder S, Yu Y-C, Santiella D, D Frommer W, Martinoia E.** SWEET17, a facilitative transporter, mediates fructose transport across the tonoplast of Arabidopsis roots and leaves. Plant Physiol 2013:**164**:777-789

**Izawa T.** (What is going on with the hormonal control of flowering in plants? Plant J. 2021:**105**:431–445

**Jin S, Ahn JH.** Regulation of flowering time by ambient temperature: repressing the repressors and activating the activators. New Phytol 2021:**230**:938–942.

**Klemens PAW, Patzke K, Deitmer J, Spinner L, Le Hir R, Bellini C, Bedu M, hardon F, Krapp A, Neuhaus HE.** Overexpression of the vacuolar sugar carrier AtSWEET16 modifies germination, growth, and stress tolerance in Arabidopsis. Plant Physiol 2013:**163:**1338-1352

**Li S, Tian Y, Wu K, Ye Y, Yu J, Zhang, Liu Q, Hu3, Li H, Tong Y, Harberd NP, Fu X.** Modulating plant growth–metabolism coordination for sustainable agriculture. Nature 2018:**560**(7720):595-600 doi.org/10.1038/s41586-018-0415-

**Liu L, Liu Chang, Hou X, Wanyan Xi, Shen L, Tao Z, Wang Y, Yu H.** FTIP1 is an essential regulator required for florigen transport. PLoS Biol 2012:**10**:e1001313

**Quiroz S, Yusti JCs, Chávez-Hernández EC, Martínez T, de la Paz Sanchez M, Arroyo AG, Álvarez-Buylla E-R, García-Ponce B** Beyond the Genetic Pathways, Flowering Regulation Complexity in Arabidopsis thaliana. J. Mol. Sci.2021:**22**,5716

**Shim JS, Kubota A, Imaizumi T.** Circadian Clock and Photoperiodic Flowering in Arabidopsis: CONSTANS Is a Hub for Signal Integration. Plant Physiol 2017:**173**:5-15

**Takehishi I, Okada K, Fukazawa J, Yohsuke Takahashi Y.** DELLA-dependent and -independent gibberellin signaling. Plant Signal Behav 2021:**13**: e1445933.

**Turck F, Fornara F, Coupland G.** Regulation and identity of florigen: FLOWERING LOCUS T moves center stage. Annu Rev Plant Biol 2008:**59**:573-594.

**Wang JW.** Regulation of flowering time by the miR156-mediated age pathway. J Exp Bot 2014:**65**: 4723-4730

**Yin S, Ahn JH**. Regulation of flowering time by ambient temperature: repressing the repressors and activating the activators. New Phytol. 2021:30:938-942. DOI10.1111/nph.17217

**Zhong C, Y Xu H, Ye S, Wang S, Li L, Zhang S, Wang X.** Gibberellic Acid-Stimulated Arabidopsis6 serves as an integrator of gibberellin, abscisic acid, and glucose signaling during seed germination in Arabidopsis. Plant Physiol 2015:**169**:2288-2303
